# Anchorage-independent cell proliferation promoted by fascin’s F-actin bundling

**DOI:** 10.1101/2024.05.04.592404

**Authors:** Tadamoto Isogai, Vasanth Siruvallur Murali, Felix Zhou, Xinxin Wang, Divya Rajendran, Lizbeth Perez-Castro, Niranjan Venkateswaran, Maralice Conacci-Sorrell, Gaudenz Danuser

**Affiliations:** Lyda Hill Department of Bioinformatics, UT Southwestern Medical Center, Dallas, TX, USA; Cecil H. and Ida Green Center for Systems Biology, UT Southwestern Medical Center, Dallas, TX, USA; Department of Cell Biology, UT Southwestern Medical Center, Dallas, TX, USA

**Keywords:** actin, anchorage-independent, proliferation, cancer, signaling

## Abstract

The actin filament (F-actin) bundling protein fascin-1 is highly enriched in many metastatic cancers. Fascin’s contribution to metastasis have been ascribed to its enhancement of cell migration and invasion. However, mouse genetic studies clearly point to functions also in tumorigenesis, yet without mechanistic underpinnings. Here, we show that fascin expression promotes the formation of a non-canonical signaling complex that enables anchorage-independent proliferation. This complex shares similarities to focal adhesions and we refer to them as pseudo-adhesion signaling scaffolds (PASS). PASS are enriched with tyrosine phosphorylated proteins and require fascin’s F-actin-bundling activity for its assembly. PASS serve as hubs for the Rac1/PAK/JNK proliferation signaling axis, driven by PASS-associated Rac-specific GEFs. Experimental disruption of either fascin or RacGEF function abrogates sustained proliferation of aggressive cancers *in vitro* and *in vivo*. These results add a new molecular element to the growing arsenal of metabolic and oncogenic signaling programs regulated by the cytoskeleton architecture.

## Introduction

The expression of actin filament (F-actin) bundling protein fascin-1 (fascin) is directly correlated with increased risk of metastasis and poor clinical outcome ^1–7^. In normal physiology, fascin expression is high during early embryogenesis and largely absent from healthy epithelial tissue ^8^. Mice lacking fascin are viable but have melanoblast cell migration defects resulting in a ‘white-spot’ phenotype ^9^. When overexpressed in cancer cells, fascin is found enriched in filopodia ^10^, invadopodia ^11^ and focal adhesions ^12,13^, and shown to stimulate cancer cell migration and cell invasion through its actin-bundling activity ^4,5,7,14–18^. These observations have widely established fascin as a pro-migration factor and on this ground, fascin expression has been asserted to promote cancer cell metastasis *in vivo* ^1–5^. Under this premise, a series of fascin actin-bundling inhibitors have been developed with the goal of blocking metastatic spread by blocking migration ^14–16^. However, the data accompanying more recent evaluations of fascin inhibitors clearly indicate effects on the growth of primary breast cancer xenografts as well ^15,16^. This observation was later confirmed in specific subtypes of solid tumors ^19^. Fascin’s contribution to tumor growth has been underappreciated, perhaps because fascin perturbation in a variety of cancer cell lines showed robust effects on migration but mixed effects on proliferation. Surveyed in a meta-study by Liu et al. ^7^, 96% of studies (54/56) reported a positive correlation between fascin expression and enhanced cell migration or invasion, whereas only 70% (26/37) report a correlation between fascin expression and proliferation (Table S1). Notably, most of these studies relied on cell culture on 2D plastic substrates and metabolic activity-based assays to determine fascin’s association with proliferation ^20,21^. Given that fascin was shown to stimulate mitochondrial respiration ^22^, these results may be confounded by secondary and indirect effects ^23^.

In contrast, studies employing genetically engineered cancer mouse models (GEMMs) for pancreatic ductal adenocarcinoma and colorectal carcinoma indicated a clear role of fascin primary tumor growth ^24,25^; perplexingly histological analyses in both studies led to the conclusion that neither proliferation nor survival were modulated by the presence or absence of fascin. Thus, despite its remarkable effects, fascin’s tumorigenic role has remained unexplained.

The following results resolve the conundrum surrounding fascin’s function in tumor growth by elucidating how its F-actin bundling activity facilitates the assembly of signaling scaffolds that activate proliferation pathways independently of cell anchorage. While signals originating from these scaffolds are nonessential in cells firmly attached to substrates as encountered in cell culture *in vitro*, they gain importance in environments with weaker mechanical support.

## Results

### Fascin’s penetrance as a proliferation factor depends on the environment

To investigate fascin’s potential roles in cancer cell proliferation and survival, we first examined whether genetic depletion of fascin by CRISPR/Cas9 in a human colorectal adenocarcinoma cell line (HCT116) xenograft would reproduce the outcomes in GEMMs ^24,25^. Fascin-1 is the predominant fascin expressed in HCT116 (Fig. S1a) and knockout of fascin (*FSCN1*) did not alter the expression of fascin-2 (*FSCN2*) and fascin-3 (*FSCN3*) (Fig. S1b). When subcutaneously injected into immunodeficient (NOD/SCID) mice, wild-type (WT) and fascin knockout (KO) cells formed tumors, however, WT tumors grew faster than KO tumors (Fig. 1a), which was confirmed with an independent colon adenocarcinoma cell line (DLD-1). Hence, these cell lines provided an appropriate model systems to explore the contribution of fascin in cancer cell proliferation and survival. Strikingly, in agreement with previous reports ^7^, we found that under standard 2D conditions, fascin KO cell cultures grew similarly to their wild-type counterpart (Fig. 1b). We conjectured that the discrepancy in growth between *in vivo* xenografts and *in vitro* cell culture may originate in the embedding of cells in a much softer and more complex environment compared to placement on a flat and infinitely stiff 2D substrate. To test this, we seeded wild-type and fascin KO cells on top of a pre-polymerized collagen gel (2.5D) and monitored their growth. Cell counts over time indicated that fascin KO cell cultures had a growth disadvantage when compared to the wild-type cell cultures, akin to the situation in tumor xenografts (Fig. 1c). We confirmed this result with an independent KO using a different targeting sgRNA (Fig. S1c). The difference in cell count between wild-type and fascin KO cells could be attributed to a decrease in cell proliferation or survival under fascin depletion. An imaging-based proliferation assay ^26^ showed that fascin was dispensable for proliferation in 2D (Fig.1d) but not in 2.5D (Fig.1e). Fascin KO cells proliferated even less when fully embedded in 3D (Fig.1f). Importantly, the number of dead cells did not differ between wild-type and fascin KO cells embedded in 3D (Fig.1g). These results indicate that fascin expression promotes proliferation in an environment-dependent manner. Since our 2D assay relied on collagen coating of the glass substrate, the differences in proliferation between 2D, 2.5D and 3D are likely related to differences in ECM geometry and/or stiffness rather than ECM chemistry.

**Fig. 1:**
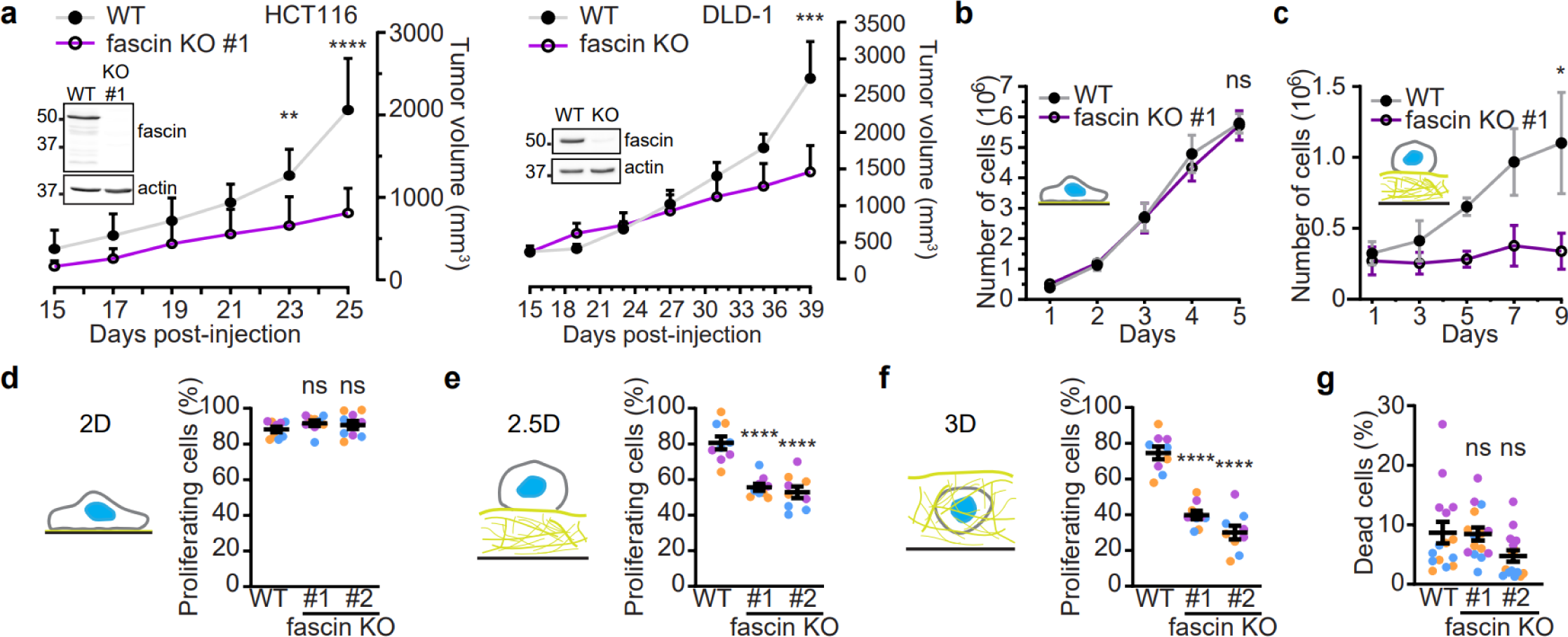
Fascin is a context-dependent proliferation factor. **a,** Fascin expression promotes tumor growth *in vivo*. Volume of tumors derived from subcutaneously injected wild-type (WT) and fascin knockout (KO) HCT116 and DLD-1 cells were followed over time. *n* = 5, 6, 8 & 9 for HCT116 WT, KO, DLD-1 WT, KO, respectively. **b,** Fascin expression is dispensable for cell growth in 2D. Cell number was counted at indicated time. *n* = 18 for each cell line. **c,** Loss of fascin attenuates cell growth in 2.5D. Cells seeded on top of a pre-polymerized collagen gel were detached at indicated time to count cell numbers. *n* = 16 for each cell line. **d,** Fascin expression is dispensable for proliferation in 2D. Cells were incubated with Edu for 18 hours and proliferating cells were determined using an imaging-based method ^26^. Proliferating cells (%) were determined by the fraction of Edu-positive cells among all cells detected by DAPI counterstain. *n* **=** 9 per cell line. **e,f,** Loss of fascin attenuates cell proliferation in 2.5D and 3D. Cells were either seeded on top of pre-polymerized collagen gel (**e**) or fully embedded in collagen gel (**f**) and proliferating cells as determined in **e.** *n* **=** 9 per cell line and per condition. **g,** Cell survival is not affected upon fascin KO. Cells embedded in 3D collagen gel were stained live for membrane impermeable Ethidium Homodimer (EthD-1) and percentage of EthD-1-positive dead cells over total number of cells were determined by imaging. *n* **=** 15 per cell line. Scale bars, 100 µm. For **a**, 2way ANOVA, for **b-c**, paired two-tailed Student’s t-tests, for **d-g**, unpaired two-tailed Student’s t-tests were performed as statistical analysis. ns, *P* > 0.05; \**P* < 0.05, \*\**P* < 0.01, \*\*\**P* < 0.001, \*\*\*\**P* < 0.0001. Data are mean ± s.e.m. (**a, d-g**) or mean ± s.d. (**b** and **c**). Each color represents a biological replicate.

### Fascin nucleates the formation of anchorage-independent signaling scaffolds

Considering the sensitivity of proliferation rates to changes in the ECM, we focused on fascin’s association with focal adhesions (FAs) ^12,13^. FAs are multicomponent signaling complexes that bridge the extracellular matrix and the actin cytoskeleton and are involved in cell proliferation and survival ^27,28^. The density and subcellular distribution of FAs are strongly affected by the ECM, especially when cells are cultured in soft 3D collagen gels ^29^. Therefore, we hypothesized that fascin expression may be an important factor in sustaining FA-mediated proliferation signaling in soft 2.5D and 3D environments. Indeed, by imaging the canonical FA component paxillin we found that wild-type cells formed significantly denser punctate structures in 3D collagen gels than the fascin KO cells (Fig. 2a; Supplemental Movie 1-2). We developed software to quantify the density of mNeonGreen-paxillin puncta at the cell surface (Fig. 2b). Co-staining of cells with antibody recognizing phospho-Tyrosine (pY) residues in proteins displayed a similar punctate pattern at the cell surface, whose density was reduced upon fascin KO. (Figs. 2c,d; Supplemental Movies 3-6). Hence, we concluded that fascin expression maintains a high-density pool of FA-like adhesions that are signaling competent even in a soft 3D environment with low mechanical engagement by the ECM.

**Fig. 2:**
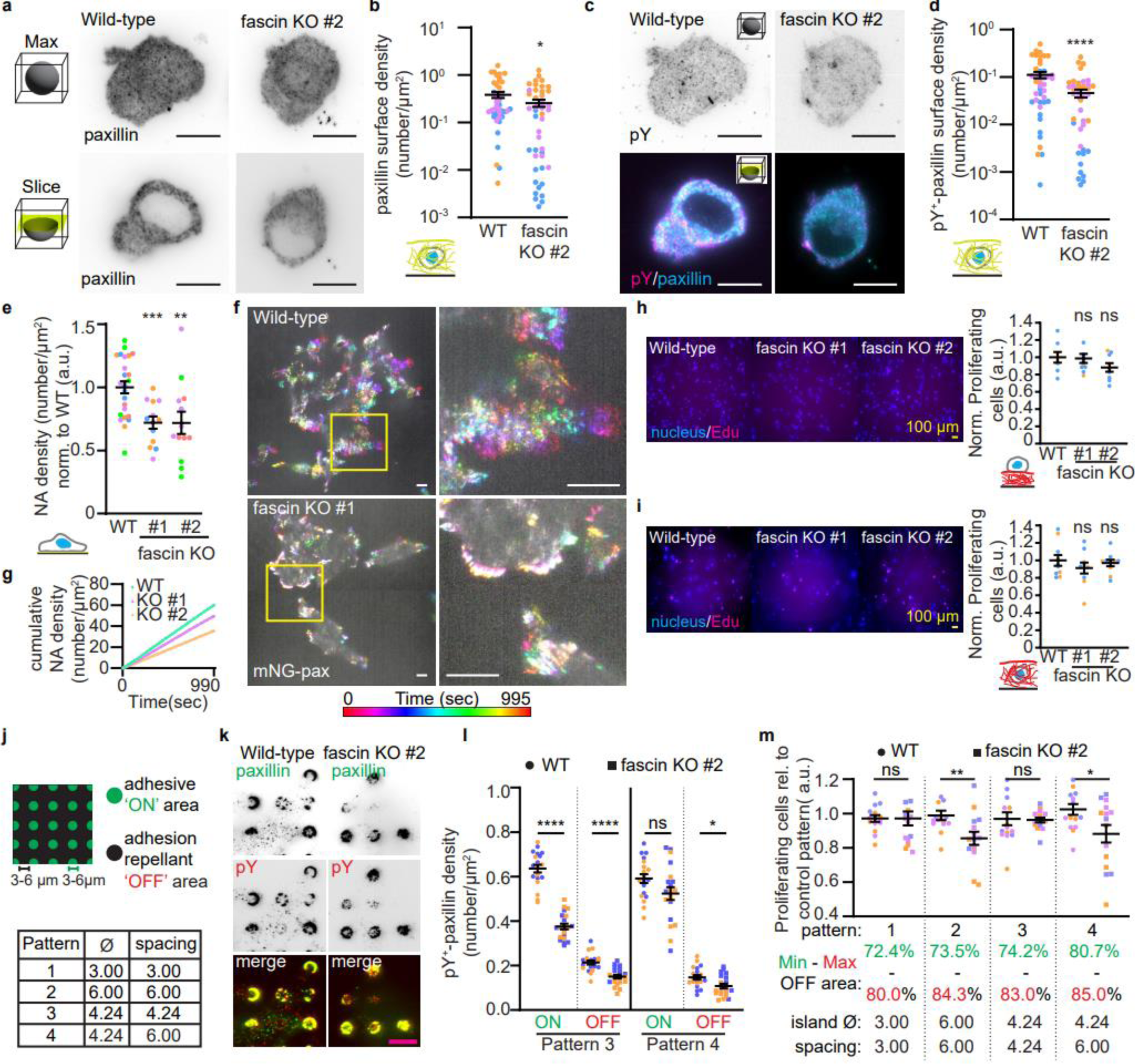
Fascin nucleates focal adhesions not bound to the ECM. **a**, Fascin expression stimulates the formation of nascent adhesion (NA)-like structures in 3D. Cells embedded in 3D collagen gels were stained with anti-paxillin antibodies and imaged using LSFM. Maximal intensity projections or a single slice in the stack are shown **b**, Fascin KO decreases surface paxillin density of cells in 3D. Paxillin speckles at the cell surface of cells from **a** were detected and quantified as number per area (density). *n* = 45 per cell line. **c**,**d**, Fascin expression stimulates the formation of Tyrosine phosphorylated signaling complexes. Cells from **a** were co-stained with anti-phospho-Tyrosine (pY) antibodies. pY-positive (pY^+^) paxillin density at the surface are plotted in **d**. *n* = 45 per cell line. **e**, Fascin KO cells have defects in NA dynamics. NAs were detected as described in Method and normalized per cell area. Mean NA density over time per live cell movie are plotted. *n* = 25 for WT, 13 for both KOs. **f**, Fascin expression nucleates transient NAs. Live-cell images of mNeonGreen-paxillin expressing wild-type and fascin KO cells were projected over time with the indicated color LUT. See also Figs. S2b,c for quantification of NA dynamics. **g**, Fascin expression stimulate nucleation of NA. Cumulative NA density was obtained from e and the mean of all cells is plotted over time. **h,i** Representative images and normalized proliferation quantification of cells seeded on 2.5D (h) and 3D (**i**) collagen gels polymerized with telocollagen (see also Fig. 1e,f for additional details). Scale bar, 100 µm. **j**, Micropattern design. Collagen islands are represented in green, whereas adhesion-repellant area depicted in black. **k,l**, Fascin expressing cells contain more paxillin, pY and pY^+^-paxillin NA-like speckles independent of ECM engagement. Cells seeded on pattern 4 were stained for paxillin and pY. On and OFF pattern pY^+^-paxillin density is plotted in **l**. *n* = 20 per cell line. **m**, Fascin KO cells proliferate less by restricting adhesion area. Cells seeded on micropatterns overnight and pulsed for 1 hour with Edu. Percentage proliferating cells were determined as in Fig. 1d. n = 15 per condition. Scale bars, 10 µm. For b,d, one-sided Wilcoxon Rank Sum test, for e,l,m, unpaired two-tailed Student’s t-tests were performed as statistical analysis. ns, *P* > 0.05; \**P* < 0.05, \*\**P* < 0.01, \*\*\**P* < 0.001, \*\*\*\**P* < 0.0001. Data are mean ± s.e.m.. Each color represents a biological replicate.

Based on these results and given that fascin was dispensable for proliferation in 2D (Fig. 1d), we expected that in 2D the density of mNeonGreen-paxillin puncta would be the same in the presence or absence of fascin. To be consistent with the measurements in 3D, we only considered nascent state FAs, commonly referred to as nascent adhesions (NAs), that appear as diffraction-limited punctate structures when imaged using Total Internal Reflection Fluorescence microscopy (TIRF). Contrary to our initial conjecture, fascin KO cells displayed a decrease in NA density also on collagen-coated glass (Fig. 2e; Fig. S2a). These results seemed to suggest that the marked difference in proliferation between fascin wild-type and KO cells in 3D collagen gels may not be related to the difference in NA formation between these cells. However, while in 3D the adhesion-like structures were homogeneously punctate, in 2D the NAs were accompanied by extended FAs that were not affected by fascin expression (Fig. 2f). Comparison of fascin wild-type and KO cells by live-cell TIRF imaging indicated that fascin depletion abrogated the pool of dynamic NAs that rapidly nucleated and disassembled (Fig. 2f; Fig. S2b,c; Movies S7-12). Previous work indicated that about half of NAs do not exert traction on the substrate, suggesting that they are not mechanically engaged with the ECM ^30^. Based on this observation and considering that FA formation is more restricted in soft environments, we hypothesized that a significant fraction of fascin-dependent, paxillin-containing structures in wild-type cells in 2.5D and 3D environments may exist in absence of anchorage to the ECM. Upon fascin depletion these structures disappear and, with them, the signals that promote proliferation. On 2D substrates, these anchorage-independent NAs are complemented by assembly of mechanically engaged NAs and FAs, regardless of the level of fascin expression. The proliferation signal produced by these canonical adhesion structures is sufficient to make up for the loss of signal associated with fascin depletion.

To formally test this model, we pursued a two-pronged approach: First, as a corollary to the fascin-independence of proliferation on 2D substrates, we expected that a stiffening of the collagen gel would rescue the formation of canonical adhesions in fascin KO cells, thus promoting fascin-independent proliferation. Conversely, in 2D, fascin KO cells would also display reduced proliferation when the density of ECM-engaged canonical adhesions is limited. To test the first hypothesis, we seeded cells in 2.5D and 3D gels polymerized from non-pepsinized collagen, which are markedly denser and slightly stiffer than gels made with pepsinized collagen ^31,32^. Our results confirmed that non-pepsinized collagen gels of the same concentrations as the pepsinized collagen gels in Figs. 1e and 1f rescued proliferation of fascin KO cells to wild-type levels (Fig. 2h,i). To test the second hypothesis, we used micro-patterned substrates with collagen-coated adhesive islands in a background of oxidation-resistant adhesion repellant polymers that block cell attachment up to 14 days. We designed four patterns with varying island diameter and island spacing (Fig. 2j, Fig. S2d,e). For both wild-type and fascin KO cells we did not observe a notable difference in cell spreading (Fig. S2f). We then used paxillin and pY staining to study the ECM-engaged (ON) and anchorage-independent (OFF) NA distribution (Fig. S2f). Due to technical limitations of antibody cross-reactivity, we were not able to image the micropatterns simultaneously with the paxillin and pY stains. However, the approximate location of the micropatterns was readily visible from both paxillin and pY staining as cells tended to favor attachment to the collagen islands. Therefore, we decided to computationally overlay the micropattern area on the paxillin and pY images using independently obtained micropattern images as a guide (Fig. S2e,g,h). Fascin KO cells consistently showed decreased paxillin, pY and pY-positive paxillin puncta on both ON and OFF patterns (Fig.2j,k; Fig. S2g-m). As the adhesion-repellant area increased, restricting ECM-engaged FAs, fascin KO cells showed an attenuated proliferation, whereas wild-type cells continued to proliferate independently of micropattern configurations (Fig. 2m). This observation was independently validated using lung adenocarcinoma cell lines that were depleted of fascin using two independent sgRNAs (Fig. S2n,o). Altogether, our results demonstrate that fascin nucleates the formation of a non-canonical type of anchorage-independent NAs that promote cancer cell proliferation in highly compliant environments nonpermissive of mechanical engagement of adhesions. We termed these anchorage-independent, proliferation-stimulating structures as pseudo-adhesion signaling scaffolds (PASS).

### Actin filament bundles support proliferation in suspension

To further validate that PASS are formed independent of anchorage to the ECM, we imaged cells cultured in suspension. These experiments were challenged by cellular drift and a PASS size typically below the diffraction-limit of light microscopy. To overcome these technical hurdles, we embedded cells in abiotic and inert hydrogels that simulate suspension culture conditions while still holding cells in place for imaging extremely weak signals. Paxillin and phospho-tyrosine imaging confirmed that wild-type cells had significantly elevated PASS density compared to fascin KO cells (Figs. 3a, b; Fig. S3a). Proliferation analysis of cells sparsely cultured in suspension confirmed that fascin KO cells proliferated less compared to the wild-type cells (Fig. 3c; Fig. S3b), while cell survival rates remained similar (Fig. 3d). These results were reproduced in lung adenocarcinoma cells (Fig. S3c-e). Conversely, re-expression of EGFP-tagged fascin in the KO cells rescued proliferation in suspension (Fig. S3f,g). Small differences in the surface active β1 integrin levels relative to total β1 integrin were observed albeit randomly depending on the KO clone. This suggested that our observations were not triggered by changes in integrin activation (Fig. S3h). Moreover, chemical inhibition of myosin activity did not affect proliferation in suspension, suggesting that the phenotype is contraction-independent (Fig. S3i). This is consistent with the notion that proliferation differences between wild-type and fascin KO cells are unrelated to mechanical interactions promoted by anchorage to the substrate and/or force generation.

**Fig. 3:**
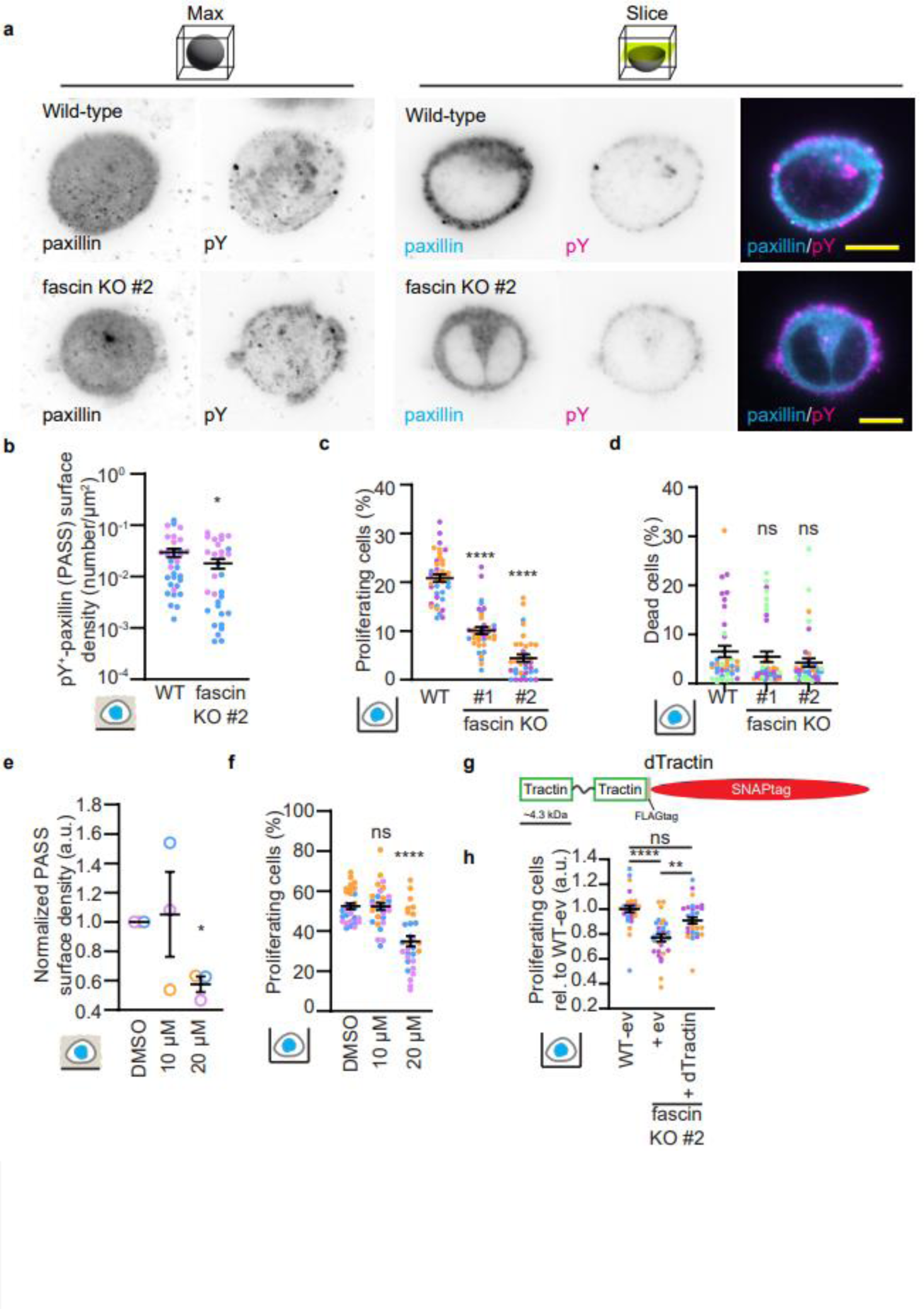
PASS requires F-actin bundles to promote proliferation. **a**, Fascin expression stimulates the formation of PASS in suspension. Wild-type and fascin KO cells embedded in hydrogels were stained with anti-paxillin and anti-pY antibodies and imaged using LSFM. Maximal intensity projections or a single slice of the stacks are shown. **b**, Fascin expression stimulates the formation of PASS. pY-positive (pY^+^) paxillin (PASS) density at the surface are plotted. *n* = 35 for WT and 36 for KO. **c**, Fascin promotes proliferation in suspension. Cells were cultured in suspension for 18 hours and pulsed with 1 hour Edu. Percentage proliferating cells were determined as in Fig. 1d. n = 40 for WT, 37 for KO #1 and 36 for KO #2. **d**, Cell survival in suspension is independent of fascin expression. Cells cultured in suspension overnight were incubated with EthD-1 and percentage of EthD-1-positive dead cells over total number of cells was determined by imaging. n = 40 per cell line. **e**, F-actin bundling activity is required for PASS formation. Cells were embedded in hydrogels and incubated overnight with the indicated concentrations of NP-G2-044. PASS was quantified as in **b** and normalized to DMSO control per repeat. n = 3. **f**, F-actin bundling activity of fascin stimulates proliferation in suspension. In suspension cultures were incubated overnight with the indicated concentration of NP-G2-044 and pulsed with 1 hour Edu. Percentage proliferating cells determined as in Fig. 1d. n = 30 per condition. **g**, Graphical representation of dTractin. Drawn to scale. Scale bar is about 4.3 kDa. **h**, dTractin expression rescues proliferation defects of fascin in suspension. Cells cultured overnight in suspension were pulsed for 1 hour with Edu; percentage proliferating cells was determined as in Fig. 1d and normalized to WT-ev control. N = 29-30. Scale bars, 10 µm. For **b**, one-sided Wilcoxon Rank Sum test, for **c,d,e,** unpaired two-tailed Student’s t-tests, for **f**, paired two-tailed Student’s t-test, for **h**, One-Way ANOVA were performed as statistical analysis. ns, *P* > 0.05; \**P* < 0.05, \*\**P* < 0.01, \*\*\*\**P* < 0.0001. Data are mean ± s.e.m.. Each color represents a biological replicate.

Since fascin crosslinks F-actin into bundles, we investigated whether this bundling activity was necessary for both PASS formation and proliferation in suspension. Indeed, chemical inhibition of fascin’s F-actin bundling activity by NP-G2-044 attenuated both PASS density and proliferation in suspension (Figs. 3e,f). NP-G2-044 did not induce cell death (Fig.S3j) and application of this reagent to fascin KO cells had minimal to no further attenuation of proliferation in suspension (Fig. S3k). Together, these data show that NP-G2-044 blocks fascin’s F-actin bundling with minimal off-target effects and that it is this function of fascin that is responsible for PASS assembly and anchorage-independent proliferation.

Motivated by these findings, we wondered whether induction of F-actin bundles alone would be sufficient to promote proliferation. To investigate this, we designed a minimal bundling construct using a tandem of the 44 amino acid F-actin binding peptide, Tractin, derived from the rat *Itpka* gene ^33^ separated by an 18 amino acid flexible linker. This choice was made to avoid confounding effects triggered by potential signaling elements present in other F-actin crosslinkers. Intriguingly, overexpression of the construct was sufficient to restore proliferation of fascin KO cells to similar levels as wild-type cells (Fig.3i). These data led us to conclude that F-actin bundles enhance proliferation under low-adhesion conditions.

### PASS is a Rac signaling hub

Our experiments thus far show that the formation of PASS in response to F-actin bundling provides cells with a mechanism for proliferation signaling independent of ECM anchorage. To investigate the proteins involved in PASS-based signaling, we conducted an unbiased proteomic screen of the PASS-proximal interactome (proxisome) in cells cultured in suspension using paxillin fused to a small and fast promiscuous biotin ligase as a bait ^34^. To validate the suitability of paxillin as a bait, we investigated paxillin’s contribution to cell proliferation in both 2D and suspension. Our results showed that knocking out paxillin had small effects on proliferation in 2D but led to more pronounced proliferation defects in suspension, akin to fascin KOs (Fig. S4a-c). Our proteomic analysis confirmed that fascin is among the significantly enriched components in the proxisome. Moreover, we found numerous FA-related proteins including GITs and several ARHGEFs, specifically RacGEFs that target Rac GTPases for activation (Fig. 4a; Table S2). Rac1 activity has been associated with the regulation of proliferation ^35^. Peculiarly, independent gene interruption of fascin or Rac1 in mice not only led to the same ‘white spotting’ phenotype due to melanocyte migration defects, they both also showed cell cycle defects ^9,36^. Based on this finding we conjectured that PASS may be a center of sustained Rac1 activation specifically in low-adhesion environments. This led us to two hypotheses: First, perturbation of RacGEFs should specifically attenuate proliferation in low-adhesion conditions but not on 2D substrates akin to the penetration of fascin perturbation; second, Rac1 activity levels in fascin KO cells should be decreased when compared to the wild-type cells in low-adhesion conditions, but not on 2D substrates.

**Fig. 4:**
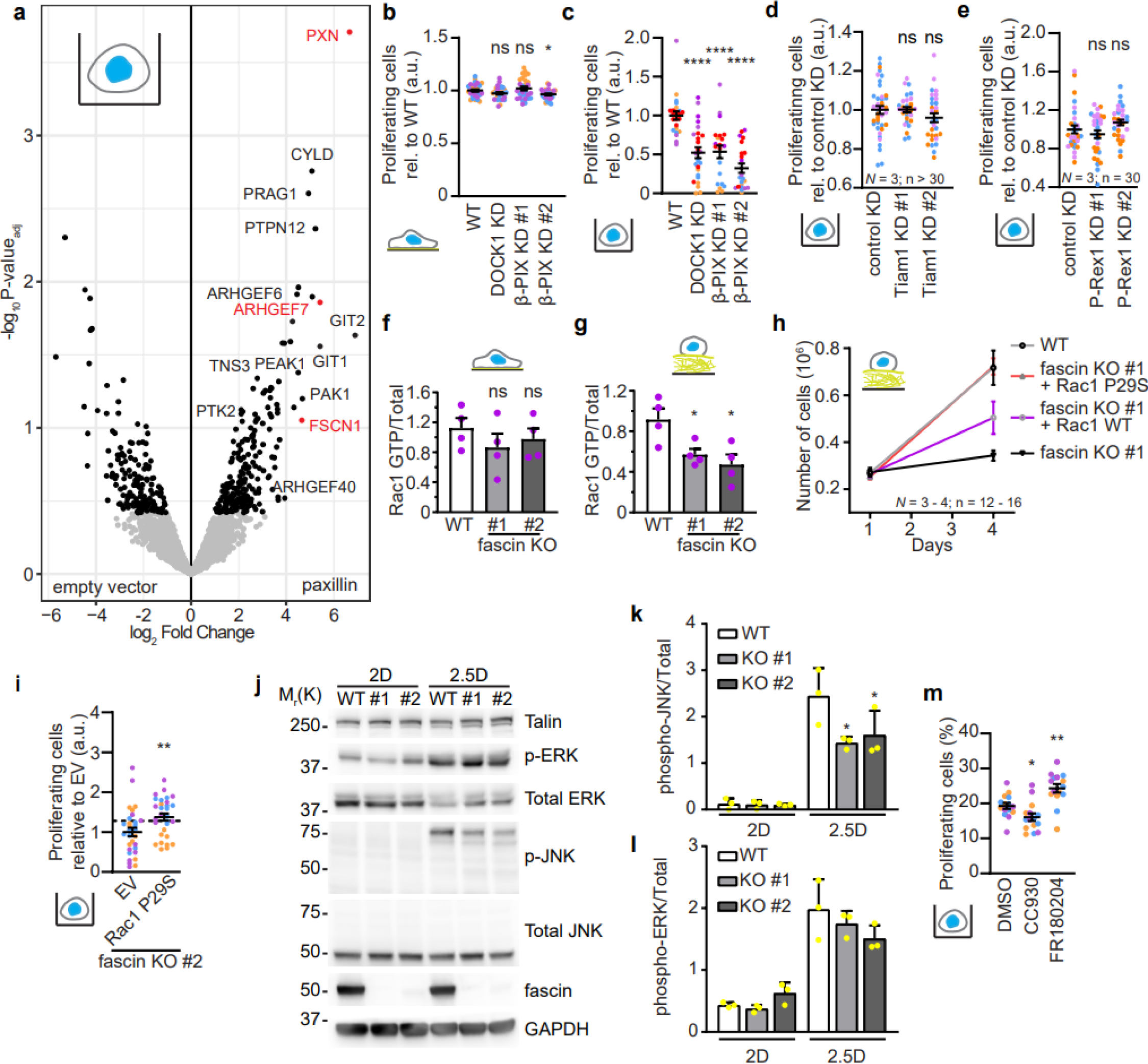
PASS is a signaling hub for Rac1. **a,** PASS proxisome identifies several canonical FA-associated proteins and RacGEFs **b,** DOCK1 and β-PIX expression is dispensable for proliferation in 2D. Percentage proliferating cells were determined as in Fig. 1d and normalized to wild-type controls. *n* = 30 for each cell line. **c,** DOCK1 and β-PIX KD attenuates proliferation in suspension. *n* = 21-25 per cell line. **d,e** Tiam1 (**d**) and P-Rex1 (**e**) expression is dispensable for proliferation in suspension. *n* = 30-40 for each cell line **f,g,** Fascin depletion does not affect Rac1 GTP levels in 2D (**f**) but decreases Rac1 GTP levels in 2.5D (**g**). Rac1 pull-downed by GST-PAK beads (Rac1 GTP) were normalized to total Rac1 levels (Total). *n* = 4 for each cell line and condition. **h,** Exogenous Rac1 expression rescues cell growth defect of fascin KO cells in low-adhesion environments. Cells were seeded and counted as in Fig. 1c. *n* = 12-16 per condition. i, Exogenous Rac1 expression rescues proliferation defect of fascin KO cells in suspension. Percentage proliferating cells were normalized to EV control. Dotted line indicates the normalized mean proliferating cells of wild-type control. *n* = 30 for each cell line. **j**, Fascin expression attenuates JNK phosphorylation in 2.5D but not in 2D. Cells cultured in 2D and 2.5D were lysed and subjected for the indicated antibodies for western blot analysis. Quantifications are presented in **k** and **l**. *n* = 3. **m,** JNK inhibition but not ERK inhibition attenuates proliferation in suspension. Proliferation of cells cultured in suspension were cultured in the presence of 10 µM CC930 (JNK inhibitor), 10 µM FR180204 (ERK inhibitor), or DMSO. *n* = 15 for each condition. For **b-i** and **k-m**, unpaired two-tailed Student’s t-tests, were performed as statistical analysis. ns, *P* > 0.05; \**P* < 0.05, \*\**P* < 0.01, \*\*\**P* < 0.001, \*\*\*\**P* < 0.0001. Data are mean ± s.e.m.. Each color represents a biological replicate except for **f,g,k,l** where each point represents a biological replicate.

To test the first hypothesis, we knocked down the highest PASS proxisome ARHGEF hit, ARHGEF7/β-PIX. β-PIX knockdown (KD) was dispensable for proliferation in 2D but attenuated proliferation in low-adhesion environments (Figs. 4b-c; Fig. S4d,e). Among the additionally tested GEFs, the KD of DOCK1 but not that of Tiam1 or P-Rex1 attenuated proliferation in suspension (Figs. 4b,d,e; Fig. S4d,f,g). These results suggests that previously FA-annotated β-PIX and DOCK1 ^37^ are PASS-associated RacGEFs, and Rac1 activity in PASS rather than global Rac1 activation sustains proliferation in low-adhesion environments.

To test the second hypothesis, we determined the Rac1 GTP levels of wild-type and fascin KO cell lines cultured in 2D, 2.5D or in suspension by pull-down assays. Western blot analysis showed that fascin KO cells had significantly attenuated Rac1 GTP levels compared to wild-type cells in 2.5D as well as in suspension but not in 2D (Figs. 4f,g; Fig. S4h,i). Importantly, modest overexpression of wild-type Rac1 (Rac1 WT) and the hyper-activatable mutant Rac1^P29S^ (Rac1 P29S; Fig. S4j) rescued population growth of fascin KO cells on collagen gels (Fig. 4h) and proliferation in suspension (Fig. 4i) to similar levels to that of wild-type cells. Of note, Rac1 overexpression in the wild-type cells in 2D and 2.5D environments did not have any aberrant effects on cell population growth (Fig. S4k,l). Collectively, our results show that Rac1 activation in PASS via the PASS-associated RacGEFs DOCK1 and β-PIX promotes cell proliferation specifically in low-adhesion environments.

Rac1 has been shown to stimulate JNK activity through PAK/MEK ^38^. Fascin KO did not alter the phosphorylation levels of JNK in 2D, but in 2.5D fascin KO cells showed a significant decrease in phosphorylated JNK (Fig. 4j,k). To further control the specificity of these culture-conditional effects of fascin KO, we confirmed that the same bands representing phosphorylated JNK disappeared upon chemical inhibition of Rac or JNK in wild-type cells cultured in 2.5D (Fig. S4m). ERK activity was not altered in any of the conditions tested (Fig. 4l). Additionally, JNK inhibition but not ERK inhibition attenuated proliferation in suspension (Fig. 4m). Thus, our data suggest that PASS signals activate JNK to promote proliferation in low-adhesion environments independent of ERK activity.

### Inhibition of F-actin bundling regresses MYC/fascin-induced hepatocellular carcinoma

To address fascin’s function in F-actin bundling as a driver of cancer proliferation *in vivo*, we leveraged a mouse model of hepatocellular carcinoma (HCC) under the control of the proto-oncogene MYC ^39^. Ectopic MYC expression has been shown to increase fascin expression in colon cancer cells including HCT116 and DLD1 ^40^. Indeed, we reproduced this MYC-driven elevated fascin expression in HCC by conditional induction of the MYC transgene in hepatocytes (Fig. S5a,b), causing aggressive tumor growth with high penetrance. Due to the rapid nature of tumor progression, this model minimally implicates tumor cell migration and distant metastasis, allowing us to disentangle fascin’s function in invasion from the proposed function in proliferation. Activating MYC at birth leads to the development of tumors in 100% of the mice. Mice carrying MYC-induced liver cancer were treated with the F-actin bundling activity inhibitor NP-G2-044 starting from day 35 post-MYC induction, 5 days a week by gavage, 11 total doses until the mice were sacrificed at 51 days post-MYC induction (Fig. 5a). The body weights of untreated mice were significantly heavier than those of the treated cohort (Fig. 5b), solely due to an increase in liver weight (Fig. 5c,d). Untreated livers developed HCC as previously described ^39^ and macroscopically it was apparent that the increased liver weight was due to tumorigenic tissue development (Fig. 5e). In contrast, livers from mice treated with NP-G2-044 showed almost complete tumor regression with most livers looking normal (Fig. 5e). NP-G2-044 treatment did not alter fascin levels in these tumors (Fig. S5c). Immunohistochemical analysis confirmed a significant decrease in tumorigenic lesions upon inhibition of F-actin bundling via fascin inhibition (Fig. 5f; Fig. S5d) and reduced proliferation (Fig. 5g,h; Fig. S5e). Some treated tumors had heterogenous response and mixed proliferative areas (Fig. S5f presenting green and red dots of incomplete proliferation suppression by fascin treatment), indicating that NP-G2-044 treatment did not fully regress HCC. Tumorigenic lesions were usually accompanied by high rates of apoptosis (Fig. S5g), but NP-G2-044 treatment did not alter cell death (Fig. S5h). Thus, inhibition of fascin’s F-actin bundling activity regressed MYC-induced HCC tumor development by reducing tumor cell proliferation.

**Fig. 5:**
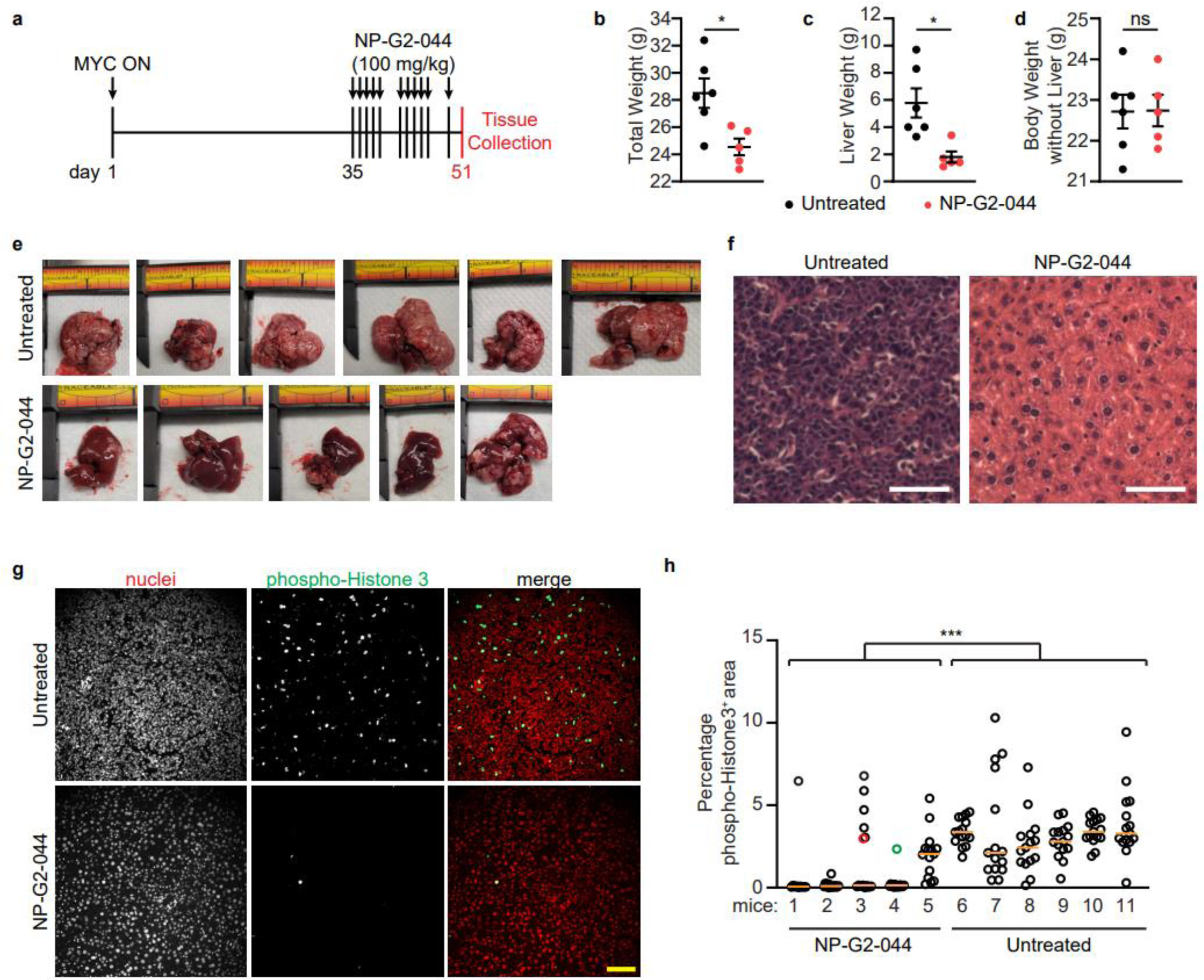
Inhibition of F-actin bundling regresses MYC-induced liver tumors. **a**, Treatment diagram of MYC-induced Hepatocellular Carcinoma mouse model with the fascin activity inhibitor NP-G2-044. MYC expression was induced on day 1 after removal of doxycycline containing water. From day 35, NP-G2-044 (100 mg/kg) was administered by gavage on the indicated days. Mice were sacrificed at day 51. **b,** Body weights of treated and untreated mice were measured at day 51. **c,** Isolated liver weights of treated and untreated mice were measured at day 51. **d,** The body weight without the liver were calculated by subtracting the total body weight from **b** with the liver weight from **c**. For **b,c,d**: *n* = 6 for untreated and 5 for treated. **e**, Images of all the isolated livers from treated and untreated mice. **f,** H&E stains of untreated and NP-G2-044-treated liver tissues. **g,** NP-G2-044 treatment attenuates proliferation *in vivo.* Tissue slices were stained for phospho-Histone 3 (pH3) antibodies and nuclei. **h**, Percentage of pH3-positive cells per tissue area was quantified. Images of red and green dots are presented in Extended Data Fig. 5c. Orange line represents median. n > 10 per mice. Scale bars, 100 µm. For **b-d**, unpaired two-tailed Student’s t-tests, for **h**, 2-way ANOVA was performed as statistical analysis. ns, *P* > 0.05; \**P* < 0.05, \*\*\**P* < 0.001. Data are mean ± s.e.m.

## Discussion

Since its discovery in 1978 ^41^, fascin has been widely studied both in the context of cytoskeleton and adhesion regulation ^10,12,13,42^ and in the context of cancer ^24,25,43^. These studies were motivated by observations of fascin as a pro-migration/pro-invasion factor *in vitro* ^4,10,11,14,16–18,40^ and the striking correlation between fascin gene expression and metastasis formation *in vivo* ^1–3^. Studies utilizing GEMMs have suggested a role for fascin also in tumorigenesis ^24,25^, but the underlying mechanism and contribution of fascin remained a mystery as both studies did not detect a difference in proliferation or survival in the presence or absence of fascin. Adding to this conundrum, *in vitro* studies have shown inconclusive results when defining the role of fascin in cell proliferation ^7^. The present study now solves the puzzle.

Acquisition of anchorage-independent growth is one of the early hallmarks of neoplasia ^44^. Integrin receptors lie at the basis of regulating anchorage-dependent growth by transducing chemical signals through the assembly of focal adhesions including tyrosine kinase adaptors ^27,28^. These integrin-driven signals modulate tumorigenesis in response to composition and stiffness of the extracellular environment ^45,46^. Cancer cells gain growth signaling that bypass the need of integrins through mutations in proto-oncogenes and tumor suppressor genes. The discovery of PASS prescribes an alternative strategy for cells to organize their intracellular signaling landscape to gain proliferative advantage in the absence of anchorage cues. We propose that the inconclusiveness of *in vitro* studies regarding fascin’s role as a proliferation factor stems from the fact that this function fully penetrates only in soft 3D environments and/or in suspension. Our study benefits from several technical advances in 3D imaging and micro-patterning of substrates that were not leveraged in previous work.

We envision that the formation of PASS provides proliferation signals at multiple stages of cancer progression. Our data directly demonstrates a role of PASS in liver tumor development, in line with the GEMM studies ^24,25^. However, in contrast to these studies we could document a direct increase in the proliferation rate of fascin expressing and thus PASS containing tumors over the rates in tumors treated with fascin inhibitors. A plausible explanation for this difference is that mature solid tumors acquire fibrotic behavior, which diminishes the need for anchorage-independent proliferation signals. The decrease of fascin- and PASS-dependence of proliferation with cells cultured in stiff as opposed to soft collagen (Fig. 2h,i) offers an *in vitro* demonstration of this conjecture. Thus, while previous studies ^24,25^ did document the advantages fascin expression generates for colorectal tumor growth, their histological analyses of proliferation rates were likely conducted at a stage in which tumor fibrosis negated the role of fascin as promotor of proliferation signals. Our discovery is also distinct from previous associations of fascin with proliferation via its secondary effects on metabolism ^20–23^. As our direct manipulations of PASS and downstream signals demonstrate, fascin’s primary contribution to tumor cell proliferation ensues from its function as a signaling organizer.

Fascin’s ability to support anchorage-independent proliferation very likely contributes also to critical stages of metastasis including proliferation during circulation and initial proliferation during colonization of metastatic niches. How much fascin promotes metastatic spreading of cancers as a proliferation factor *versus* as an enhancer of cell migration and invasion remains to be determined in future studies. Regardless, given the pro-invasive and pro-proliferative advantages fascin provides, it is not surprising that fascin levels have been found elevated in so many metastatic cancer types ^1–3,6,7^.

More recently, fascin in the nucleus has also been described to promote cancer cell survival ^47^, highlighting the versatility of this 55 kDa protein. Although our *in vitro* and *in vivo* data did not ascribe a major role in cell survival to fascin, the entirety of published observations vis-à-vis the structural simplicity of this protein suggests that the functional versatility stems from the emerging roles of F-actin bundles as signaling scaffolds. Previous work in our lab showed that cancer cells use F-actin bundles also to maintain high glycolysis in soft environments ^48^. Others have shown how F-actin bundles in the nucleus mediate efficient DNA damage response ^49^. Our current findings add an important new element to this repertoire: Fascin-induced F-actin bundles stimulate the formation of PASS as a potent proliferation signaling platform. Our observation that fascin’s function in PASS formation and cell proliferation could be replaced by a synthetic tandem F-actin binding peptide reinforces the idea that the organization of F-actin structures is a major factor in the regulation of cell signaling. This perspective significantly expands the potential ways in which cancer cells can exploit the actin cytoskeleton to acutely modulate cellular behaviors during cancer disease progression ^50^.

## Methods

### Chemicals and Reagents

High-glucose Dulbecco’s modified Eagle’s medium (DMEM) supplemented with pyruvate and stable glutamine was purchased from Invitrogen; NP-G2-044 (fascin inhibitor; CAS No. 1807454-59-6) from MedChemExpress; FR180204 (ERK inhibitor used at 10 µM) and CC-930 (pan-JNK inhibitor used at 10 µM) from SelleckChem. NSC23766 (Rac inhibitor used at 50 µM) was purchased from Tocris and has been characterized to primarily target RacGEFs Tiam1 and Trio ^51^, of which the latter was described to localize to focal adhesions ^37^. Bovine Collagen I solutions PureCol and TeloCol were purchased from Advanced Biomatrix; Halt™ Protease and Phosphatase inhibitor EDTA-free cocktail from Thermo.

### Expression Vectors

The following plasmid backbones were ordered from Addgene: pSpCas9(BB)-2A-GFP (pX458; #48138); pCW57.1 (#41393), pGEXTK-Pak1 70-117 (#12217). pEGFP-C1 and pLVX-IRES-Puro were purchased from ClonTech. Guide RNA sequences were cloned into pSpCas9(BB)-2A-GFP (pX458). Human fascin cDNA was PCR amplified and subcloned into pEGFP-C1 using BamHI/XhoI and compatible BglII/SalI sites in the vector. EGFP-fascin was subcloned into pCW57.1. pLVX-iPuro-Rac1 wild-type and hyperactive Rac1^P29S^ mutant, and pLVXCMV100-mNeonGreen-22-paxillin were previously described ^52^. miniTurbo ^34^ was N-terminally fused to human paxillin cDNA with a 22 amino acid linker and cloned into pLVX-IRES-Puro by HiFi assembly. All new constructs were sequence verified. Primer sequences are listed in Table S3.

### Cell lines and Cell culture

HCT116, DLD-1 and HEK293T cells were purchased from American Type Culture Collection. HCC4087 cell line was a kind gift from Dr. John Minna. All cell lines were regularly tested negative for mycoplasma contamination using MycoScope PCR Mycoplasma Detection Kit (Genlantis).

HCT116, DLD1 and HEK293T cells were cultured in DMEM supplemented with 10% fetal bovine serum (FBS). 293T cells were transfected with polyethylenimine (PEI) with a 3:1 PEI/DNA ratio. Lentiviral particles were produced by VSV envelope vector psPAX2 (7 µg), pMD2.g viral packaging construct (5 µg) and lentiviral expression plasmids (5 µg). Viral media was harvested, filtered (0.45 µm), and used to infect cells in the presence of 2 µg/ml polybrene. Infected cells were enriched either through antibiotic selection or fluorescence-activated cell sorting (FACS).

Gene knockout cell lines were generated by transiently transfecting cells with gene-specific pX458 together with a modified self-cleaving donor vector ^53^ to deliver a blasticidin S resistant cassette into the genomic cut site. Cells were selected with 5 µg/ml blasticidin S and surviving colonies were isolated using 6 mm Pyrex® cloning cylinder (Sigma-Aldrich). Gene knockout was verified by western blotting of whole cell lysates with appropriate antibodies.

### Cell Growth, Proliferation and Survival Assays

To determine cell number over time, for 2D planar surfaces 250,000 cells were seeded on 2 µg/ml bovine collagen I coated 6 wells plate. For soft collagen gels, 250 µl 2 mg/ml bovine collagen I was polymerized in 24 wells plate and 250,000 cells were seeded in duplicates. For each time point, cells were trypsinized from tissue culture plates or harvested from collagen gels by disassociating the gel with 500 µl of PBS containing 1 U/ml collagenase type I (Gibco). Cells were washed once with PBS, and the pellet was re-suspended in 200 µl PBS. 20 µl of the reconstituted cell suspension was mixed with 20 µl of Trypan Blue (Sigma) and the live cell count was determined using Cellometer Auto 1000 Bright Field Cell Counter (Nexcelom). Samples were counted in duplicates.

For EdU proliferation assays on 2D planar surfaces (2D), 250,000 cells were seeded in a 6 wells plate with 2 µg/ml collagen coated #1.5 glass coverslips and allowed to spread overnight. For EdU proliferation assays on soft collagen gels (2.5D), 250 µl 2 mg/ml bovine collagen I was polymerized in a 24 wells plate and 300,000 cells were plated. For EdU proliferation assays with cells fully embedded in collagen gels (3D), 100,000 cells were resuspended with 2 mg/ml bovine collagen I gel and allowed to polymerize in 24 wells plate. Proliferation of cells cultured in suspension (suspension) was determined by culturing 300,000 cells on Ultra-Low Attachment 6 well plate (Corning) in 3 ml of media. In all cases, cells were cultured for 18 hours in the incubator, followed by incubation with 5-ethynyl-2’-deoxyuridine (EdU; 10 µM), mostly for 1 hour unless otherwise indicated. After a brief wash with PBS, samples were fixed for 10 (2D), 15 (suspension), and 30 minutes (2.5D and 3D) with 4% paraformaldehyde at 37 °C. The 2D, 2.5D and 3D samples were processed as previously described ^54^. Briefly, cells were permeabilized with 0.1% Triton X-100 for 10 minutes at room temperature, and subsequently incubated for 30 minutes with a home-made Click-iT labeling mix (8 µM Sulfo-Cy3-Azide or Sulfo-Cy5-Azide, 2 mM CuSO_4_, 20 mg/ml ascorbic acid in PBS). The suspension samples were processed using the Click-iT Plus Edu Alexa Fluor 488 Flow Cytometry Assay Kit (Thermo; C10419) according to manufacturer’s instructions. In all cases, cell nuclei were labeled with 300 nM 4’,6-diamidino-2-phenylindole (DAPI). Edu (proliferative) and DAPI (total) stained cells were imaged using a Nikon Ti-Eclipse epifluorescence microscope and images were analyzed using cell profiler as previously described ^54,55^. For survival assays, cells cultured in 2D, 2.5D, 3D or in suspension as described above were incubated with cell-impermeable 4 µM ethidium homodimer 1 (EthD-1; Invitrogen catalogue no. E1169) and 10 µg/ml cell-permeable Hoechst (Thermo Fisher catalogue no. H3570) for 30 min before imaging. Dead cells were identified by positive EthD-1 staining.

### Protein Purification and Active Rac1 Pull-Down Assays

CRIB pulldown assays were carried out by incubating 400-500 µg of total cell lysates with 30 µg of GST-CRIB as previously described ^56^. Total and active Rac1 was determined by western blotting.

### Live-Cell TIRF Microscopy (TIRFM)

Cells were seeded and allowed to spread for 36 hours on 2 µg/ml collagen-coated 35 mm glass bottom dishes. Images were acquired on a Ti-Eclipse inverted microscope equipped with Perfect Focus System (Nikon), a 60x 1.49 NA Apo TIRF objective (Nikon) with an additional 1.8x tube lens, yielding a total 108x magnification, and a sCMOS camera with 60 nm pixel size (pco.edge). TIRF illumination was achieved using a Diskovery Platform (Andor Technology). TIRF time-lapse images were acquired at a penetration depth of 80 nm and every 5 seconds. Imaging was performed at 37°C, 5% carbon dioxide, and 70% humidity.

### Preparation of µPatterned Surfaces

Custom-designed micropatterns (Fig. S2d,e; MP-S-02-C) were produced by 4DCell, Montreuil, France, to create adhesive and adhesion-repellant surfaces on a 25 mm #1 coverslip. Oxidation resistance anti-adhesive polymer served to repel cell attachment to the coverslip for prolonged duration (14+ days).

Cells were seeded and allowed to spread for at least 18 hours on 2 µg/ml collagen-coated *µ*Patterned glass coverslips. Cells were imaged with TIRF microscopy, as described above.

### Detection and Quantification of Nascent Adhesions in 2D

Nascent adhesions (NAs) were detected and segmented from the images using a previously described software ^30^. For live cell imaging, diffraction-limited objects from cells expressing low levels of mNeonGreen-paxillin under a truncated promoter were detected using the point source detection used in single particle tracking ^57^. In brief, images were filtered using a Laplacian of Gaussian filter, local maxima were detected and fitted with an isotropic Gaussian function (standard deviation: 2.1 pixel). Outliers were removed using a goodness of fit test (*p* = 0.05). NA density is the number of NA divided by the cell area. All densities were normalized to the corresponding wild-type control per experimental day. For fixed samples, diffraction-limited paxillin and phospho-Tyrosine (pY) puncta were detected as described above. pY-positive paxillin were determined by optimally pairing paxillin and pY foci to minimize the total Euclidean distance using bipartite matching ^58^. Subsequently, any matched pairs with a distance larger than 1 µm in Euclidean distance were removed from the subsequent analyses.

For cells seeded on micropatterns, the NAs were detected as described above. Cells were manually segmented. The micropatterns were detected as follows: We first determined the radius (r) and the distance (d) between nearest neighbors of a given pattern by utilizing the Matlab-internal circle detection tool *imfindcircles*. Next, focal adhesions (FAs) were segmented using a previously described analysis tool ^30^. These segmentations discriminated noise from signal belonging to FAs and were used as a binary mask to match and refine the alignment of the micropatterns with the cell images. Specifically, patterns were rotated and translationally shifted to maximize the overlap between the pattern and segmented adhesions. We also added a manual adjustment step to fine-tune the position of the patterns, if needed. We then used the point sources of detected NAs within a segmented cell to determine NA counts ‘ON’ (ECM-engaged) and ‘OFF’ (anchorage-independent) the adhesive pattern. The NAs ON and OFF pattern were normalized to the respective areas within the field of view to determine ON and OFF pattern NA densities. An image typically contained 2-5 cells in the field-of-observation, and each dot in the plots Figs. 2l,m and Figs. S2i-o represents the measurement of one image.

### Embedding into Hydrogels

400,000 cells (45 µl) were embedded in xeno-free hydrogel (VitroGel 3D from The Well Biosciences; TWG001) at a 1:4 cell suspension to undiluted VitroGel (180 µl) ratio. The mixture was incubated at 37 °C for 30 minutes to allow gel formation and incubated overnight in media in a humidified incubator (37 °C and 5% CO_2_). The following day, gels were fixed and stained with antibodies prior to imaging.

### High-Resolution Adhesion Imaging in 3D

For 3D imaging, a home-built high-resolution version of the Axially Swept Light-Sheet Microscope (ASLM) was used ^59^. In brief, a 25x NA 1.1 water-immersion objective (CFI75 Apo LWD 25XW, Nikon Instruments) and a 28.6x NA 0.71 water-immersion objective lens (54-10-7, Special Optics) were used for detection and light-sheet generation, respectively. This yielded a resolution of approximately 450 nm and voxel size length of 104 nm. The microscope was controlled by the custom LabVIEW software (Coleman Technologies, National Instruments).

### Detection and Quantification of Adhesions in 3D

Cells were cropped and volumes resampled to isotropic voxel resolution. pY and paxillin channels were co-registered using rigid body registration with Mattes’ mutual information metric ^60^ and 16 intensity bins using the Python SimpleITK library. Individual foci in pY and paxillin channels were detected as Gaussian-like blobs of radii 5 voxels (= 0.52μm) with minimal separation of 5 voxels and of a minimum integrated brightness using the trackpy.locate function from the Python trackpy library ^61^. This also detects spot-like entities outside the cell. In the absence of a membrane marker, we constructed an approximate binary mask of the cell volume by Gaussian smoothing of the Otsu thresholded binary of the paxillin channel following local contrast enhancement and Wiener deconvolution. The mask was then applied to exclude out of cell foci. To assess colocalization, paxillin and pY foci were optimally paired to minimize the total Euclidean distance using bipartite matching ^58^. Only pairs separated by a distance less than or equal to 10 voxels (=1.04μm) were retained for analysis. To compute the cell surface area, the binary volume was triangularly meshed using Marching cubes. The surface area was then computed as the sum of the individual triangle face areas.

### Proximity biotin labeling screen

HCT116 wild-type and fascin KO #2 cells expressing paxillin fused to a small promiscuous biotin ligase were seeded on 100mm ultra-low attachment petri dishes (Corning; CLS3262) for 24 hours before beginning the experiment. Cells were incubated with 50 µM biotin for 1 hour and lysed in JS buffer (50 mM Tris HCl pH7.5, 150 mM NaCl, 5 mM EGTA, 1.5 mM MgCl2, 1% Glycerol and 1% Triton X-100) supplemented with protease and phosphatase inhibitors (Halt Protease and Phosphatase Inhibitor, Thermo). Cleared lysates were incubated with Protein A Sepharose beads to remove proteins with nonspecific binding to the beads. The resulting supernatant (500 µg) was incubated with 100 µl of streptavidin-coated magnetic beads (Dynabeads MyOne Streptavidin T1 beads, Invitrogen) overnight at 4 °C with rotation to enrich for biotinylated proteins. Beads were washed three times with ice-cold JS buffer and in-gel digested for Mass Spectrometry analysis. Two independent experiments were performed.

Proteomics data was analyzed using the DEP package for differential enrichment analysis of proteomics data in R ^62^. In brief, abundance data was background-corrected and normalized by variance stabilizing transformation, and any remaining missing values were imputed as draws from a Gaussian distribution centered on quantile 0.005 of the observed values with standard deviation equal to the median standard deviation of the observed values. Subsequently, differential enrichment was estimated based on protein-wise linear models and empirical Bayes statistics.

### Mouse experiments

500.000 HCT116 and DLD-1 cells resuspended in 100 µl of PBS were injected subcutaneously into the dorsal flank of 6–8-week-old immunocompromised NOD/SCID female mice (purchased for Jackson laboratory). Tumor size was measured twice a week using calipers, and the mice were euthanized before tumors reach 2 cm^3^ in volume, according to animal protocol approvals. The MYC-driven liver tumor model was generated by crossing TRE-MYC with LAP-tTA mice. The liver-specific transactivator Tg mouse LAP-tTA drives TRE-MYC expression in the absence of Dox (Tet-Off system). The TRE-MYC mice were generated by D.W. Felsher ^39^ and LAP-tTA by H. Bujard ^63^. Both mice were obtained from Dr. Hao Zhu (UTSW). Male breeders with 2 copies of LAP-tTA and a single copy of TRE-MYC were crossed with wild-type FVB females to generate LAP-tTA/TRE-MYC mice used for experiments. The breeders were maintained on doxycycline water (1 mg/ml). The females with the litters were transferred to a fresh cage on the day of birth with doxycycline-free water to activate MYC overexpression. Because human MYC transgene randomly integrated in the Y chromosome in this model, only male offspring develop liver cancer upon MYC induction as described previously ^39^. Animals were maintained on a regular chow diet. NP-G2-044 was dissolved in 10% DMSO and 90% corn oil and administered to mice at 100 mg/kg by gavage. Treatment with NP-G2-044 was initiated on at P35 and treated in total for 11 days (administered only on weekdays and not on weekends). Body weight of mice was measured on day 51 before harvest of liver tissues. Subsequently, the liver tissues were isolated and weighted. All animal experiments were performed under the approved animal protocol: APN2017-101798 (UTSW).

### Immunohistochemistry

Following overnight fixation in 10% neutral-buffered formalin, subgross tumor samples were paraffin processed, embedded, and sectioned by members of UT Southwestern’s Histo Pathology Core according to standard procedures ^64,65^. Paraffin sections were concomitantly prepared and checked by dark-field microscopy ^66^. Resulting serial sections were stained by hematoxylin & eosin (H&E), Terminal deoxynucleotidyl transferase-mediated UTP End Labeling (TUNEL; DeadEnd Fluorometric TUNEL System; Promega, G3250), and Ki-67 (Leica Biosystems, # KI67-MM1-L-CE) or phospho-histone3 (phosphoH3; Millipore, # 06-570-AF488) antigen immunohistochemistry (IHC). Briefly, sections were deparaffinized and Ki-67 or phosphoH3 antigens were retrieved by heating in pH 6.0 citrate-buffer or in citraconic-anhydride, respectively. Sections were blocked with commercially available blocking reagents for Ki67 (Vector Mouse on Mouse “MOM” Kit, Vector Laboratories, # BMK-2202) or with 0.5% BSA for phosphoH3. Sections were incubated with primary antibodies (Ki67 at 1:300; phosphoH3 at 1:250) overnight at 4°C. The following day, Ki67 positive cells were detected by biotin/streptavidin-fluorescein incubation according to the MOM kit instructions. Finally, nuclei were counterstained with propidium iodide and mounted on glass slides using Vectashield (Vector Laboratories, # H1000). phophoH3-positive (pH3^+^) and apoptotic cell nuclei from tissue images were segmented using Pixel Classification in ilastik ^67^.

### Total RNA Isolation and qPCR Analyses

Total RNA was isolated using RNeasy mini kit (Qiagen) and complementary DNA (cDNA) was transcribed by SuperScript™ IV Reverse Transcriptase PCR (Invitrogen) according to manufacturer’s guidelines. Quantitative real-time PCR (qRT-PCR) reactions were set up using 5–10 ng of cDNA as a template and gene-specific primers (200 nM) in a CFX96 Touch™ Real-Time PCR Detection System (Bio-Rad). All reactions produced single amplicons, which allowed us to equate one threshold cycle difference. Results were normalized with respect to EF1α expression. mRNA levels were quantified according to the 2−ΔΔCt method (Ct, cycle threshold). Primer sequences are listed in Table S3.

### Statistical Analysis

Statistical analysis and plotting of data were performed using GraphPad Prism 9.0, as indicated in the figure legends. Counts and the meaning of *n* is stated in the figure legend. In all cases: \**P* < 0.05; \*\**P* < 0.01; \*\*\**P* < 0.001; \*\*\*\**P* < 0.0001.

## Supporting information

Supplemental Figures with Legends

## Acknowledgements

We thank Drs. Dana Reed, Kushal Bhatt and Mr. Joseph Chi (UT Southwestern Medical Center) for logistical support. Computational resources were provided by the BioHPC facility located in the Lyda Hill Department of Bioinformatics. The Danuser lab software repositories are maintained by Dr. Qiongjing Zou. Funding in the Danuser lab is provided by the grant R35GM136428. Funding in the Conacci-Sorrell lab is provided by grants Cancer Prevention and Research Institute of Texas (CPRIT) RP220046, American Cancer Society 724003, Welch foundation I-2058-20210327 and NCI R01CA245548. The content of the manuscript is solely the responsibility of the authors and does not represent the official views of the funding agencies.

## Author Contributions

Conceptualization, T.I. and G.D; methodology, T.I., V.S.M., F.Z., X.W., D.R., L.P.-C. and N.V.; formal analysis, T.I., V.S.M., F.Z. and X.W.; validation, T.I., V.S.M..; resources, T.I., M.C.-S. and G.D.; investigation, T.I. and G.D.; supervision, G.D.; funding acquisition, M.C.-S. and G.D.; writing—original draft, T.I., and G.D.

## Declaration of interests

T.I and G.D. have filed a patent to target F-actin bundles to restrict the growth of tumors.

## Notes

### Competing Interest Statement

The authors have declared no competing interest.

