## Supplemental Figures with Legends for "Anchorage-independent cell proliferation promoted by fascin’s F-actin bundling"

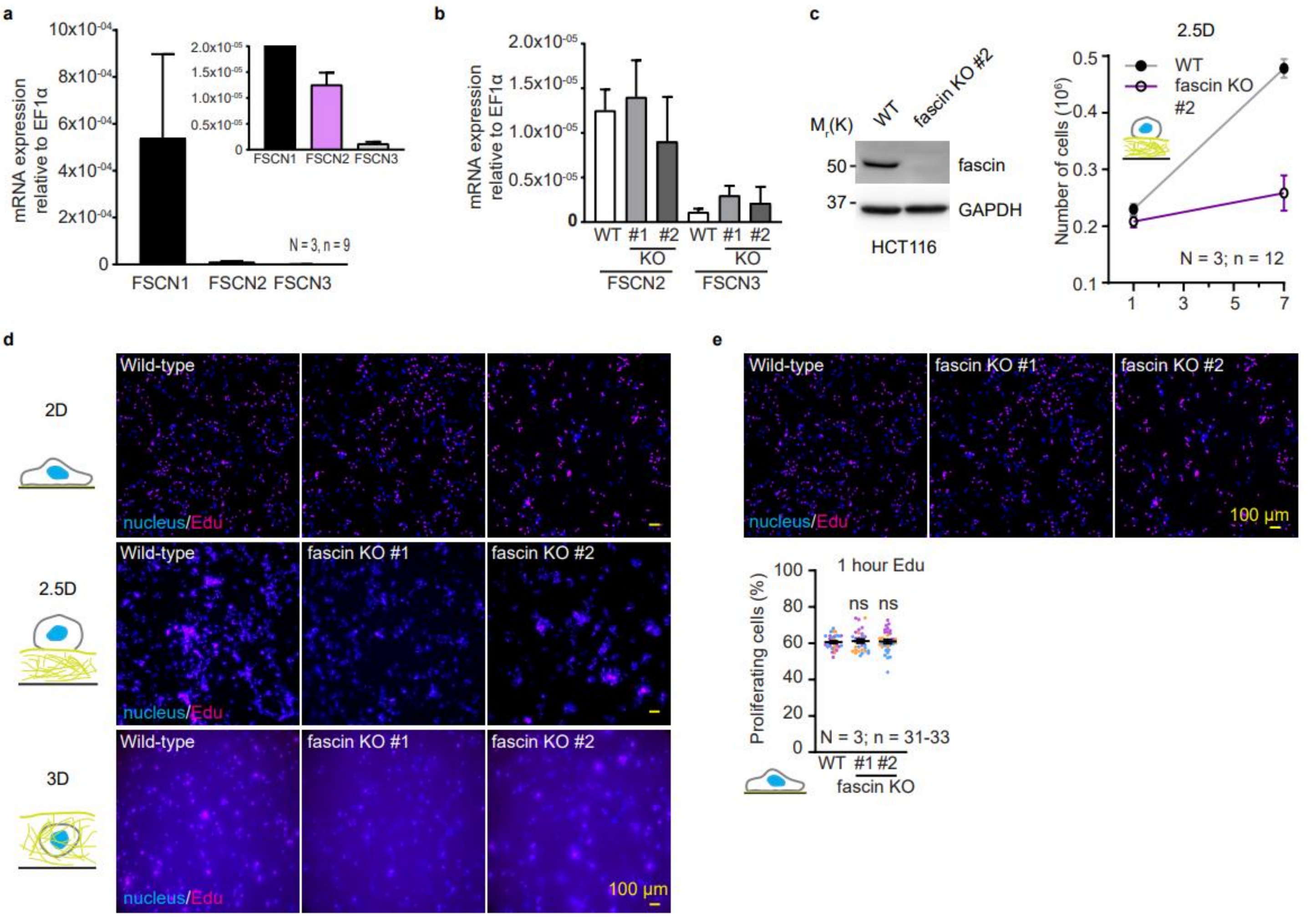


**Figure S1 Fascin promotes cell proliferation in an ECM-dependent manner.**

**a**, HCT116 cells predominantly express fascin-1 (FSCN1) and not fascin-2 (FSCN2) or fascin-3 (FSCN3). $n$ = 9. **b**, Fascin KO does not lead to compensatory expression of fascin-2 or 3. FSCN2 and FSCN3 mRNA levels were determined by qPCR from cDNA isolated from wild-type and fascin KO #1 and #2 HCT116 cells. $n$ = 9. **c**, An independent guide RNA was used to generate a second fascin KO clone and shown to have similar growth defects in 2.5D (see Fig. 1c for additional details). $n$ = 12. **d,** Representative images of our imaging-based Edu proliferations assay for 2D (top), 2.5D (middle) and 3D (bottom) (see Figs. 1d-f for the quantifications). For 2.5D and 3D, maximal projections are shown. **e**, Short Edu pulse reproduced the overnight Edu incubation (compare with Fig. 1d) and overnight Edu incubation did not saturate proliferation. Representative images and quantifications are shown. $n$ = 33 for WT and 32 for KOs #1 and 2. Scale bars are 100 µm. For **e**, unpaired two-tailed Student's $t$-tests were performed; ns , $P > 0.05$. Data are mean ± s.e.m.**.** Each color represents a biological replicate.

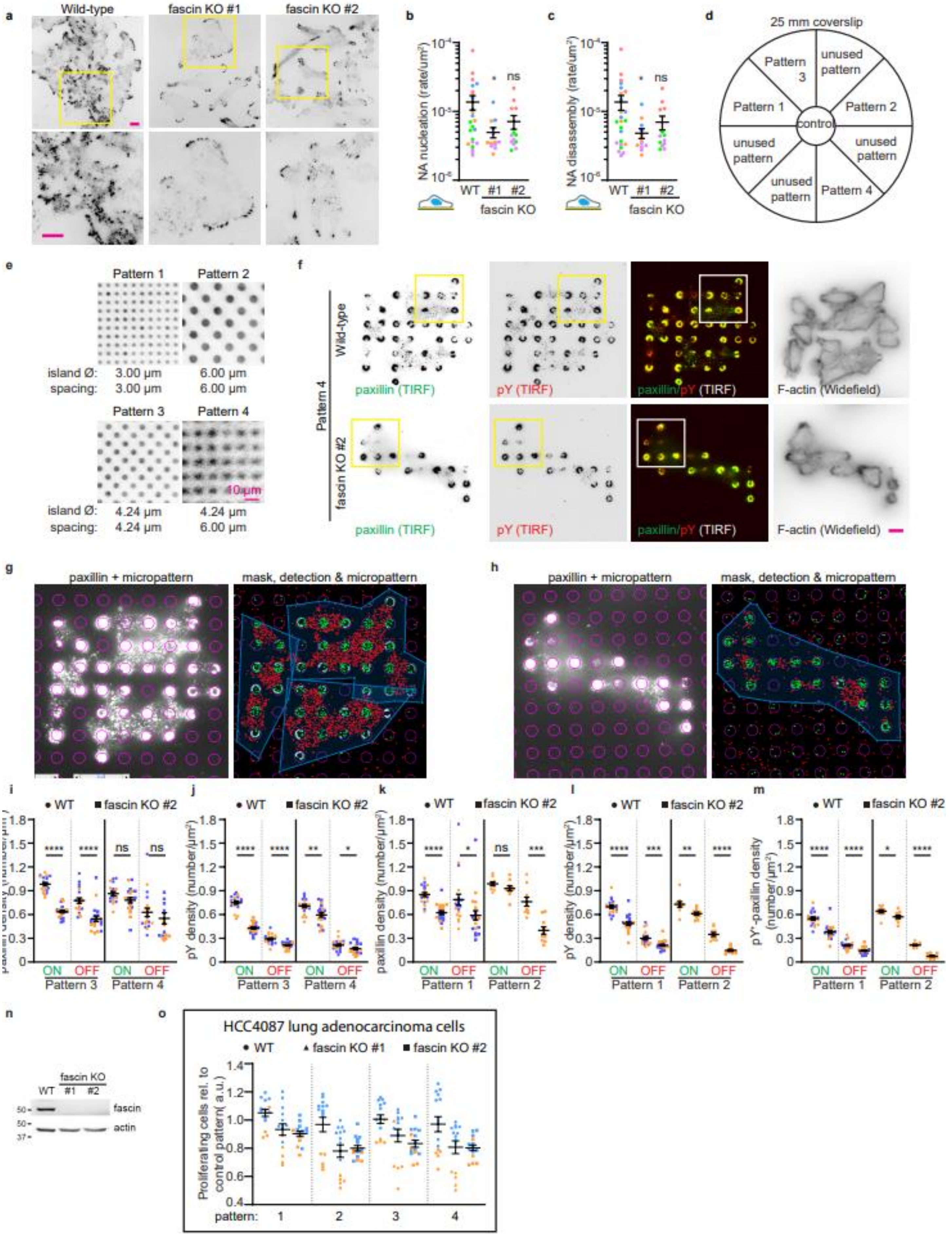

a
Wild-type
fascin KO #1
fascin KO #2
b
NA nucleation (rate/um²)
WT #1 #2
fascin KO
ns
c
NA disassembly (rate/um²)
WT #1 #2
fascin KO
ns
d
25 mm coverslip
Pattern 3
unused pattern
Pattern 1
Pattern 2
control
unused pattern
Pattern 4
e
Pattern 1
Pattern 2
island Ø: 3.00 µm 6.00 µm
spacing: 3.00 µm 6.00 µm
Pattern 3
Pattern 4
10 µm
island Ø: 4.24 µm 4.24 µm
spacing: 4.24 µm 6.00 µm
f
Pattern 4
Wild-type
fascin KO #2
paxillin (TIRF)
pY (TIRF)
paxillin/pY (TIRF)
F-actin (Widefield)
g
paxillin + micropattern
mask, detection & micropattern
h
paxillin + micropattern
mask, detection & micropattern
i
• WT ■ fascin KO #2
ON OFF ON OFF
Pattern 3 Pattern 4
j
pY density (number/µm²)
Pattern 3 Pattern 4
k
paxillin density (number/µm²)
Pattern 1 Pattern 2
l
pY density (number/µm²)
Pattern 1 Pattern 2
m
pY⁺-paxillin density (number/µm²)
Pattern 1 Pattern 2
n
fascin KO
WT #1 #2
fascin
actin
o
HCC4087 lung adenocarcinoma cells
• WT ▲ fascin KO #1 ■ fascin KO #2
Proliferating cells rel. to control pattern( a.u.)
pattern: 1 2 3 4

**Figure S2 Fascin nucleates anchorage-independent nascent adhesion-like structures.**

**a,** Fascin depletion leads to decreased nascent-adhesion (NA) formation. Cells expressing mNeonGreen-tagged paxillin were TIRF imaged. Bottom panels are zoom in of the insets in the top panel. See also Supplementary Movies 7-12 for live cell movies. Scale bar is 10 µm. **b**, Fascin expression is associated with high NA nucleation. **c**, Fascin expression is associated with high NA disassembly. For **b,c**, $n = 25$ for wild-type and 13 for KOs #1 and 2. **d**, Overall micropattern design on a 25 mm glass coverslip used in Figs. 1k-m. **e**, Micropatterned surfaces coated with collagen I and labeled with cy3. Scale bar is 10 µm. **f**, Representative images of cells seeded on micropatterns and stained for indicated antibodies and phalloidin (F-actin). Paxillin and pY were imaged using TIRF. F-actin was imaged using epifluorescence. **g,h**, Micropattern quantification pipeline. Detected micropatterns (magenta circle plots) were overlayed onto the paxillin image. Point sources within the segmented cells (blue box) were detected and classified as ON pattern (green dots) versus OFF pattern (red dots). **i-m**, ON and OFF pattern paxillin (i,k), pY (j,l) and pY-positive ($pY^+$) paxillin densities for the indicated patterns. **n**, Generation of fascin KO HCC4087 lung adenocarcinoma cells using CRISPR/Cas9 with two independent guide RNAs. **o**, Fascin KO cells proliferate less by restricting adhesion area. See Fig. 2l for additional details. For **a,e,f**, scale bars are 10 µm. For **b,d,** one-sided Wilcoxon Rank Sum tests, for i-m,o, two-tailed student *t-tests* were performed. ns, $P > 0.05$; $*P \leq 0.05$, $**P \leq 0.01$, $***P \leq 0.001$, $****P \leq 0.0001$. Data are mean ± s.e.m.**.** Each color represents a biological replicate.

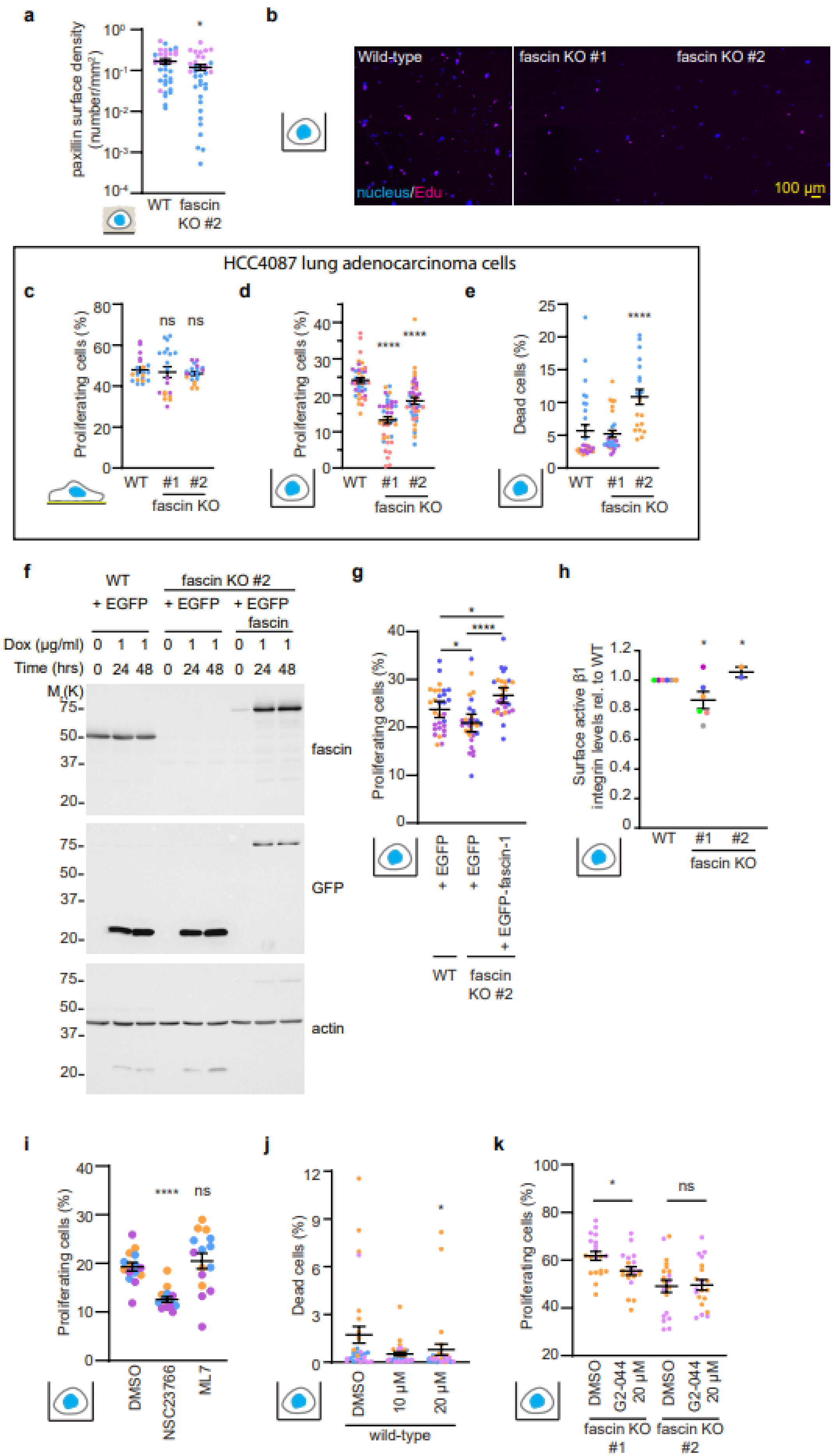
a
paxillin surface density (number/mm²)
WT
fascin KO #2
b
Wild-type
fascin KO #1
fascin KO #2
nucleus/Edu
100 µm
HCC4087 lung adenocarcinoma cells
c
Proliferating cells (%)
WT
#1
#2
fascin KO
d
Proliferating cells (%)
e
Dead cells (%)
f
WT
+ EGFP
fascin KO #2
+ EGFP
+ EGFP fascin
Dox (µg/ml)
Time (hrs)
M (K)
fascin
GFP
actin
g
Proliferating cells (%)
+ EGFP
+ EGFP-fascin-1
WT
fascin KO #2
h
Surface active β1 integrin levels rel. to WT
i
Proliferating cells (%)
DMSO
NSC23766
ML7
j
Dead cells (%)
10 µM
20 µM
wild-type
k
Proliferating cells (%)
G2-044 20 µM
fascin KO #1
fascin KO #2

**Figure S3 Fascin expression supports proliferation in suspension cultures.**

**a,** Fascin KO decreases surface paxillin density of cells cultured in suspension. Cells embedded in Hydrogels were stained with anti-paxillin antibodies and imaged using LSFM. Paxillin speckles at the cell surface were detected and quantified as number per area (density). See Figs. 2a,b, and 3a for additional details. $n$ = 35 for WT and 36 for KO. **b,** Representative images of our imaging-based Edu proliferations assay in suspension. (see Fig. 3c for the quantifications). **c,d,e,** Fascin KO decreases proliferation of HCC4087 lung adenocarcinoma cells in suspension (**d**) but not in 2D (**c**). Cells were incubated with Edu for 1 hour and proliferating cells were determined by our imaging-based proliferation assay. Dead cells were determined by EthD-1 exclusion (**e**). **f**, Tet-inducible rescue of fascin KO cells with EGFP-fascin. Cells were incubated with indicated Doxycycline concentration and time and cell lysates were subjected for western blot analysis with the indicated antibodies. **g**, Ectopic fascin expression rescues proliferation in suspension. Proliferation in suspension of cells from **f** were analyzed using our imaging-based Edu proliferation assay. **h**, Surface active and total beta1 integrin levels were analyzed by FACS. Active beta1 integrin was normalized to total beta1 integrin levels and subsequently normalized to the wild-type controls for plotting. $n$ = 6 for WT and fascin KO #1 and 2 for KO #2. **i**, Cells cultured in suspension were treated with the indicated inhibitors overnight and proliferation determined using our imaging-based method. $n$ = 15. **j**, Wild-type cells cultured in suspension were incubated with indicated concentration of NP-G2-044 overnight and dead cells determined by EthD-1 exclusion assay. $n$ = 30. k, Fascin KO #1 and #2 cells cultured in suspension were treated with 20 µM NP-G2-044 overnight and proliferation determined using our imaging-based method. $n$ = 20. For **a,** one-sided Wilcoxon Rank Sum test, for **c,d,g,h,i,k**, two-tailed student $t$-tests, for **e,j**, two-sided Wilcoxon Rank sum tests were performed. ns, $P > 0.05$; *$P \leq 0.05$, **$P \leq 0.01$, ***$P \leq 0.001$, ****$P \leq 0.0001$. Data are mean ± s.e.m.. Each color represents a biological replicate.

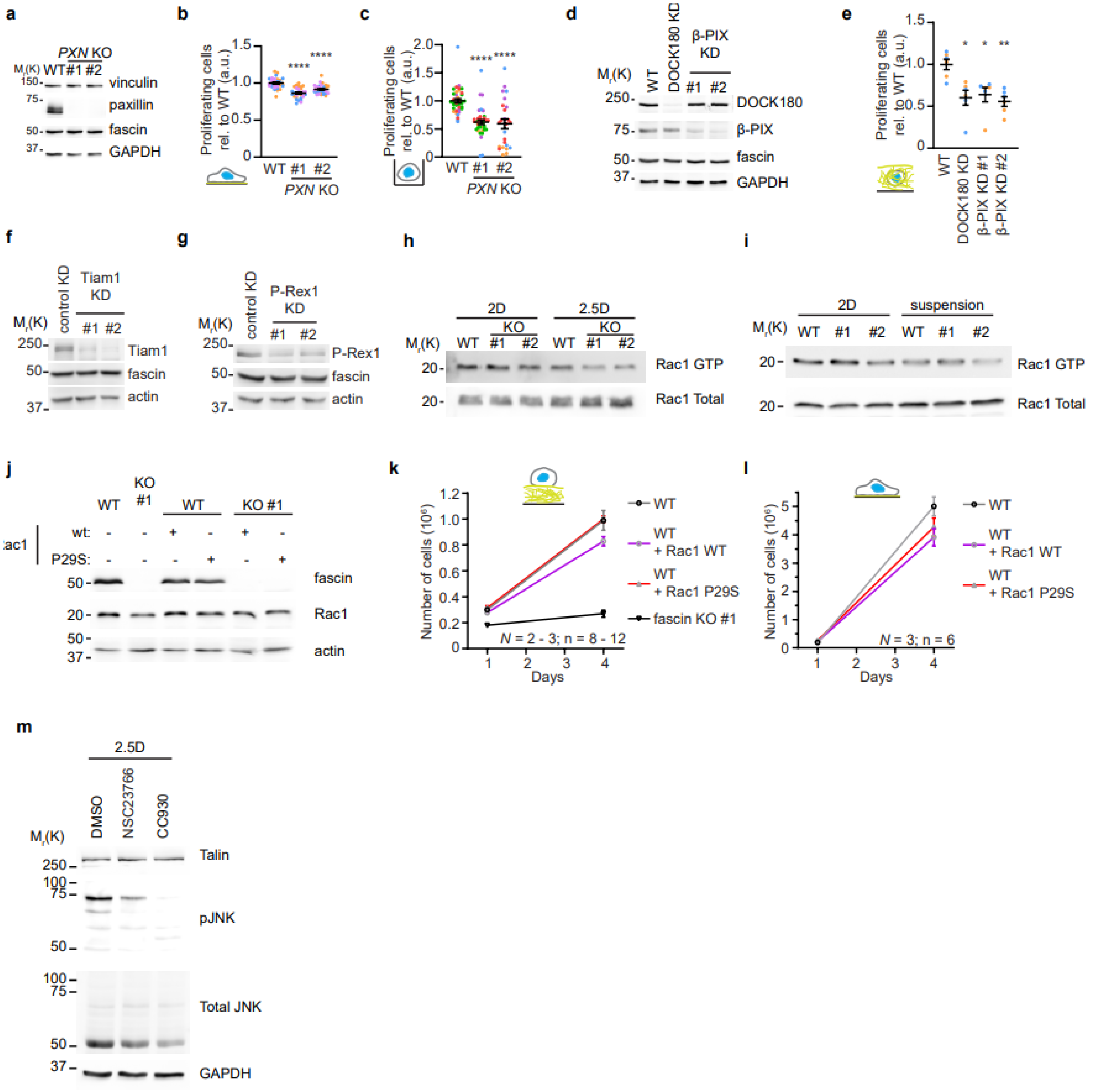

a
PXN KO
Mr(K) WT #1 #2
150
75
50
37
vinculin
paxillin
fascin
GAPDH
b
Proliferating cells rel. to WT (a.u.)
1.5
1.0
0.5
0
WT #1 #2
PXN KO
****
c
2.0
1.5
1.0
0.5
0
d
WT
DOCK180 KD
β-PIX KD
#1 #2
250
75
50
37
DOCK180
β-PIX
fascin
GAPDH
e
*
**
WT
DOCK180 KD
β-PIX KD #1
β-PIX KD #2
f
control KD
Tiam1 KD
Tiam1
actin
g
P-Rex1 KD
P-Rex1
h
2D
2.5D
KO
Rac1 GTP
Rac1 Total
i
suspension
j
KO #1
WT
wt:
P29S:
Rac1
k
Number of cells (10⁶)
Days
WT + Rac1 WT
WT + Rac1 P29S
fascin KO #1
N = 2 - 3; n = 8 - 12
l
N = 3; n = 6
m
DMSO
NSC23766
CC930
Talin
pJNK
Total JNK
100

**Figure S4 PASS is a signaling hub for Rac1**

**a**, Paxillin KO HCT116 cells were generated using two independent guide RNAs. **b**, Paxillin KO has minimal effects on proliferation in 2D. $n = 30$. **c**, Paxillin KO strongly attenuates proliferation in suspension. $n = 39$ for WT and KO #1 and 25 for KO #2. **d,** Generation of DOCK180 and β-PIX knockdown (KD) HCT116 cells using shRNAs. **e,** DOCK180 and β-PIX KD attenuates proliferation in 3D (see Fig. 1f for additional details). $n = 21$-25. **f**, Tiam1 KD HCT116 cells were generated using two independent shRNAs. **g**, P-Rex1 KD HCT116 cells were generated using two independent shRNAs. **h**, Active Rac1 (Rac1 GTP) from cells cultured in 2D or suspension was pulled down using PAK-CRIB beads (see Figs. 4f,g for additional details). **i**, Rac1 GTP from cells cultured in 2D or suspension was pulled down using PAK-CRIB beads. **j**, Western blot characterization of HCT116 cells expressing ectopic Rac1 wild-type (wt) or P29S hyperactive mutant. **k**, Cell growth of wild-type cells expressing either WT or P29S Rac1 over time in 2.5D (see Fig. 4h for additional details). $n > 8$. **l**, Cell growth of wild-type cells expressing either WT or P29S Rac1 over time in 2D. $n = 6$. **m**, Cells cultured in 2.5D were treated with Rac1 (NSC23766) or JNK (CC930) inhibitors overnight and subjected for western blot analysis with the indicated antibodies (see also Fig. 4j for additional details). For **b,c,e,** two-tailed Student *t*-tests were performed. $P > 0.05$; $*P \leq 0.05$, $**P \leq 0.01$, $****P \leq 0.0001$. Data are mean ± s.e.m.**.** Each color represents a biological replicate.

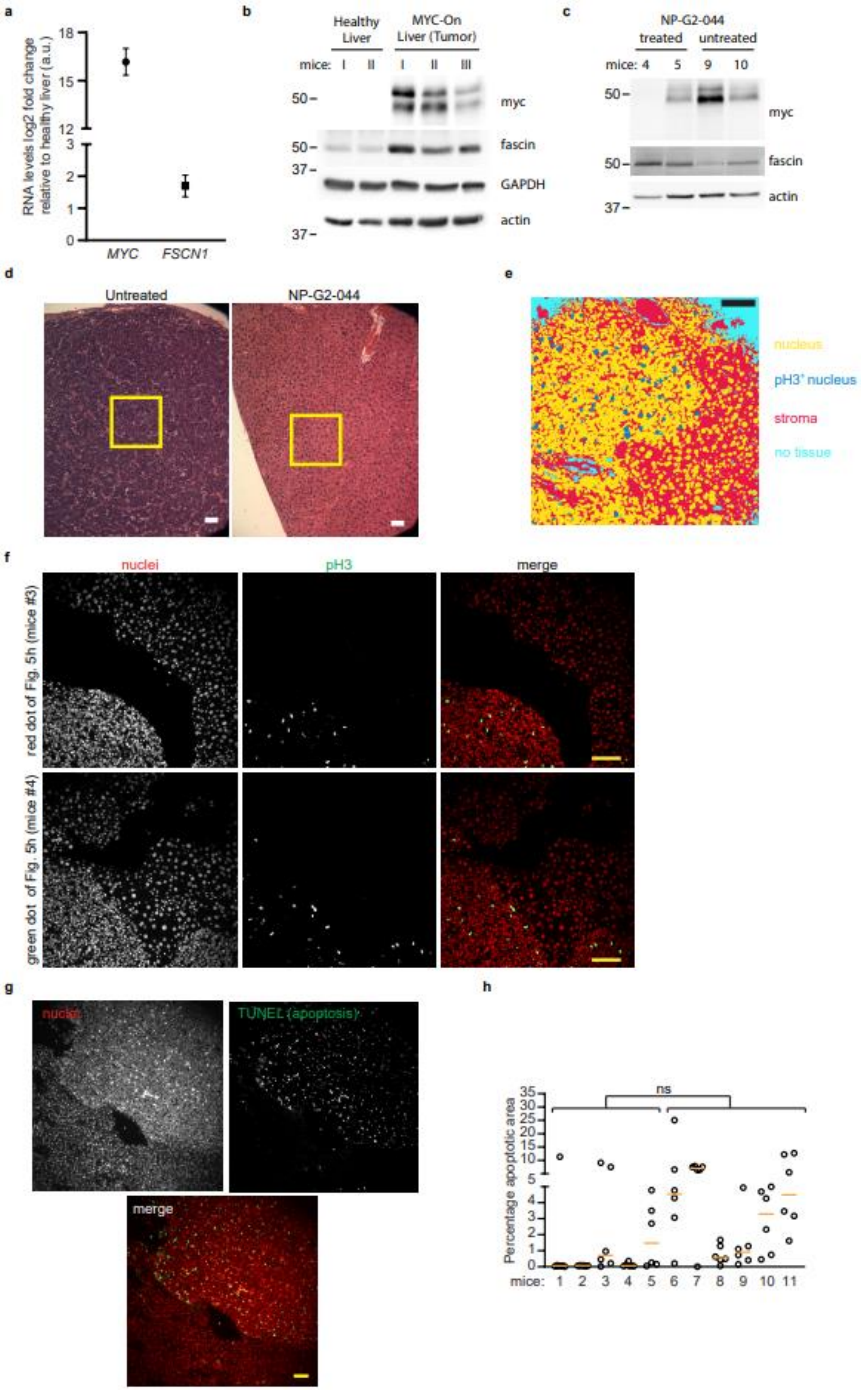
a
RNA levels log2 fold change relative to healthy liver (a.u.)
18
15
12
3
2
1
0
MYC
FSCN1
b
Healthy Liver
MYC-On Liver (Tumor)
mice: I II I II III
50
50
37
37
myc
fascin
GAPDH
actin
c
NP-G2-044
treated
untreated
mice: 4 5 9 10
50
50
37
myc
fascin
actin
d
Untreated
NP-G2-044
e
nucleus
pH3⁺ nucleus
stroma
no tissue
f
nuclei
pH3
merge
red dot of Fig. 5h (mice #3)
green dot of Fig. 5h (mice #4)
g
nuclei
TUNEL (apoptosis)
merge
h
ns
Percentage apoptotic area
35
30
25
20
15
10
5
4
3
2
1
0
mice: 1 2 3 4 5 6 7 8 9 10 11

**Figure S5 NP-G2-044 treatment attenuates proliferation of MYC-induced HCC**

**a**, mRNA expressions of MYC (*MYC*) and fascin (*FSCN1*) isolated from MYC-induced liver tumors normalized to healthy liver controls. n = 3. **b** Western blot analysis of isolated healthy and MYC-On livers for MYC and fascin expression. Each lane represents independent samples. **c**, Western blot analysis of MYC-induced liver tumors with or without NP-G2-044 treatment for MYC and fascin expression. **d,** Uncropped images of H&E-stained tissues. Cropped regions demarked with yellow boxes are presented in Fig. 5f. **e,** An example segmentation result to determine the nucleus (yellow), phospho-Histone3 positive ($pH3^+$) nucleus (dark blue), stroma (red) or no tissue area (cyan) of liver tumor IHCs for the quantification of $pH3^+$ tumor area using ilastik. **f,** Examples of IHC results of liver tumor samples with heterogeneous response (see Fig. 5h for quantification results). Red and green dots of Fig. 5h corresponds to samples from mice #3 and #4, respectively. **g**, A representative image of TUNEL-stained IHC sample. **h**, Quantification of percentage apoptotic cell area per sampled tissue. *n* = 6 per mice. Orange line represents median. All scale bars, 100 µm. For **i**, 2-way ANOVA was performed as statistical analysis. ns, $P > 0.05$.
