## Supplemental Table 1 for "Anchorage-independent cell proliferation promoted by fascin’s F-actin bundling"

**Table S1: Summary of the meta-analysis of fascin's role in migration and proliferation based on Liu et al., 2022**

| | Migration/Invasion | | | Proliferation | | |
|---|---|---|---|---|---|---|
| | Inhibited | No Effect | Unknown | Inhibited | No Effect | Unknown |
| Fascin Depletion | 37 | 1 | 8 | 20 | 7 | 19 |
| | Enhanced | No Effect | Unknown | Enhanced | No Effect | Unknown |
| Fascin Overexpression | 17 | 1 | 2 | 6 | 4 | 10 |
| Total per condition | 54 | 2 | 10 | 26 | 11 | 29 |
| Total of all tested conditions | 56 | | | 37 | | |
