## Supplemental Table 2 for "Anchorage-independent cell proliferation promoted by fascin’s F-actin bundling"

**Table S2: PASS proxisome of wild-type cells cultured in suspension**

| name | ID | Adjusted p-value | Log2 FC |
|---|---|---|---|
| GIT2 | Q14161 | 1.49E-13 | 6.89 |
| PXN | A0A1B0GTU4 | 8.11E-14 | 6.66 |
| SORBS1 | Q9BX66 | 8.11E-14 | 5.73 |
| ARHGEF7 | B1ALK7 | 8.11E-14 | 5.42 |
| GIT1 | Q9Y2X7 | 2.72E-12 | 5.42 |
| PTPN12 | Q05209 | 8.11E-14 | 5.23 |
| CYLD | Q9NQC7 | 8.11E-14 | 5.09 |
| SYNRG | Q9UMZ2 | 8.11E-14 | 5.09 |
| PRAG1 | Q86YV5 | 8.11E-14 | 4.94 |
| PAK1 | Q13153 | 3.85E-08 | 4.68 |
| FSCN1 | Q16658 | 9.78E-07 | 4.65 |
| TNS3 | Q68CZ2 | 8.11E-14 | 4.5 |
| PHLDB1.1 | A0A087WZL0 | 0.000149 | 4.31 |
| PTK2B | Q14289 | 8.11E-14 | 4.16 |
| BRCC3 | P46736 | 1.33E-08 | 4.12 |
| PEAK1 | Q9H792 | 3.50E-10 | 4.08 |
| HEATR5B | Q9P2D3 | 1.66E-12 | 3.83 |
| ABRAXAS2 | Q15018 | 1.42E-05 | 3.72 |
| ZCCHC3 | Q9NUD5 | 0.000227 | 3.72 |
| AHNAK2 | Q8IVF2 | 2.54E-09 | 3.68 |
| UHRF1BP1L | A0JNW5 | 4.99E-09 | 3.6 |
| RPF2 | Q9H7B2 | 5.51E-09 | 3.56 |
| ARFIP2 | A0A087X1E4 | 4.57E-05 | 3.47 |
| YEATS2 | Q9ULM3 | 4.33E-09 | 3.38 |
| SPATA2 | Q9UM82 | 5.25E-05 | 3.37 |
| ARHGEF10L | Q9HCE6 | 4.55E-06 | 3.36 |
| NEDD1 | Q8NHV4 | 0.0176 | 3.35 |
| ATP6V1E1 | P36543 | 0.0405 | 3.35 |
| H0Y858 | H0Y858 | 2.65E-09 | 3.32 |
| SVIL | O95425 | 3.73E-06 | 3.29 |
| AFTPH | Q6ULP2 | 1.66E-07 | 3.22 |
| MPP5 | Q8N3R9 | 3.95E-06 | 3.17 |
| PIP | P12273 | 7.95E-09 | 3.15 |
| DCP1B | Q8IZD4 | 0.000247 | 3.15 |
| FGFR1OP | O95684 | 2.99E-07 | 3.14 |
| ZKSCAN3 | Q9BRR0 | 0.0019 | 3.13 |
| PIMREG | I3L156 | 1.61E-06 | 3.12 |
| SRSF3 | P84103 | 1.07E-05 | 3.12 |
| NCOA3 | Q9Y6Q9 | 0.0182 | 3 |
| GLYR1 | Q49A26 | 0.036 | 3 |
| PPAT | Q06203 | 0.153 | 2.95 |
| SCO2 | O43819 | 0.000146 | 2.94 |
| BCAR3 | O75815 | 0.0022 | 2.94 |
| GAR1 | Q9NY12 | 1.57E-05 | 2.91 |
| ARHGEF40 | Q8TER5 | 0.0626 | 2.91 |

| | | | |
|---|---|---|---|
| ALMS1 | Q8TCU4 | 4.43E-05 | 2.9 |
| GIT2.1 | R4GNG3 | 0.000103 | 2.89 |
| MIB1 | Q86YT6 | 0.000555 | 2.87 |
| FERMT2 | Q96AC1 | 0.024 | 2.87 |
| CDK5RAP2 | Q96SN8 | 0.00557 | 2.86 |
| MACC1 | Q6ZN28 | 1.10E-05 | 2.83 |
| PTBP3 | O95758 | 1.39E-07 | 2.8 |
| NUP58 | Q9BVL2 | 6.08E-05 | 2.8 |
| SSX2IP | Q9Y2D8 | 0.00192 | 2.8 |
| TPRN | Q4KMQ1 | 2.29E-05 | 2.77 |
| TACC3 | Q9Y6A5 | 0.00651 | 2.76 |
| TRIR | Q9BQ61 | 3.58E-08 | 2.75 |
| GIT1.1 | A0A0C4DGN6 | 1.05E-05 | 2.75 |
| ZBTB20 | Q9HC78 | 0.00121 | 2.75 |
| SGO2 | Q562F6 | 0.0685 | 2.74 |
| TANK | Q92844 | 2.62E-06 | 2.73 |
| TNK1 | Q13470 | 0.00012 | 2.72 |
| KNL1 | Q8NG31 | 3.11E-07 | 2.71 |
| ALDOC | P09972 | 0.246 | 2.71 |
| RNF168 | Q8IYW5 | 0.0398 | 2.66 |
| EPHX1 | P07099 | 0.243 | 2.66 |
| DECR1 | Q16698 | 0.0216 | 2.65 |
| GDAP1 | Q8TB36 | 1.87E-05 | 2.64 |
| CEMIP | Q8WUJ3 | 0.00681 | 2.62 |
| CRYBG1 | A0A0J9YWL0 | 8.68E-09 | 2.61 |
| AP2S1 | M0QYZ2 | 0.00395 | 2.61 |
| LMO7 | Q8WWI1 | 1.55E-05 | 2.6 |
| PCM1 | Q15154 | 0.000259 | 2.6 |
| GCC2 | Q8IWJ2 | 3.91E-06 | 2.59 |
| GJC1 | P36383 | 0.000117 | 2.59 |
| EXOC3 | O60645 | 0.00958 | 2.59 |
| MSI2 | Q96DH6 | 0.0171 | 2.59 |
| EIF3K | Q9UBQ5 | 0.232 | 2.59 |
| MYH10 | P35580 | 1.10E-05 | 2.58 |
| PLOD2 | O00469 | 0.0111 | 2.58 |
| CCZ1B | P86790 | 0.00205 | 2.54 |
| RGPD5 | Q99666 | 8.39E-06 | 2.51 |
| DSTN | P60981 | 0.0147 | 2.5 |
| MTERF4 | Q7Z6M4 | 0.00221 | 2.49 |
| CNOT2 | Q9NZN8 | 0.0104 | 2.48 |
| HECW1 | Q76N89 | 0.0248 | 2.47 |
| CTNNA2 | P26232 | 2.75E-05 | 2.46 |
| ABHD10 | Q9NUJ1 | 0.237 | 2.45 |
| CEP131 | Q9UPN4 | 0.000496 | 2.44 |
| PIBF1 | Q8WXW3 | 0.0041 | 2.44 |
| PSTPIP2 | Q9H939 | 0.000952 | 2.42 |
| HIST1H2BJ | P06899 | 0.0106 | 2.42 |

| | | | |
|---|---|---|---|
| DHPS | P49366 | 0.0243 | 2.42 |
| REPS2 | Q8NFH8 | 0.000802 | 2.41 |
| PRKAR2A | P13861 | 0.000102 | 2.4 |
| FAM193B | Q96PV7 | 0.161 | 2.4 |
| PARVB | Q9HBI1 | 0.00015 | 2.39 |
| GNAZ | P19086 | 0.000211 | 2.39 |
| BCAS3 | Q9H6U6 | 2.33E-06 | 2.37 |
| CEP170.3 | H0YB92 | 0.0489 | 2.36 |
| MGA.1 | H3BU53 | 0.0631 | 2.36 |
| SPECC1L | Q69YQ0 | 0.105 | 2.35 |
| DCP1A | Q9NPI6 | 0.000585 | 2.31 |
| NEK1 | Q96PY6 | 0.0107 | 2.3 |
| LIMS4 | A0A0J9YXC7 | 0.02 | 2.3 |
| ARHGEF6 | Q15052 | 0.00119 | 2.29 |
| NCK2 | O43639 | 0.00321 | 2.28 |
| PCNX3 | Q9H6A9 | 0.000678 | 2.27 |
| TNFRSF10A | O00220 | 0.00403 | 2.27 |
| HAUS3 | Q68CZ6 | 0.00426 | 2.27 |
| WDR70 | Q9NW82 | 0.000327 | 2.24 |
| FBXO34 | Q9NWN3 | 0.282 | 2.24 |
| SKIV2L | Q15477 | 0.000138 | 2.23 |
| EVPL | Q92817 | 0.00279 | 2.23 |
| ARID4A | P29374 | 0.379 | 2.23 |
| TAPT1 | Q6NXT6 | 3.77E-05 | 2.22 |
| SPECC1 | Q5M775 | 0.00754 | 2.22 |
| RRAS2 | P62070 | 0.315 | 2.22 |
| SUMO1 | B8ZZN6 | 0.000509 | 2.21 |
| C2CD3 | Q4AC94 | 0.00585 | 2.21 |
| CCP110 | O43303 | 0.0298 | 2.21 |
| STIM1 | G0XQ39 | 0.0711 | 2.21 |
| DIS3L | Q8TF46 | 6.96E-05 | 2.2 |
| ITSN2 | Q9NZM3 | 0.00506 | 2.2 |
| CYR61 | O00622 | 0.00181 | 2.19 |
| CCDC14 | Q49A88 | 0.0136 | 2.19 |
| CEP120 | Q8N960 | 0.00388 | 2.18 |
| GDF15 | Q99988 | 3.18E-07 | 2.17 |
| HMGXB4 | Q9UGU5 | 0.276 | 2.17 |
| SMAD4 | Q13485 | 0.00144 | 2.16 |
| AGFG1 | P52594 | 0.0152 | 2.16 |
| PSMA6 | G3V5Z7 | 0.505 | 2.16 |
| MYH9 | P35579 | 0.0307 | 2.15 |
| RBM15B | Q8NDT2 | 6.93E-06 | 2.13 |
| LPP | Q93052 | 0.0196 | 2.13 |
| SYTL4 | Q96C24 | 0.0524 | 2.13 |
| SEC16A.1 | J3KNL6 | 0.0507 | 2.12 |
| ERCC1 | P07992 | 0.00698 | 2.11 |
| TP53BP2 | Q13625 | 0.00963 | 2.11 |

| | | | |
|---|---|---|---|
| SKA3 | Q8IX90 | 2.66E-07 | 2.1 |
| PTK2 | E7ESA6 | 4.67E-07 | 2.1 |
| CEP350 | Q5VT06 | 0.00084 | 2.1 |
| AMOTL1 | Q8IY63 | 0.00137 | 2.1 |
| HAUS8 | Q9BT25 | 0.00322 | 2.1 |
| MED16.1 | B9TX12 | 0.0484 | 2.09 |
| EDC3 | Q96F86 | 5.81E-05 | 2.08 |
| ITIH2 | P19823 | 0.362 | 2.08 |
| TPI1 | P60174 | 0.411 | 2.07 |
| ARAP3 | Q8WWN8 | 0.021 | 2.04 |
| AZGP1 | P25311 | 0.0621 | 2.04 |
| INTS4 | Q96HW7 | 0.151 | 2.04 |
| LGALS3 | P17931 | 0.282 | 2.04 |
| SSBP1 | Q04837 | 0.136 | 2.04 |
| DOCK6 | Q96HP0 | 0.000219 | 2.03 |
| TEP1 | Q99973 | 0.00108 | 2.02 |
| AHNAK | Q09666 | 0.00429 | 2.02 |
| ERRFI1 | Q9UJM3 | 0.0307 | 2.02 |
| PAWR | Q96IZ0 | 1.85E-05 | 2.01 |
| MPRIP.1 | H0Y2S9 | 0.00121 | 2.01 |
| S100A9 | P06702 | 0.00367 | 2.01 |
| EBNA1BP2 | H7C2Q8 | 0.0784 | 2.01 |
| TPD52 | P55327 | 0.254 | 2.01 |
| ISY1 | Q9ULR0 | 0.00815 | 2 |
| HAUS5 | O94927 | 0.00883 | 2 |
| ATG4B | Q9Y4P1 | 0.00837 | 1.99 |
| OFD1 | O75665 | 0.0534 | 1.99 |
| E2F7 | Q96AV8 | 3.86E-06 | 1.98 |
| HAUS4 | Q9H6D7 | 0.227 | 1.98 |
| RELL1 | Q8IUW5 | 0.00359 | 1.97 |
| KIAA1551 | Q9HCM1 | 0.281 | 1.97 |
| CEP97 | Q8IW35 | 9.10E-05 | 1.96 |
| PLEK2 | Q9NYT0 | 0.0628 | 1.96 |
| TPP1 | O14773 | 0.0143 | 1.95 |
| PAK2 | Q13177 | 0.0231 | 1.95 |
| ARHGAP10.1 | H0Y8W5 | 0.0962 | 1.95 |
| CD2AP | Q9Y5K6 | 0.273 | 1.95 |
| GOLGA3 | Q08378 | 0.0363 | 1.93 |
| C12orf10 | Q9HB07 | 0.278 | 1.93 |
| ITPR2 | Q14571 | 0.000551 | 1.92 |
| NUP37 | Q8NFH4 | 0.00582 | 1.91 |
| EXOSC5 | Q9NQT4 | 0.107 | 1.91 |
| MARF1 | Q9Y4F3 | 0.0135 | 1.9 |
| WRAP73 | Q9P2S5 | 0.00887 | 1.89 |
| RANGAP1 | P46060 | 0.0421 | 1.89 |
| CCDC137 | I3L0U5 | 0.0784 | 1.89 |
| NBAS | A2RRP1 | 0.0761 | 1.89 |

| | | | |
|---|---|---|---|
| KANK1 | Q14678 | 4.24E-06 | 1.88 |
| AKAP9 | Q99996 | 0.00553 | 1.88 |
| HIST1H2BN | U3KQK0 | 0.0318 | 1.87 |
| RFX5 | P48382 | 0.136 | 1.87 |
| POLR2C | P19387 | 0.0463 | 1.85 |
| ZNF724 | A8MTY0 | 0.375 | 1.85 |
| SRCIN1 | A0A087X165 | 0.00108 | 1.84 |
| PATJ | Q8NI35 | 0.0162 | 1.84 |
| CCDC138 | Q96M89 | 0.139 | 1.84 |
| MBTD1 | Q05BQ5 | 0.0647 | 1.84 |
| COG2 | Q14746 | 0.464 | 1.83 |
| TRUB1 | Q8WWH5 | 0.205 | 1.83 |
| H3F3A | P84243 | 9.27E-05 | 1.81 |
| CASKIN2 | Q8WXE0 | 0.0012 | 1.81 |
| SYNJ2 | O15056 | 0.0387 | 1.81 |
| IPPK | Q9H8X2 | 0.297 | 1.81 |
| PIK3C3 | A0A1W2PS05 | 0.308 | 1.8 |
| RANBP2 | P49792 | 2.07E-05 | 1.78 |
| MAGI1.2 | H7C4S7 | 0.00605 | 1.78 |
| SOWAHC | Q53LP3 | 0.00793 | 1.76 |
| RUFY1 | Q96T51 | 0.0334 | 1.76 |
| MYH14 | Q7Z406 | 0.00918 | 1.75 |
| CDC42 | P60953 | 0.41 | 1.75 |
| SCRIB.1 | H0YDF9 | 0.0142 | 1.74 |
| PSAP | C9JIZ6 | 0.415 | 1.74 |
| TSC1 | Q92574 | 0.000178 | 1.73 |
| ZNF680 | Q8NEM1 | 0.00166 | 1.73 |
| TBK1 | Q9UHD2 | 0.125 | 1.73 |
| TSC2 | P49815 | 0.29 | 1.73 |
| ARMC9 | Q7Z3E5 | 0.0197 | 1.72 |
| RPL22L1 | H0Y8C2 | 0.0375 | 1.72 |
| MED4 | Q9NPJ6 | 0.0792 | 1.72 |
| ZNF251 | Q9BRH9 | 0.206 | 1.72 |
| CDC42BPG | Q6DT37 | 0.000246 | 1.71 |
| 7-Mar | Q9H992 | 0.158 | 1.71 |
| TRIP4 | Q15650 | 0.0841 | 1.71 |
| BABAM2 | Q9NXR7 | 0.0209 | 1.7 |
| PPOX | P50336 | 0.0652 | 1.7 |
| SCRIB | A0A0G2JPP5 | 0.0231 | 1.69 |
| CASC3 | O15234 | 0.183 | 1.69 |
| HLA-A.1 | A0A140T913 | 0.154 | 1.69 |
| NPHP4 | O75161 | 0.253 | 1.69 |
| PDCD4 | Q53EL6 | 0.361 | 1.69 |
| PURB | Q96QR8 | 0.13 | 1.69 |
| PPAN | H7C446 | 0.00081 | 1.68 |
| PDE4DIP | A0A075B749 | 0.0177 | 1.68 |
| MTDH.1 | E5RJU9 | 0.287 | 1.68 |

| | | | |
|---|---|---|---|
| FAM50A | Q14320 | 0.000145 | 1.67 |
| PLAUR | Q03405 | 0.22 | 1.66 |
| SRP14 | P37108 | 0.283 | 1.66 |
| PLEKHA6 | A0A1B0GUN5 | 0.0109 | 1.65 |
| MPRIP | Q6WCQ1 | 0.0124 | 1.65 |
| MPRIP.2 | A0A1W2PPU9 | 0.0297 | 1.65 |
| RPLP1 | P05386 | 0.186 | 1.65 |
| TMEM259 | Q4ZIN3 | 0.185 | 1.65 |
| IBTK | Q9P2D0 | 0.195 | 1.64 |
| TRIM41 | Q8WV44 | 0.535 | 1.64 |
| ASAP1 | Q9ULH1 | 0.00336 | 1.63 |
| LARP1B | Q659C4 | 0.0199 | 1.63 |
| AMOTL2 | Q9Y2J4 | 0.121 | 1.63 |
| TNRC18 | H9KVB4 | 0.466 | 1.63 |
| CNOT7 | Q9UIV1 | 0.155 | 1.62 |
| NOL10 | Q9BSC4 | 0.636 | 1.62 |
| ITM2C | Q9NQX7 | 0.00648 | 1.61 |
| GULP1 | Q9UBP9 | 0.335 | 1.61 |
| MRPS15 | P82914 | 0.171 | 1.61 |
| PEX14 | O75381 | 0.657 | 1.6 |
| WDR46.1 | A0A0G2JJL1 | 0.0927 | 1.59 |
| DOCK7 | Q96N67 | 0.00371 | 1.58 |
| ZFYVE16 | Q7Z3T8 | 0.104 | 1.58 |
| ZNF468 | Q5VIY5 | 0.0324 | 1.57 |
| LAP3 | P28838 | 0.513 | 1.57 |
| MIDN | Q504T8 | 0.0869 | 1.57 |
| ERCC6L | Q2NKX8 | 0.0152 | 1.56 |
| GAPVD1 | Q14C86 | 0.291 | 1.56 |
| SSFA2 | P28290 | 0.0784 | 1.56 |
| TBC1D2 | Q9BYX2 | 0.00516 | 1.55 |
| ATP6AP1 | Q15904 | 0.626 | 1.55 |
| CRKL | P46109 | 0.389 | 1.55 |
| KIAA1217 | Q5T5P2 | 0.102 | 1.55 |
| RALY | Q9UKM9 | 0.086 | 1.55 |
| PPL | O60437 | 0.0484 | 1.54 |
| MISP | Q8IVT2 | 0.0943 | 1.54 |
| STRN | O43815 | 0.0613 | 1.54 |
| TROAP | F8W130 | 0.147 | 1.54 |
| ARHGAP1 | Q07960 | 0.56 | 1.53 |
| MYL12A | J3QRS3 | 0.285 | 1.53 |
| WDCP | Q9H6R7 | 0.419 | 1.53 |
| COTL1 | Q14019 | 0.272 | 1.52 |
| PCNT | O95613 | 0.114 | 1.52 |
| VPS33B | Q9H267 | 0.185 | 1.52 |
| ZW10 | O43264 | 0.468 | 1.52 |
| CNOT11 | Q9UKZ1 | 0.015 | 1.51 |
| ITPR3 | Q14573 | 0.0215 | 1.51 |

| | | | |
|---|---|---|---|
| VCPIP1 | Q96JH7 | 0.402 | 1.51 |
| ZNF622 | Q969S3 | 0.485 | 1.51 |
| C11orf98 | A0A0B4J220 | 0.132 | 1.5 |
| MYC.1 | A0A087WUS5 | 0.5 | 1.5 |
| HIST1H4A | P62805 | 0.00357 | 1.49 |
| LRRFIP2 | Q9Y608 | 0.0261 | 1.49 |
| AASDHPPT | Q9NRN7 | 0.137 | 1.49 |
| DCD | P81605 | 0.504 | 1.49 |
| HNRNPD.1 | D6RBQ9 | 0.176 | 1.49 |
| SLC39A10 | Q9ULF5 | 0.366 | 1.49 |
| KANK2 | Q63ZY3 | 0.0151 | 1.48 |
| GIGYF2 | Q6Y7W6 | 0.0311 | 1.48 |
| FRYL | O94915 | 0.258 | 1.47 |
| GPN1 | Q9HCN4 | 0.155 | 1.47 |
| LRRC40 | Q9H9A6 | 0.581 | 1.47 |
| SNAPC3 | Q92966 | 0.0974 | 1.47 |
| STRN3 | Q13033 | 0.309 | 1.47 |
| METTL8 | B3KW44 | 0.444 | 1.46 |
| PPP1R9A | Q9ULJ8 | 0.157 | 1.46 |
| TNRC6B | Q9UPQ9 | 0.155 | 1.46 |
| ZNF708 | P17019 | 0.176 | 1.46 |
| EDC4 | Q6P2E9 | 0.379 | 1.45 |
| GAK | O14976 | 0.0775 | 1.45 |
| H2AFZ | P0C0S5 | 0.155 | 1.45 |
| SH2D3A | Q9BRG2 | 0.319 | 1.45 |
| NHS | Q6T4R5 | 0.00279 | 1.44 |
| BCAR1 | P56945 | 0.00968 | 1.44 |
| ARHGAP21 | Q5T5U3 | 0.0446 | 1.44 |
| ATAT1 | Q5SQI0 | 0.0698 | 1.44 |
| TACC2 | O95359 | 0.0836 | 1.44 |
| SEC16A | F1T0I1 | 0.0329 | 1.43 |
| BABAM1 | Q9NWV8 | 0.142 | 1.43 |
| KPRP | Q5T749 | 0.196 | 1.43 |
| BRMS1L | Q5PSV4 | 0.189 | 1.42 |
| FABP5 | Q01469 | 0.644 | 1.42 |
| SREK1.1 | E5RFV3 | 0.731 | 1.42 |
| CRYBG3 | Q68DQ2 | 0.154 | 1.41 |
| PSMA1 | P25786 | 0.673 | 1.41 |
| SPAG9 | O60271 | 0.492 | 1.41 |
| TOP2B | Q02880 | 0.293 | 1.41 |
| DYNC1LI2 | O43237 | 0.384 | 1.4 |
| MARK2.1 | Q7KZI7 | 0.361 | 1.4 |
| RGS12 | O14924 | 0.36 | 1.4 |
| WASHC5 | Q12768 | 0.44 | 1.4 |
| ADSS | P30520 | 0.687 | 1.39 |
| AQR | O60306 | 0.248 | 1.39 |
| MAPRE1 | Q15691 | 0.454 | 1.39 |

| | | | |
|---|---|---|---|
| PURA | Q00577 | 0.598 | 1.39 |
| CNOT1 | A5YKK6 | 0.0395 | 1.38 |
| AGO3 | Q9H9G7 | 0.413 | 1.38 |
| MGA | Q8IWI9 | 0.452 | 1.38 |
| NHEJ1 | Q9H9Q4 | 0.236 | 1.38 |
| 10-Sep | B5ME97 | 0.232 | 1.38 |
| HABP4 | Q5JVS0 | 0.191 | 1.37 |
| IDH2 | P48735 | 0.288 | 1.37 |
| ZNF7 | P17097 | 0.314 | 1.37 |
| DDX60 | Q8IY21 | 0.678 | 1.36 |
| KIF11 | P52732 | 0.39 | 1.36 |
| MCRS1 | Q96EZ8 | 0.465 | 1.36 |
| S100A14 | Q9HCY8 | 0.0796 | 1.36 |
| SRP9 | P49458 | 0.133 | 1.36 |
| MYO5B | Q9ULV0 | 0.00409 | 1.35 |
| LGALS8.1 | Q5T3P9 | 0.439 | 1.35 |
| MED23 | Q5JWT2 | 0.502 | 1.35 |
| SACS | Q9NZJ4 | 0.0951 | 1.35 |
| PDLIM5 | Q96HC4 | 0.548 | 1.34 |
| RNF220 | Q5VTB9 | 0.00551 | 1.33 |
| COG6 | Q9Y2V7 | 0.596 | 1.33 |
| DMAP1 | Q9NPF5 | 0.391 | 1.33 |
| GPC4 | O75487 | 0.42 | 1.33 |
| SIPA1L3 | O60292 | 0.3 | 1.33 |
| TBCE | Q15813 | 0.464 | 1.33 |
| ASCC3 | Q8N3C0 | 0.00768 | 1.32 |
| GPHN | F5H039 | 0.632 | 1.32 |
| LRPAP1 | P30533 | 0.698 | 1.32 |
| MAT2B | Q9NZL9 | 0.644 | 1.32 |
| PSMB5 | P28074 | 0.49 | 1.32 |
| ZBTB7B | O15156 | 0.742 | 1.32 |
| CEP55 | Q53EZ4 | 0.573 | 1.31 |
| MYADM | Q96S97 | 0.39 | 1.31 |
| SLC2A1 | P11166 | 0.198 | 1.31 |
| SOD1 | P00441 | 0.691 | 1.31 |
| STXBP2 | Q15833 | 0.582 | 1.31 |
| WNK1 | Q9H4A3 | 0.336 | 1.31 |
| ASCC2 | Q9H1I8 | 0.0488 | 1.3 |
| ABHD12 | Q8N2K0 | 0.114 | 1.3 |
| CENPE | Q02224 | 0.435 | 1.3 |
| FAM193A | A0A1B0GVL4 | 0.494 | 1.3 |
| BOD1L1 | Q8NFC6 | 0.0884 | 1.29 |
| DAB2IP | Q5VWQ8 | 0.686 | 1.29 |
| NCOA4 | Q13772 | 0.431 | 1.29 |
| USP19 | O94966 | 0.558 | 1.29 |
| CRAMP1 | Q96RY5 | 0.759 | 1.28 |
| OGFR | Q9NZT2 | 0.629 | 1.28 |

| | | | |
|---|---|---|---|
| USP14 | P54578 | 0.63 | 1.28 |
| CDK11A | Q9UQ88 | 0.709 | 1.27 |
| INPP5K | Q9BT40 | 0.203 | 1.27 |
| MINPP1 | Q9UNW1 | 0.139 | 1.27 |
| MUS81 | Q96NY9 | 0.56 | 1.26 |
| NUP88 | Q99567 | 0.463 | 1.26 |
| WDR46 | O15213 | 0.298 | 1.26 |
| RPL22 | P35268 | 0.049 | 1.25 |
| CSNK1G2 | P78368 | 0.142 | 1.25 |
| FAM204A | Q9H8W3 | 0.391 | 1.25 |
| RPL36AL | Q969Q0 | 0.0181 | 1.24 |
| N4BP3 | O15049 | 0.12 | 1.24 |
| LRBA.1 | H0YAC6 | 0.34 | 1.23 |
| MRPS23 | Q9Y3D9 | 0.224 | 1.23 |
| RPGRIP1L | Q68CZ1 | 0.523 | 1.23 |
| USP12 | O75317 | 0.0689 | 1.23 |
| FGD3 | Q5JSP0 | 0.115 | 1.22 |
| GNAS | Q5JWF2 | 0.743 | 1.22 |
| MRPL19 | P49406 | 0.413 | 1.22 |
| THADA | Q6YHU6 | 0.431 | 1.22 |
| TEDC2 | Q7L2K0 | 0.0245 | 1.21 |
| FAM98B | Q52LJ0 | 0.646 | 1.21 |
| MPST | P25325 | 0.267 | 1.21 |
| CFAP97 | Q9P2B7 | 0.616 | 1.2 |
| H1FX | Q92522 | 0.553 | 1.2 |
| MRRF | Q96E11 | 0.199 | 1.2 |
| PATL1 | Q86TB9 | 0.556 | 1.2 |
| RPL21 | P46778 | 0.624 | 1.2 |
| TLN1 | Q9Y490 | 0.0591 | 1.2 |
| TNPO2 | O14787 | 0.696 | 1.2 |
| MAP4.1 | P27816 | 0.031 | 1.19 |
| CIP2A | A0A087X211 | 0.602 | 1.19 |
| GPRC5A | Q8NFJ5 | 0.713 | 1.19 |
| PI4K2A | Q9BTU6 | 0.653 | 1.19 |
| POLR3H | Q9Y535 | 0.128 | 1.19 |
| LGALS8 | F6V2D4 | 0.741 | 1.17 |
| NSDHL | Q15738 | 0.687 | 1.17 |
| 11-Sep | D6RER5 | 0.68 | 1.17 |
| TMOD3 | Q9NYL9 | 0.442 | 1.17 |
| AKAP13 | Q12802 | 0.565 | 1.16 |
| COG1 | Q8WTW3 | 0.56 | 1.16 |
| MFGE8 | Q08431 | 0.543 | 1.16 |
| PLXNB2 | O15031 | 0.663 | 1.16 |
| PPP1R15B | Q5SWA1 | 0.306 | 1.16 |
| AIMP2 | Q13155 | 0.582 | 1.15 |
| C21orf2 | O43822 | 0.476 | 1.15 |
| CEP290 | J3KNF5 | 0.188 | 1.15 |

| | | | |
|---|---|---|---|
| NDC80 | O14777 | 0.548 | 1.15 |
| SHROOM3 | Q8TF72 | 0.411 | 1.15 |
| SLC12A2 | P55011 | 0.685 | 1.15 |
| DIAPH3 | Q9NSV4 | 0.0415 | 1.14 |
| ATP2B1 | P20020 | 0.715 | 1.14 |
| CNKSR2 | Q8WXI2 | 0.115 | 1.14 |
| TAF4B | Q92750 | 0.592 | 1.14 |
| FLOT2 | E7EMK3 | 0.0415 | 1.13 |
| GSS | P48637 | 0.633 | 1.13 |
| HOOK1 | Q9UJC3 | 0.581 | 1.13 |
| PSMD6 | Q15008 | 0.63 | 1.13 |
| RAB3GAP1 | Q15042 | 0.542 | 1.13 |
| SRSF7 | Q16629 | 0.575 | 1.13 |
| H1F0 | P07305 | 0.262 | 1.12 |
| HSD17B11 | Q8NBQ5 | 0.739 | 1.12 |
| KIAA1671 | Q9BY89 | 0.287 | 1.12 |
| KIN | O60870 | 0.0794 | 1.12 |
| PEX6 | Q13608 | 0.138 | 1.12 |
| SLC9A3R2 | Q15599 | 0.647 | 1.12 |
| COIL | P38432 | 0.726 | 1.11 |
| LRCH1 | Q9Y2L9 | 0.619 | 1.11 |
| CENPJ | Q9HC77 | 0.517 | 1.1 |
| CEP95 | Q96GE4 | 0.427 | 1.1 |
| FLNA | P21333 | 0.284 | 1.1 |
| GCDH | Q92947 | 0.484 | 1.1 |
| OXSM | Q9NWU1 | 0.302 | 1.1 |
| RAI14 | Q9P0K7 | 0.422 | 1.1 |
| SCYL2 | Q6P3W7 | 0.616 | 1.1 |
| TTF2 | Q9UNY4 | 0.113 | 1.1 |
| ZMYM4 | Q5VZL5 | 0.759 | 1.1 |
| ZNF568 | A0A087WZ84 | 0.585 | 1.1 |
| SNRPE | P62304 | 0.0416 | 1.09 |
| CEBPZ | Q03701 | 0.771 | 1.09 |
| EPHB2 | P29323 | 0.371 | 1.09 |
| MAPT.1 | A0A0G2JMX7 | 0.723 | 1.09 |
| FAM83H | Q6ZRV2 | 0.316 | 1.08 |
| HLA-C.1 | Q95604 | 0.466 | 1.08 |
| PLEKHG1 | Q9ULL1 | 0.275 | 1.08 |
| TADA1 | Q96BN2 | 0.689 | 1.08 |
| TNKS1BP1 | Q9C0C2 | 0.269 | 1.08 |
| ABCB10 | Q9NRK6 | 0.438 | 1.07 |
| AGO1 | Q9UL18 | 0.203 | 1.07 |
| INO80 | Q9ULG1 | 0.794 | 1.07 |
| JMJD1C | Q15652 | 0.662 | 1.07 |
| RAI1 | Q7Z5J4 | 0.761 | 1.07 |
| YLPM1 | P49750 | 0.52 | 1.07 |
| ACSF3 | Q4G176 | 0.71 | 1.06 |

| | | | |
|---|---|---|---|
| CHMP2A | O43633 | 0.669 | 1.06 |
| PHLDB2 | Q86SQ0 | 0.484 | 1.06 |
| SPTAN1 | Q13813 | 0.0964 | 1.06 |
| ST7 | H7BXS2 | 0.658 | 1.06 |
| AGO2 | Q9UKV8 | 0.431 | 1.05 |
| IGHG1 | A0A0A0MS08 | 0.388 | 1.05 |
| MED15 | Q96RN5 | 0.573 | 1.05 |
| NT5C2 | P49902 | 0.67 | 1.05 |
| RAB5A | P20339 | 0.608 | 1.05 |
| SH3KBP1 | Q96B97 | 0.672 | 1.05 |
| SPRYD3 | F8VWW7 | 0.464 | 1.05 |
| VWA8 | A3KMH1 | 0.509 | 1.05 |
| CCDC9 | Q9Y3X0 | 0.189 | 1.04 |
| CLNS1A | P54105 | 0.665 | 1.04 |
| DCPS | Q96C86 | 0.635 | 1.04 |
| DYNLL1 | P63167 | 0.129 | 1.04 |
| MED13L | Q71F56 | 0.724 | 1.04 |
| PLPP2 | O43688 | 0.601 | 1.04 |
| PPP1R9B | D3DTX6 | 0.533 | 1.04 |
| TCF25.2 | H3BSS3 | 0.69 | 1.04 |
| DVL3 | Q92997 | 0.0412 | 1.03 |
| ANAPC1 | Q9H1A4 | 0.126 | 1.03 |
| DIP2C | A0A0U1RQW6 | 0.141 | 1.03 |
| DNAJC25 | Q9H1X3 | 0.277 | 1.03 |
| DOCK8 | Q8NF50 | 0.695 | 1.03 |
| IFT57 | Q9NWB7 | 0.279 | 1.03 |
| LIMCH1 | Q9UPQ0 | 0.5 | 1.03 |
| PARD6B | Q9BYG5 | 0.663 | 1.03 |
| SH2D4A | Q9H788 | 0.639 | 1.03 |
| XAB2 | Q9HCS7 | 0.136 | 1.03 |
| ZNF195 | A0A087WT67 | 0.484 | 1.03 |
| GNB1 | P62873 | 0.83 | 1.02 |
| MRM1 | Q6IN84 | 0.695 | 1.02 |
| MTUS1 | Q9ULD2 | 0.223 | 1.02 |
| VRK3 | Q8IV63 | 0.367 | 1.02 |
| ARHGAP29 | Q52LW3 | 0.13 | 1.01 |
| CPNE1 | B0QZ18 | 0.653 | 1.01 |
| EIF4E | D6RBW1 | 0.774 | 1.01 |
| FBXW4 | P57775 | 0.69 | 1.01 |
| FRG1 | Q14331 | 0.519 | 1.01 |
| LRCH3 | Q96II8 | 0.142 | 1.01 |
| PUF60 | Q9UHX1 | 0.593 | 1.01 |
| TNRC6C | A0A1B0GU24 | 0.604 | 1.01 |
| TOMM40 | O96008 | 0.591 | 1.01 |
| VTN | P04004 | 0.203 | 1 |
| KCNN4 | O15554 | 0.484 | 0.998 |
| DVL2 | O14641 | 0.105 | 0.996 |

| | | | |
|---|---|---|---|
| DDX21 | Q9NR30 | 0.392 | 0.994 |
| PFKFB3 | A0A1W2PR17 | 0.559 | 0.994 |
| PGAM5 | Q96HS1 | 0.243 | 0.994 |
| PLCB1 | Q9NQ66 | 0.471 | 0.994 |
| MYO18A | Q92614 | 0.133 | 0.993 |
| MYO5C | Q9NQX4 | 0.106 | 0.993 |
| GOLGB1 | Q14789 | 0.439 | 0.992 |
| GTF3C6 | Q969F1 | 0.722 | 0.991 |
| PRRC2B | Q5JSZ5 | 0.576 | 0.991 |
| ZNF106 | Q9H2Y7 | 0.681 | 0.991 |
| MED14 | O60244 | 0.764 | 0.985 |
| STRIP1 | Q5VSL9 | 0.555 | 0.985 |
| FLG2 | Q5D862 | 0.629 | 0.984 |
| HDHD5 | Q9BXW7 | 0.74 | 0.98 |
| RPL9 | P32969 | 0.385 | 0.98 |
| ARHGEF18 | Q6ZSZ5 | 0.508 | 0.979 |
| PHLDB3 | Q6NSJ2 | 0.568 | 0.979 |
| LZTS2 | Q9BRK4 | 0.484 | 0.978 |
| MYL6 | B7Z6Z4 | 0.633 | 0.974 |
| ZNF121 | P58317 | 0.471 | 0.973 |
| KIF15 | Q9NS87 | 0.51 | 0.972 |
| PRKAA1 | Q13131 | 0.636 | 0.972 |
| SHKBP1 | Q8TBC3 | 0.343 | 0.971 |
| TCF12 | Q99081 | 0.735 | 0.971 |
| FASTKD3 | Q14CZ7 | 0.0946 | 0.97 |
| NHSL1 | Q5SYE7 | 0.695 | 0.968 |
| SMCHD1 | A6NHR9 | 0.206 | 0.967 |
| GPRC5C | A0A0C4DFY5 | 0.602 | 0.966 |
| RRP1 | P56182 | 0.623 | 0.966 |
| ZNF326 | Q5BKZ1 | 0.216 | 0.966 |
| AAK1.1 | A0A096LP25 | 0.481 | 0.962 |
| TNS2 | Q63HR2 | 0.09 | 0.962 |
| CLINT1 | Q14677 | 0.726 | 0.96 |
| SMN1 | E7EQZ4 | 0.406 | 0.96 |
| NAPA | P54920 | 0.465 | 0.959 |
| ZNF787 | Q6DD87 | 0.775 | 0.958 |
| PAPSS1 | O43252 | 0.737 | 0.953 |
| SORBS1.1 | A0A0U1RQI5 | 0.534 | 0.952 |
| PIK3C2A | O00443 | 0.171 | 0.951 |
| HPS6 | Q86YV9 | 0.37 | 0.95 |
| SNTB2 | Q13425 | 0.186 | 0.949 |
| URB2 | Q14146 | 0.787 | 0.948 |
| ZBTB5 | O15062 | 0.753 | 0.948 |
| ZNF57 | Q68EA5 | 0.524 | 0.947 |
| QPCTL | Q9NXS2 | 0.295 | 0.943 |
| RPS15A | P62244 | 0.371 | 0.942 |
| UTP23 | Q9BRU9 | 0.15 | 0.942 |

| | | | |
|---|---|---|---|
| COG8 | Q96MW5 | 0.781 | 0.941 |
| SPTBN1 | Q01082 | 0.226 | 0.938 |
| HIST2H2AA3 | Q6FI13 | 0.495 | 0.935 |
| TNRC6B.1 | H0Y720 | 0.304 | 0.935 |
| CFDP1 | Q9UEE9 | 0.601 | 0.934 |
| ORC5 | O43913 | 0.592 | 0.934 |
| MKI67 | P46013 | 0.691 | 0.93 |
| CDK7 | P50613 | 0.456 | 0.928 |
| BARD1 | Q99728 | 0.66 | 0.927 |
| DNAJB14 | Q8TBM8 | 0.358 | 0.927 |
| HIP1 | O00291 | 0.417 | 0.927 |
| CNOT3 | O75175 | 0.449 | 0.925 |
| HLA-B.1 | P30466 | 0.485 | 0.924 |
| PARP4 | Q9UKK3 | 0.236 | 0.924 |
| ZNF277 | Q9NRM2 | 0.532 | 0.924 |
| MYBL2 | P10244 | 0.566 | 0.923 |
| BICRA | Q9NZM4 | 0.812 | 0.917 |
| C6orf132 | Q5T0Z8 | 0.715 | 0.917 |
| CAV1 | Q03135 | 0.648 | 0.917 |
| CHD2 | O14647 | 0.698 | 0.917 |
| RPS24 | P62847 | 0.281 | 0.917 |
| STRN4 | Q9NRL3 | 0.407 | 0.917 |
| EFCAB7 | A8K855 | 0.717 | 0.916 |
| MED12 | Q93074 | 0.696 | 0.914 |
| ZNF525 | J3KR51 | 0.434 | 0.913 |
| CCT7 | Q99832 | 0.765 | 0.911 |
| ZYX | Q15942 | 0.748 | 0.911 |
| KRIT1 | O00522 | 0.787 | 0.91 |
| PDS5B.1 | A9IYQ1 | 0.474 | 0.91 |
| MPHOSPH8 | Q99549 | 0.79 | 0.909 |
| ERBIN | Q96RT1 | 0.533 | 0.907 |
| DDX51 | Q8N8A6 | 0.782 | 0.903 |
| TANC1 | Q9C0D5 | 0.73 | 0.903 |
| IFIT5 | Q13325 | 0.603 | 0.902 |
| GLE1 | Q53GS7 | 0.501 | 0.899 |
| CEP170 | Q5SW79 | 0.643 | 0.898 |
| FKBP3 | Q00688 | 0.684 | 0.896 |
| RBM12B | Q8IXT5 | 0.332 | 0.896 |
| UBN1 | Q9NPG3 | 0.709 | 0.896 |
| PPP1R12A | O14974 | 0.13 | 0.895 |
| PCM1.1 | E7EV93 | 0.356 | 0.891 |
| COG7 | P83436 | 0.735 | 0.889 |
| USP42 | Q9H9J4 | 0.787 | 0.887 |
| DLG5 | Q8TDM6 | 0.694 | 0.885 |
| POLR1E.1 | E7EX70 | 0.649 | 0.885 |
| RC3H1 | Q5TC82 | 0.305 | 0.885 |
| VPS51 | Q9UID3 | 0.751 | 0.88 |

| | | | |
|---|---|---|---|
| VPS39 | Q96JC1 | 0.51 | 0.877 |
| ADNP2 | Q6IQ32 | 0.775 | 0.874 |
| GNG12 | Q9UBI6 | 0.788 | 0.874 |
| HMG20B | Q9P0W2 | 0.53 | 0.871 |
| HERC5 | Q9UII4 | 0.434 | 0.865 |
| OCLN | A0A0G2JMZ8 | 0.777 | 0.863 |
| ZC3H18 | E7ERS3 | 0.667 | 0.862 |
| H2AFX | P16104 | 0.437 | 0.86 |
| ARHGEF5 | Q12774 | 0.211 | 0.858 |
| SON | P18583 | 0.711 | 0.858 |
| CABIN1 | Q9Y6J0 | 0.607 | 0.857 |
| DDX28 | Q9NUL7 | 0.45 | 0.855 |
| SPICE1 | Q8N0Z3 | 0.258 | 0.854 |
| S100A6 | P06703 | 0.778 | 0.851 |
| EDRF1 | Q3B7T1 | 0.698 | 0.85 |
| CAMSAP1 | Q5T5Y3 | 0.612 | 0.849 |
| CMTR2 | A0A0U1RRB3 | 0.644 | 0.849 |
| CSNK1A1 | P48729 | 0.262 | 0.849 |
| TPR | P12270 | 0.422 | 0.849 |
| RPS17 | P08708 | 0.372 | 0.848 |
| RPS27 | P42677 | 0.498 | 0.848 |
| CCDC186 | Q7Z3E2 | 0.54 | 0.847 |
| RPL11 | P62913 | 0.559 | 0.845 |
| CTTNBP2NL | Q9P2B4 | 0.793 | 0.842 |
| FLII | Q13045 | 0.162 | 0.841 |
| GNA13 | Q14344 | 0.783 | 0.84 |
| HAUS6 | Q7Z4H7 | 0.138 | 0.84 |
| MAGEB2 | O15479 | 0.222 | 0.839 |
| FARP1.1 | A0A1B0GV68 | 0.567 | 0.838 |
| SSR4 | P51571 | 0.62 | 0.838 |
| CCT6A | P40227 | 0.759 | 0.837 |
| CCT8 | P50990 | 0.754 | 0.836 |
| CAPZA2 | P47755 | 0.654 | 0.832 |
| TAP1 | A0A140T9T7 | 0.627 | 0.831 |
| SREBF1 | P36956 | 0.695 | 0.83 |
| DOT1L | Q8TEK3 | 0.739 | 0.829 |
| HNRNPDL | O14979 | 0.589 | 0.829 |
| RPL31 | B7Z4C8 | 0.375 | 0.827 |
| SPEN | Q96T58 | 0.711 | 0.827 |
| ING1 | A0A0C4DFW2 | 0.75 | 0.826 |
| SENP5 | Q96HI0 | 0.679 | 0.822 |
| ARHGAP23 | Q9P227 | 0.777 | 0.821 |
| MEN1 | O00255 | 0.747 | 0.821 |
| UBAP1 | A0A0D9SG79 | 0.695 | 0.821 |
| DUS1L | Q6P1R4 | 0.666 | 0.819 |
| PPP2CA | P67775 | 0.671 | 0.819 |
| NSMCE3 | Q96MG7 | 0.502 | 0.817 |

| | | | |
|---|---|---|---|
| DNMBP | Q6XZF7 | 0.196 | 0.813 |
| SLC38A2 | Q96QD8 | 0.684 | 0.811 |
| SMG8 | Q8ND04 | 0.403 | 0.811 |
| RIPK1 | Q13546 | 0.768 | 0.809 |
| ATRX | P46100 | 0.428 | 0.807 |
| ZPR1 | O75312 | 0.754 | 0.806 |
| ACIN1 | Q9UKV3 | 0.49 | 0.804 |
| E2F4 | Q16254 | 0.713 | 0.804 |
| FNBP1L | S4R347 | 0.572 | 0.804 |
| RAD17 | A0A0G2JPT5 | 0.711 | 0.803 |
| EPN2 | O95208 | 0.768 | 0.802 |
| NAXE | Q8NCW5 | 0.635 | 0.801 |
| CASK | O14936 | 0.538 | 0.8 |
| HIVEP2 | P31629 | 0.733 | 0.795 |
| IRS2 | Q9Y4H2 | 0.784 | 0.794 |
| GTPBP4 | Q9BZE4 | 0.451 | 0.792 |
| ZMYM3 | A6NHB5 | 0.781 | 0.792 |
| DNAJC13 | O75165 | 0.204 | 0.791 |
| APC | P25054 | 0.56 | 0.79 |
| CCT5 | P48643 | 0.776 | 0.787 |
| LUZP1 | Q86V48 | 0.655 | 0.785 |
| CEP85 | Q6P2H3 | 0.619 | 0.784 |
| ZBTB7A | O95365 | 0.806 | 0.782 |
| DUSP11 | O75319 | 0.483 | 0.781 |
| WFS1 | O76024 | 0.676 | 0.778 |
| MRPS35 | P82673 | 0.58 | 0.776 |
| PTCD3 | Q96EY7 | 0.538 | 0.776 |
| TPM4 | P67936 | 0.787 | 0.775 |
| DCTN4 | Q9UJW0 | 0.686 | 0.773 |
| TCP1 | P17987 | 0.788 | 0.773 |
| NUMA1.1 | A0A087WY61 | 0.833 | 0.772 |
| SH3GL1 | Q99961 | 0.693 | 0.772 |
| DNAJC8 | O75937 | 0.337 | 0.771 |
| RPS27L | H0YMV8 | 0.422 | 0.77 |
| GRB2 | P62993 | 0.539 | 0.767 |
| PSMD4 | P55036 | 0.674 | 0.767 |
| FGFRL1 | Q8N441 | 0.802 | 0.765 |
| RTCB | Q9Y3I0 | 0.448 | 0.765 |
| RRAS | P10301 | 0.683 | 0.763 |
| ANAPC2 | Q9UJX6 | 0.672 | 0.761 |
| ATP6V1C1 | P21283 | 0.803 | 0.761 |
| GPRIN1 | Q7Z2K8 | 0.785 | 0.761 |
| NOP58 | Q9Y2X3 | 0.787 | 0.761 |
| UTRN | P46939 | 0.553 | 0.761 |
| NAB2 | Q15742 | 0.264 | 0.76 |
| TAF1B | Q53T94 | 0.689 | 0.76 |
| FAM117B | Q6P1L5 | 0.742 | 0.758 |

| | | | |
|---|---|---|---|
| RPS11 | P62280 | 0.649 | 0.757 |
| UBR4 | Q5T4S7 | 0.641 | 0.757 |
| MIER1 | Q8N108 | 0.74 | 0.755 |
| NEPRO | Q6NW34 | 0.794 | 0.755 |
| PLEKHN1 | Q494U1 | 0.556 | 0.755 |
| FKBP15 | A0A0A0MT60 | 0.764 | 0.754 |
| NUP35 | C9IYQ7 | 0.412 | 0.754 |
| NUP160 | Q12769 | 0.475 | 0.753 |
| RPA2 | P15927 | 0.711 | 0.753 |
| PEX5 | B4E0T2 | 0.708 | 0.751 |
| TOP2A | P11388 | 0.597 | 0.751 |
| SEL1L3 | Q68CR1 | 0.763 | 0.75 |
| YWHAG | P61981 | 0.816 | 0.749 |
| BUB1 | O43683 | 0.614 | 0.748 |
| NECTIN2 | Q92692 | 0.802 | 0.748 |
| ZC2HC1A | Q96GY0 | 0.724 | 0.748 |
| ARRB1.1 | H0YET9 | 0.611 | 0.747 |
| TBCB | Q99426 | 0.818 | 0.747 |
| CAV2 | P51636 | 0.737 | 0.746 |
| NEURL4 | Q96JN8 | 0.745 | 0.746 |
| BLVRB | P30043 | 0.555 | 0.744 |
| KIAA0753 | Q2KHM9 | 0.697 | 0.742 |
| MINK1 | Q8N4C8 | 0.742 | 0.742 |
| RPL35 | P42766 | 0.231 | 0.741 |
| UNK | Q9C0B0 | 0.373 | 0.741 |
| NDUFS1 | P28331 | 0.752 | 0.74 |
| WDR76 | Q9H967 | 0.793 | 0.74 |
| E9PLN8 | E9PLN8 | 0.679 | 0.737 |
| VMP1 | Q96GC9 | 0.767 | 0.737 |
| BRCA1 | P38398 | 0.775 | 0.736 |
| MFN2 | O95140 | 0.836 | 0.736 |
| SET | Q01105 | 0.709 | 0.735 |
| MYH14.1 | M0QY43 | 0.371 | 0.734 |
| PCGF6 | Q9BYE7 | 0.679 | 0.733 |
| PSAT1 | Q9Y617 | 0.845 | 0.732 |
| ROCK2 | O75116 | 0.729 | 0.732 |
| MVP | Q14764 | 0.705 | 0.731 |
| CTTN | Q14247 | 0.78 | 0.73 |
| SCAPER | Q9BY12 | 0.64 | 0.73 |
| RPL17-C18orf32 | A0A0A6YYL6 | 0.358 | 0.729 |
| CRTC3 | Q6UUV7 | 0.79 | 0.727 |
| PRDM15 | P57071 | 0.792 | 0.725 |
| TTC3 | P53804 | 0.651 | 0.724 |
| GMPR2 | H0YNJ6 | 0.56 | 0.723 |
| PLEKHA7.1 | E9PKC0 | 0.761 | 0.722 |
| CCT3 | P49368 | 0.773 | 0.721 |
| ROCK1 | Q13464 | 0.608 | 0.721 |

| | | | |
|---|---|---|---|
| MRTO4 | Q9UKD2 | 0.514 | 0.718 |
| RAPGEF6 | Q8TEU7 | 0.717 | 0.718 |
| HMGN2 | P05204 | 0.486 | 0.717 |
| RPL23 | P62829 | 0.327 | 0.717 |
| TAOK3 | Q9H2K8 | 0.781 | 0.715 |
| CDC42EP4 | Q9H3Q1 | 0.389 | 0.712 |
| GEN1 | Q17RS7 | 0.825 | 0.711 |
| NAA30 | Q147X3 | 0.291 | 0.709 |
| BRAP | Q7Z569 | 0.637 | 0.707 |
| PDLIM7 | Q9NR12 | 0.584 | 0.705 |
| DNAJA4.1 | F5H300 | 0.597 | 0.704 |
| TRA2A | Q13595 | 0.633 | 0.704 |
| EPN1 | Q9Y6I3 | 0.788 | 0.702 |
| MROH6 | A6NGR9 | 0.738 | 0.702 |
| RPS9 | A0A024R4M0 | 0.712 | 0.699 |
| SLTM | Q9NWH9 | 0.798 | 0.697 |
| RPS14 | P62263 | 0.679 | 0.695 |
| DOCK9 | A0A088AWN3 | 0.723 | 0.694 |
| DDX50 | Q9BQ39 | 0.786 | 0.693 |
| TMF1 | P82094 | 0.558 | 0.693 |
| CDC5L | Q99459 | 0.78 | 0.692 |
| MRPS31 | Q92665 | 0.426 | 0.692 |
| EME1 | Q96AY2 | 0.721 | 0.691 |
| HERC2 | O95714 | 0.592 | 0.691 |
| NME1 | P15531 | 0.827 | 0.688 |
| DCAF13 | A0A087WT20 | 0.623 | 0.687 |
| PLS1 | Q14651 | 0.847 | 0.687 |
| PLEKHG6 | Q3KR16 | 0.722 | 0.686 |
| SH3BP4 | Q9P0V3 | 0.683 | 0.685 |
| CA2 | P00918 | 0.74 | 0.684 |
| CCT4 | P50991 | 0.788 | 0.684 |
| RPS13 | P62277 | 0.573 | 0.682 |
| ATAD5 | Q96QE3 | 0.473 | 0.681 |
| NEK9 | Q8TD19 | 0.704 | 0.681 |
| TCTN3 | A0A0C4DFN5 | 0.579 | 0.681 |
| NLE1 | Q9NVX2 | 0.71 | 0.678 |
| RPL37A | P61513 | 0.421 | 0.678 |
| MYO5A | F8W6H6 | 0.704 | 0.677 |
| RPL35A | P18077 | 0.777 | 0.676 |
| A0A0B4J2A0 | A0A0B4J2A0 | 0.778 | 0.672 |
| A0A1W2PNV4 | A0A1W2PNV4 | 0.799 | 0.671 |
| ARFGAP2 | Q8N6H7 | 0.814 | 0.671 |
| EIF2AK4 | Q9P2K8 | 0.762 | 0.671 |
| GSTO1 | P78417 | 0.824 | 0.669 |
| SYTL1 | Q8IYJ3 | 0.657 | 0.668 |
| ZFP64 | Q9NPA5 | 0.765 | 0.668 |
| ADAR | P55265 | 0.359 | 0.666 |

| | | | |
|---|---|---|---|
| RASA2 | A0A0A0MSJ9 | 0.83 | 0.663 |
| CALCOCO2 | Q13137 | 0.783 | 0.661 |
| RPS10 | P46783 | 0.624 | 0.661 |
| SACM1L | Q9NTJ5 | 0.83 | 0.659 |
| TJP1 | G3V1L9 | 0.723 | 0.659 |
| ARHGEF12 | Q9NZN5 | 0.692 | 0.657 |
| RPL36 | Q9Y3U8 | 0.564 | 0.657 |
| RPS25 | P62851 | 0.757 | 0.657 |
| PNKD | Q8N490 | 0.701 | 0.656 |
| RSL1D1 | O76021 | 0.765 | 0.655 |
| CCT2 | P78371 | 0.813 | 0.653 |
| CUX1 | P39880 | 0.583 | 0.653 |
| FLII.1 | J3KS54 | 0.584 | 0.653 |
| KAT5 | Q92993 | 0.802 | 0.653 |
| MAP7D1 | Q3KQU3 | 0.756 | 0.653 |
| GNAI3 | P08754 | 0.859 | 0.652 |
| DYRK1A | Q13627 | 0.708 | 0.648 |
| hCG_1984214 | I3L0E3 | 0.685 | 0.648 |
| IRS1 | P35568 | 0.828 | 0.648 |
| HMMR | O75330 | 0.763 | 0.647 |
| RPS7 | P62081 | 0.542 | 0.647 |
| POLR2A | P24928 | 0.781 | 0.644 |
| SYTL2 | A0A0U1RR07 | 0.449 | 0.644 |
| CACNA2D2 | C9JVC9 | 0.74 | 0.643 |
| RPS8 | P62241 | 0.783 | 0.643 |
| PPP1CC | F8W0W8 | 0.444 | 0.64 |
| CFL1 | E9PK25 | 0.82 | 0.639 |
| DDX27 | Q96GQ7 | 0.796 | 0.639 |
| EPS15 | P42566 | 0.434 | 0.639 |
| ASF1A | Q9Y294 | 0.789 | 0.638 |
| SH3PXD2B | A1X283 | 0.731 | 0.638 |
| MRPL44 | Q9H9J2 | 0.5 | 0.637 |
| RPS19 | P39019 | 0.74 | 0.637 |
| RPS23 | P62266 | 0.696 | 0.637 |
| NOP53 | Q9NZM5 | 0.771 | 0.636 |
| ASCC1 | Q8N9N2 | 0.773 | 0.635 |
| CNOT10 | Q9H9A5 | 0.469 | 0.635 |
| IGF2BP2 | F8W930 | 0.797 | 0.635 |
| METTL17 | Q9H7H0 | 0.719 | 0.635 |
| ATAD3B | Q5T9A4 | 0.624 | 0.634 |
| INO80B-WBP1 | J3KQ70 | 0.748 | 0.634 |
| RPS6 | P62753 | 0.739 | 0.634 |
| USP11 | P51784 | 0.775 | 0.634 |
| PLEKHA5 | Q9HAU0 | 0.749 | 0.633 |
| TAF3 | Q5VWG9 | 0.849 | 0.633 |
| RPL26 | P61254 | 0.403 | 0.632 |
| TG | P01266 | 0.552 | 0.632 |

| | | | |
|---|---|---|---|
| MADD | A0A0A0MRB5 | 0.489 | 0.629 |
| TAX1BP1 | B8ZZD4 | 0.615 | 0.629 |
| AXIN1 | O15169 | 0.821 | 0.627 |
| LAS1L | Q9Y4W2 | 0.821 | 0.627 |
| ILF2 | Q12905 | 0.668 | 0.626 |
| KAT8 | Q9H7Z6 | 0.801 | 0.626 |
| SLC25A3 | Q00325 | 0.768 | 0.626 |
| ARHGEF7.1 | Q14155 | 0.757 | 0.625 |
| RPL13 | P26373 | 0.777 | 0.62 |
| POLB | P06746 | 0.839 | 0.619 |
| SLC12A7 | A0A0G2JNW7 | 0.764 | 0.619 |
| TBC1D10A | Q9BXI6 | 0.805 | 0.618 |
| CDC16 | Q13042 | 0.512 | 0.617 |
| KIAA2013 | Q8IYS2 | 0.796 | 0.617 |
| PHF20 | Q9BVI0 | 0.837 | 0.616 |
| POLL | Q9UGP5 | 0.627 | 0.616 |
| WDR59 | Q6PJI9 | 0.579 | 0.616 |
| PDXDC1 | H3BND4 | 0.83 | 0.613 |
| DNAJB6 | O75190 | 0.787 | 0.612 |
| SLU7 | O95391 | 0.808 | 0.612 |
| ADRM1 | Q16186 | 0.819 | 0.611 |
| BCR | P11274 | 0.516 | 0.61 |
| DNAJA4 | Q8WW22 | 0.568 | 0.61 |
| ERCC5 | P28715 | 0.846 | 0.61 |
| KLF13 | Q9Y2Y9 | 0.703 | 0.61 |
| RBM42 | Q9BTD8 | 0.796 | 0.61 |
| SEC23IP | Q9Y6Y8 | 0.789 | 0.609 |
| NUCB1 | Q02818 | 0.813 | 0.608 |
| C18orf25 | Q96B23 | 0.684 | 0.607 |
| IMPA1 | P29218 | 0.811 | 0.605 |
| MED26 | O95402 | 0.845 | 0.605 |
| TJP1.1 | G5E9E7 | 0.758 | 0.604 |
| PSMC6 | A0A087X2I1 | 0.824 | 0.603 |
| SNAPC2 | Q13487 | 0.821 | 0.603 |
| ARFGEF2 | Q9Y6D5 | 0.672 | 0.602 |
| MRPS5 | P82675 | 0.779 | 0.602 |
| RPL36A | J3KQN4 | 0.47 | 0.602 |
| TSPYL2 | Q9H2G4 | 0.841 | 0.602 |
| TANC2 | Q9HCD6 | 0.773 | 0.601 |
| EPPK1 | P58107 | 0.778 | 0.6 |
| LMNA.1 | Q3BDU5 | 0.801 | 0.599 |
| ZC3H13 | Q5T200 | 0.816 | 0.599 |
| RBM4 | Q9BWF3 | 0.816 | 0.598 |
| RPS4X | P62701 | 0.767 | 0.598 |
| VPS45 | Q9NRW7 | 0.784 | 0.598 |
| TCF7L2 | E2GH26 | 0.781 | 0.597 |
| SLC35E1 | Q96K37 | 0.727 | 0.596 |

| | | | |
|---|---|---|---|
| TGS1 | Q96RS0 | 0.799 | 0.596 |
| CHUK | O15111 | 0.812 | 0.595 |
| RAB3GAP2 | Q9H2M9 | 0.79 | 0.595 |
| 2-Sep | Q15019 | 0.766 | 0.595 |
| THYN1 | Q9P016 | 0.73 | 0.595 |
| NUFIP2 | Q7Z417 | 0.559 | 0.594 |
| SUMO3 | A8MU27 | 0.565 | 0.593 |
| DSC3 | Q14574 | 0.81 | 0.592 |
| ZNF329 | Q86UD4 | 0.635 | 0.592 |
| GDI1 | P31150 | 0.834 | 0.591 |
| TAF2 | Q6P1X5 | 0.796 | 0.59 |
| PSME1 | Q06323 | 0.823 | 0.589 |
| CMTR1 | Q8N1G2 | 0.736 | 0.588 |
| DDI2 | Q5TDH0 | 0.765 | 0.588 |
| PFN1 | P07737 | 0.836 | 0.588 |
| RAPGEF2 | Q9Y4G8 | 0.793 | 0.587 |
| HTRA2 | O43464 | 0.755 | 0.586 |
| SLC38A1 | F8VX04 | 0.828 | 0.586 |
| COG5 | Q9UP83 | 0.84 | 0.585 |
| MECP2 | P51608 | 0.787 | 0.585 |
| BBX | Q8WY36 | 0.847 | 0.584 |
| RPL24 | P83731 | 0.58 | 0.584 |
| MED13 | Q9UHV7 | 0.832 | 0.581 |
| NKTR | P30414 | 0.826 | 0.579 |
| EIF4E2 | O60573 | 0.815 | 0.578 |
| TBC1D24 | Q9ULP9 | 0.795 | 0.577 |
| TAF8 | Q7Z7C8 | 0.501 | 0.576 |
| RAB11FIP5 | A0A1B0GTL5 | 0.799 | 0.575 |
| DHX40 | Q8IX18 | 0.589 | 0.574 |
| ECHDC1 | Q9NTX5 | 0.811 | 0.574 |
| MON2 | A0A286YFF8 | 0.656 | 0.574 |
| C12orf43 | F5H7W8 | 0.818 | 0.573 |
| CCDC77 | Q9BR77 | 0.739 | 0.572 |
| TAF5L | O75529 | 0.785 | 0.572 |
| NUP107 | P57740 | 0.732 | 0.571 |
| RINT1 | Q6NUQ1 | 0.821 | 0.571 |
| DDX6 | P26196 | 0.608 | 0.569 |
| MYO6 | A0A0A0MRM8 | 0.513 | 0.569 |
| HNRNPC | G3V4C1 | 0.687 | 0.567 |
| FIZ1 | Q96SL8 | 0.836 | 0.566 |
| MED1 | Q15648 | 0.819 | 0.566 |
| TUBGCP2 | Q9BSJ2 | 0.717 | 0.566 |
| RPS3A | P61247 | 0.75 | 0.565 |
| SAMHD1 | Q9Y3Z3 | 0.75 | 0.565 |
| YTHDC2 | Q9H6S0 | 0.515 | 0.563 |
| ATF6 | P18850 | 0.766 | 0.561 |
| PISD | Q9UG56 | 0.67 | 0.56 |

| | | | |
|---|---|---|---|
| RPS16 | P62249 | 0.844 | 0.56 |
| PABPC4.1 | H0Y5F5 | 0.811 | 0.559 |
| RPL29 | P47914 | 0.766 | 0.559 |
| ADAM10 | O14672 | 0.844 | 0.558 |
| CHAMP1 | Q96JM3 | 0.833 | 0.558 |
| ICE2 | Q659A1 | 0.812 | 0.558 |
| RPL38 | P63173 | 0.585 | 0.557 |
| DYNC1LI1 | Q9Y6G9 | 0.818 | 0.556 |
| H2AFY | O75367 | 0.632 | 0.556 |
| 1-Mar | Q5VT66 | 0.754 | 0.556 |
| UACA | Q9BZF9 | 0.808 | 0.555 |
| ATAD3A | H0Y2W2 | 0.682 | 0.554 |
| SQLE | Q14534 | 0.657 | 0.554 |
| MERTK | Q12866 | 0.613 | 0.553 |
| MAP4 | E7EVA0 | 0.547 | 0.552 |
| ELAVL1 | Q15717 | 0.788 | 0.55 |
| GON4L | Q3T8J9 | 0.834 | 0.549 |
| DSP | P15924 | 0.562 | 0.548 |
| FAM111A | Q96PZ2 | 0.76 | 0.548 |
| MAPKAPK5 | Q8IW41 | 0.795 | 0.548 |
| PMPCB | G3V0E4 | 0.832 | 0.548 |
| VAC14 | Q08AM6 | 0.825 | 0.545 |
| XRCC6 | P12956 | 0.755 | 0.544 |
| CAMSAP2 | Q08AD1 | 0.831 | 0.543 |
| BMP4 | P12644 | 0.741 | 0.542 |
| MRPS22 | P82650 | 0.582 | 0.542 |
| RPL10 | P27635 | 0.672 | 0.542 |
| NFS1 | Q9Y697 | 0.738 | 0.541 |
| SLC25A4 | P12235 | 0.788 | 0.541 |
| PLRG1 | O43660 | 0.796 | 0.54 |
| USP34 | Q70CQ2 | 0.759 | 0.54 |
| ZNF205 | O95201 | 0.841 | 0.54 |
| OTUD4 | Q01804 | 0.567 | 0.538 |
| TRIOBP | Q9H2D6 | 0.733 | 0.538 |
| CAVIN3 | E9PIE3 | 0.696 | 0.537 |
| SPAST | Q9UBP0 | 0.657 | 0.537 |
| ASAP2 | O43150 | 0.832 | 0.535 |
| ATP6V1A | P38606 | 0.778 | 0.535 |
| COG3 | Q96JB2 | 0.79 | 0.535 |
| POLR3C | Q9BUI4 | 0.659 | 0.535 |
| EP400 | Q96L91 | 0.854 | 0.534 |
| SCAI | Q8N9R8 | 0.801 | 0.533 |
| FAM114A1 | Q8IWE2 | 0.833 | 0.532 |
| PIK3R4 | Q99570 | 0.622 | 0.532 |
| RPL27A | P46776 | 0.53 | 0.532 |
| PRPS1 | P60891 | 0.839 | 0.531 |
| RSRC2 | Q7L4I2 | 0.803 | 0.531 |

| | | | |
|---|---|---|---|
| ZNF598 | Q86UK7 | 0.711 | 0.53 |
| SUGP2 | M0R2Z9 | 0.798 | 0.529 |
| PPP1R12C | Q9BZL4 | 0.77 | 0.528 |
| ZBTB48 | P10074 | 0.717 | 0.528 |
| EXOC5 | A0A0A0MSI8 | 0.789 | 0.527 |
| RALGAPB | Q86X10 | 0.83 | 0.527 |
| SHC1 | P29353 | 0.787 | 0.525 |
| GNAI2 | P04899 | 0.854 | 0.524 |
| CORO1C | Q9ULV4 | 0.809 | 0.522 |
| ACSL5 | Q9ULC5 | 0.683 | 0.521 |
| BRSK2 | Q8IWQ3 | 0.778 | 0.521 |
| DCXR | Q7Z4W1 | 0.845 | 0.52 |
| ZBTB43 | O43298 | 0.831 | 0.52 |
| IQSEC2 | Q5JU85 | 0.826 | 0.519 |
| LIG3 | P49916 | 0.713 | 0.519 |
| RAB14 | P61106 | 0.571 | 0.519 |
| ZBTB1 | Q9Y2K1 | 0.84 | 0.516 |
| DNA2 | P51530 | 0.843 | 0.515 |
| L3MBTL2 | Q969R5 | 0.642 | 0.515 |
| MAP1B | P46821 | 0.663 | 0.515 |
| THRAP3 | Q9Y2W1 | 0.545 | 0.514 |
| RPL19 | P84098 | 0.679 | 0.512 |
| USP22 | Q9UPT9 | 0.836 | 0.511 |
| FAM161A | Q3B820 | 0.813 | 0.51 |
| POLQ | A0A087WXL3 | 0.792 | 0.51 |
| SMS | P52788 | 0.843 | 0.508 |
| MCM3AP | O60318 | 0.829 | 0.505 |
| MCM10 | Q7L590 | 0.675 | 0.503 |
| PABPC4 | Q13310 | 0.786 | 0.503 |
| PPP1CB | P62140 | 0.596 | 0.503 |
| OSBPL8 | Q9BZF1 | 0.814 | 0.502 |
| RPL13A | M0QYS1 | 0.856 | 0.501 |
| STK3 | Q13188 | 0.706 | 0.499 |
| FLNB | O75369 | 0.773 | 0.497 |
| NT5DC2 | Q9H857 | 0.752 | 0.497 |
| PDCD11 | Q14690 | 0.86 | 0.497 |
| AFDN | J3KN01 | 0.811 | 0.496 |
| KIF18B.1 | A0A0C4DGP2 | 0.809 | 0.495 |
| AMFR | Q9UKV5 | 0.791 | 0.494 |
| HELZ | P42694 | 0.847 | 0.492 |
| RICTOR | Q6R327 | 0.829 | 0.491 |
| UPF2 | Q9HAU5 | 0.844 | 0.491 |
| MED16 | Q9Y2X0 | 0.788 | 0.49 |
| RPS26 | P62854 | 0.753 | 0.49 |
| ATP5F1 | P24539 | 0.864 | 0.489 |
| FAM160B1 | Q5W0V3 | 0.835 | 0.489 |
| FAM83D | A0A087WXK8 | 0.774 | 0.489 |

| | | | |
|---|---|---|---|
| PRDX1 | Q06830 | 0.848 | 0.489 |
| EXOC4 | Q96A65 | 0.78 | 0.487 |
| NXN | Q6DKJ4 | 0.852 | 0.487 |
| KDM4B | F5GX28 | 0.847 | 0.484 |
| UTP3 | Q9NQZ2 | 0.837 | 0.484 |
| MICAL2 | O94851 | 0.843 | 0.483 |
| MRPS2 | Q9Y399 | 0.719 | 0.483 |
| SNRPB | P14678 | 0.717 | 0.483 |
| WTAP | Q15007 | 0.826 | 0.483 |
| MCC | P23508 | 0.841 | 0.482 |
| GOLGA4 | Q13439 | 0.793 | 0.481 |
| ANXA2 | P07355 | 0.704 | 0.479 |
| DNM1L.1 | G8JLD5 | 0.811 | 0.479 |
| PTPN23 | Q9H3S7 | 0.791 | 0.479 |
| NSUN5 | Q96P11 | 0.794 | 0.478 |
| CTNNB1 | P35222 | 0.785 | 0.477 |
| ILK | A0A0A0MTH3 | 0.693 | 0.477 |
| MRPS7 | J3QLS3 | 0.751 | 0.476 |
| PPP2R2A | P63151 | 0.799 | 0.476 |
| EPB41L1.3 | X6RC15 | 0.786 | 0.475 |
| IGF2BP3 | O00425 | 0.815 | 0.475 |
| PLIN3 | O60664 | 0.834 | 0.475 |
| ZNF746 | Q6NUN9 | 0.843 | 0.475 |
| ARRDC1 | Q8N5I2 | 0.844 | 0.474 |
| IFT172 | Q9UG01 | 0.754 | 0.474 |
| PLOD3 | O60568 | 0.745 | 0.473 |
| ATP6V0A2 | Q9Y487 | 0.643 | 0.471 |
| CSNK1D | H7BYT1 | 0.685 | 0.471 |
| GTF2E2 | P29084 | 0.735 | 0.471 |
| RAE1 | P78406 | 0.749 | 0.471 |
| EHBP1 | Q8NDI1 | 0.859 | 0.47 |
| NR4A3 | Q92570 | 0.705 | 0.47 |
| PXK | Q7Z7A4 | 0.826 | 0.47 |
| KLC2 | Q9H0B6 | 0.85 | 0.469 |
| DHX29 | A0A087WYN9 | 0.658 | 0.468 |
| RPL30 | P62888 | 0.828 | 0.468 |
| SLC9A3R1 | O14745 | 0.763 | 0.466 |
| CRNKL1 | Q9BZJ0 | 0.728 | 0.464 |
| RBBP6 | Q7Z6E9 | 0.861 | 0.464 |
| FAM83G | A6ND36 | 0.68 | 0.463 |
| PSMA2 | P25787 | 0.857 | 0.463 |
| TM9SF3 | Q9HD45 | 0.82 | 0.463 |
| CDK2 | G3V5T9 | 0.773 | 0.462 |
| LRBA | P50851 | 0.785 | 0.461 |
| RFWD3 | Q6PCD5 | 0.695 | 0.461 |
| RPL28 | P46779 | 0.837 | 0.46 |
| GPS1 | Q13098 | 0.849 | 0.458 |

| | | | |
|---|---|---|---|
| NFYA | P23511 | 0.841 | 0.458 |
| PPP1CA | P62136 | 0.663 | 0.458 |
| TRAPPC9 | Q96Q05 | 0.764 | 0.458 |
| MICAL3 | Q7RTP6 | 0.727 | 0.457 |
| MRPL4 | Q9BYD3 | 0.813 | 0.457 |
| RPS15 | K7ELC2 | 0.837 | 0.456 |
| ANKLE2 | Q86XL3 | 0.773 | 0.455 |
| HNRNPA0 | Q13151 | 0.818 | 0.455 |
| HRNR | Q86YZ3 | 0.845 | 0.455 |
| PLCG2 | P16885 | 0.817 | 0.454 |
| RPL23A | P62750 | 0.681 | 0.454 |
| THAP4 | Q8WY91 | 0.826 | 0.454 |
| PAK6 | Q9NQU5 | 0.78 | 0.452 |
| ZNF384 | Q8TF68 | 0.745 | 0.452 |
| ACADSB | P45954 | 0.845 | 0.451 |
| EIF3A | Q14152 | 0.792 | 0.451 |
| PPP4R1 | Q8TF05 | 0.841 | 0.451 |
| RBM14 | Q96PK6 | 0.757 | 0.45 |
| SNX8 | Q9Y5X2 | 0.752 | 0.45 |
| EHBP1L1 | Q8N3D4 | 0.783 | 0.449 |
| WDR82 | Q6UXN9 | 0.74 | 0.449 |
| PNO1 | Q9NRX1 | 0.806 | 0.448 |
| TUBGCP3 | Q96CW5 | 0.802 | 0.448 |
| UPF1 | Q92900 | 0.677 | 0.448 |
| RPL6 | Q02878 | 0.831 | 0.447 |
| CD109 | Q6YHK3 | 0.822 | 0.445 |
| CLPTM1L | Q96KA5 | 0.688 | 0.445 |
| CSK | P41240 | 0.857 | 0.445 |
| PRKAG1 | P54619 | 0.809 | 0.444 |
| RPA1 | P27694 | 0.734 | 0.443 |
| TMED7 | Q9Y3B3 | 0.845 | 0.442 |
| VASP | P50552 | 0.831 | 0.441 |
| AURKB | Q96GD4 | 0.705 | 0.439 |
| ATXN2L | Q8WWM7 | 0.737 | 0.438 |
| LYN | P07948 | 0.86 | 0.438 |
| GNB2 | P62879 | 0.869 | 0.437 |
| MXRA5 | Q9NR99 | 0.863 | 0.437 |
| ARHGAP35 | Q9NRY4 | 0.784 | 0.436 |
| CAPZA1 | P52907 | 0.814 | 0.435 |
| FAM83B | Q5T0W9 | 0.778 | 0.435 |
| NUP214 | P35658 | 0.724 | 0.435 |
| USP31 | Q70CQ4 | 0.819 | 0.435 |
| KHSRP | Q92945 | 0.797 | 0.434 |
| KIF1C | O43896 | 0.763 | 0.434 |
| LIG4 | P49917 | 0.794 | 0.434 |
| SOGA1 | O94964 | 0.851 | 0.434 |
| STAT3 | P40763 | 0.672 | 0.433 |

| | | | |
|---|---|---|---|
| TEDC1 | Q86SX3 | 0.816 | 0.433 |
| PRKAR1A | P10644 | 0.845 | 0.432 |
| ARAF | Q96II5 | 0.768 | 0.43 |
| DNAJA3 | Q96EY1 | 0.756 | 0.43 |
| HSPB1 | P04792 | 0.85 | 0.43 |
| RPL8 | P62917 | 0.838 | 0.43 |
| YWHAB | P31946 | 0.873 | 0.427 |
| HP1BP3 | Q5SSJ5 | 0.859 | 0.426 |
| SMARCC2 | Q8TAQ2 | 0.86 | 0.426 |
| NXF1 | Q9UBU9 | 0.793 | 0.425 |
| 8-Sep | Q92599 | 0.831 | 0.425 |
| BAG2 | O95816 | 0.834 | 0.424 |
| HNRNPUL2 | Q1KMD3 | 0.727 | 0.423 |
| CSTF1 | Q05048 | 0.812 | 0.42 |
| HNRNPA2B1 | P22626 | 0.813 | 0.42 |
| ZFC3H1 | O60293 | 0.853 | 0.42 |
| ATXN7L1 | Q9ULK2 | 0.828 | 0.419 |
| GNL3 | Q9BVP2 | 0.823 | 0.418 |
| UBXN4 | Q92575 | 0.84 | 0.418 |
| TRIM33 | Q9UPN9 | 0.806 | 0.417 |
| CDC42BPA | Q5VT25 | 0.865 | 0.416 |
| KPNB1 | Q14974 | 0.799 | 0.415 |
| SRP54 | P61011 | 0.751 | 0.415 |
| TRAF3 | Q13114 | 0.794 | 0.414 |
| VPS26B | Q4G0F5 | 0.836 | 0.414 |
| EIF3F | O00303 | 0.817 | 0.413 |
| TIMM50 | Q3ZCQ8 | 0.766 | 0.413 |
| AHCTF1 | Q8WYP5 | 0.855 | 0.412 |
| EIF2B4.1 | A0A087WTA5 | 0.812 | 0.412 |
| SNRPF | P62306 | 0.783 | 0.411 |
| UIMC1 | Q96RL1 | 0.858 | 0.411 |
| BCLAF1 | Q9NYF8 | 0.701 | 0.41 |
| TEX10 | Q9NXF1 | 0.834 | 0.41 |
| PLEKHG3 | A1L390 | 0.798 | 0.409 |
| NMRAL1 | Q9HBL8 | 0.849 | 0.408 |
| PAK1IP1 | Q9NWT1 | 0.811 | 0.407 |
| GTF2B | Q00403 | 0.807 | 0.406 |
| LDHA | P00338 | 0.866 | 0.405 |
| NOP14 | P78316 | 0.846 | 0.405 |
| SPOUT1 | Q5T280 | 0.802 | 0.405 |
| GEMIN5 | Q8TEQ6 | 0.711 | 0.403 |
| LRRIQ1 | Q96JM4 | 0.724 | 0.403 |
| PLAU | P00749 | 0.835 | 0.403 |
| XRCC1 | P18887 | 0.732 | 0.403 |
| ACO1 | P21399 | 0.845 | 0.402 |
| AREG | P15514 | 0.798 | 0.402 |
| CAPZB | B1AK88 | 0.853 | 0.402 |

| | | | |
|---|---|---|---|
| FLOT1 | O75955 | 0.855 | 0.401 |
| MED24 | A0A0B4J1W0 | 0.835 | 0.401 |
| GTSE1 | Q9NYZ3 | 0.776 | 0.399 |
| NOL6 | Q9H6R4 | 0.878 | 0.399 |
| XRCC5 | P13010 | 0.775 | 0.399 |
| NUP62 | P37198 | 0.756 | 0.398 |
| PLOD1 | Q02809 | 0.786 | 0.397 |
| SETX | Q7Z333 | 0.85 | 0.397 |
| HPSE | Q9Y251 | 0.801 | 0.395 |
| SAFB | Q15424 | 0.687 | 0.395 |
| CD55 | H7BY55 | 0.87 | 0.394 |
| EIF4G1.1 | C9K073 | 0.824 | 0.394 |
| GNAI1 | P63096 | 0.874 | 0.393 |
| PARP2.1 | E9PJ27 | 0.779 | 0.393 |
| WASHC2C | A0A096LPC5 | 0.843 | 0.393 |
| RPL34 | P49207 | 0.865 | 0.392 |
| CPSF4 | O95639 | 0.835 | 0.391 |
| GTPBP10 | A4D1E9 | 0.847 | 0.389 |
| ING2 | Q9H160 | 0.818 | 0.389 |
| TWNK | Q96RR1 | 0.782 | 0.389 |
| BRCA2 | P51587 | 0.81 | 0.388 |
| NPM1 | P06748 | 0.81 | 0.388 |
| RPL27 | P61353 | 0.816 | 0.388 |
| GCOM1 | H8Y6P7 | 0.801 | 0.387 |
| MRPL10 | Q7Z7H8 | 0.807 | 0.387 |
| HMGA1 | P17096 | 0.781 | 0.386 |
| RNF40 | O75150 | 0.831 | 0.385 |
| RNF214 | Q8ND24 | 0.843 | 0.384 |
| CDC42BPB.1 | H0YLY0 | 0.833 | 0.383 |
| IMPDH2 | P12268 | 0.859 | 0.383 |
| NACC1 | Q96RE7 | 0.798 | 0.382 |
| WASHC4 | A0A087X256 | 0.805 | 0.382 |
| SLC25A1 | P53007 | 0.789 | 0.381 |
| ASPM | Q8IZT6 | 0.824 | 0.38 |
| EPHA2 | P29317 | 0.858 | 0.379 |
| PKN3 | Q6P5Z2 | 0.855 | 0.379 |
| SEC24B | O95487 | 0.782 | 0.379 |
| YAP1 | P46937 | 0.846 | 0.379 |
| NBR1 | Q14596 | 0.823 | 0.378 |
| NDUFV1 | P49821 | 0.833 | 0.377 |
| 7-Sep | E7EPK1 | 0.79 | 0.377 |
| TIMM44 | O43615 | 0.82 | 0.377 |
| MRPS18B | Q9Y676 | 0.819 | 0.375 |
| WDR92 | Q96MX6 | 0.829 | 0.375 |
| CCM2 | Q9BSQ5 | 0.806 | 0.374 |
| CEP170.1 | H0Y2V6 | 0.825 | 0.374 |
| DDX52 | Q9Y2R4 | 0.859 | 0.374 |

| | | | |
|---|---|---|---|
| PATZ1 | Q9HBE1 | 0.847 | 0.373 |
| UTP15 | Q8TED0 | 0.842 | 0.373 |
| DDX39A | O00148 | 0.841 | 0.372 |
| AMPD2 | Q01433 | 0.811 | 0.371 |
| ANK3 | Q12955 | 0.839 | 0.371 |
| IDH1 | O75874 | 0.873 | 0.371 |
| KEAP1 | Q14145 | 0.719 | 0.371 |
| CSNK1G1 | U3KQB3 | 0.79 | 0.37 |
| FBXO31 | Q5XUX0 | 0.849 | 0.37 |
| MAP7D1.1 | H0YF21 | 0.848 | 0.37 |
| CHD3 | Q12873 | 0.765 | 0.369 |
| FMR1.1 | Q06787 | 0.749 | 0.368 |
| NUP133 | Q8WUM0 | 0.774 | 0.368 |
| QSER1 | Q2KHR3 | 0.724 | 0.367 |
| SBDS | Q9Y3A5 | 0.849 | 0.367 |
| BCORL1 | Q5H9F3 | 0.86 | 0.366 |
| CARM1 | Q86X55 | 0.848 | 0.366 |
| OGFOD1 | Q8N543 | 0.848 | 0.365 |
| ANAPC5 | Q9UJX4 | 0.784 | 0.364 |
| SERBP1 | Q8NC51 | 0.838 | 0.364 |
| LUC7L2 | Q9Y383 | 0.84 | 0.363 |
| PNN | Q9H307 | 0.788 | 0.362 |
| POTEI | P0CG38 | 0.85 | 0.361 |
| TEFM | Q96QE5 | 0.827 | 0.361 |
| DNAJB5 | O75953 | 0.826 | 0.36 |
| FNDC3B | Q53EP0 | 0.806 | 0.36 |
| PIGT | A0A1W2PNP0 | 0.798 | 0.36 |
| MDH2 | P40926 | 0.869 | 0.358 |
| ACAD11 | Q709F0 | 0.803 | 0.357 |
| RGL3 | A0A0A0MRX4 | 0.801 | 0.357 |
| VIRMA | Q69YN4 | 0.863 | 0.356 |
| GNA11 | P29992 | 0.797 | 0.355 |
| GRHL2 | Q6ISB3 | 0.853 | 0.355 |
| PRKACA | P17612 | 0.847 | 0.355 |
| JUND | P17535 | 0.808 | 0.354 |
| TRIO | O75962 | 0.841 | 0.354 |
| BRD8 | H7C127 | 0.867 | 0.353 |
| FAM98A | Q8NCA5 | 0.822 | 0.353 |
| NELFCD | H0UI80 | 0.849 | 0.352 |
| ZFX | P17010 | 0.861 | 0.352 |
| TFB2M | Q9H5Q4 | 0.853 | 0.351 |
| TRIM46 | Q7Z4K8 | 0.845 | 0.351 |
| MDC1 | Q14676 | 0.865 | 0.35 |
| TPM3 | A0A087WWU8 | 0.87 | 0.35 |
| AP3S1 | Q92572 | 0.851 | 0.349 |
| MSTO1 | Q9BUK6 | 0.841 | 0.349 |
| GGCT | O75223 | 0.857 | 0.348 |

| | | | |
|---|---|---|---|
| MRPL11 | Q9Y3B7 | 0.745 | 0.348 |
| NSD2 | O96028 | 0.872 | 0.348 |
| KIF21A | Q7Z4S6 | 0.861 | 0.347 |
| UNC45A | A0A1W2PNX8 | 0.835 | 0.347 |
| SPATS2L.1 | F8W6C2 | 0.759 | 0.346 |
| VPS35.1 | A0A1W2PP10 | 0.868 | 0.345 |
| BLM | P54132 | 0.853 | 0.344 |
| SRPK2.1 | C9JWF7 | 0.838 | 0.344 |
| DEF6 | Q9H4E7 | 0.822 | 0.343 |
| CD3EAP | O15446 | 0.854 | 0.342 |
| GEMIN4 | P57678 | 0.782 | 0.342 |
| KANSL2 | F8VX10 | 0.874 | 0.342 |
| PARVA | J3KNQ4 | 0.814 | 0.342 |
| CALU | O43852 | 0.846 | 0.341 |
| NGDN | Q8NEJ9 | 0.846 | 0.341 |
| TNRC6A | Q8NDV7 | 0.808 | 0.341 |
| GOLGA2 | Q08379 | 0.851 | 0.34 |
| SORBS3 | O60504 | 0.818 | 0.34 |
| CRTC2 | Q53ET0 | 0.836 | 0.339 |
| COL18A1 | P39060 | 0.797 | 0.338 |
| ACADVL | P49748 | 0.848 | 0.337 |
| CMSS1 | Q9BQ75 | 0.831 | 0.337 |
| DDX19B | H3BQK0 | 0.848 | 0.337 |
| EXOSC2 | Q13868 | 0.835 | 0.337 |
| NCBP3 | Q53F19 | 0.832 | 0.337 |
| RPL7A | P62424 | 0.866 | 0.337 |
| TDRD3 | Q9H7E2 | 0.799 | 0.337 |
| RPL18A | Q02543 | 0.87 | 0.336 |
| RPS20 | P60866 | 0.86 | 0.336 |
| STAU1 | O95793 | 0.754 | 0.336 |
| CHM | P24386 | 0.844 | 0.335 |
| F2RL1 | P55085 | 0.758 | 0.335 |
| PLEC | Q15149 | 0.798 | 0.335 |
| PCYOX1 | Q9UHG3 | 0.843 | 0.334 |
| RSBN1L | Q6PCB5 | 0.849 | 0.334 |
| TACC1 | O75410 | 0.855 | 0.334 |
| ALDH18A1 | P54886 | 0.844 | 0.333 |
| BUB1B | O60566 | 0.761 | 0.333 |
| EEA1 | Q15075 | 0.866 | 0.333 |
| LRRC8A | Q8IWT6 | 0.815 | 0.333 |
| SRSF6 | Q13247 | 0.795 | 0.333 |
| INCENP | Q9NQS7 | 0.834 | 0.332 |
| NUMA1 | Q14980 | 0.874 | 0.332 |
| TKFC | Q3LXA3 | 0.868 | 0.332 |
| CD59 | E9PR17 | 0.871 | 0.331 |
| NUDT21 | O43809 | 0.856 | 0.331 |
| TTC31 | Q49AM3 | 0.837 | 0.331 |

| | | | |
|---|---|---|---|
| TRIM15 | A0A0G2JJP1 | 0.787 | 0.33 |
| HECTD1 | Q9ULT8 | 0.823 | 0.329 |
| PBRM1 | Q86U86 | 0.878 | 0.328 |
| AP1G1 | O43747 | 0.845 | 0.327 |
| CLTC | A0A087WVQ6 | 0.822 | 0.326 |
| MTO1 | Q9Y2Z2 | 0.815 | 0.326 |
| NLRP2 | A0A0G2JPQ2 | 0.816 | 0.326 |
| PELP1 | C9JFV4 | 0.875 | 0.326 |
| BRIP1 | Q9BX63 | 0.809 | 0.325 |
| TRIM25 | Q14258 | 0.843 | 0.325 |
| ARHGAP5 | Q13017 | 0.822 | 0.324 |
| ERCC6 | Q03468 | 0.866 | 0.324 |
| UBR5 | O95071 | 0.84 | 0.324 |
| CERK | Q8TCT0 | 0.867 | 0.323 |
| PPP1R13L | Q8WUF5 | 0.826 | 0.323 |
| RPS5 | P46782 | 0.845 | 0.322 |
| PHF10 | Q8WUB8 | 0.871 | 0.321 |
| SULF2 | Q8IWU5 | 0.812 | 0.321 |
| AFF1 | P51825 | 0.763 | 0.32 |
| EPS8L2 | Q9H6S3 | 0.866 | 0.32 |
| PKP2 | Q99959 | 0.795 | 0.32 |
| TJP3 | O95049 | 0.863 | 0.319 |
| ANKRD28 | O15084 | 0.837 | 0.318 |
| MLEC | Q14165 | 0.876 | 0.318 |
| MOV10 | Q9HCE1 | 0.78 | 0.318 |
| TRAF3IP1 | Q8TDR0 | 0.865 | 0.318 |
| DENND2D | Q9H6A0 | 0.867 | 0.317 |
| RALGAPA1 | A0A1B0GUI1 | 0.858 | 0.316 |
| TYK2 | P29597 | 0.778 | 0.315 |
| LAD1 | O00515 | 0.834 | 0.314 |
| RPL7 | P18124 | 0.866 | 0.314 |
| HNRNPAB | D6RBZ0 | 0.821 | 0.313 |
| AAK1 | Q2M2I8 | 0.821 | 0.311 |
| EEF1B2 | P24534 | 0.844 | 0.311 |
| GOLGA5 | Q8TBA6 | 0.863 | 0.311 |
| MED17.1 | A0A1W2PRB8 | 0.841 | 0.311 |
| MTAP | B4DUC8 | 0.876 | 0.311 |
| APEH | C9JIF9 | 0.861 | 0.31 |
| CDK5RAP1 | Q96SZ6 | 0.837 | 0.308 |
| RCN2 | Q14257 | 0.789 | 0.307 |
| SEMA3C | Q99985 | 0.824 | 0.307 |
| TBC1D8B | Q0IIM8 | 0.84 | 0.306 |
| NUP85 | Q9BW27 | 0.846 | 0.305 |
| NUP205 | Q92621 | 0.78 | 0.304 |
| PRDX4 | Q13162 | 0.864 | 0.304 |
| LARP1 | Q6PKG0 | 0.784 | 0.303 |
| DDX18 | Q9NVP1 | 0.862 | 0.302 |

| | | | |
|---|---|---|---|
| MDH1 | P40925 | 0.88 | 0.302 |
| ANKRD17 | O75179 | 0.817 | 0.301 |
| UBR2 | Q8IWV8 | 0.857 | 0.301 |
| DCTN1 | E7EX90 | 0.859 | 0.3 |
| RPL10A | P62906 | 0.873 | 0.3 |
| SNX2 | O60749 | 0.873 | 0.3 |
| FNDC3A | Q9Y2H6 | 0.832 | 0.299 |
| ING3 | Q9NXR8 | 0.865 | 0.299 |
| HJURP | Q8NCD3 | 0.824 | 0.298 |
| SRSF1 | J3KTL2 | 0.816 | 0.297 |
| RACK1 | P63244 | 0.868 | 0.296 |
| MED17 | Q9NVC6 | 0.846 | 0.295 |
| MYO9A | B2RTY4 | 0.858 | 0.295 |
| PPIA | P62937 | 0.835 | 0.295 |
| HIRA | P54198 | 0.877 | 0.293 |
| NCSTN | Q92542 | 0.859 | 0.291 |
| SPTLC1 | O15269 | 0.854 | 0.291 |
| TRMT10C | Q7L0Y3 | 0.819 | 0.291 |
| COLGALT1 | Q8NBJ5 | 0.859 | 0.29 |
| KDM2A | Q9Y2K7 | 0.821 | 0.29 |
| RBMX | P38159 | 0.857 | 0.29 |
| RPRD1A | Q96P16 | 0.819 | 0.29 |
| RRS1 | Q15050 | 0.818 | 0.29 |
| DDX20 | Q9UHI6 | 0.819 | 0.288 |
| DTL | Q9NZJ0 | 0.847 | 0.288 |
| CDKAL1 | Q5VV42 | 0.834 | 0.286 |
| DAP3 | P51398 | 0.803 | 0.286 |
| FBXO11 | Q86XK2 | 0.87 | 0.286 |
| MAGT1 | A0A087WU53 | 0.802 | 0.286 |
| VAPB | O95292 | 0.862 | 0.286 |
| CDC40 | O60508 | 0.848 | 0.285 |
| CEP78 | A8MST6 | 0.853 | 0.285 |
| MMS22L | Q6ZRQ5 | 0.857 | 0.285 |
| PYM1 | Q9BRP8 | 0.849 | 0.284 |
| SMNDC1 | O75940 | 0.868 | 0.284 |
| UFD1 | Q92890 | 0.851 | 0.284 |
| RFTN1 | Q14699 | 0.877 | 0.283 |
| TSG101 | Q99816 | 0.819 | 0.283 |
| NIFK | Q9BYG3 | 0.848 | 0.282 |
| YME1L1 | Q96TA2 | 0.84 | 0.282 |
| ZSWIM9 | Q86XI8 | 0.853 | 0.282 |
| CCNB2 | O95067 | 0.844 | 0.281 |
| CDC42BPB | Q9Y5S2 | 0.847 | 0.277 |
| G3BP1 | Q13283 | 0.837 | 0.277 |
| GZF1 | Q9H116 | 0.865 | 0.277 |
| LIMD1 | Q9UGP4 | 0.855 | 0.277 |
| WRN | Q14191 | 0.862 | 0.277 |

| | | | |
|---|---|---|---|
| USP15 | Q9Y4E8 | 0.862 | 0.276 |
| CYBA | P13498 | 0.857 | 0.275 |
| UHRF2 | Q96PU4 | 0.866 | 0.275 |
| ZNF330 | Q9Y3S2 | 0.87 | 0.275 |
| FTSJ3 | Q8IY81 | 0.873 | 0.274 |
| FBL | P22087 | 0.856 | 0.273 |
| MATR3 | A8MXP9 | 0.807 | 0.272 |
| C8orf33 | Q9H7E9 | 0.827 | 0.27 |
| CSNK2A2 | P19784 | 0.825 | 0.269 |
| RPS27A | P62979 | 0.85 | 0.269 |
| SUPT6H | Q7KZ85 | 0.861 | 0.267 |
| TRIP11 | Q15643 | 0.856 | 0.267 |
| SDSL | Q96GA7 | 0.821 | 0.266 |
| ZNHIT6 | Q9NWK9 | 0.862 | 0.266 |
| CLPX | O76031 | 0.834 | 0.265 |
| LARP4B | Q92615 | 0.847 | 0.265 |
| PABPC1 | P11940 | 0.849 | 0.265 |
| UGGT2 | Q9NYU1 | 0.875 | 0.265 |
| CARMIL1 | Q5VZK9 | 0.855 | 0.263 |
| HNRNPD | Q14103 | 0.84 | 0.261 |
| PPAN-P2RY11 | A0A0B4J1V8 | 0.871 | 0.261 |
| TMA7 | Q9Y2S6 | 0.877 | 0.259 |
| ZNF629 | Q9UEG4 | 0.87 | 0.259 |
| C3orf58 | Q8NDZ4 | 0.833 | 0.258 |
| CHPF2 | Q9P2E5 | 0.824 | 0.258 |
| COPE | M0QXB4 | 0.871 | 0.258 |
| VDAC2 | P45880 | 0.871 | 0.258 |
| MAP3K7 | O43318 | 0.874 | 0.257 |
| DPYSL2 | A0A1C7CYX9 | 0.873 | 0.256 |
| MICB | A0A0G2JH58 | 0.871 | 0.256 |
| PHRF1 | Q9P1Y6 | 0.865 | 0.256 |
| USP10 | Q14694 | 0.834 | 0.255 |
| LMNB1 | P20700 | 0.836 | 0.254 |
| NMT2 | O60551 | 0.861 | 0.254 |
| ECD | O95905 | 0.853 | 0.252 |
| MACO1 | Q8N5G2 | 0.841 | 0.252 |
| YARS2 | Q9Y2Z4 | 0.863 | 0.252 |
| CARS | P49589 | 0.865 | 0.25 |
| EIF2A | Q9BY44 | 0.845 | 0.25 |
| GTF3C1 | Q12789 | 0.88 | 0.25 |
| LRRFIP1 | Q32MZ4 | 0.846 | 0.247 |
| ZC3HAV1.1 | C9J6P4 | 0.856 | 0.247 |
| NQO1 | P15559 | 0.881 | 0.246 |
| RPL5 | P46777 | 0.874 | 0.246 |
| ZKSCAN1 | P17029 | 0.856 | 0.246 |
| RPL15 | P61313 | 0.879 | 0.245 |
| GSTP1 | P09211 | 0.883 | 0.244 |

| | | | |
|---|---|---|---|
| TECR | Q9NZ01 | 0.815 | 0.244 |
| ZMPSTE24 | O75844 | 0.883 | 0.244 |
| TWF1 | Q12792 | 0.875 | 0.243 |
| ALDH1A3 | P47895 | 0.878 | 0.24 |
| HUWE1 | Q7Z6Z7 | 0.848 | 0.24 |
| IGHMBP2 | P38935 | 0.869 | 0.24 |
| ZCCHC7 | Q8N3Z6 | 0.88 | 0.24 |
| RPTOR | Q8N122 | 0.812 | 0.239 |
| TJP2 | A0A1B0GTW1 | 0.857 | 0.239 |
| FOXM1 | Q08050 | 0.873 | 0.237 |
| LYAR | Q9NX58 | 0.88 | 0.237 |
| PTDSS1 | P48651 | 0.873 | 0.237 |
| MTCH2 | Q9Y6C9 | 0.877 | 0.235 |
| POLE | Q07864 | 0.82 | 0.235 |
| GBF1 | Q92538 | 0.841 | 0.234 |
| LYZ | P61626 | 0.874 | 0.234 |
| ANK1 | P16157 | 0.87 | 0.233 |
| TSFM | P43897 | 0.865 | 0.233 |
| URB1 | O60287 | 0.881 | 0.23 |
| ZNF219 | Q9P2Y4 | 0.848 | 0.23 |
| MINDY4 | Q4G0A6 | 0.859 | 0.229 |
| NUMBL | Q9Y6R0 | 0.874 | 0.229 |
| CEP170B | J3KQR7 | 0.839 | 0.228 |
| CYFIP2 | E7EVJ5 | 0.859 | 0.228 |
| ODF2 | Q5BJF6 | 0.872 | 0.227 |
| CSNK1G3 | Q9Y6M4 | 0.826 | 0.226 |
| ICE1 | Q9Y2F5 | 0.868 | 0.226 |
| PDAP1 | Q13442 | 0.848 | 0.226 |
| RNPC3 | Q96LT9 | 0.883 | 0.226 |
| SNX9 | Q9Y5X1 | 0.838 | 0.226 |
| SPATS2 | Q86XZ4 | 0.854 | 0.226 |
| DTWD1 | Q8N5C7 | 0.854 | 0.224 |
| HNRNPM | P52272 | 0.861 | 0.224 |
| ANKHD1 | Q8IWZ3 | 0.836 | 0.223 |
| COQ8A | Q8NI60 | 0.867 | 0.223 |
| LEMD2 | Q8NC56 | 0.871 | 0.223 |
| RPSA | A0A0C4DG17 | 0.857 | 0.223 |
| TRA2B | P62995 | 0.858 | 0.223 |
| BTF3 | P20290 | 0.872 | 0.222 |
| CCDC86 | Q9H6F5 | 0.868 | 0.222 |
| FDFT1 | A0A1W2PQ47 | 0.877 | 0.222 |
| PLS3 | P13797 | 0.879 | 0.222 |
| SPATS2L | Q9NUQ6 | 0.853 | 0.222 |
| SPTBN2 | O15020 | 0.842 | 0.222 |
| IQSEC1 | A0A087WWK8 | 0.871 | 0.221 |
| ZFR | Q96KR1 | 0.854 | 0.221 |
| TOP3B | O95985 | 0.859 | 0.22 |

| | | | |
|---|---|---|---|
| BAHCC1 | Q9P281 | 0.874 | 0.218 |
| DPP3 | G3V180 | 0.878 | 0.218 |
| TMPO | P42166 | 0.842 | 0.218 |
| TWF2 | Q6IBS0 | 0.882 | 0.218 |
| USP43 | Q70EL4 | 0.863 | 0.218 |
| VDAC1 | P21796 | 0.878 | 0.218 |
| ALDH2 | P05091 | 0.882 | 0.217 |
| KIAA0319L | Q8IZA0 | 0.859 | 0.217 |
| RRP9 | O43818 | 0.864 | 0.217 |
| SLFN5 | Q08AF3 | 0.845 | 0.217 |
| PAPD7 | A0A0X1KG68 | 0.826 | 0.216 |
| PHKB | Q93100 | 0.859 | 0.216 |
| SERPINH1 | P50454 | 0.87 | 0.216 |
| FIP1L1 | Q6UN15 | 0.867 | 0.215 |
| RPS3 | P23396 | 0.874 | 0.215 |
| ACO2 | A2A274 | 0.88 | 0.214 |
| ATP6V1B2 | P21281 | 0.847 | 0.214 |
| UBN2 | Q6ZU65 | 0.881 | 0.214 |
| KATNAL2 | Q8IYT4 | 0.87 | 0.213 |
| NUMB | P49757 | 0.879 | 0.213 |
| SCAF11 | Q99590 | 0.882 | 0.213 |
| DYNC1I2 | Q13409 | 0.874 | 0.212 |
| EHMT1 | Q9H9B1 | 0.884 | 0.212 |
| OXSR1 | O95747 | 0.855 | 0.212 |
| CPS1 | P31327 | 0.863 | 0.211 |
| EIF2S2 | P20042 | 0.867 | 0.211 |
| ELF2 | Q15723 | 0.867 | 0.211 |
| OSBPL5 | Q9H0X9 | 0.874 | 0.211 |
| RASAL2 | Q9UJF2 | 0.878 | 0.21 |
| MRPS30 | Q9NP92 | 0.832 | 0.208 |
| DLST | P36957 | 0.88 | 0.206 |
| HSPA2 | P54652 | 0.863 | 0.206 |
| EIF4G1 | Q04637 | 0.858 | 0.205 |
| FRA10AC1 | Q70Z53 | 0.865 | 0.205 |
| G3BP2 | Q9UN86 | 0.867 | 0.205 |
| MFAP1 | P55081 | 0.855 | 0.205 |
| TBC1D4 | O60343 | 0.846 | 0.205 |
| SRP68 | Q9UHB9 | 0.842 | 0.204 |
| CENPF | P49454 | 0.876 | 0.203 |
| MAP1A.1 | E9PGC8 | 0.88 | 0.203 |
| SMG7 | E9PD50 | 0.87 | 0.202 |
| DHX33 | Q9H6R0 | 0.878 | 0.201 |
| SRC | P12931 | 0.879 | 0.201 |
| ZNF462 | Q96JM2 | 0.881 | 0.2 |
| RPL18 | J3QQ67 | 0.88 | 0.199 |
| TRAF7 | Q6Q0C0 | 0.854 | 0.199 |
| FAM207A | Q9NSI2 | 0.865 | 0.198 |

| | | | |
|---|---|---|---|
| SFXN1 | Q9H9B4 | 0.885 | 0.198 |
| SERPINB1 | P30740 | 0.868 | 0.197 |
| UPF3B | Q9BZI7 | 0.874 | 0.196 |
| AIMP1 | Q12904 | 0.873 | 0.195 |
| RPAP2 | Q8IXW5 | 0.853 | 0.195 |
| COPS4 | Q9BT78 | 0.878 | 0.194 |
| MKL2 | Q9ULH7 | 0.868 | 0.194 |
| OSTC | Q9NRP0 | 0.878 | 0.194 |
| PDP1 | Q9P0J1 | 0.86 | 0.194 |
| EXO1 | Q9UQ84 | 0.882 | 0.193 |
| RPL4 | P36578 | 0.865 | 0.192 |
| TEX2 | A0A087WVA8 | 0.876 | 0.192 |
| CAMSAP3 | Q9P1Y5 | 0.877 | 0.191 |
| DDX5 | J3KTA4 | 0.848 | 0.191 |
| RAN | J3KQE5 | 0.875 | 0.191 |
| BUD23 | O43709 | 0.861 | 0.19 |
| ZC3H6 | P61129 | 0.877 | 0.19 |
| TBL2 | Q9Y4P3 | 0.863 | 0.189 |
| IFT74 | Q96LB3 | 0.876 | 0.188 |
| OCRL | Q01968 | 0.868 | 0.188 |
| TOP3A | Q13472 | 0.862 | 0.188 |
| FAM120A | Q9NZB2 | 0.855 | 0.187 |
| FASN | P49327 | 0.864 | 0.187 |
| KLC1 | G3V5R9 | 0.872 | 0.187 |
| SIN3B | O75182 | 0.884 | 0.187 |
| SLK | Q9H2G2 | 0.857 | 0.187 |
| DBT | P11182 | 0.881 | 0.186 |
| INVS | Q9Y283 | 0.867 | 0.186 |
| HSPA1B | A0A0G2JIW1 | 0.862 | 0.185 |
| PPIB | P23284 | 0.842 | 0.185 |
| XPNPEP1 | Q9NQW7 | 0.882 | 0.185 |
| APTX | Q7Z2E3 | 0.86 | 0.184 |
| RPP30 | P78346 | 0.868 | 0.184 |
| SSH1 | Q8WYL5 | 0.877 | 0.184 |
| FAM92A | A1XBS5 | 0.87 | 0.183 |
| ACTN4 | O43707 | 0.877 | 0.182 |
| CLCA3P | Q9Y6N3 | 0.865 | 0.181 |
| DHX30 | Q7L2E3 | 0.846 | 0.181 |
| MTDH | Q86UE4 | 0.873 | 0.18 |
| PLK1 | P53350 | 0.866 | 0.18 |
| CANX.1 | D6RB85 | 0.878 | 0.179 |
| ERC1 | X6RLX0 | 0.864 | 0.179 |
| TAF9 | Q16594 | 0.872 | 0.179 |
| VAV2 | P52735 | 0.876 | 0.179 |
| CSTF3 | Q12996 | 0.872 | 0.178 |
| HLA-B | P30483 | 0.88 | 0.178 |
| LARP4 | Q71RC2 | 0.869 | 0.178 |

| | | | |
|---|---|---|---|
| NUP188 | Q5SRE5 | 0.853 | 0.178 |
| PCCB | E9PDR0 | 0.887 | 0.178 |
| NOA1 | Q8NC60 | 0.848 | 0.177 |
| PPFIA3 | O75145 | 0.869 | 0.177 |
| ITGA2 | P17301 | 0.867 | 0.176 |
| ZNF574 | A0A0C4DFM2 | 0.881 | 0.176 |
| RSBN1 | Q5VWQ0 | 0.882 | 0.175 |
| MAPKBP1 | O60336 | 0.868 | 0.173 |
| PRPF19 | Q9UMS4 | 0.877 | 0.173 |
| RAVER1 | E9PAU2 | 0.876 | 0.173 |
| CAVIN1 | Q6NZI2 | 0.865 | 0.172 |
| GPRASP1 | Q5JY77 | 0.88 | 0.172 |
| MAGED1 | Q9Y5V3 | 0.882 | 0.172 |
| RPUSD3 | Q6P087 | 0.871 | 0.172 |
| SUPT5H | O00267 | 0.877 | 0.172 |
| GAS8 | O95995 | 0.872 | 0.171 |
| SNRPD1 | P62314 | 0.846 | 0.17 |
| ADIG | H0Y829 | 0.874 | 0.169 |
| SNRPB2 | P08579 | 0.863 | 0.168 |
| U2AF1 | Q01081 | 0.878 | 0.168 |
| CSTF2T | Q9H0L4 | 0.858 | 0.166 |
| PRKCD | Q05655 | 0.869 | 0.166 |
| TTC17 | Q96AE7 | 0.879 | 0.166 |
| ARHGAP17 | Q68EM7 | 0.885 | 0.163 |
| DOCK1 | A0A096LNH6 | 0.856 | 0.163 |
| NOC4L | Q9BVI4 | 0.88 | 0.163 |
| EXOSC8 | Q96B26 | 0.882 | 0.162 |
| RPAP3 | Q9H6T3 | 0.854 | 0.161 |
| V9GY48 | V9GY48 | 0.865 | 0.161 |
| CALM2 | E7EMB3 | 0.885 | 0.16 |
| MARK1 | A0A087X0I6 | 0.885 | 0.16 |
| CDK8 | P49336 | 0.867 | 0.159 |
| AURKA | O14965 | 0.872 | 0.158 |
| UBTF | P17480 | 0.879 | 0.158 |
| EIF4G2 | P78344 | 0.874 | 0.157 |
| GPI | A0A0A0MTS2 | 0.883 | 0.157 |
| HSD17B10 | Q99714 | 0.86 | 0.157 |
| TCF20 | Q9UGU0 | 0.879 | 0.156 |
| CEP89 | Q96ST8 | 0.877 | 0.155 |
| HIST1H1E | P10412 | 0.867 | 0.155 |
| PNKP | Q96T60 | 0.863 | 0.155 |
| RBM11 | P57052 | 0.861 | 0.155 |
| DPF2 | Q92785 | 0.882 | 0.154 |
| NAV3 | Q8IVL0 | 0.871 | 0.154 |
| DHX9 | Q08211 | 0.858 | 0.153 |
| HIST1H1C | P16403 | 0.866 | 0.153 |
| LRCH4 | O75427 | 0.873 | 0.153 |

| | | | |
|---|---|---|---|
| PLD3 | Q8IV08 | 0.875 | 0.153 |
| PYGL | P06737 | 0.884 | 0.153 |
| ZNF335 | Q9H4Z2 | 0.889 | 0.153 |
| PSMD3 | O43242 | 0.884 | 0.152 |
| RBFOX2 | O43251 | 0.871 | 0.152 |
| 9-Sep | Q9UHD8 | 0.872 | 0.151 |
| DVL1 | O14640 | 0.878 | 0.15 |
| RRBP1 | Q9P2E9 | 0.877 | 0.15 |
| BUD13 | Q9BRD0 | 0.885 | 0.149 |
| ESCO2 | Q56NI9 | 0.88 | 0.149 |
| KCMF1 | Q9P0J7 | 0.885 | 0.149 |
| NSF | P46459 | 0.885 | 0.149 |
| NUP43 | Q8NFH3 | 0.881 | 0.149 |
| MAP4K4 | E7EN19 | 0.885 | 0.148 |
| RRP1B | Q14684 | 0.882 | 0.148 |
| SUPT7L | O94864 | 0.884 | 0.148 |
| AP3M1 | Q9Y2T2 | 0.88 | 0.147 |
| KRI1 | Q8N9T8 | 0.887 | 0.146 |
| MAGI1 | Q96QZ7 | 0.88 | 0.146 |
| RIPK4 | P57078 | 0.865 | 0.146 |
| TRIP6 | Q15654 | 0.87 | 0.146 |
| ADH5 | P11766 | 0.888 | 0.144 |
| DDX1 | Q92499 | 0.871 | 0.144 |
| NCKIPSD | Q9NZQ3 | 0.879 | 0.144 |
| GSE1 | Q14687 | 0.89 | 0.142 |
| RPS2 | P15880 | 0.886 | 0.142 |
| TBRG4 | Q969Z0 | 0.888 | 0.142 |
| DAG1 | Q14118 | 0.887 | 0.141 |
| SMARCAL1 | Q9NZC9 | 0.863 | 0.141 |
| ZBTB41 | Q5SVQ8 | 0.86 | 0.14 |
| DNAJC14 | Q6Y2X3 | 0.886 | 0.139 |
| THOC1 | A0A087WWS1 | 0.891 | 0.139 |
| FBXL6 | Q8N531 | 0.88 | 0.138 |
| R3HDM1 | Q15032 | 0.869 | 0.138 |
| ZC3HAV1 | Q7Z2W4 | 0.877 | 0.138 |
| CEPT1 | Q9Y6K0 | 0.863 | 0.137 |
| POLR1A | O95602 | 0.887 | 0.137 |
| RECQL4 | O94761 | 0.886 | 0.137 |
| SRSF2 | Q01130 | 0.875 | 0.137 |
| AP2M1 | E9PFW3 | 0.873 | 0.136 |
| CCDC174 | Q6PII3 | 0.882 | 0.136 |
| PRPF3 | O43395 | 0.882 | 0.136 |
| DPM1 | H0Y368 | 0.866 | 0.135 |
| PSMB4 | P28070 | 0.887 | 0.135 |
| TTK | P33981 | 0.864 | 0.135 |
| NAV1.1 | H0Y6F6 | 0.877 | 0.133 |
| POLDIP3 | F6VRR5 | 0.878 | 0.133 |

| | | | |
|---|---|---|---|
| MTMR10 | Q9NXD2 | 0.887 | 0.132 |
| SUPT16H | Q9Y5B9 | 0.873 | 0.132 |
| ASXL1 | Q8IXJ9 | 0.889 | 0.131 |
| DNAJC10 | Q8IXB1 | 0.877 | 0.131 |
| HACD3 | Q9P035 | 0.884 | 0.131 |
| LRRC1 | Q9BTT6 | 0.884 | 0.131 |
| DCAF7 | P61962 | 0.871 | 0.13 |
| ITSN1 | Q15811 | 0.87 | 0.13 |
| SP6 | Q3SY56 | 0.871 | 0.13 |
| DNAJC9 | Q8WXX5 | 0.886 | 0.129 |
| MICU1 | A0A286YF11 | 0.885 | 0.129 |
| EIF6 | P56537 | 0.885 | 0.128 |
| RFC1 | P35251 | 0.885 | 0.128 |
| ERI1 | Q8IV48 | 0.886 | 0.127 |
| KIF23 | Q02241 | 0.884 | 0.127 |
| MORC3 | Q14149 | 0.874 | 0.126 |
| MRPL39 | Q9NYK5 | 0.884 | 0.126 |
| NFIA | Q12857 | 0.887 | 0.126 |
| CPNE7 | Q9UBL6 | 0.869 | 0.123 |
| FXR1 | P51114 | 0.869 | 0.123 |
| LGALS3BP | Q08380 | 0.883 | 0.123 |
| PDCD2 | Q16342 | 0.872 | 0.123 |
| YWHAQ | P27348 | 0.891 | 0.123 |
| ARMC6 | Q6NXE6 | 0.883 | 0.122 |
| CTPS1 | P17812 | 0.888 | 0.122 |
| SUN1 | E9PHI4 | 0.882 | 0.122 |
| TRRAP.1 | H0Y4W2 | 0.888 | 0.122 |
| A0A0A0MRY4 | A0A0A0MRY4 | 0.89 | 0.121 |
| DAZAP1 | Q96EP5 | 0.875 | 0.121 |
| NUP153 | P49790 | 0.872 | 0.121 |
| SLC38A5 | Q8WUX1 | 0.879 | 0.121 |
| NOC3L | Q8WTT2 | 0.891 | 0.12 |
| NPTN | Q9Y639 | 0.887 | 0.12 |
| NUDC | Q9Y266 | 0.887 | 0.12 |
| SHPRH | A0A0D9SFM0 | 0.884 | 0.12 |
| TTC34 | A0A1C7CYW7 | 0.888 | 0.12 |
| DDB2 | Q92466 | 0.882 | 0.119 |
| RNF10 | Q8N5U6 | 0.887 | 0.119 |
| SCFD2 | Q8WU76 | 0.876 | 0.119 |
| ATXN2L.1 | H3BUF6 | 0.876 | 0.118 |
| CILP2 | K7EPJ4 | 0.872 | 0.118 |
| RMI1 | Q9H9A7 | 0.882 | 0.117 |
| KIF5B | P33176 | 0.874 | 0.115 |
| NR1H2 | P55055 | 0.888 | 0.115 |
| SLC25A5 | P05141 | 0.877 | 0.115 |
| COPG2 | Q9UBF2 | 0.889 | 0.114 |
| ERAL1 | O75616 | 0.874 | 0.114 |

| | | | |
|---|---|---|---|
| CGN | Q9P2M7 | 0.884 | 0.113 |
| DIEXF | Q68CQ4 | 0.885 | 0.113 |
| FBXO3 | Q9UK99 | 0.875 | 0.113 |
| ATXN7 | O15265 | 0.886 | 0.111 |
| BMP2K | Q9NSY1 | 0.886 | 0.11 |
| VTA1 | Q9NP79 | 0.883 | 0.11 |
| HGS | O14964 | 0.889 | 0.109 |
| POMGNT2 | Q8NAT1 | 0.881 | 0.109 |
| RALBP1 | Q15311 | 0.888 | 0.109 |
| ILF3 | Q12906 | 0.88 | 0.108 |
| KNOP1 | Q1ED39 | 0.889 | 0.107 |
| TRMT1L | Q7Z2T5 | 0.871 | 0.107 |
| CTNNA1 | P35221 | 0.875 | 0.106 |
| FECH | P22830 | 0.88 | 0.105 |
| KIFC3 | Q9BVG8 | 0.872 | 0.104 |
| SMARCA1 | B7ZLQ5 | 0.884 | 0.104 |
| ARRB1 | P49407 | 0.88 | 0.103 |
| NFRKB | Q6P4R8 | 0.891 | 0.103 |
| SUPT20H | Q8NEM7 | 0.888 | 0.103 |
| ZNF292 | O60281 | 0.89 | 0.103 |
| SHMT2 | P34897 | 0.884 | 0.102 |
| SPIRE1 | Q08AE8 | 0.889 | 0.101 |
| SRSF11 | Q05519 | 0.891 | 0.101 |
| PPA1 | Q15181 | 0.892 | 0.0998 |
| MAP4K4.1 | C9J840 | 0.892 | 0.0995 |
| TOR1AIP2 | Q8NFQ8 | 0.89 | 0.0991 |
| SNRPD2 | P62316 | 0.875 | 0.0985 |
| MTDH.2 | H0YBJ8 | 0.886 | 0.0983 |
| JPH1 | Q9HDC5 | 0.889 | 0.0961 |
| NFIC | P08651 | 0.889 | 0.0961 |
| ZDHHC5 | Q9C0B5 | 0.893 | 0.0951 |
| NF2 | P35240 | 0.886 | 0.095 |
| RRP7A | Q9Y3A4 | 0.891 | 0.095 |
| CLK3 | P49761 | 0.889 | 0.0947 |
| TFB1M | Q8WVM0 | 0.886 | 0.0947 |
| POLR1E | Q9GZS1 | 0.887 | 0.0941 |
| SIRT6 | Q8N6T7 | 0.892 | 0.0938 |
| RCL1 | Q9Y2P8 | 0.884 | 0.093 |
| SRPK2 | P78362 | 0.89 | 0.0927 |
| ARHGAP12 | Q8IWW6 | 0.877 | 0.0925 |
| TES | Q9UGI8 | 0.891 | 0.0923 |
| GTF2F2 | P13984 | 0.888 | 0.0916 |
| YARS | P54577 | 0.887 | 0.0916 |
| ZCCHC14 | Q8WYQ9 | 0.884 | 0.0916 |
| G6PD | P11413 | 0.891 | 0.0915 |
| WDR36 | Q8NI36 | 0.891 | 0.0908 |
| PKN1 | Q16512 | 0.884 | 0.0897 |

| | | | |
|---|---|---|---|
| FOXO3 | O43524 | 0.883 | 0.0895 |
| HIP1R | O75146 | 0.887 | 0.0892 |
| NIPBL | Q6KC79 | 0.89 | 0.0886 |
| LMNA | P02545 | 0.881 | 0.0884 |
| ANKRD11 | Q6UB99 | 0.888 | 0.0881 |
| CXXC1 | Q9P0U4 | 0.886 | 0.0879 |
| GGA2 | Q9UJY4 | 0.881 | 0.0877 |
| EGLN1 | Q9GZT9 | 0.879 | 0.0875 |
| EIF3L | B0QY89 | 0.886 | 0.0871 |
| SLC25A6 | P12236 | 0.889 | 0.0868 |
| CDK17 | Q00537 | 0.881 | 0.0867 |
| PRPSAP1 | Q14558 | 0.89 | 0.0855 |
| ARHGEF26 | Q96DR7 | 0.884 | 0.0852 |
| ECT2 | Q9H8V3 | 0.891 | 0.0849 |
| GSPT1 | P15170 | 0.887 | 0.0847 |
| MAST1 | Q9Y2H9 | 0.893 | 0.0833 |
| TRIM56 | Q9BRZ2 | 0.887 | 0.0824 |
| RUVBL1 | Q9Y265 | 0.878 | 0.0816 |
| SRSF5 | Q13243 | 0.884 | 0.0815 |
| SRP72 | O76094 | 0.881 | 0.081 |
| LMNB2 | Q03252 | 0.882 | 0.0803 |
| PRPF40B | F8VU11 | 0.893 | 0.08 |
| ELP2 | Q6IA86 | 0.889 | 0.0794 |
| KLHL7 | Q8IXQ5 | 0.888 | 0.0791 |
| NUCKS1 | Q9H1E3 | 0.885 | 0.079 |
| EIF3D | O15371 | 0.88 | 0.0789 |
| TTC28 | Q96AY4 | 0.888 | 0.0787 |
| SMPD4 | Q9NXE4 | 0.88 | 0.0785 |
| ANLN | Q9NQW6 | 0.882 | 0.0782 |
| PDIA5 | Q14554 | 0.892 | 0.0781 |
| ARHGAP32 | A7KAX9 | 0.88 | 0.0778 |
| DHX37 | Q8IY37 | 0.892 | 0.0777 |
| ARHGAP19 | Q14CB8 | 0.888 | 0.0774 |
| FRMD8 | Q9BZ67 | 0.886 | 0.0772 |
| TAF6 | J3KR72 | 0.891 | 0.0772 |
| CTBP2 | Q5SQP8 | 0.886 | 0.0769 |
| RNGTT | O60942 | 0.882 | 0.0768 |
| UCKL1 | Q9NWZ5 | 0.882 | 0.0765 |
| CEP72 | Q9P209 | 0.889 | 0.0761 |
| RSPRY1 | Q96DX4 | 0.892 | 0.0761 |
| WDR33 | Q9C0J8 | 0.893 | 0.0759 |
| ZC3H15 | Q8WU90 | 0.891 | 0.0758 |
| ELOA | Q14241 | 0.88 | 0.0754 |
| ELP3 | Q9H9T3 | 0.881 | 0.0731 |
| INTS1 | Q8N201 | 0.89 | 0.0723 |
| GSK3A | P49840 | 0.887 | 0.0722 |
| CDC37 | Q16543 | 0.888 | 0.071 |

| | | | |
|---|---|---|---|
| XRN1 | Q8IZH2 | 0.882 | 0.0706 |
| ZNF148 | Q9UQR1 | 0.89 | 0.07 |
| STAT2 | P52630 | 0.886 | 0.0695 |
| DIMT1 | Q9UNQ2 | 0.886 | 0.0677 |
| ARHGEF10 | O15013 | 0.888 | 0.0671 |
| ZNF281 | Q9Y2X9 | 0.891 | 0.0669 |
| F5H5P2 | F5H5P2 | 0.893 | 0.0668 |
| NOP56 | O00567 | 0.889 | 0.0666 |
| CAPRIN1 | Q14444 | 0.888 | 0.0641 |
| EIF4B | E7EX17 | 0.891 | 0.064 |
| GTF2H1 | P32780 | 0.894 | 0.0637 |
| KDELR2 | P33947 | 0.888 | 0.0637 |
| SPAG5 | Q96R06 | 0.887 | 0.0634 |
| VPS16 | Q9H269 | 0.891 | 0.0634 |
| XRCC1.1 | F5H8D7 | 0.884 | 0.0632 |
| ADD3 | Q9UEY8 | 0.89 | 0.0626 |
| FMR1 | G3V0J0 | 0.884 | 0.0624 |
| CABLES2 | Q9BTV7 | 0.892 | 0.0618 |
| BLVRA | P53004 | 0.892 | 0.0617 |
| PPP4R3A | Q6IN85 | 0.893 | 0.0609 |
| VRK2 | Q86Y07 | 0.892 | 0.0603 |
| MAP1A | P78559 | 0.886 | 0.06 |
| DAPK3 | O43293 | 0.89 | 0.0597 |
| PLCB4 | Q15147 | 0.893 | 0.0587 |
| GNS | F6S8M0 | 0.894 | 0.0586 |
| CD47 | Q08722 | 0.893 | 0.0573 |
| IQCE | A0A087WX45 | 0.893 | 0.057 |
| KRR1 | Q13601 | 0.891 | 0.0568 |
| MID1 | O15344 | 0.891 | 0.0563 |
| YY1AP1 | Q9H869 | 0.893 | 0.0563 |
| RCC1L | Q96I51 | 0.891 | 0.0561 |
| RAB11FIP1 | Q6WKZ4 | 0.89 | 0.0541 |
| YBX1 | P67809 | 0.893 | 0.0539 |
| LRRC8E | Q6NSJ5 | 0.888 | 0.0538 |
| MYO10 | A0A0A0MQX1 | 0.893 | 0.0535 |
| ZNF692 | Q9BU19 | 0.894 | 0.0532 |
| JUP | P14923 | 0.894 | 0.0529 |
| ANAPC4 | Q9UJX5 | 0.89 | 0.0528 |
| ANKRD26 | E7ESJ3 | 0.894 | 0.0523 |
| ZNF627 | Q7L945 | 0.89 | 0.052 |
| RPLP0 | P05388 | 0.894 | 0.051 |
| CTC1 | Q2NKJ3 | 0.893 | 0.0508 |
| GTF2F1 | P35269 | 0.895 | 0.0506 |
| MAP4K2 | Q12851 | 0.892 | 0.0506 |
| GGH | Q92820 | 0.889 | 0.0499 |
| TRMT10A | Q8TBZ6 | 0.894 | 0.0491 |
| CLIP1 | P30622 | 0.892 | 0.0482 |

| | | | |
|---|---|---|---|
| TRIM47 | Q96LD4 | 0.891 | 0.0481 |
| PDCD7 | Q8N8D1 | 0.895 | 0.0466 |
| CC2D1A | Q6P1N0 | 0.891 | 0.0456 |
| UBQLN4 | Q9NRR5 | 0.893 | 0.0451 |
| MSL2 | Q9HCI7 | 0.895 | 0.0448 |
| SCYL1 | Q96KG9 | 0.888 | 0.0444 |
| LUC7L3 | J3KPP4 | 0.894 | 0.044 |
| STOM | P27105 | 0.893 | 0.0433 |
| EIF2AK2 | P19525 | 0.891 | 0.0431 |
| BAIAP2.1 | I3L4C2 | 0.894 | 0.0429 |
| ORC1 | Q13415 | 0.895 | 0.0425 |
| PSMC2 | P35998 | 0.895 | 0.0423 |
| SRGAP1 | Q7Z6B7 | 0.896 | 0.0423 |
| ATP6V0A1 | Q93050 | 0.894 | 0.0414 |
| TGM3 | Q08188 | 0.893 | 0.0412 |
| ZMYM1 | Q5SVZ6 | 0.895 | 0.0412 |
| THOC5 | Q13769 | 0.896 | 0.0407 |
| KDM3B | Q7LBC6 | 0.89 | 0.0404 |
| NUFIP1 | Q9UHK0 | 0.894 | 0.0403 |
| MYBBP1A | Q9BQG0 | 0.896 | 0.04 |
| REPS1 | Q96D71 | 0.895 | 0.0398 |
| SENP6 | Q9GZR1 | 0.894 | 0.0397 |
| PPP6R2 | O75170 | 0.894 | 0.0391 |
| MAP3K20 | Q9NYL2 | 0.893 | 0.0383 |
| TSEN54 | Q7Z6J9 | 0.892 | 0.0376 |
| DIAPH1 | A0A0G2JH68 | 0.893 | 0.0373 |
| SETD1A | O15047 | 0.895 | 0.0371 |
| ZNF24 | P17028 | 0.894 | 0.037 |
| PACSIN2 | Q9UNF0 | 0.893 | 0.0367 |
| HADH | E9PF18 | 0.892 | 0.0364 |
| POMK | Q9H5K3 | 0.894 | 0.0363 |
| HNRNPH1 | G8JLB6 | 0.893 | 0.0357 |
| NRIP2 | Q9BQI9 | 0.895 | 0.0352 |
| GYG1 | P46976 | 0.896 | 0.0342 |
| HNRNPA1 | P09651 | 0.895 | 0.034 |
| RXRB | P28702 | 0.894 | 0.0334 |
| SF3A3 | Q12874 | 0.893 | 0.0334 |
| RPL3 | P39023 | 0.894 | 0.0333 |
| MTOR | P42345 | 0.895 | 0.0332 |
| OSBPL9 | Q96SU4 | 0.895 | 0.0332 |
| POLR2B | P30876 | 0.892 | 0.033 |
| POM121B | A6NF01 | 0.894 | 0.0329 |
| HSPH1 | Q92598 | 0.894 | 0.0326 |
| MCMBP | Q9BTE3 | 0.896 | 0.0326 |
| GSN | P06396 | 0.894 | 0.0325 |
| RPS6KB2 | Q9UBS0 | 0.894 | 0.0324 |
| NOL9 | Q5SY16 | 0.896 | 0.0321 |

| | | | |
|---|---|---|---|
| SDC4 | P31431 | 0.894 | 0.0321 |
| HDGFL2 | Q7Z4V5 | 0.895 | 0.0317 |
| BRIX1 | Q8TDN6 | 0.895 | 0.0307 |
| HNRNPR | O43390 | 0.894 | 0.0307 |
| SPTY2D1 | Q68D10 | 0.893 | 0.0307 |
| NUP155 | O75694 | 0.894 | 0.0306 |
| EPC2 | Q52LR7 | 0.896 | 0.0305 |
| TAF15 | Q92804 | 0.895 | 0.03 |
| AFF4 | Q9UHB7 | 0.896 | 0.0298 |
| GRSF1 | Q12849 | 0.895 | 0.0293 |
| KIF2A.1 | D6R9M0 | 0.896 | 0.029 |
| PHC2 | Q8IXK0 | 0.896 | 0.0288 |
| WDR83 | Q9BRX9 | 0.896 | 0.0286 |
| SRPK1 | Q96SB4 | 0.895 | 0.0283 |
| SYNCRIP | O60506 | 0.895 | 0.0279 |
| SRSF4 | Q08170 | 0.894 | 0.0269 |
| MYO1D | O94832 | 0.894 | 0.0267 |
| TFG | Q92734 | 0.895 | 0.0261 |
| RING1 | Q06587 | 0.895 | 0.026 |
| HNRNPL | P14866 | 0.893 | 0.0255 |
| TRAP1 | Q12931 | 0.897 | 0.0248 |
| ANKMY2 | Q8IV38 | 0.896 | 0.0247 |
| UBAP2 | Q5T6F2 | 0.894 | 0.0236 |
| ANKRD50 | Q9ULJ7 | 0.896 | 0.0235 |
| HS2ST1 | Q7LGA3 | 0.897 | 0.023 |
| USP53 | Q70EK8 | 0.896 | 0.0229 |
| TRIP12 | Q14669 | 0.896 | 0.0227 |
| RTKN | Q9BST9 | 0.895 | 0.0226 |
| NDUFA10 | E7ESZ7 | 0.895 | 0.0224 |
| MRPL45 | A0A087X2D5 | 0.896 | 0.0223 |
| SLC16A3 | O15427 | 0.895 | 0.0222 |
| NFIX | Q14938 | 0.896 | 0.0219 |
| PNPT1 | Q8TCS8 | 0.897 | 0.0208 |
| ARFGAP1 | Q8N6T3 | 0.896 | 0.0202 |
| CAP1 | Q01518 | 0.897 | 0.0201 |
| CDYL | Q9Y232 | 0.896 | 0.019 |
| MRPS28 | Q9Y2Q9 | 0.895 | 0.0185 |
| YWHAE | P62258 | 0.897 | 0.0169 |
| USP9X | Q93008 | 0.895 | 0.0168 |
| LRWD1 | Q9UFC0 | 0.896 | 0.0167 |
| SBF1 | O95248 | 0.897 | 0.0163 |
| PHLDB1 | Q86UU1 | 0.897 | 0.0161 |
| PA2G4 | Q9UQ80 | 0.895 | 0.0159 |
| RBFA | Q8N0V3 | 0.895 | 0.0157 |
| ASH2L | Q9UBL3 | 0.896 | 0.015 |
| PJA2 | O43164 | 0.896 | 0.0147 |
| AKAP17A | Q02040 | 0.896 | 0.0146 |

| | | | |
|---|---|---|---|
| RBM28 | Q9NW13 | 0.897 | 0.0145 |
| ATXN2 | Q99700 | 0.896 | 0.0141 |
| SNRNP70 | P08621 | 0.896 | 0.0136 |
| TUBB3 | Q13509 | 0.896 | 0.0136 |
| RPP38 | P78345 | 0.896 | 0.0131 |
| PRRC2C | Q9Y520 | 0.896 | 0.0118 |
| SF3A2 | Q15428 | 0.897 | 0.0109 |
| ACOT7 | O00154 | 0.898 | 0.0103 |
| AKAP11 | Q9UKA4 | 0.897 | 0.0102 |
| DDX11 | Q96FC9 | 0.897 | 0.00984 |
| RPL12 | P30050 | 0.897 | 0.0098 |
| ACACB | O00763 | 0.897 | 0.00894 |
| VCL | P18206 | 0.898 | 0.00885 |
| KIF20B | Q96Q89 | 0.898 | 0.00874 |
| TUBB | P07437 | 0.897 | 0.00872 |
| UTP4 | H3BSH7 | 0.898 | 0.0082 |
| RTN4RL2 | Q86UN3 | 0.897 | 0.00713 |
| HNRNPLL | D6W592 | 0.897 | 0.00689 |
| WDR1 | O75083 | 0.898 | 0.00655 |
| CBL | P22681 | 0.897 | 0.00641 |
| NAV1 | Q8NEY1 | 0.897 | 0.0063 |
| CACYBP | Q9HB71 | 0.898 | 0.00626 |
| INF2 | Q27J81 | 0.897 | 0.00586 |
| MRPS9 | P82933 | 0.897 | 0.00575 |
| STK24 | Q9Y6E0 | 0.897 | 0.00564 |
| CIAO1 | O76071 | 0.898 | 0.00558 |
| AHDC1 | Q5TGY3 | 0.898 | 0.00554 |
| POM121 | Q96HA1 | 0.898 | 0.00505 |
| PHF8 | Q9UPP1 | 0.898 | 0.0047 |
| MARCKS | P29966 | 0.898 | 0.00464 |
| PSMC1 | P62191 | 0.898 | 0.00463 |
| BIRC6 | Q9NR09 | 0.898 | 0.00461 |
| IARS2 | Q9NSE4 | 0.898 | 0.00352 |
| SDHA | P31040 | 0.898 | 0.00333 |
| ARF4 | P18085 | 0.898 | 0.00323 |
| PTBP1 | P26599 | 0.898 | 0.00317 |
| POM121C | A8CG34 | 0.898 | 0.00301 |
| AP1B1 | Q10567 | 0.898 | 0.00297 |
| B4GALT7 | Q9UBV7 | 0.898 | 0.00288 |
| TAF1C | Q15572 | 0.898 | 0.00223 |
| OR5K3 | A6NET4 | 0.898 | 0.00191 |
| NOP9 | Q86U38 | 0.898 | 0.00171 |
| EMD | P50402 | 0.898 | 0.00134 |
| ZMYND8 | Q9ULU4 | 0.898 | 0.00126 |
| DRG2 | P55039 | 0.898 | 0.00103 |
| RBM39 | Q14498 | 0.898 | 0.00063 |
| HDLBP | A0A024R4E5 | 0.898 | 3.30E-05 |

| | | | |
|---|---|---|---|
| ZNF407 | Q9C0G0 | 0.898 | -0.000819 |
| SND1 | Q7KZF4 | 0.898 | -0.000823 |
| RMDN3 | Q96TC7 | 0.898 | -0.000965 |
| DNPEP | E7ETB3 | 0.898 | -0.00101 |
| PACSIN3 | Q9UKS6 | 0.898 | -0.00126 |
| TBC1D10B | Q4KMP7 | 0.898 | -0.00138 |
| CAD | P27708 | 0.898 | -0.00156 |
| DSG2 | Q14126 | 0.898 | -0.00234 |
| EPS15L1 | Q9UBC2 | 0.898 | -0.00241 |
| ZNF688 | P0C7X2 | 0.898 | -0.00327 |
| DDX3X | A0A0D9SF53 | 0.898 | -0.00432 |
| CHD5 | Q8TDI0 | 0.898 | -0.00499 |
| PDIA6 | Q15084 | 0.898 | -0.00524 |
| ABCF2 | Q9UG63 | 0.897 | -0.00613 |
| GTF3C5 | Q9Y5Q8 | 0.898 | -0.00723 |
| NLRP10 | Q86W26 | 0.897 | -0.00724 |
| YES1 | J3QRU1 | 0.898 | -0.00758 |
| ZBED6 | P86452 | 0.898 | -0.00763 |
| TOP1 | P11387 | 0.897 | -0.00824 |
| SBNO1 | A3KN83 | 0.897 | -0.0086 |
| RRP12 | Q5JTH9 | 0.897 | -0.00886 |
| CUL5 | Q93034 | 0.897 | -0.00912 |
| ERICH1 | Q86X53 | 0.898 | -0.00942 |
| LRSAM1 | Q6UWE0 | 0.897 | -0.00975 |
| TAF7 | Q15545 | 0.897 | -0.00993 |
| PRPF8 | Q6P2Q9 | 0.898 | -0.0103 |
| MTPAP | Q9NVV4 | 0.897 | -0.0108 |
| ZNFX1 | Q9P2E3 | 0.896 | -0.0118 |
| GGA3 | Q9NZ52 | 0.896 | -0.0141 |
| REXO1 | Q8N1G1 | 0.897 | -0.0141 |
| ASS1 | P00966 | 0.897 | -0.0156 |
| SRPRB | Q9Y5M8 | 0.896 | -0.0174 |
| TUBG1 | P23258 | 0.896 | -0.0174 |
| LDHB | P07195 | 0.897 | -0.0178 |
| BAIAP2L1 | Q9UHR4 | 0.896 | -0.0179 |
| TMPO.1 | P42167 | 0.896 | -0.0182 |
| PPP2R1A | P30153 | 0.896 | -0.0183 |
| TPD52L2 | O43399 | 0.897 | -0.0183 |
| AP1M2 | Q9Y6Q5 | 0.894 | -0.0184 |
| CKAP2 | Q8WWK9 | 0.896 | -0.0185 |
| DPYSL3 | Q14195 | 0.897 | -0.0186 |
| ANKRD12 | Q6UB98 | 0.897 | -0.0189 |
| EMG1 | Q92979 | 0.895 | -0.0197 |
| RNMT | O43148 | 0.897 | -0.0198 |
| EIF2S3 | P41091 | 0.896 | -0.02 |
| C1QBP | Q07021 | 0.896 | -0.0201 |
| MELK | Q14680 | 0.896 | -0.0202 |

| | | | |
|---|---|---|---|
| GGA1 | Q9UJY5 | 0.894 | -0.0207 |
| DEPDC1B | Q8WUY9 | 0.897 | -0.0211 |
| PPP6R1 | Q9UPN7 | 0.896 | -0.0215 |
| CAB39 | Q9Y376 | 0.897 | -0.0216 |
| IQGAP3 | Q86VI3 | 0.897 | -0.0241 |
| RAB11FIP2 | Q7L804 | 0.896 | -0.0245 |
| AP3D1 | O14617 | 0.896 | -0.025 |
| EIF2S3L | Q2VIR3 | 0.895 | -0.025 |
| CCNK | O75909 | 0.896 | -0.0258 |
| BTRC | Q9Y297 | 0.895 | -0.0271 |
| CHEK1 | E7EPP6 | 0.893 | -0.0272 |
| ESS2 | Q96DF8 | 0.895 | -0.0277 |
| GNPAT | O15228 | 0.895 | -0.0278 |
| NOP2 | P46087 | 0.896 | -0.028 |
| ATP1B1 | P05026 | 0.896 | -0.0283 |
| CLK2 | P49760 | 0.894 | -0.0283 |
| DTX3L | Q8TDB6 | 0.895 | -0.0287 |
| TRAM1 | Q15629 | 0.895 | -0.0287 |
| HM13 | Q8TCT9 | 0.896 | -0.029 |
| NUDT19 | A8MXV4 | 0.896 | -0.0297 |
| CDC23 | Q9UJX2 | 0.892 | -0.0298 |
| ASMTL | O95671 | 0.896 | -0.0304 |
| PTMA | H7C2N1 | 0.896 | -0.0305 |
| NFKB2 | Q00653 | 0.896 | -0.0306 |
| UGGT1 | Q9NYU2 | 0.893 | -0.0308 |
| PDIA3.1 | H7BZJ3 | 0.895 | -0.031 |
| RFC3 | P40938 | 0.892 | -0.0318 |
| SNRPD3 | P62318 | 0.891 | -0.032 |
| ARHGEF16 | Q5VV41 | 0.896 | -0.0325 |
| GNL2 | Q13823 | 0.894 | -0.0336 |
| EIF3C | Q99613 | 0.893 | -0.0339 |
| C2CD4C | Q8TF44 | 0.895 | -0.0344 |
| CHPF | Q8IZ52 | 0.89 | -0.0349 |
| MIOS | Q9NXC5 | 0.892 | -0.035 |
| RFC4 | P35249 | 0.892 | -0.035 |
| YIPF5 | Q969M3 | 0.894 | -0.0351 |
| CD63 | P08962 | 0.896 | -0.0357 |
| LENG1 | Q96BZ8 | 0.893 | -0.0357 |
| FUBP1.1 | C9JSZ1 | 0.895 | -0.0359 |
| FERMT1 | Q9BQL6 | 0.891 | -0.0363 |
| SKP1 | P63208 | 0.895 | -0.0365 |
| EIF2S1 | P05198 | 0.893 | -0.0369 |
| SENP3 | Q9H4L4 | 0.895 | -0.0377 |
| EIF3I | Q13347 | 0.895 | -0.0378 |
| HNRNPH2 | P55795 | 0.894 | -0.0385 |
| GTF3C2 | Q8WUA4 | 0.895 | -0.0386 |
| KATNB1 | Q9BVA0 | 0.893 | -0.0386 |

| | | | |
|---|---|---|---|
| DKC1 | O60832 | 0.894 | -0.0388 |
| RTRAF | Q9Y224 | 0.895 | -0.0396 |
| EPRS | P07814 | 0.89 | -0.0401 |
| RAB7A | P51149 | 0.895 | -0.0403 |
| KDM2B | Q8NHM5 | 0.894 | -0.0405 |
| SMG9 | Q9H0W8 | 0.893 | -0.0413 |
| USO1 | O60763 | 0.895 | -0.0414 |
| AARS | P49588 | 0.895 | -0.0421 |
| AGAP1.1 | Q9UPQ3 | 0.896 | -0.0427 |
| MLLT6 | A0A087WW39 | 0.893 | -0.0428 |
| EPB41L5 | Q9HCM4 | 0.889 | -0.0433 |
| THOC2 | Q8NI27 | 0.896 | -0.0434 |
| SPINT2 | O43291 | 0.894 | -0.0436 |
| AKAP12 | Q02952 | 0.894 | -0.044 |
| CHERP.1 | J3QK89 | 0.895 | -0.0442 |
| PNPLA6 | Q8IY17 | 0.895 | -0.0443 |
| GTF2H3 | Q13889 | 0.891 | -0.0444 |
| SMARCA5 | O60264 | 0.894 | -0.0445 |
| MYO9B | Q13459 | 0.894 | -0.0453 |
| CDK12 | Q9NYV4 | 0.893 | -0.0458 |
| CERS2 | Q96G23 | 0.89 | -0.0459 |
| AHR | P35869 | 0.887 | -0.046 |
| AKAP1 | Q92667 | 0.892 | -0.0461 |
| MKRN2 | Q9H000 | 0.891 | -0.0468 |
| CPSF1 | Q10570 | 0.895 | -0.0477 |
| PRPF4 | O43172 | 0.893 | -0.0477 |
| SEC31A | D6REX3 | 0.894 | -0.0481 |
| TAF6L | Q9Y6J9 | 0.892 | -0.0482 |
| GTF3C4 | Q9UKN8 | 0.894 | -0.0486 |
| ANXA11 | P50995 | 0.895 | -0.0487 |
| CDK1 | A0A024QZP7 | 0.89 | -0.0487 |
| ZNF431 | Q8TF32 | 0.891 | -0.049 |
| PTPN20 | Q4JDL3 | 0.888 | -0.0492 |
| ZNF638 | Q14966 | 0.894 | -0.0498 |
| PDE4A | P27815 | 0.89 | -0.0499 |
| SMU1 | Q2TAY7 | 0.893 | -0.0503 |
| CDKN2AIP | Q9NXV6 | 0.891 | -0.0504 |
| SNX1 | Q13596 | 0.894 | -0.0505 |
| MAP3K21 | Q5TCX8 | 0.893 | -0.0509 |
| PRICKLE3 | O43900 | 0.891 | -0.0509 |
| NUP98 | P52948 | 0.889 | -0.051 |
| GRK2 | P25098 | 0.887 | -0.0512 |
| SIK3 | H0Y4E8 | 0.891 | -0.0512 |
| ERICH5 | Q6P6B1 | 0.893 | -0.0513 |
| NUSAP1 | Q9BXS6 | 0.891 | -0.0514 |
| CSPP1 | Q1MSJ5 | 0.891 | -0.0515 |
| FAM208B | Q5VWN6 | 0.89 | -0.0515 |

| | | | |
|---|---|---|---|
| CLASP2.1 | E3W994 | 0.894 | -0.0518 |
| GTF2I | P78347 | 0.888 | -0.0521 |
| LTA4H | P09960 | 0.894 | -0.0522 |
| TMEM43 | Q9BTV4 | 0.889 | -0.0522 |
| ZC3H14 | G3V4R5 | 0.895 | -0.0527 |
| EFR3A | A0A1B0GUZ7 | 0.888 | -0.0529 |
| EXOC1 | Q9NV70 | 0.888 | -0.0542 |
| CPNE3 | O75131 | 0.893 | -0.0543 |
| TARS | P26639 | 0.893 | -0.0554 |
| SHTN1 | A0MZ66 | 0.892 | -0.0555 |
| ZNF566 | Q969W8 | 0.892 | -0.0556 |
| SLAIN2.1 | D6RIF6 | 0.889 | -0.0559 |
| RNF2 | Q99496 | 0.891 | -0.0561 |
| USP5 | P45974 | 0.894 | -0.0563 |
| EIF5A | P63241 | 0.886 | -0.0564 |
| HK2 | P52789 | 0.893 | -0.0564 |
| TPM2 | Q5TCU8 | 0.89 | -0.0566 |
| CPSF7 | Q8N684 | 0.889 | -0.0574 |
| EIF5B | A0A087WUT6 | 0.894 | -0.0577 |
| KAT2A | Q92830 | 0.892 | -0.0577 |
| SCAF1 | Q9H7N4 | 0.895 | -0.0579 |
| ADD1 | P35611 | 0.89 | -0.0582 |
| FAM133A | Q8N9E0 | 0.89 | -0.0585 |
| SGPL1 | O95470 | 0.891 | -0.0587 |
| NAT10 | Q9H0A0 | 0.894 | -0.059 |
| WDR48 | Q8TAF3 | 0.891 | -0.0595 |
| KIF18A | Q8NI77 | 0.893 | -0.0597 |
| AP2A1 | O95782 | 0.892 | -0.0606 |
| ZNF592 | Q92610 | 0.894 | -0.0606 |
| DYNC1H1 | Q14204 | 0.887 | -0.0609 |
| ETF1 | P62495 | 0.883 | -0.0611 |
| MED27 | Q6P2C8 | 0.888 | -0.0625 |
| DDX56 | Q9NY93 | 0.888 | -0.0636 |
| EPB41L1.1 | H7C2K6 | 0.891 | -0.0648 |
| SRPRA | P08240 | 0.887 | -0.0651 |
| RECQL | P46063 | 0.89 | -0.0653 |
| NPAT | Q14207 | 0.89 | -0.0654 |
| SNRPA1 | P09661 | 0.886 | -0.066 |
| FMNL3 | Q8IVF7 | 0.889 | -0.0662 |
| UNC5D | C9J2B6 | 0.893 | -0.0663 |
| ZNF644 | Q9H582 | 0.892 | -0.0664 |
| WAPL | Q7Z5K2 | 0.89 | -0.0669 |
| MPP6 | Q9NZW5 | 0.891 | -0.0671 |
| NOTCH2 | Q04721 | 0.889 | -0.0672 |
| BAIAP2 | Q9UQB8 | 0.889 | -0.0675 |
| SLC5A6 | Q9Y289 | 0.89 | -0.0676 |
| FDPS | P14324 | 0.894 | -0.0677 |

| | | | |
|---|---|---|---|
| HNRNPU | Q00839 | 0.887 | -0.0679 |
| TOR4A | Q9NXH8 | 0.889 | -0.0679 |
| NKAP | Q8N5F7 | 0.889 | -0.0681 |
| GTF3C5.1 | Q5T7U1 | 0.89 | -0.0694 |
| RPS18 | P62269 | 0.891 | -0.0705 |
| SSRP1 | Q08945 | 0.885 | -0.0707 |
| UBE2S | Q16763 | 0.886 | -0.0707 |
| CYFIP1 | Q7L576 | 0.882 | -0.071 |
| CARS.1 | E9PLP0 | 0.886 | -0.0711 |
| TXLNG | Q9NUQ3 | 0.889 | -0.0711 |
| SMAD3 | P84022 | 0.889 | -0.0712 |
| WDR17 | Q8IZU2 | 0.891 | -0.0721 |
| MAGED2 | Q9UNF1 | 0.883 | -0.0725 |
| SNRNP200 | O75643 | 0.892 | -0.0735 |
| CDC123 | O75794 | 0.881 | -0.0736 |
| ARGLU1 | Q9NWB6 | 0.884 | -0.0737 |
| YWHAZ | P63104 | 0.893 | -0.074 |
| SMARCE1 | Q969G3 | 0.891 | -0.0741 |
| DDX55 | Q8NHQ9 | 0.889 | -0.0742 |
| GHDC | Q8N2G8 | 0.884 | -0.0745 |
| KATNA1 | O75449 | 0.888 | -0.0746 |
| ARMC5 | J3KQ26 | 0.887 | -0.0747 |
| CDH3 | P22223 | 0.889 | -0.0751 |
| EHHADH | Q08426 | 0.889 | -0.0752 |
| PUM3 | Q15397 | 0.893 | -0.076 |
| RAC2 | P15153 | 0.892 | -0.076 |
| FUBP3 | Q96I24 | 0.89 | -0.0761 |
| NSD1 | Q96L73 | 0.886 | -0.0763 |
| EIF5 | P55010 | 0.89 | -0.0765 |
| MARS | P56192 | 0.886 | -0.0773 |
| SLC1A5 | Q15758 | 0.89 | -0.0775 |
| DHX8 | Q14562 | 0.893 | -0.0777 |
| DICER1 | Q9UPY3 | 0.888 | -0.0784 |
| SFXN4 | Q6P4A7 | 0.892 | -0.0788 |
| GTPBP2 | Q9BX10 | 0.891 | -0.0789 |
| SRRM1 | A9Z1X7 | 0.89 | -0.079 |
| PUM2 | A0A0A0MR59 | 0.89 | -0.0793 |
| MYOF | Q9NZM1 | 0.889 | -0.0797 |
| NACA | E9PAV3 | 0.888 | -0.0797 |
| ZNF318 | Q5VUA4 | 0.892 | -0.08 |
| MELTF | P08582 | 0.891 | -0.0804 |
| EXOSC3 | Q9NQT5 | 0.887 | -0.0809 |
| EWSR1.1 | H7BY36 | 0.884 | -0.0816 |
| ZNF770 | Q6IQ21 | 0.885 | -0.0823 |
| TTI1 | O43156 | 0.886 | -0.0828 |
| MTCL1 | Q9Y4B5 | 0.891 | -0.0836 |
| RABL6 | Q3YEC7 | 0.893 | -0.0837 |

| | | | |
|---|---|---|---|
| WDR6 | Q9NNW5 | 0.887 | -0.0845 |
| PRKCI | P41743 | 0.89 | -0.0846 |
| VAPA | Q9P0L0 | 0.883 | -0.0849 |
| SPATA5 | Q8NB90 | 0.879 | -0.0852 |
| RDH13 | Q8NBN7 | 0.891 | -0.0853 |
| P4HA2 | O15460 | 0.882 | -0.0856 |
| RCOR3 | Q9P2K3 | 0.888 | -0.086 |
| KIF14 | Q15058 | 0.879 | -0.0863 |
| CALD1 | E9PGZ1 | 0.892 | -0.0881 |
| GCN1 | Q92616 | 0.883 | -0.0883 |
| FAT1 | Q14517 | 0.88 | -0.0887 |
| CAST | A0A0C4DGB5 | 0.89 | -0.089 |
| LAMC2 | Q13753 | 0.883 | -0.0895 |
| HSPD1 | P10809 | 0.89 | -0.0898 |
| SLC25A3.1 | F8VVM2 | 0.879 | -0.0898 |
| TUBB6 | Q9BUF5 | 0.882 | -0.0898 |
| ZNF768 | Q9H5H4 | 0.891 | -0.0902 |
| OGT | O15294 | 0.883 | -0.0903 |
| USP16 | Q9Y5T5 | 0.89 | -0.0909 |
| PKM | P14618 | 0.89 | -0.0912 |
| SEC23B | Q15437 | 0.88 | -0.0923 |
| FARSA | K7ER00 | 0.887 | -0.0927 |
| SNW1 | Q13573 | 0.886 | -0.0927 |
| PARP2 | Q9UGN5 | 0.884 | -0.093 |
| EIF4ENIF1 | Q9NRA8 | 0.875 | -0.0935 |
| RBM17 | Q96I25 | 0.883 | -0.0935 |
| SENP1 | Q9P0U3 | 0.885 | -0.0941 |
| POLR3A | O14802 | 0.89 | -0.0943 |
| ANKHD1-EIF4EBP3 | H7C0V5 | 0.878 | -0.0945 |
| AKAP8L | Q9ULX6 | 0.873 | -0.0946 |
| TOMM34 | Q15785 | 0.883 | -0.0947 |
| STAU1.1 | Q5JW28 | 0.883 | -0.0949 |
| WASL | O00401 | 0.88 | -0.0949 |
| RACGAP1 | Q9H0H5 | 0.886 | -0.0953 |
| UMPS | P11172 | 0.888 | -0.0953 |
| BAZ1A | Q9NRL2 | 0.89 | -0.0954 |
| KLF16 | Q9BXK1 | 0.891 | -0.0954 |
| BAP1 | Q92560 | 0.887 | -0.0955 |
| INTS14 | B4DJL6 | 0.875 | -0.0965 |
| EIF3H | B3KS98 | 0.89 | -0.0969 |
| TRRAP | F2Z2U4 | 0.891 | -0.0969 |
| ZCCHC6 | Q5VYS8 | 0.884 | -0.0975 |
| METAP1 | P53582 | 0.888 | -0.0995 |
| TMEM183A | Q8IXX5 | 0.882 | -0.0996 |
| DDX47 | Q9H0S4 | 0.887 | -0.0998 |
| ZCCHC8 | Q6NZY4 | 0.875 | -0.0999 |
| SGTA | O43765 | 0.889 | -0.1 |

| | | | |
|---|---|---|---|
| TAOK2 | Q9UL54 | 0.89 | -0.1 |
| CCDC88C | Q9P219 | 0.883 | -0.101 |
| CKAP5 | Q14008 | 0.888 | -0.101 |
| ALCAM | Q13740 | 0.89 | -0.102 |
| BTBD9 | Q96Q07 | 0.885 | -0.102 |
| EML4 | B5MBZ0 | 0.883 | -0.102 |
| NKRF | O15226 | 0.875 | -0.102 |
| ELL | P55199 | 0.873 | -0.104 |
| ENO1 | P06733 | 0.889 | -0.104 |
| ISOC1 | Q96CN7 | 0.885 | -0.104 |
| POP1 | Q99575 | 0.888 | -0.105 |
| TUBA1A | Q71U36 | 0.881 | -0.105 |
| CCNA2 | P20248 | 0.882 | -0.106 |
| MARK3 | P27448 | 0.884 | -0.106 |
| IDE | P14735 | 0.887 | -0.107 |
| NCBP1 | Q09161 | 0.878 | -0.107 |
| WRNIP1 | Q96S55 | 0.885 | -0.107 |
| FUBP1 | Q96AE4 | 0.887 | -0.108 |
| MDN1 | Q9NU22 | 0.876 | -0.108 |
| SKI | P12755 | 0.881 | -0.108 |
| TDP2 | O95551 | 0.889 | -0.108 |
| AKT2 | P31751 | 0.884 | -0.109 |
| ATL3 | Q6DD88 | 0.889 | -0.109 |
| EPS8 | Q12929 | 0.886 | -0.109 |
| NFATC1 | O95644 | 0.887 | -0.109 |
| TUBB4B | P68371 | 0.877 | -0.109 |
| UXS1 | Q8NBZ7 | 0.883 | -0.109 |
| RBM22 | Q9NW64 | 0.881 | -0.11 |
| UBA2 | Q9UBT2 | 0.89 | -0.11 |
| FKBP8 | Q14318 | 0.881 | -0.111 |
| KIAA0391 | O15091 | 0.879 | -0.111 |
| SLC20A2 | Q08357 | 0.885 | -0.111 |
| FARSB | Q9NSD9 | 0.885 | -0.112 |
| TMEM30A | Q9NV96 | 0.883 | -0.113 |
| DEPDC1 | Q5TB30 | 0.889 | -0.114 |
| TNPO1 | Q92973 | 0.887 | -0.114 |
| ARHGAP26 | Q9UNA1 | 0.876 | -0.116 |
| MCTP2 | Q6DN12 | 0.89 | -0.116 |
| NOM1 | Q5C9Z4 | 0.89 | -0.117 |
| DEK | P35659 | 0.888 | -0.118 |
| TP53BP1 | Q12888 | 0.872 | -0.118 |
| MACF1.1 | Q9UPN3 | 0.888 | -0.119 |
| PDIA3 | P30101 | 0.882 | -0.119 |
| CALR | P27797 | 0.887 | -0.12 |
| CDK6 | Q00534 | 0.883 | -0.12 |
| NMT1 | P30419 | 0.882 | -0.12 |
| ZC3H11A | O75152 | 0.881 | -0.121 |

| | | | |
|---|---|---|---|
| POGLUT1 | Q8NBL1 | 0.882 | -0.122 |
| BAZ1B | Q9UIG0 | 0.883 | -0.123 |
| PRKD2 | Q9BZL6 | 0.874 | -0.123 |
| SNX27 | Q96L92 | 0.887 | -0.123 |
| STT3A | P46977 | 0.882 | -0.123 |
| C2CD5 | Q86YS7 | 0.886 | -0.124 |
| ELMSAN1 | A0A1C7CYX1 | 0.889 | -0.124 |
| MAPKAP1 | Q9BPZ7 | 0.884 | -0.125 |
| RBBP5 | Q15291 | 0.874 | -0.125 |
| GPD2 | P43304 | 0.885 | -0.126 |
| MAOB | P27338 | 0.887 | -0.126 |
| SAMD9 | Q5K651 | 0.873 | -0.127 |
| AIFM1 | O95831 | 0.879 | -0.129 |
| EHD1 | A0A024R571 | 0.867 | -0.13 |
| MARK2 | E9PC69 | 0.88 | -0.131 |
| UTP18 | Q9Y5J1 | 0.88 | -0.131 |
| CCDC124 | Q96CT7 | 0.884 | -0.132 |
| MRPS34 | C9JJ19 | 0.879 | -0.132 |
| ZBTB2 | Q8N680 | 0.888 | -0.132 |
| ARHGEF2 | V9GYM8 | 0.887 | -0.133 |
| LRRC59 | Q96AG4 | 0.879 | -0.133 |
| PHF20L1 | A8MW92 | 0.887 | -0.133 |
| RUVBL2 | Q9Y230 | 0.859 | -0.133 |
| AUH | Q13825 | 0.875 | -0.134 |
| FXR2 | P51116 | 0.87 | -0.134 |
| GPATCH8 | Q9UKJ3 | 0.891 | -0.134 |
| HMGCS1 | Q01581 | 0.876 | -0.134 |
| PRNP | P04156 | 0.886 | -0.134 |
| RBM15 | Q96T37 | 0.883 | -0.134 |
| ATP5MF-PTCD1 | G3V325 | 0.868 | -0.135 |
| BMS1 | Q14692 | 0.887 | -0.135 |
| DENND4B | O75064 | 0.885 | -0.135 |
| TXNL1 | O43396 | 0.875 | -0.135 |
| DNAJC11 | Q9NVH1 | 0.868 | -0.136 |
| SH3BP1 | Q9Y3L3 | 0.862 | -0.136 |
| TAB1 | Q15750 | 0.873 | -0.136 |
| ARHGAP10 | A1A4S6 | 0.883 | -0.137 |
| PRC1 | O43663 | 0.883 | -0.137 |
| PRPF4B | Q13523 | 0.889 | -0.137 |
| PLD1 | Q13393 | 0.882 | -0.138 |
| SRGAP2 | O75044 | 0.888 | -0.138 |
| KTN1 | Q86UP2 | 0.884 | -0.139 |
| PRR14L | Q5THK1 | 0.881 | -0.139 |
| ARHGEF3 | E9PG37 | 0.876 | -0.14 |
| DSC2 | Q02487 | 0.888 | -0.14 |
| AKAP2 | Q9Y2D5 | 0.88 | -0.141 |
| FNBP4 | Q8N3X1 | 0.887 | -0.141 |

| | | | |
|---|---|---|---|
| MTHFD2 | P13995 | 0.882 | -0.141 |
| PPIL2 | Q13356 | 0.872 | -0.141 |
| TSPYL1 | Q9H0U9 | 0.883 | -0.141 |
| ZNF451 | Q9Y4E5 | 0.885 | -0.141 |
| LARS2 | Q15031 | 0.885 | -0.142 |
| ORC3 | Q9UBD5 | 0.866 | -0.142 |
| PAPOLA | P51003 | 0.879 | -0.142 |
| PARP1 | P09874 | 0.884 | -0.142 |
| RBM33 | H0Y3K4 | 0.88 | -0.142 |
| SRRM2 | Q9UQ35 | 0.871 | -0.142 |
| ZNF791 | Q3KP31 | 0.88 | -0.142 |
| MRPL38 | Q96DV4 | 0.855 | -0.143 |
| REPIN1 | Q9BWE0 | 0.884 | -0.143 |
| SECISBP2L | Q93073 | 0.885 | -0.143 |
| ARAP1 | Q96P48 | 0.882 | -0.144 |
| ARID1B | Q8NFD5 | 0.873 | -0.144 |
| ATP2A2 | P16615 | 0.866 | -0.144 |
| ERLIN1 | O75477 | 0.883 | -0.144 |
| STIP1 | P31948 | 0.883 | -0.144 |
| EXOSC9 | Q06265 | 0.863 | -0.145 |
| HYKK | A2RU49 | 0.884 | -0.145 |
| NCAPD3 | P42695 | 0.884 | -0.145 |
| NOS1AP | O75052 | 0.875 | -0.145 |
| SLMAP | Q14BN4 | 0.873 | -0.145 |
| EFCAB14 | O75071 | 0.877 | -0.146 |
| EIF2B2 | P49770 | 0.867 | -0.146 |
| FASTKD2 | Q9NYY8 | 0.881 | -0.146 |
| HSPA9 | P38646 | 0.875 | -0.146 |
| CLPB | H0YGM0 | 0.856 | -0.147 |
| DDX54 | Q8TDD1 | 0.886 | -0.147 |
| OAT | P04181 | 0.884 | -0.147 |
| OSBPL3 | Q9H4L5 | 0.886 | -0.147 |
| PVR | A0A0C4DG49 | 0.887 | -0.147 |
| ST14 | Q9Y5Y6 | 0.882 | -0.147 |
| YBX3 | P16989 | 0.883 | -0.147 |
| UGP2 | E7EUC7 | 0.89 | -0.148 |
| EIF3E | P60228 | 0.869 | -0.149 |
| AP1M1 | Q9BXS5 | 0.852 | -0.15 |
| BUB3 | O43684 | 0.871 | -0.15 |
| EMC2 | Q15006 | 0.882 | -0.15 |
| ITGA3 | P26006 | 0.882 | -0.15 |
| HSPA8 | P11142 | 0.87 | -0.151 |
| MYO1E | Q12965 | 0.862 | -0.151 |
| PPFIA1 | Q13136 | 0.884 | -0.151 |
| PUM1 | Q5T1Z8 | 0.882 | -0.151 |
| WDR11 | Q9BZH6 | 0.878 | -0.151 |
| NOC2L | Q9Y3T9 | 0.873 | -0.152 |

| | | | |
|---|---|---|---|
| PWP2 | Q15269 | 0.886 | -0.153 |
| MYNN | Q9NPC7 | 0.871 | -0.154 |
| NELFA | A0A0C4DFX9 | 0.87 | -0.154 |
| PIK3CB | P42338 | 0.868 | -0.154 |
| SARS | Q5T5C7 | 0.885 | -0.154 |
| GLA | P06280 | 0.884 | -0.155 |
| MDK | E9PLM6 | 0.859 | -0.155 |
| ZNF567 | Q8N184 | 0.869 | -0.155 |
| APP | P05067 | 0.869 | -0.156 |
| CLASRP | A0A0A0MQS2 | 0.871 | -0.156 |
| MAFF | Q9ULX9 | 0.872 | -0.156 |
| MYO1C.1 | I3L204 | 0.883 | -0.156 |
| SERPINC1 | P01008 | 0.881 | -0.156 |
| ABHD14B | Q96IU4 | 0.872 | -0.157 |
| PANX1 | Q96RD7 | 0.869 | -0.157 |
| PTPN3 | P26045 | 0.884 | -0.157 |
| URI1 | O94763 | 0.867 | -0.157 |
| CHD8 | Q9HCK8 | 0.883 | -0.159 |
| PRIM1 | P49642 | 0.858 | -0.159 |
| PSD4 | Q8NDX1 | 0.874 | -0.159 |
| ARCN1 | B0YIW6 | 0.868 | -0.16 |
| FZD6 | O60353 | 0.876 | -0.16 |
| SEC24C | P53992 | 0.875 | -0.16 |
| TXLNA | P40222 | 0.875 | -0.16 |
| MTHFD1L | B7ZM99 | 0.881 | -0.162 |
| MRM3 | Q9HC36 | 0.88 | -0.163 |
| RCC2 | Q9P258 | 0.872 | -0.163 |
| TP53 | A0A0U1RQC9 | 0.882 | -0.163 |
| NRDE2 | Q9H7Z3 | 0.883 | -0.164 |
| PRIM2 | P49643 | 0.859 | -0.164 |
| SLC19A1 | P41440 | 0.887 | -0.164 |
| AGAP1 | E7EUN2 | 0.885 | -0.165 |
| ASNS | P08243 | 0.88 | -0.165 |
| GARS | P41250 | 0.886 | -0.165 |
| HNRNPK | P61978 | 0.874 | -0.165 |
| ITCH | Q96J02 | 0.85 | -0.165 |
| KPNA2 | P52292 | 0.868 | -0.165 |
| MAP7 | Q14244 | 0.863 | -0.165 |
| PCBP2 | Q15366 | 0.878 | -0.165 |
| RPRD2 | Q5VT52 | 0.878 | -0.165 |
| TAF1 | P21675 | 0.881 | -0.166 |
| SART3 | Q15020 | 0.87 | -0.167 |
| TMA16 | H0Y9X1 | 0.849 | -0.167 |
| GALNT18 | Q6P9A2 | 0.868 | -0.168 |
| HIF1AN | Q9NWT6 | 0.875 | -0.168 |
| TUBA4A | P68366 | 0.865 | -0.168 |
| ALDH9A1 | P49189 | 0.846 | -0.169 |

| | | | |
|---|---|---|---|
| NCAPD2 | Q15021 | 0.86 | -0.169 |
| CBX4 | O00257 | 0.878 | -0.17 |
| COG4 | J3KNI1 | 0.884 | -0.17 |
| LAMA5 | O15230 | 0.879 | -0.17 |
| RBAK | Q9NYW8 | 0.873 | -0.17 |
| SDCBP | G5EA09 | 0.872 | -0.17 |
| UVSSA | Q2YD98 | 0.878 | -0.17 |
| KIF18B | Q86Y91 | 0.886 | -0.171 |
| ABCD3 | P28288 | 0.859 | -0.172 |
| LETMD1 | Q6P1Q0 | 0.858 | -0.172 |
| NAA35 | Q5VZE5 | 0.854 | -0.172 |
| CHD4 | A0A0C4DGG9 | 0.884 | -0.173 |
| MRNIP | Q6NTE8 | 0.869 | -0.173 |
| RAD50 | Q92878 | 0.881 | -0.173 |
| SMC1A | Q14683 | 0.855 | -0.173 |
| ZNF207 | J3QRS9 | 0.885 | -0.173 |
| WDR43 | Q15061 | 0.868 | -0.174 |
| CNN2 | B4DUT8 | 0.883 | -0.175 |
| EGFR | P00533 | 0.878 | -0.175 |
| SNRPG | C9JVQ0 | 0.849 | -0.175 |
| GOLT1B | Q9Y3E0 | 0.869 | -0.176 |
| OSBPL11 | Q9BXB4 | 0.882 | -0.176 |
| SSB | P05455 | 0.871 | -0.176 |
| DHX36 | Q9H2U1 | 0.841 | -0.177 |
| TBL3 | Q12788 | 0.88 | -0.177 |
| THUMPD1 | Q9NXG2 | 0.886 | -0.177 |
| TRMT2A | F2Z2W7 | 0.874 | -0.177 |
| ALYREF | E9PB61 | 0.866 | -0.178 |
| DSC1 | Q08554 | 0.877 | -0.178 |
| NOL11 | Q9H8H0 | 0.854 | -0.178 |
| NSUN4 | Q96CB9 | 0.862 | -0.178 |
| PKN2 | Q16513 | 0.859 | -0.178 |
| TRIM72 | Q6ZMU5 | 0.864 | -0.178 |
| ARAP2 | Q8WZ64 | 0.873 | -0.179 |
| KIF20A | O95235 | 0.868 | -0.179 |
| MYCBP2 | O75592 | 0.865 | -0.179 |
| TTC13 | Q8NBP0 | 0.869 | -0.18 |
| NUF2 | Q9BZD4 | 0.875 | -0.181 |
| TUBA1C | Q9BQE3 | 0.845 | -0.181 |
| CHD6 | Q8TD26 | 0.879 | -0.182 |
| LRPPRC | P42704 | 0.862 | -0.182 |
| PDPR | Q8NCN5 | 0.875 | -0.182 |
| PGK1 | P00558 | 0.883 | -0.184 |
| CFAP20 | Q9Y6A4 | 0.842 | -0.185 |
| CNOT8 | Q9UFF9 | 0.872 | -0.185 |
| AP3B1 | O00203 | 0.881 | -0.186 |
| SF3A1 | Q15459 | 0.856 | -0.186 |

| | | | |
|---|---|---|---|
| DCUN1D3 | Q8IWE4 | 0.872 | -0.187 |
| DLD | P09622 | 0.877 | -0.187 |
| PES1 | O00541 | 0.876 | -0.187 |
| UBA1 | P22314 | 0.877 | -0.187 |
| CNDP2 | Q96KP4 | 0.881 | -0.188 |
| PLD2 | O14939 | 0.848 | -0.188 |
| POLR1B | Q9H9Y6 | 0.877 | -0.188 |
| RPS6KA1 | Q15418 | 0.877 | -0.188 |
| CDC7 | O00311 | 0.871 | -0.189 |
| SAMM50 | Q9Y512 | 0.873 | -0.189 |
| SQSTM1 | Q13501 | 0.884 | -0.189 |
| CUTC | Q9NTM9 | 0.863 | -0.19 |
| FAM120B | F5GY05 | 0.86 | -0.19 |
| PRKDC | P78527 | 0.838 | -0.19 |
| TUBB4A | P04350 | 0.845 | -0.19 |
| PLEKHG5 | O94827 | 0.877 | -0.191 |
| SEC24A | O95486 | 0.868 | -0.191 |
| DARS | P14868 | 0.866 | -0.192 |
| DLAT | P10515 | 0.886 | -0.192 |
| RBM10 | P98175 | 0.872 | -0.192 |
| NAMPT | P43490 | 0.865 | -0.193 |
| PUF60.1 | A0A0J9YVP6 | 0.872 | -0.193 |
| CSNK1E | P49674 | 0.85 | -0.194 |
| TMEM131 | Q92545 | 0.875 | -0.194 |
| GLI4 | P10075 | 0.879 | -0.195 |
| GPKOW | Q92917 | 0.866 | -0.195 |
| STK26 | B4E0Y9 | 0.857 | -0.195 |
| USP48 | Q86UV5 | 0.856 | -0.195 |
| CCNB1 | P14635 | 0.846 | -0.196 |
| GLG1 | Q92896 | 0.87 | -0.196 |
| ALDOA | P04075 | 0.88 | -0.197 |
| CPSF6 | Q16630 | 0.866 | -0.197 |
| EHD4 | Q9H223 | 0.841 | -0.197 |
| MAP7D3 | Q8IWC1 | 0.868 | -0.197 |
| MRPS27 | Q92552 | 0.84 | -0.197 |
| HADHA | P40939 | 0.868 | -0.198 |
| PIP4K2C | Q8TBX8 | 0.838 | -0.198 |
| PTPN11 | Q06124 | 0.88 | -0.198 |
| RBM19 | Q9Y4C8 | 0.866 | -0.198 |
| SPTLC2 | O15270 | 0.865 | -0.198 |
| TPP2 | P29144 | 0.865 | -0.198 |
| SURF4 | O15260 | 0.86 | -0.199 |
| ASPH | Q12797 | 0.877 | -0.2 |
| MAST2 | Q6P0Q8 | 0.876 | -0.2 |
| POLD1 | M0R2B7 | 0.838 | -0.2 |
| CPOX | P36551 | 0.878 | -0.201 |
| LANCL1 | O43813 | 0.863 | -0.201 |

| | | | |
|---|---|---|---|
| NUP93 | Q8N1F7 | 0.859 | -0.201 |
| DNAJA2 | O60884 | 0.836 | -0.202 |
| MED25 | Q71SY5 | 0.869 | -0.202 |
| TICRR | Q7Z2Z1 | 0.884 | -0.202 |
| TIPRL | O75663 | 0.852 | -0.202 |
| ABI1 | Q8IZP0 | 0.841 | -0.203 |
| ANKS1A | Q92625 | 0.868 | -0.203 |
| FAM129B | Q96TA1 | 0.87 | -0.203 |
| MCAT | Q8IVS2 | 0.867 | -0.203 |
| TOMM20 | Q15388 | 0.849 | -0.203 |
| PRR5 | P85299 | 0.873 | -0.204 |
| SIN3A | Q96ST3 | 0.883 | -0.204 |
| SLC25A12 | O75746 | 0.87 | -0.204 |
| CLTA | P09496 | 0.86 | -0.205 |
| GTF2E1 | P29083 | 0.872 | -0.205 |
| EIF3B | P55884 | 0.856 | -0.206 |
| VRK1 | Q99986 | 0.874 | -0.206 |
| ATP6V1H | Q9UI12 | 0.879 | -0.207 |
| ERCC3 | P19447 | 0.879 | -0.207 |
| KIF3A | E9PES4 | 0.854 | -0.207 |
| POLR3D | P05423 | 0.866 | -0.208 |
| GORASP1 | Q9BQQ3 | 0.875 | -0.209 |
| RBM34 | P42696 | 0.875 | -0.209 |
| SNAPC1 | Q16533 | 0.874 | -0.209 |
| MBOAT7 | Q96N66 | 0.876 | -0.21 |
| RPL14 | P50914 | 0.881 | -0.21 |
| SF1 | Q15637 | 0.831 | -0.21 |
| PARG | Q86W56 | 0.868 | -0.211 |
| SYPL1 | Q16563 | 0.853 | -0.211 |
| FARP1 | Q9Y4F1 | 0.866 | -0.212 |
| KAT7 | O95251 | 0.866 | -0.212 |
| NOLC1 | A0A0A0MRM9 | 0.876 | -0.212 |
| PKP4 | Q99569 | 0.839 | -0.212 |
| PSMD12 | O00232 | 0.869 | -0.212 |
| SLC3A2 | J3KPF3 | 0.873 | -0.212 |
| CHMP4C | Q96CF2 | 0.855 | -0.213 |
| HMGB1 | P09429 | 0.881 | -0.213 |
| ZBTB11 | O95625 | 0.874 | -0.213 |
| OGDH | E9PCR7 | 0.876 | -0.214 |
| SETD3 | Q86TU7 | 0.871 | -0.214 |
| USP39 | Q53GS9 | 0.876 | -0.214 |
| EIF3J | O75822 | 0.868 | -0.215 |
| LETM1 | O95202 | 0.865 | -0.215 |
| PDIA4 | P13667 | 0.867 | -0.215 |
| RIOK1 | Q9BRS2 | 0.876 | -0.215 |
| SSR1 | P43307 | 0.837 | -0.216 |
| CNP | P09543 | 0.831 | -0.217 |

| | | | |
|---|---|---|---|
| DDX49 | Q9Y6V7 | 0.877 | -0.217 |
| MAD1L1 | Q9Y6D9 | 0.869 | -0.217 |
| DDX17 | A0A1W2PQ51 | 0.839 | -0.218 |
| HYOU1 | Q9Y4L1 | 0.866 | -0.218 |
| POLRMT | O00411 | 0.877 | -0.218 |
| UBE2O | Q9C0C9 | 0.853 | -0.218 |
| UHRF1 | A0A087WTW0 | 0.869 | -0.218 |
| MORF4L1 | B3KTM8 | 0.864 | -0.219 |
| RAB11A | P62491 | 0.861 | -0.22 |
| STXBP3 | O00186 | 0.845 | -0.22 |
| HSPA5 | P11021 | 0.851 | -0.221 |
| ZNF777 | Q9ULD5 | 0.877 | -0.221 |
| GFPT1 | Q06210 | 0.875 | -0.222 |
| SRRT | Q9BXP5 | 0.831 | -0.222 |
| BCCIP | Q9P287 | 0.866 | -0.223 |
| NFX1 | Q12986 | 0.859 | -0.223 |
| ELMO3 | F8W9E7 | 0.851 | -0.224 |
| G3V3G9 | G3V3G9 | 0.863 | -0.224 |
| C2CD2L | O14523 | 0.855 | -0.225 |
| FAM83F | Q8NEG4 | 0.857 | -0.225 |
| HCFC1.1 | P51610 | 0.849 | -0.225 |
| RP2 | O75695 | 0.878 | -0.225 |
| ZBTB21 | Q9ULJ3 | 0.877 | -0.225 |
| ALDH7A1 | A0A1B0GW77 | 0.875 | -0.226 |
| ASXL2 | Q76L83 | 0.872 | -0.226 |
| CSNK2A1 | P68400 | 0.822 | -0.226 |
| CUL1 | Q13616 | 0.833 | -0.226 |
| PARD6G | Q9BYG4 | 0.871 | -0.226 |
| SMC3 | Q9UQE7 | 0.841 | -0.226 |
| NCL | P19338 | 0.862 | -0.227 |
| PACS2 | Q86VP3 | 0.872 | -0.227 |
| PPFIBP1 | Q86W92 | 0.861 | -0.227 |
| ANXA5 | P08758 | 0.876 | -0.228 |
| DDX24 | Q9GZR7 | 0.875 | -0.228 |
| PALLD | Q8WX93 | 0.86 | -0.228 |
| TFDP1 | Q14186 | 0.862 | -0.228 |
| PPP6R3 | Q5H9R7 | 0.844 | -0.229 |
| ACOT9 | Q9Y305 | 0.872 | -0.23 |
| GAPDH | P04406 | 0.875 | -0.23 |
| HCFC1 | A6NEM2 | 0.859 | -0.23 |
| OSBPL6 | Q9BZF3 | 0.881 | -0.23 |
| PTPN14 | Q15678 | 0.865 | -0.23 |
| P4HB | P07237 | 0.866 | -0.231 |
| RANBP1 | P43487 | 0.872 | -0.231 |
| SRFBP1 | Q8NEF9 | 0.874 | -0.231 |
| IQGAP1 | P46940 | 0.834 | -0.232 |
| PRPSAP2 | O60256 | 0.87 | -0.232 |

| | | | |
|---|---|---|---|
| EIF4A1 | P60842 | 0.871 | -0.233 |
| GLO1 | Q04760 | 0.858 | -0.233 |
| NCDN | Q9UBB6 | 0.863 | -0.233 |
| SCFD1 | Q8WVM8 | 0.876 | -0.233 |
| STX5 | Q13190 | 0.849 | -0.233 |
| AKAP8 | O43823 | 0.859 | -0.234 |
| IARS | P41252 | 0.819 | -0.234 |
| MCM9 | Q9NXL9 | 0.876 | -0.234 |
| FBXO7 | Q9Y3I1 | 0.865 | -0.235 |
| IGF2R | P11717 | 0.864 | -0.235 |
| KANSL1 | A0A0G2JNB1 | 0.852 | -0.235 |
| PGAM1 | P18669 | 0.881 | -0.235 |
| SV2C | Q496J9 | 0.862 | -0.235 |
| ZNF264 | O43296 | 0.851 | -0.235 |
| MAK16 | Q9BXY0 | 0.879 | -0.236 |
| TGOLN2 | O43493 | 0.876 | -0.236 |
| THOP1 | P52888 | 0.873 | -0.236 |
| ARRB2 | P32121 | 0.87 | -0.237 |
| ATP2C1 | B4E2Q0 | 0.849 | -0.237 |
| DDX19A | Q9NUU7 | 0.878 | -0.237 |
| DNAJC16 | Q9Y2G8 | 0.866 | -0.237 |
| EZR | E7EQR4 | 0.838 | -0.237 |
| GSK3B | P49841 | 0.82 | -0.237 |
| L3MBTL3 | Q96JM7 | 0.881 | -0.237 |
| MORF4L2 | Q15014 | 0.869 | -0.237 |
| PIH1D1 | Q9NWS0 | 0.855 | -0.237 |
| BRD4 | O60885 | 0.88 | -0.238 |
| CHD1L | A0A0A0MSH9 | 0.812 | -0.238 |
| COPA | P53621 | 0.818 | -0.238 |
| DHX34 | Q14147 | 0.877 | -0.239 |
| ERP44 | Q9BS26 | 0.849 | -0.239 |
| CDCP1 | Q9H5V8 | 0.87 | -0.24 |
| RIOX1 | Q9H6W3 | 0.881 | -0.24 |
| TRMT1 | Q9NXH9 | 0.874 | -0.24 |
| BPTF | Q12830 | 0.863 | -0.241 |
| FBXO21 | O94952 | 0.883 | -0.241 |
| MSN | P26038 | 0.852 | -0.241 |
| SMAD5 | Q99717 | 0.872 | -0.241 |
| TBC1D5 | Q92609 | 0.868 | -0.241 |
| ACTN1 | P12814 | 0.874 | -0.242 |
| HSPA4 | P34932 | 0.858 | -0.242 |
| ABCB6 | Q9NP58 | 0.866 | -0.243 |
| GANAB.1 | E9PKU7 | 0.852 | -0.243 |
| PRPF38A | Q8NAV1 | 0.871 | -0.243 |
| SUGP1 | Q8IWZ8 | 0.838 | -0.243 |
| ATP5F1A | P25705 | 0.866 | -0.244 |
| NFXL1 | Q6ZNB6 | 0.865 | -0.244 |

| | | | |
|---|---|---|---|
| WDR3 | Q9UNX4 | 0.88 | -0.244 |
| HNRNPUL1 | Q9BUJ2 | 0.833 | -0.245 |
| NVL | O15381 | 0.869 | -0.245 |
| PELO | Q9BRX2 | 0.842 | -0.246 |
| UBE3A | Q05086 | 0.858 | -0.246 |
| XIAP | P98170 | 0.874 | -0.246 |
| ABR | A0A1C7CYZ0 | 0.864 | -0.247 |
| BAG3 | O95817 | 0.865 | -0.247 |
| DDX31 | Q9H8H2 | 0.871 | -0.247 |
| TADA2B | Q86TJ2 | 0.848 | -0.247 |
| ZNF512B | Q96KM6 | 0.877 | -0.247 |
| TXNDC5 | Q8NBS9 | 0.852 | -0.248 |
| DDX23 | Q9BUQ8 | 0.874 | -0.249 |
| MT-CYB | P00156 | 0.864 | -0.249 |
| PALD1 | Q9ULE6 | 0.828 | -0.249 |
| U2AF2 | P26368 | 0.871 | -0.249 |
| C1orf226 | A1L170 | 0.801 | -0.25 |
| NECTIN3 | Q9NQS3 | 0.838 | -0.25 |
| DLG1.1 | Q12959 | 0.835 | -0.251 |
| RARS | P54136 | 0.852 | -0.251 |
| RDX | P35241 | 0.861 | -0.251 |
| TMEM33 | P57088 | 0.866 | -0.251 |
| EIF3M | Q7L2H7 | 0.863 | -0.252 |
| ERCC2 | P18074 | 0.841 | -0.252 |
| OAS3 | Q9Y6K5 | 0.825 | -0.252 |
| THAP12 | O43422 | 0.881 | -0.252 |
| VPS35 | Q96QK1 | 0.869 | -0.252 |
| ABCE1 | P61221 | 0.826 | -0.253 |
| DGAT1 | O75907 | 0.873 | -0.253 |
| SECISBP2 | Q96T21 | 0.873 | -0.253 |
| INTS6 | Q9UL03 | 0.868 | -0.254 |
| PHF6 | Q8IWS0 | 0.853 | -0.255 |
| MPP7 | Q5T2T1 | 0.854 | -0.257 |
| ANKFY1 | Q9P2R3 | 0.847 | -0.258 |
| OLA1 | J3KQ32 | 0.872 | -0.258 |
| POLG | P54098 | 0.857 | -0.258 |
| HELLS | Q9NRZ9 | 0.865 | -0.259 |
| P3H1 | Q32P28 | 0.849 | -0.259 |
| PER1 | O15534 | 0.87 | -0.259 |
| ZNF668 | Q96K58 | 0.833 | -0.26 |
| SNAPC4 | Q5SXM2 | 0.881 | -0.261 |
| SUCLG1 | P53597 | 0.868 | -0.261 |
| TNFRSF10D | Q9UBN6 | 0.837 | -0.261 |
| YTHDC1 | J3QR07 | 0.829 | -0.261 |
| FKBP4 | Q02790 | 0.871 | -0.262 |
| GAA | P10253 | 0.869 | -0.262 |
| KPNA6 | O60684 | 0.803 | -0.262 |

| | | | |
|---|---|---|---|
| SMC2 | O95347 | 0.805 | -0.262 |
| CCNL1 | Q9UK58 | 0.854 | -0.263 |
| ECPAS | J3KN16 | 0.856 | -0.263 |
| P4HA1 | P13674 | 0.817 | -0.263 |
| QARS | P47897 | 0.819 | -0.263 |
| TRMT6 | Q9UJA5 | 0.829 | -0.263 |
| TSR1 | Q2NL82 | 0.863 | -0.263 |
| UTP20 | O75691 | 0.879 | -0.263 |
| DST.1 | Q5T0V7 | 0.851 | -0.265 |
| GYS1 | P13807 | 0.867 | -0.265 |
| PRPF40A | O75400 | 0.876 | -0.265 |
| SUB1 | P53999 | 0.868 | -0.265 |
| WIZ | M0QXA7 | 0.879 | -0.265 |
| YWHAH | Q04917 | 0.852 | -0.265 |
| HADHB | P55084 | 0.846 | -0.266 |
| PIAS1 | O75925 | 0.859 | -0.266 |
| PRKAB1 | Q9Y478 | 0.845 | -0.266 |
| SHB | Q15464 | 0.865 | -0.266 |
| VDAC3 | Q9Y277 | 0.877 | -0.266 |
| CNOT4 | O95628 | 0.879 | -0.267 |
| EIF2B5 | Q13144 | 0.852 | -0.267 |
| PAICS | P22234 | 0.853 | -0.267 |
| PRPF6 | O94906 | 0.871 | -0.267 |
| TRIM32 | Q13049 | 0.865 | -0.267 |
| CAMK2D | E9PF82 | 0.87 | -0.268 |
| IMP4 | Q96G21 | 0.8 | -0.268 |
| SUPV3L1 | Q8IYB8 | 0.86 | -0.268 |
| TMCO1 | J9JIE6 | 0.856 | -0.268 |
| ATP1A1 | P05023 | 0.862 | -0.269 |
| RBPJ | Q06330 | 0.869 | -0.269 |
| SMARCA4.1 | Q9HBD4 | 0.878 | -0.269 |
| OXCT1 | P55809 | 0.874 | -0.27 |
| ABCF1 | Q8NE71 | 0.857 | -0.271 |
| CCDC68 | Q9H2F9 | 0.85 | -0.271 |
| PIK3R2 | O00459 | 0.814 | -0.271 |
| ACAP2 | Q15057 | 0.86 | -0.272 |
| DCAF1 | Q9Y4B6 | 0.814 | -0.272 |
| GFM1 | Q96RP9 | 0.86 | -0.272 |
| HBS1L | Q9Y450 | 0.851 | -0.272 |
| HSP90AA1 | P07900 | 0.862 | -0.272 |
| GTF3C3 | Q9Y5Q9 | 0.876 | -0.273 |
| CLPTM1 | O96005 | 0.864 | -0.274 |
| CD44 | P16070 | 0.856 | -0.275 |
| OR5K4 | A6NMS3 | 0.856 | -0.275 |
| PRDM10 | Q9NQV6 | 0.872 | -0.275 |
| DIS3 | Q9Y2L1 | 0.844 | -0.276 |
| PSMA7 | O14818 | 0.844 | -0.277 |

| | | | |
|---|---|---|---|
| SLAIN2 | Q9P270 | 0.826 | -0.277 |
| STAT1 | P42224 | 0.867 | -0.277 |
| AP2B1 | P63010 | 0.864 | -0.279 |
| DSG1 | Q02413 | 0.87 | -0.279 |
| NCKAP1 | Q9Y2A7 | 0.822 | -0.279 |
| TELO2 | Q9Y4R8 | 0.842 | -0.279 |
| PSME2 | A0A087X1Z3 | 0.874 | -0.28 |
| CUL4B | Q13620 | 0.87 | -0.281 |
| PSMD2 | Q13200 | 0.858 | -0.281 |
| SEC63 | Q9UGP8 | 0.835 | -0.281 |
| EPB41L2 | O43491 | 0.827 | -0.282 |
| GLUD1 | P00367 | 0.863 | -0.283 |
| LARS | Q9P2J5 | 0.789 | -0.283 |
| ZNF324 | O75467 | 0.867 | -0.283 |
| CAPN5 | E7EV01 | 0.863 | -0.284 |
| DST | F6QMI7 | 0.853 | -0.284 |
| IK | Q13123 | 0.863 | -0.284 |
| TUBB2A | Q13885 | 0.837 | -0.284 |
| PPM1G | O15355 | 0.865 | -0.286 |
| CAT | P04040 | 0.869 | -0.287 |
| AP1G2 | O75843 | 0.845 | -0.288 |
| CNPY3 | Q9BT09 | 0.809 | -0.288 |
| COPB1 | P53618 | 0.806 | -0.288 |
| RNPEP | Q9H4A4 | 0.801 | -0.288 |
| BRI3BP | Q8WY22 | 0.782 | -0.29 |
| BYSL | Q13895 | 0.841 | -0.29 |
| RELA | Q04206 | 0.872 | -0.29 |
| U2SURP | O15042 | 0.868 | -0.29 |
| ZWINT | O95229 | 0.831 | -0.29 |
| DPP9 | Q86TI2 | 0.857 | -0.291 |
| HARS | P12081 | 0.85 | -0.291 |
| SORD | Q00796 | 0.871 | -0.291 |
| STRAP | Q9Y3F4 | 0.84 | -0.291 |
| CHST7 | Q9NS84 | 0.806 | -0.292 |
| CORO1B | Q9BR76 | 0.789 | -0.292 |
| ACAD9 | Q9H845 | 0.877 | -0.293 |
| DBR1 | Q9UK59 | 0.793 | -0.293 |
| GMPS | P49915 | 0.869 | -0.293 |
| HSPA4L | O95757 | 0.838 | -0.293 |
| ALB | P02768 | 0.877 | -0.294 |
| DIS3L2 | Q8IYB7 | 0.858 | -0.294 |
| PCK2 | Q16822 | 0.869 | -0.294 |
| PHB2 | Q99623 | 0.871 | -0.294 |
| GRK6 | P43250 | 0.849 | -0.295 |
| HSP90AB1 | P08238 | 0.856 | -0.295 |
| WDR18 | Q9BV38 | 0.813 | -0.295 |
| DBF4 | Q9UBU7 | 0.874 | -0.296 |

| | | | |
|---|---|---|---|
| GNL3L | Q9NVN8 | 0.835 | -0.296 |
| HNRNPU.1 | A0A1W2PP35 | 0.868 | -0.296 |
| KARS | Q15046 | 0.816 | -0.296 |
| PAFAH1B1 | P43034 | 0.8 | -0.296 |
| AGPS | O00116 | 0.799 | -0.297 |
| APMAP | Q9HDC9 | 0.863 | -0.298 |
| DNAJC7 | Q99615 | 0.849 | -0.298 |
| NEDD8 | Q15843 | 0.786 | -0.298 |
| POLR2H | P52434 | 0.798 | -0.298 |
| TFCP2 | Q12800 | 0.839 | -0.298 |
| CARD10 | Q9BWT7 | 0.858 | -0.299 |
| DDX39B | Q13838 | 0.832 | -0.299 |
| FOXC1 | Q12948 | 0.833 | -0.299 |
| RRM2 | P31350 | 0.827 | -0.299 |
| WDR12 | Q9GZL7 | 0.863 | -0.299 |
| ATG9A | Q7Z3C6 | 0.837 | -0.3 |
| KHDRBS1 | Q07666 | 0.806 | -0.3 |
| ARID1A | O14497 | 0.878 | -0.301 |
| PSIP1 | O75475 | 0.848 | -0.301 |
| NUP54 | Q7Z3B4 | 0.873 | -0.302 |
| ZNF518A | Q6AHZ1 | 0.854 | -0.302 |
| PRRC2A | P48634 | 0.793 | -0.303 |
| SDAD1 | Q9NVU7 | 0.862 | -0.303 |
| SLC25A13 | Q9UJS0 | 0.836 | -0.303 |
| SYNGR2 | O43760 | 0.851 | -0.304 |
| AATF | Q9NY61 | 0.88 | -0.305 |
| GRB10 | Q13322 | 0.822 | -0.305 |
| VPS4A | Q9UN37 | 0.815 | -0.305 |
| SF3B1 | O75533 | 0.851 | -0.306 |
| TAF4 | O00268 | 0.87 | -0.306 |
| TSTA3 | Q13630 | 0.87 | -0.307 |
| GPATCH4 | Q5T3I0 | 0.821 | -0.308 |
| TAF10 | Q12962 | 0.828 | -0.308 |
| TIAM1 | Q13009 | 0.843 | -0.309 |
| CDC27 | G5EA36 | 0.833 | -0.31 |
| CORO2A | Q92828 | 0.87 | -0.31 |
| LSM14B | Q9BX40 | 0.833 | -0.31 |
| PGM2 | Q96G03 | 0.867 | -0.31 |
| SMARCA2 | P51531 | 0.875 | -0.31 |
| POLDIP2 | Q9Y2S7 | 0.842 | -0.311 |
| TERF1 | P54274 | 0.869 | -0.311 |
| NTHL1 | P78549 | 0.841 | -0.313 |
| EPB41L1 | A0A0C4DH22 | 0.849 | -0.314 |
| PLCD3 | Q8N3E9 | 0.803 | -0.314 |
| DAXX | Q9UER7 | 0.871 | -0.315 |
| MVB12A | E9PQA6 | 0.872 | -0.315 |
| UGDH | O60701 | 0.86 | -0.315 |

| | | | |
|---|---|---|---|
| SMARCA4 | A0A0A0MT49 | 0.88 | -0.316 |
| MAN1B1 | Q9UKM7 | 0.789 | -0.317 |
| PHC3 | Q8NDX5 | 0.858 | -0.317 |
| PCID2 | Q5JVF3 | 0.774 | -0.318 |
| ARID2 | Q68CP9 | 0.878 | -0.319 |
| CPSF3 | Q9UKF6 | 0.847 | -0.319 |
| MALT1 | Q9UDY8 | 0.86 | -0.319 |
| PREP | P48147 | 0.861 | -0.32 |
| SUCLG2 | Q96I99 | 0.829 | -0.32 |
| TPX2 | Q9ULW0 | 0.86 | -0.32 |
| EEF1D.1 | E9PL71 | 0.853 | -0.322 |
| WAC | A0A0A0MRT2 | 0.857 | -0.322 |
| RAB11B | Q15907 | 0.87 | -0.323 |
| HNRNPA3 | P51991 | 0.841 | -0.324 |
| PI4KA | P42356 | 0.839 | -0.324 |
| RASA3 | Q14644 | 0.822 | -0.324 |
| SAMD1 | Q6SPF0 | 0.853 | -0.325 |
| LONP1 | P36776 | 0.769 | -0.326 |
| PCIF1 | Q9H4Z3 | 0.83 | -0.326 |
| KANSL3 | Q9P2N6 | 0.874 | -0.327 |
| SF3B3 | Q15393 | 0.85 | -0.327 |
| SBF2 | Q86WG5 | 0.859 | -0.328 |
| STIM2 | A0A1W2PRA1 | 0.856 | -0.328 |
| ATP5F1B | P06576 | 0.853 | -0.329 |
| LLGL2 | Q6P1M3 | 0.866 | -0.329 |
| NUB1 | H3BM74 | 0.841 | -0.329 |
| PTPN13 | Q12923 | 0.857 | -0.329 |
| PTGES2 | Q9H7Z7 | 0.85 | -0.331 |
| UBP1 | Q9NZI7 | 0.84 | -0.331 |
| ARHGEF1 | M0QZR4 | 0.815 | -0.332 |
| CDC73 | Q6P1J9 | 0.802 | -0.332 |
| CSTF2 | E7EWR4 | 0.828 | -0.332 |
| EIF3G | O75821 | 0.857 | -0.332 |
| ETV3 | P41162 | 0.839 | -0.332 |
| POLR3B | Q9NW08 | 0.83 | -0.332 |
| USP54 | Q70EL1 | 0.825 | -0.332 |
| MAPK3 | P27361 | 0.846 | -0.333 |
| XPO5 | Q9HAV4 | 0.849 | -0.333 |
| RIOK2 | Q9BVS4 | 0.841 | -0.334 |
| ADSL | P30566 | 0.855 | -0.335 |
| ALDH3A2 | P51648 | 0.858 | -0.335 |
| DHX35 | Q9H5Z1 | 0.826 | -0.335 |
| PPIG | Q13427 | 0.859 | -0.335 |
| ZNF460 | Q14592 | 0.857 | -0.335 |
| CCAR2 | Q8N163 | 0.778 | -0.336 |
| MACF1.3 | A0A0A6YYJ5 | 0.761 | -0.336 |
| MTREX | P42285 | 0.851 | -0.336 |

| | | | |
|---|---|---|---|
| PPIP5K2 | A0A087WZV0 | 0.747 | -0.336 |
| YTHDF2 | Q9Y5A9 | 0.841 | -0.336 |
| CHD1L.1 | Q86WJ1 | 0.852 | -0.337 |
| CHAF1B | Q13112 | 0.806 | -0.338 |
| M6PR | P20645 | 0.866 | -0.338 |
| AGAP2 | F8VVT9 | 0.86 | -0.339 |
| RPN2 | P04844 | 0.831 | -0.339 |
| DDX41 | J3KNN5 | 0.8 | -0.34 |
| INAVA | Q3KP66 | 0.835 | -0.34 |
| PKP3 | Q9Y446 | 0.748 | -0.34 |
| VCP | P55072 | 0.834 | -0.34 |
| ZNF609 | O15014 | 0.82 | -0.34 |
| CDCA2 | Q69YH5 | 0.866 | -0.342 |
| IPO8 | O15397 | 0.776 | -0.342 |
| ITGB1 | P05556 | 0.851 | -0.342 |
| KIAA1522 | Q9P206 | 0.865 | -0.342 |
| TCERG1 | O14776 | 0.874 | -0.342 |
| WDR5 | P61964 | 0.823 | -0.343 |
| ZNF48 | Q96MX3 | 0.866 | -0.343 |
| ADAM9 | Q13443 | 0.844 | -0.344 |
| EIF2B4 | E7ERK9 | 0.838 | -0.344 |
| LEO1 | Q8WVC0 | 0.808 | -0.344 |
| CANX | P27824 | 0.856 | -0.345 |
| CUL3 | Q13618 | 0.809 | -0.345 |
| KIF2A | O00139 | 0.821 | -0.345 |
| SMPDL3B | Q92485 | 0.874 | -0.345 |
| APEX1 | P27695 | 0.843 | -0.346 |
| ETFB | P38117 | 0.867 | -0.346 |
| PPP3CA | Q08209 | 0.864 | -0.346 |
| AP2A2 | O94973 | 0.856 | -0.347 |
| CBX6 | O95503 | 0.748 | -0.347 |
| EXOSC10 | Q01780 | 0.775 | -0.347 |
| MACF1 | H3BQK9 | 0.848 | -0.347 |
| DKK1 | O94907 | 0.846 | -0.348 |
| ITGB4 | P16144 | 0.837 | -0.349 |
| LIMA1 | Q9UHB6 | 0.868 | -0.349 |
| NACAP1 | Q9BZK3 | 0.823 | -0.349 |
| PPFIBP1.1 | A0A0A0MTP2 | 0.853 | -0.349 |
| PTPRG | P23470 | 0.837 | -0.35 |
| HEXIM1 | O94992 | 0.853 | -0.351 |
| PAK4 | O96013 | 0.84 | -0.351 |
| RAB21 | Q9UL25 | 0.844 | -0.351 |
| RPL32 | F8W727 | 0.867 | -0.351 |
| SEMA7A | O75326 | 0.86 | -0.351 |
| ESYT2 | H7BXI1 | 0.806 | -0.352 |
| LSR | Q86X29 | 0.862 | -0.352 |
| SHOC2 | Q9UQ13 | 0.841 | -0.352 |

| | | | |
|---|---|---|---|
| ST13 | P50502 | 0.859 | -0.352 |
| TRIP10 | Q15642 | 0.787 | -0.352 |
| TXNRD1 | Q16881 | 0.834 | -0.352 |
| ZNF100 | A0A075B7G6 | 0.829 | -0.352 |
| IDH3B | A0A087WZN1 | 0.853 | -0.353 |
| LEMD3 | Q9Y2U8 | 0.725 | -0.353 |
| MAP1S | Q66K74 | 0.831 | -0.353 |
| DLG3 | Q5JUW8 | 0.848 | -0.354 |
| PBDC1 | Q9BVG4 | 0.811 | -0.354 |
| DHX15 | O43143 | 0.813 | -0.355 |
| EXOC7 | Q9UPT5 | 0.788 | -0.355 |
| RAD54B | Q9Y620 | 0.847 | -0.355 |
| DDX46.1 | Q7L014 | 0.834 | -0.356 |
| PTDSS2 | Q9BVG9 | 0.808 | -0.356 |
| YTHDF3 | A0A087WY31 | 0.859 | -0.356 |
| ANAPC7 | Q9UJX3 | 0.826 | -0.357 |
| CYB5B | J3KNF8 | 0.856 | -0.357 |
| NSMCE4A | Q9NXX6 | 0.74 | -0.357 |
| PGAP1 | Q75T13 | 0.838 | -0.357 |
| B4DLN1 | B4DLN1 | 0.845 | -0.358 |
| BOP1 | Q14137 | 0.865 | -0.358 |
| FIBP | O43427 | 0.849 | -0.358 |
| CTNND1 | O60716 | 0.729 | -0.359 |
| TPM4.1 | K7ENT6 | 0.794 | -0.359 |
| XPO1 | O14980 | 0.834 | -0.359 |
| SPATA5L1 | Q9BVQ7 | 0.814 | -0.36 |
| TKT | P29401 | 0.854 | -0.36 |
| TTLL4 | Q14679 | 0.833 | -0.36 |
| USP8 | P40818 | 0.844 | -0.36 |
| CDSN | G8JLG2 | 0.846 | -0.361 |
| CLUH | O75153 | 0.79 | -0.361 |
| CYC1 | P08574 | 0.755 | -0.361 |
| TCF7L2.1 | Q9NQB0 | 0.775 | -0.361 |
| UTP14A | Q9BVJ6 | 0.849 | -0.361 |
| BSG | P35613 | 0.852 | -0.362 |
| PDCL3 | Q9H2J4 | 0.869 | -0.362 |
| SNX18 | Q96RF0 | 0.849 | -0.362 |
| XPOT | O43592 | 0.824 | -0.362 |
| HSD17B12 | Q53GQ0 | 0.842 | -0.363 |
| DNAJB11 | Q9UBS4 | 0.72 | -0.364 |
| KIF3B | O15066 | 0.833 | -0.364 |
| PSMD14 | O00487 | 0.843 | -0.364 |
| UBR1 | Q8IWV7 | 0.81 | -0.364 |
| CCDC6 | Q16204 | 0.828 | -0.365 |
| HSP90B1 | P14625 | 0.844 | -0.365 |
| MED19 | J3KR33 | 0.799 | -0.365 |
| MPHOSPH10 | O00566 | 0.867 | -0.365 |

| | | | |
|---|---|---|---|
| STXBP4 | Q6ZWJ1 | 0.839 | -0.365 |
| TCOF1.1 | E7ETY2 | 0.865 | -0.365 |
| VPS53 | Q5VIR6 | 0.812 | -0.365 |
| WEE1 | P30291 | 0.715 | -0.365 |
| AAAS | Q9NRG9 | 0.832 | -0.366 |
| CTR9 | Q6PD62 | 0.759 | -0.366 |
| DUS3L | Q96G46 | 0.834 | -0.366 |
| NEIL2 | Q969S2 | 0.864 | -0.366 |
| RB1 | P06400 | 0.822 | -0.366 |
| NUDCD1 | Q96RS6 | 0.811 | -0.367 |
| BRAT1 | Q6PJG6 | 0.842 | -0.368 |
| DHX57 | Q6P158 | 0.841 | -0.368 |
| NUP50 | Q9UKX7 | 0.818 | -0.368 |
| RAB35 | Q15286 | 0.869 | -0.368 |
| USP24 | Q9UPU5 | 0.846 | -0.368 |
| USP7 | Q93009 | 0.752 | -0.368 |
| EMSY | Q7Z589 | 0.866 | -0.369 |
| GRWD1 | Q9BQ67 | 0.717 | -0.369 |
| USP36 | Q9P275 | 0.859 | -0.37 |
| CHAF1A | Q13111 | 0.849 | -0.371 |
| CHD1 | O14646 | 0.824 | -0.372 |
| NOB1 | Q9ULX3 | 0.796 | -0.372 |
| SLC27A4 | Q6P1M0 | 0.817 | -0.372 |
| AHSA1 | O95433 | 0.856 | -0.373 |
| NAA10 | P41227 | 0.738 | -0.373 |
| SLC38A10 | Q9HBR0 | 0.775 | -0.373 |
| SEC23A | Q15436 | 0.814 | -0.374 |
| WDR60 | Q8WVS4 | 0.858 | -0.374 |
| DENND3 | E9PF32 | 0.823 | -0.375 |
| GLOD4 | F6TLX2 | 0.834 | -0.375 |
| NPC1 | O15118 | 0.854 | -0.375 |
| PACS1 | Q6VY07 | 0.853 | -0.375 |
| SMARCD2 | Q92925 | 0.852 | -0.376 |
| TROVE2 | P10155 | 0.827 | -0.376 |
| FOXK1 | P85037 | 0.767 | -0.377 |
| GRPEL1 | Q9HAV7 | 0.852 | -0.377 |
| LSG1 | Q9H089 | 0.799 | -0.377 |
| ATG16L1 | E7EVC7 | 0.82 | -0.378 |
| CDK11B | P21127 | 0.862 | -0.378 |
| MTIF2 | P46199 | 0.737 | -0.378 |
| NASP | P49321 | 0.777 | -0.378 |
| AKR1A1 | P14550 | 0.865 | -0.379 |
| CACNG6 | Q9BXT2 | 0.828 | -0.379 |
| CEP295 | Q9C0D2 | 0.776 | -0.38 |
| GART | P22102 | 0.829 | -0.38 |
| JAK1 | P23458 | 0.864 | -0.381 |
| KPNA3 | O00505 | 0.717 | -0.381 |

| | | | |
|---|---|---|---|
| PXDN | Q92626 | 0.784 | -0.382 |
| SMG6 | Q86US8 | 0.858 | -0.382 |
| VAT1 | Q99536 | 0.837 | -0.382 |
| WDR20 | A0A088AWN2 | 0.833 | -0.382 |
| BIRC2 | Q13490 | 0.858 | -0.383 |
| ESF1 | Q9H501 | 0.835 | -0.383 |
| SMARCC1 | Q92922 | 0.866 | -0.383 |
| MAPK14 | Q16539 | 0.842 | -0.384 |
| XYLT2 | Q9H1B5 | 0.845 | -0.384 |
| EEF1D | E9PRY8 | 0.778 | -0.386 |
| NAE1 | Q13564 | 0.842 | -0.386 |
| SMTN | A0A087X1R1 | 0.756 | -0.386 |
| ABCF3 | Q9NUQ8 | 0.69 | -0.387 |
| KIF1B | O60333 | 0.796 | -0.387 |
| ZNF608 | Q9ULD9 | 0.83 | -0.387 |
| ESRP1 | Q6NXG1 | 0.82 | -0.39 |
| EXD2 | Q9NVH0 | 0.814 | -0.391 |
| RAD23B | P54727 | 0.834 | -0.391 |
| EED | O75530 | 0.83 | -0.392 |
| PABPN1 | Q86U42 | 0.807 | -0.392 |
| PC | P11498 | 0.84 | -0.392 |
| PMPCA | Q10713 | 0.833 | -0.392 |
| SF1.1 | H7C0N4 | 0.811 | -0.392 |
| CD320 | Q9NPF0 | 0.73 | -0.393 |
| KLF2 | Q9Y5W3 | 0.74 | -0.393 |
| TNIK | Q9UKE5 | 0.861 | -0.393 |
| PWP1 | Q13610 | 0.82 | -0.396 |
| MTHFSD | Q2M296 | 0.789 | -0.397 |
| NOMO2 | Q5JPE7 | 0.777 | -0.398 |
| VPS26A | O75436 | 0.815 | -0.398 |
| SF3B2 | Q13435 | 0.85 | -0.399 |
| TUBB1 | Q9H4B7 | 0.725 | -0.399 |
| VANGL1 | Q8TAA9 | 0.819 | -0.399 |
| EHMT2 | A0A0G2JIR1 | 0.838 | -0.4 |
| THOC6 | Q86W42 | 0.851 | -0.4 |
| TCOF1 | Q13428 | 0.86 | -0.401 |
| TLK2 | Q86UE8 | 0.835 | -0.401 |
| TMEM201 | Q5SNT2 | 0.753 | -0.401 |
| CDH1 | A0A087WXI5 | 0.866 | -0.402 |
| EEF1A1 | P68104 | 0.778 | -0.402 |
| PLAA | Q9Y263 | 0.696 | -0.402 |
| PPP2R1B | P30154 | 0.844 | -0.402 |
| PARD3 | Q5VWV2 | 0.842 | -0.403 |
| CCAR1 | Q8IX12 | 0.84 | -0.404 |
| LASP1 | Q14847 | 0.812 | -0.404 |
| PPP2R5D | Q14738 | 0.708 | -0.404 |
| ABCC1 | P33527 | 0.821 | -0.405 |

| | | | |
|---|---|---|---|
| ABL1 | P00519 | 0.833 | -0.405 |
| IMMT | Q16891 | 0.783 | -0.405 |
| PLCB3 | Q01970 | 0.859 | -0.405 |
| TM9SF4 | Q92544 | 0.857 | -0.405 |
| MSL1 | Q68DK7 | 0.857 | -0.406 |
| PPP1R12C.1 | K7EL81 | 0.827 | -0.406 |
| DNAJB1 | P25685 | 0.831 | -0.407 |
| FHOD1 | Q9Y613 | 0.822 | -0.407 |
| SMC4 | Q9NTJ3 | 0.711 | -0.407 |
| DHCR7 | Q9UBM7 | 0.855 | -0.408 |
| DLG1 | A0A0C4DFT3 | 0.81 | -0.408 |
| MCM3 | P25205 | 0.709 | -0.409 |
| OTUD7B | Q6GQQ9 | 0.717 | -0.409 |
| ZXDC | Q2QGD7 | 0.863 | -0.409 |
| NOSIP | Q9Y314 | 0.792 | -0.41 |
| MMTAG2 | Q9BU76 | 0.734 | -0.411 |
| SCARB1 | B7ZKQ9 | 0.761 | -0.411 |
| MRPL1 | Q9BYD6 | 0.774 | -0.412 |
| NET1 | Q7Z628 | 0.85 | -0.412 |
| POC1A | Q8NBT0 | 0.842 | -0.412 |
| ARL6IP4 | Q66PJ3 | 0.802 | -0.413 |
| METTL15 | A6NJ78 | 0.721 | -0.413 |
| RBM25 | P49756 | 0.858 | -0.413 |
| ZNF398 | Q8TD17 | 0.846 | -0.413 |
| DBNL | Q9UJU6 | 0.852 | -0.414 |
| SLC7A5 | Q01650 | 0.866 | -0.414 |
| ABT1 | Q9ULW3 | 0.831 | -0.415 |
| AHCYL1 | O43865 | 0.743 | -0.415 |
| ANXA3 | P12429 | 0.845 | -0.415 |
| AGAP3 | Q96P47 | 0.843 | -0.416 |
| HDAC1 | Q13547 | 0.734 | -0.416 |
| HSD17B4 | P51659 | 0.791 | -0.416 |
| TNPO3 | Q9Y5L0 | 0.84 | -0.416 |
| ARNT | P27540 | 0.672 | -0.417 |
| MAP2K3 | P46734 | 0.735 | -0.417 |
| UGT8 | Q16880 | 0.854 | -0.417 |
| EPCAM | B5MCA4 | 0.858 | -0.418 |
| ATP2B4 | P23634 | 0.832 | -0.42 |
| KLHDC4 | Q8TBB5 | 0.848 | -0.42 |
| PPP2R5C | H0YJ75 | 0.827 | -0.421 |
| GTPBP1 | O00178 | 0.831 | -0.422 |
| KIF1BP | A0A1B0GUA3 | 0.67 | -0.422 |
| PYGB | P11216 | 0.841 | -0.422 |
| SCAF4 | O95104 | 0.856 | -0.422 |
| WDR77 | Q9BQA1 | 0.744 | -0.422 |
| CREBBP | Q92793 | 0.844 | -0.423 |
| NCAPH2 | Q6IBW4 | 0.786 | -0.423 |

| | | | |
|---|---|---|---|
| CSDE1 | O75534 | 0.819 | -0.424 |
| LTV1 | Q96GA3 | 0.837 | -0.425 |
| NT5DC1 | Q5TFE4 | 0.856 | -0.425 |
| OSBP | P22059 | 0.8 | -0.425 |
| SART1 | O43290 | 0.857 | -0.425 |
| POLR2E | P19388 | 0.724 | -0.426 |
| PSMA4 | P25789 | 0.825 | -0.426 |
| CCNT1 | O60563 | 0.841 | -0.427 |
| CYP51A1 | A0A0C4DFL7 | 0.844 | -0.427 |
| EPB41L2.1 | H0Y5B0 | 0.692 | -0.428 |
| IFRD2 | Q12894 | 0.821 | -0.428 |
| YY1 | P25490 | 0.846 | -0.428 |
| EARS2 | Q5JPH6 | 0.795 | -0.43 |
| PRCP | P42785 | 0.768 | -0.43 |
| SLC25A19 | Q9HC21 | 0.827 | -0.43 |
| XPO7 | E7ESC6 | 0.844 | -0.431 |
| MED24.1 | F5GY88 | 0.796 | -0.432 |
| POLA1 | A6NMQ1 | 0.746 | -0.432 |
| UBXN1 | Q04323 | 0.838 | -0.432 |
| ENAH | Q8N8S7 | 0.804 | -0.433 |
| NECAP1 | Q8NC96 | 0.801 | -0.434 |
| PCCA | P05165 | 0.855 | -0.435 |
| PM20D2 | Q8IYS1 | 0.847 | -0.435 |
| PWWP2B | Q6NUJ5 | 0.838 | -0.435 |
| COPB2 | P35606 | 0.788 | -0.436 |
| DDB1 | Q16531 | 0.65 | -0.436 |
| NAA15 | Q9BXJ9 | 0.701 | -0.436 |
| POGZ | Q7Z3K3 | 0.85 | -0.436 |
| SFSWAP | Q12872 | 0.865 | -0.437 |
| KSR1 | Q8IVT5 | 0.825 | -0.438 |
| MTMR1 | F8WA39 | 0.839 | -0.438 |
| AGAP3.1 | E7ESL9 | 0.777 | -0.439 |
| ARHGAP39 | Q9C0H5 | 0.839 | -0.439 |
| DIP2A | Q14689 | 0.778 | -0.439 |
| GSR | P00390 | 0.847 | -0.439 |
| DNAJA1 | P31689 | 0.701 | -0.441 |
| EEF1G | P26641 | 0.697 | -0.441 |
| WASH2P | Q6VEQ5 | 0.726 | -0.441 |
| ZC3H7B | Q9UGR2 | 0.799 | -0.441 |
| CAPN2 | P17655 | 0.832 | -0.443 |
| KIDINS220 | Q9ULH0 | 0.791 | -0.443 |
| NPEPPS | P55786 | 0.819 | -0.443 |
| ACLY | P53396 | 0.824 | -0.445 |
| GALNT14 | Q96FL9 | 0.837 | -0.445 |
| TWISTNB | Q3B726 | 0.811 | -0.445 |
| DDX11L8 | A8MPP1 | 0.699 | -0.446 |
| EEF2 | P13639 | 0.819 | -0.447 |

| | | | |
|---|---|---|---|
| LPCAT1 | Q8NF37 | 0.732 | -0.447 |
| PRPF31 | Q8WWY3 | 0.836 | -0.448 |
| SYBU | A0A0C4DG86 | 0.817 | -0.448 |
| EIF4A3 | P38919 | 0.814 | -0.449 |
| FEN1 | P39748 | 0.813 | -0.45 |
| JCAD | Q9P266 | 0.772 | -0.45 |
| LUC7L | Q9NQ29 | 0.842 | -0.45 |
| CC2D2A | Q9P2K1 | 0.84 | -0.451 |
| MYO19 | Q96H55 | 0.628 | -0.451 |
| DNAJC3 | Q13217 | 0.718 | -0.452 |
| PPP5C | P53041 | 0.792 | -0.452 |
| RHPN2 | Q8IUC4 | 0.771 | -0.452 |
| SERPINB5 | P36952 | 0.867 | -0.452 |
| GANAB | Q14697 | 0.78 | -0.453 |
| MYO1C | O00159 | 0.821 | -0.453 |
| PSMD1 | Q99460 | 0.823 | -0.453 |
| ACSL3 | O95573 | 0.819 | -0.454 |
| KIF4A | O95239 | 0.838 | -0.454 |
| MAP2 | P11137 | 0.775 | -0.456 |
| INTS11 | Q5TA45 | 0.678 | -0.457 |
| PSMC3 | P17980 | 0.779 | -0.458 |
| PSMD7 | P51665 | 0.845 | -0.458 |
| SDC1 | P18827 | 0.729 | -0.458 |
| METTL13 | Q8N6R0 | 0.821 | -0.459 |
| PTPRF | P10586 | 0.753 | -0.459 |
| SRBD1 | Q8N5C6 | 0.805 | -0.459 |
| TULP3 | O75386 | 0.729 | -0.459 |
| CD276 | Q5ZPR3 | 0.791 | -0.46 |
| PDE12 | Q6L8Q7 | 0.809 | -0.46 |
| TAF5 | Q15542 | 0.852 | -0.46 |
| TBCD | J3KR97 | 0.796 | -0.46 |
| FANCI | Q9NVI1 | 0.642 | -0.461 |
| PCBP1 | Q15365 | 0.774 | -0.461 |
| STT3B | Q8TCJ2 | 0.805 | -0.461 |
| ARID4B | Q4LE39 | 0.859 | -0.462 |
| PRPF38B | Q5VTL8 | 0.774 | -0.462 |
| FMNL2 | Q96PY5 | 0.681 | -0.463 |
| NSD3 | Q9BZ95 | 0.856 | -0.463 |
| PCNA | P12004 | 0.838 | -0.463 |
| TEC | P42680 | 0.838 | -0.463 |
| DNAH10 | A0A1C7CYW8 | 0.854 | -0.464 |
| FARP2 | O94887 | 0.838 | -0.464 |
| GNL1 | P36915 | 0.672 | -0.464 |
| LLGL1 | Q15334 | 0.846 | -0.464 |
| PPIL4 | Q8WUA2 | 0.807 | -0.464 |
| SEC62 | Q99442 | 0.798 | -0.464 |
| ZNF526 | Q8TF50 | 0.85 | -0.464 |

| | | | |
|---|---|---|---|
| SRCAP | Q6ZRS2 | 0.792 | -0.465 |
| KIFC1 | Q9BW19 | 0.766 | -0.466 |
| MTHFD1 | F5H2F4 | 0.807 | -0.466 |
| NAA15.1 | A0A0B4J1W3 | 0.722 | -0.466 |
| NEDD4L | A0A1B0GVY1 | 0.838 | -0.466 |
| TALDO1 | P37837 | 0.848 | -0.466 |
| AHCY | P23526 | 0.85 | -0.468 |
| CWC22 | Q9HCG8 | 0.603 | -0.468 |
| NEK4 | P51957 | 0.819 | -0.468 |
| NR2F1 | P10589 | 0.646 | -0.468 |
| STOML2 | Q9UJZ1 | 0.849 | -0.468 |
| DNMT1 | P26358 | 0.824 | -0.469 |
| EEFSEC | P57772 | 0.823 | -0.469 |
| CTSD | P07339 | 0.861 | -0.47 |
| ANKRD27 | Q96NW4 | 0.724 | -0.471 |
| CTBP1 | Q13363 | 0.726 | -0.471 |
| EEF1A2 | Q05639 | 0.574 | -0.471 |
| NCAPH | Q15003 | 0.802 | -0.471 |
| ABCF1.1 | Q5STZ8 | 0.795 | -0.472 |
| METAP2 | P50579 | 0.778 | -0.472 |
| PARD3.1 | Q5VWU8 | 0.843 | -0.472 |
| TMEM237 | Q96Q45 | 0.679 | -0.472 |
| PHGDH | O43175 | 0.783 | -0.474 |
| PICALM | Q13492 | 0.831 | -0.475 |
| TBL1XR1 | Q9BZK7 | 0.626 | -0.475 |
| TUBGCP4 | Q9UGJ1 | 0.675 | -0.475 |
| GIPC1 | O14908 | 0.816 | -0.476 |
| MCM5 | P33992 | 0.618 | -0.476 |
| CHD7 | Q9P2D1 | 0.781 | -0.477 |
| TOPORS | Q9NS56 | 0.852 | -0.478 |
| FBXO2 | Q9UK22 | 0.825 | -0.481 |
| SREBF2 | Q12772 | 0.774 | -0.481 |
| COL4A3BP | Q9Y5P4 | 0.819 | -0.482 |
| EIF4A2 | Q14240 | 0.84 | -0.482 |
| PRMT5 | O14744 | 0.715 | -0.482 |
| CRTAP | O75718 | 0.656 | -0.483 |
| ESYT1 | Q9BSJ8 | 0.792 | -0.484 |
| WDR75 | Q8IWA0 | 0.819 | -0.484 |
| YTHDF1 | Q9BYJ9 | 0.847 | -0.484 |
| ELAC2 | Q9BQ52 | 0.782 | -0.486 |
| POLR3E | Q9NVU0 | 0.781 | -0.486 |
| ATG5 | Q9H1Y0 | 0.775 | -0.487 |
| PSMD5 | Q16401 | 0.843 | -0.487 |
| ARMT1 | Q9H993 | 0.692 | -0.488 |
| RPS6KA4 | O75676 | 0.802 | -0.489 |
| MET | P08581 | 0.719 | -0.491 |
| MMS19 | Q96T76 | 0.692 | -0.491 |

| | | | |
|---|---|---|---|
| VARS | P26640 | 0.594 | -0.491 |
| RETREG2 | Q8NC44 | 0.732 | -0.492 |
| CTCF | P49711 | 0.84 | -0.493 |
| PTOV1 | Q86YD1 | 0.618 | -0.493 |
| CCDC144A | A2RUR9 | 0.69 | -0.494 |
| AGTPBP1 | J3KNS1 | 0.847 | -0.495 |
| ESPL1 | Q14674 | 0.842 | -0.495 |
| PPRC1 | Q5VV67 | 0.792 | -0.495 |
| RRP15 | Q9Y3B9 | 0.863 | -0.496 |
| DNAJC1 | Q96KC8 | 0.831 | -0.497 |
| DIDO1 | Q9BTC0 | 0.822 | -0.498 |
| CSE1L | P55060 | 0.802 | -0.499 |
| PTGES3L-AARSD1 | C9J5N1 | 0.837 | -0.499 |
| BAG6 | A0A0G2JK23 | 0.695 | -0.501 |
| FAM91A1 | Q658Y4 | 0.658 | -0.501 |
| LAMP1 | P11279 | 0.844 | -0.501 |
| NSUN2 | Q08J23 | 0.753 | -0.501 |
| FNTA | P49354 | 0.815 | -0.502 |
| RNF20 | Q5VTR2 | 0.809 | -0.502 |
| GATAD2A | Q86YP4 | 0.785 | -0.503 |
| RBM33.1 | Q96EV2 | 0.731 | -0.503 |
| VPS4B | O75351 | 0.778 | -0.504 |
| GPATCH1 | Q9BRR8 | 0.802 | -0.505 |
| CLUH.1 | I3L2B0 | 0.743 | -0.506 |
| NUP210 | Q8TEM1 | 0.706 | -0.508 |
| ZFP91 | Q96JP5 | 0.834 | -0.508 |
| ACACA | Q13085 | 0.706 | -0.509 |
| CKAP4 | Q07065 | 0.776 | -0.509 |
| DPF1 | E9PDV3 | 0.753 | -0.509 |
| HTATSF1 | O43719 | 0.671 | -0.509 |
| RCC1 | P18754 | 0.692 | -0.509 |
| SGF29 | Q96ES7 | 0.818 | -0.509 |
| ANXA6 | P08133 | 0.714 | -0.51 |
| CDK5 | Q00535 | 0.715 | -0.51 |
| MOGS | Q13724 | 0.835 | -0.51 |
| RIC8A | Q9NPQ8 | 0.772 | -0.51 |
| EGR1 | P18146 | 0.785 | -0.511 |
| FRMD5 | Q7Z6J6 | 0.816 | -0.511 |
| GUF1 | Q8N442 | 0.8 | -0.511 |
| ATP5O | P48047 | 0.556 | -0.512 |
| DSE | Q9UL01 | 0.826 | -0.512 |
| SP2 | Q02086 | 0.854 | -0.512 |
| DHX16 | O60231 | 0.822 | -0.513 |
| GLS | O94925 | 0.851 | -0.513 |
| MTBP | Q96DY7 | 0.833 | -0.514 |
| PUS1 | Q9Y606 | 0.656 | -0.514 |
| RSRC1 | Q96IZ7 | 0.791 | -0.514 |

| | | | |
|---|---|---|---|
| GPC1 | P35052 | 0.815 | -0.515 |
| SAE1 | Q9UBE0 | 0.835 | -0.515 |
| CLIP2 | Q9UDT6 | 0.825 | -0.516 |
| CAND1 | Q86VP6 | 0.747 | -0.518 |
| HNRNPF | P52597 | 0.722 | -0.518 |
| TRIP13 | Q15645 | 0.818 | -0.518 |
| RPN1 | P04843 | 0.776 | -0.519 |
| LRRC47 | Q8N1G4 | 0.661 | -0.52 |
| PDZD8 | Q8NEN9 | 0.822 | -0.52 |
| CD9 | G8JLH6 | 0.846 | -0.521 |
| FARP2.1 | H7C3M7 | 0.826 | -0.521 |
| FOXK2 | Q01167 | 0.662 | -0.521 |
| POR | P16435 | 0.813 | -0.521 |
| RECQL5 | O94762 | 0.829 | -0.521 |
| RPS6KA3 | P51812 | 0.713 | -0.521 |
| TCAF1 | Q9Y4C2 | 0.694 | -0.521 |
| IMPDH1 | P20839 | 0.857 | -0.522 |
| WWC1 | Q8IX03 | 0.654 | -0.522 |
| MAP4K3 | Q8IVH8 | 0.787 | -0.523 |
| EPB41 | P11171 | 0.801 | -0.524 |
| ME2 | P23368 | 0.836 | -0.526 |
| NRF1 | Q16656 | 0.712 | -0.526 |
| PSMD8 | P48556 | 0.682 | -0.526 |
| TTLL12 | Q14166 | 0.833 | -0.526 |
| USP33 | Q8TEY7 | 0.825 | -0.526 |
| ZNF628 | Q5EBL2 | 0.846 | -0.526 |
| AARS2 | Q5JTZ9 | 0.811 | -0.527 |
| KPNA1 | P52294 | 0.738 | -0.527 |
| ABCB7 | O75027 | 0.747 | -0.528 |
| EIF1AY | O14602 | 0.756 | -0.529 |
| ZNF689 | Q96CS4 | 0.764 | -0.529 |
| PLCH1 | Q4KWH8 | 0.743 | -0.53 |
| SEC61A1 | B4DR61 | 0.797 | -0.53 |
| CAMTA2 | I3L3W6 | 0.822 | -0.531 |
| ELP1 | O95163 | 0.523 | -0.531 |
| ZHX1 | Q9UKY1 | 0.772 | -0.531 |
| CLSPN | Q9HAW4 | 0.838 | -0.532 |
| COASY | Q13057 | 0.836 | -0.532 |
| EXOC2 | Q96KP1 | 0.664 | -0.532 |
| PSMD11 | O00231 | 0.839 | -0.532 |
| CWF19L1 | Q69YN2 | 0.739 | -0.533 |
| GOPC | Q9HD26 | 0.559 | -0.533 |
| MCCC1 | Q96RQ3 | 0.823 | -0.533 |
| MEPCE | Q7L2J0 | 0.779 | -0.533 |
| RGS3 | P49796 | 0.838 | -0.533 |
| TTC1 | Q99614 | 0.818 | -0.533 |
| EXT2 | Q93063 | 0.791 | -0.535 |

| | | | |
|---|---|---|---|
| CHTF18 | E7EXA6 | 0.727 | -0.536 |
| DCBLD2 | Q96PD2 | 0.809 | -0.536 |
| DERL1 | Q9BUN8 | 0.64 | -0.536 |
| PRKAR1A.1 | K7EPR5 | 0.803 | -0.536 |
| AVIL | O75366 | 0.827 | -0.537 |
| BICRAL | Q6AI39 | 0.836 | -0.537 |
| CACNB1 | Q02641 | 0.693 | -0.538 |
| GDI2 | P50395 | 0.831 | -0.538 |
| PLEKHH2 | Q8IVE3 | 0.767 | -0.539 |
| IPO11 | Q9UI26 | 0.599 | -0.54 |
| QSOX1 | O00391 | 0.785 | -0.54 |
| TRIM28 | Q13263 | 0.474 | -0.54 |
| MTA2 | O94776 | 0.742 | -0.541 |
| PROCR | Q9UNN8 | 0.824 | -0.541 |
| WDR26 | Q9H7D7 | 0.753 | -0.541 |
| ARHGAP27 | A0A0G2JNY3 | 0.662 | -0.542 |
| HEATR1 | Q9H583 | 0.788 | -0.542 |
| TMED10 | P49755 | 0.708 | -0.542 |
| DNTTIP2 | Q5QJE6 | 0.597 | -0.543 |
| ARMC1 | Q9NVT9 | 0.673 | -0.544 |
| NPLOC4 | Q8TAT6 | 0.731 | -0.544 |
| SMG5 | Q9UPR3 | 0.783 | -0.544 |
| UBA6 | A0AVT1 | 0.737 | -0.544 |
| WDHD1 | O75717 | 0.789 | -0.544 |
| JADE3 | Q92613 | 0.798 | -0.545 |
| RABGGTA | Q92696 | 0.833 | -0.545 |
| LENG8 | Q96PV6 | 0.772 | -0.547 |
| RAB1A | P62820 | 0.666 | -0.547 |
| SNX29 | Q8TEQ0 | 0.822 | -0.547 |
| CELF1 | G5EA30 | 0.769 | -0.549 |
| RBM5 | P52756 | 0.753 | -0.55 |
| EWSR1 | Q01844 | 0.494 | -0.551 |
| GTPBP6 | O43824 | 0.783 | -0.551 |
| SLC16A1 | P53985 | 0.633 | -0.551 |
| SQOR | Q9Y6N5 | 0.615 | -0.551 |
| RPS6KA5 | O75582 | 0.689 | -0.552 |
| AFG3L2 | Q9Y4W6 | 0.753 | -0.553 |
| RBM6 | P78332 | 0.579 | -0.553 |
| RFC5 | P40937 | 0.608 | -0.553 |
| RER1 | O15258 | 0.765 | -0.554 |
| PRR12 | Q9ULL5 | 0.804 | -0.555 |
| WRAP53 | Q9BUR4 | 0.696 | -0.555 |
| DONSON | Q9NYP3 | 0.835 | -0.556 |
| IPO7 | O95373 | 0.745 | -0.556 |
| MAPK1 | P28482 | 0.809 | -0.556 |
| ZBTB10 | Q96DT7 | 0.834 | -0.557 |
| JMJD6 | Q6NYC1 | 0.548 | -0.558 |

| | | | |
|---|---|---|---|
| PRCC | Q92733 | 0.699 | -0.558 |
| TOR1AIP1 | Q5JTV8 | 0.629 | -0.558 |
| DBN1 | Q16643 | 0.814 | -0.559 |
| DNAJC21 | Q5F1R6 | 0.812 | -0.559 |
| CACTIN | Q8WUQ7 | 0.796 | -0.56 |
| EHD2 | Q9NZN4 | 0.531 | -0.56 |
| TFRC | P02786 | 0.814 | -0.563 |
| BLMH | Q13867 | 0.693 | -0.564 |
| SMARCB1 | Q12824 | 0.793 | -0.564 |
| ALDH1B1 | P30837 | 0.718 | -0.565 |
| GTF2H2 | Q13888 | 0.752 | -0.566 |
| NCAPG2 | Q86XI2 | 0.819 | -0.568 |
| RHOT2 | Q8IXI1 | 0.699 | -0.568 |
| SETD2 | Q9BYW2 | 0.835 | -0.568 |
| ZBED5 | Q49AG3 | 0.828 | -0.568 |
| CMAS | Q8NFW8 | 0.763 | -0.569 |
| RRM1 | P23921 | 0.797 | -0.57 |
| SLC39A7 | Q92504 | 0.741 | -0.57 |
| FAM76A | Q8TAV0 | 0.652 | -0.572 |
| GALK1 | P51570 | 0.783 | -0.573 |
| KIF2C | Q99661 | 0.795 | -0.573 |
| ADNP | Q9H2P0 | 0.834 | -0.574 |
| FUCA2 | Q9BTY2 | 0.778 | -0.574 |
| NAP1L1 | P55209 | 0.694 | -0.577 |
| ATP13A1 | Q9HD20 | 0.778 | -0.578 |
| PTK7 | Q13308 | 0.799 | -0.578 |
| ATXN10 | Q9UBB4 | 0.808 | -0.58 |
| MCCC2 | Q9HCC0 | 0.818 | -0.58 |
| SAP130.1 | H7BXF5 | 0.84 | -0.58 |
| HEATR6 | Q6AI08 | 0.703 | -0.581 |
| WWP2 | O00308 | 0.677 | -0.581 |
| EXOSC4 | E9PI41 | 0.729 | -0.582 |
| API5 | Q9BZZ5 | 0.626 | -0.583 |
| CC2D1B | H7C1U3 | 0.689 | -0.583 |
| EYA3 | Q99504 | 0.815 | -0.584 |
| GBE1 | Q04446 | 0.822 | -0.584 |
| TCF25 | Q9BQ70 | 0.809 | -0.586 |
| PARP9 | Q8IXQ6 | 0.718 | -0.587 |
| GALM | Q96C23 | 0.783 | -0.588 |
| MAPRE2 | Q15555 | 0.458 | -0.588 |
| SDHB | P21912 | 0.67 | -0.588 |
| BCL7B | F2Z3H6 | 0.832 | -0.589 |
| PBXIP1 | Q96AQ6 | 0.824 | -0.589 |
| SUDS3 | Q9H7L9 | 0.828 | -0.589 |
| MCM7 | P33993 | 0.693 | -0.59 |
| NARS | O43776 | 0.726 | -0.591 |
| GOT1 | P17174 | 0.842 | -0.593 |

| | | | |
|---|---|---|---|
| PARN | O95453 | 0.547 | -0.594 |
| FAM167B | Q9BTA0 | 0.718 | -0.595 |
| LBR | Q14739 | 0.595 | -0.595 |
| SUN2 | Q9UH99 | 0.761 | -0.595 |
| DGCR8 | Q8WYQ5 | 0.547 | -0.597 |
| HK1 | P19367 | 0.767 | -0.597 |
| EFNB1 | P98172 | 0.754 | -0.598 |
| MCM6 | Q14566 | 0.713 | -0.598 |
| ELF3 | P78545 | 0.842 | -0.599 |
| HARS2 | P49590 | 0.764 | -0.6 |
| PDS5B | Q9NTI5 | 0.824 | -0.6 |
| AVL9 | Q8NBF6 | 0.777 | -0.602 |
| RBM12 | Q9NTZ6 | 0.748 | -0.602 |
| MRPL37 | Q9BZE1 | 0.757 | -0.604 |
| FOXRED1 | Q96CU9 | 0.607 | -0.606 |
| MAGI3 | Q5TCQ9 | 0.827 | -0.606 |
| SLC4A2 | P04920 | 0.611 | -0.606 |
| SYT1 | P21579 | 0.791 | -0.606 |
| VEZF1 | Q14119 | 0.837 | -0.606 |
| EMC1 | Q8N766 | 0.611 | -0.607 |
| XPNPEP3 | Q9NQH7 | 0.755 | -0.608 |
| KMT2A | Q03164 | 0.822 | -0.609 |
| PALB2 | Q86YC2 | 0.797 | -0.609 |
| ZBTB40 | Q9NUA8 | 0.786 | -0.609 |
| SLC25A22 | Q9H936 | 0.784 | -0.61 |
| TEAD1 | H0YE88 | 0.747 | -0.611 |
| FMNL1 | K7EK60 | 0.644 | -0.612 |
| NANS | Q9NR45 | 0.809 | -0.612 |
| PDS5A | Q29RF7 | 0.511 | -0.613 |
| RCOR1 | Q9UKL0 | 0.799 | -0.613 |
| SLC27A2 | O14975 | 0.629 | -0.613 |
| FUS | P35637 | 0.589 | -0.614 |
| ATP5F1C | P36542 | 0.499 | -0.615 |
| XRN2 | Q9H0D6 | 0.482 | -0.615 |
| PGM3 | O95394 | 0.716 | -0.616 |
| RERE | Q9P2R6 | 0.569 | -0.617 |
| WBP11 | Q9Y2W2 | 0.857 | -0.617 |
| MTA1 | Q13330 | 0.728 | -0.619 |
| ZBTB9 | Q96C00 | 0.783 | -0.619 |
| IMMT.1 | C9J406 | 0.75 | -0.62 |
| ACBD3 | Q9H3P7 | 0.83 | -0.621 |
| PFKP | Q01813 | 0.74 | -0.621 |
| PHC1 | P78364 | 0.791 | -0.621 |
| RBBP4 | Q09028 | 0.762 | -0.621 |
| ILVBL | A1L0T0 | 0.662 | -0.622 |
| ALKBH5 | Q6P6C2 | 0.761 | -0.624 |
| KIF22 | Q14807 | 0.758 | -0.625 |

| | | | |
|---|---|---|---|
| RLF | Q13129 | 0.676 | -0.625 |
| SENP2 | Q9HC62 | 0.776 | -0.625 |
| CWF19L2 | Q2TBE0 | 0.75 | -0.626 |
| IPO9 | Q96P70 | 0.539 | -0.626 |
| SPATA13 | Q96N96 | 0.708 | -0.626 |
| AEBP2 | Q6ZN18 | 0.359 | -0.627 |
| MSH6 | P52701 | 0.722 | -0.627 |
| CENPU | Q71F23 | 0.728 | -0.628 |
| MCTS1 | Q9ULC4 | 0.73 | -0.629 |
| PRKCZ | Q05513 | 0.833 | -0.629 |
| CAAP1 | Q9H8G2 | 0.669 | -0.63 |
| GTF2H4 | Q92759 | 0.819 | -0.63 |
| PDHX | O00330 | 0.594 | -0.63 |
| DHX38 | Q92620 | 0.687 | -0.631 |
| ANXA6.1 | E5RK69 | 0.803 | -0.632 |
| INO80D | Q53TQ3 | 0.831 | -0.632 |
| PSMC4 | P43686 | 0.786 | -0.632 |
| METAP2.1 | F8VQZ7 | 0.768 | -0.633 |
| MUM1 | J3KNX4 | 0.764 | -0.633 |
| NMD3 | C9JA08 | 0.423 | -0.634 |
| IKBIP | Q70UQ0 | 0.64 | -0.635 |
| IREB2 | A0A0A6YY96 | 0.623 | -0.635 |
| PFAS | O15067 | 0.773 | -0.635 |
| CCDC59 | Q9P031 | 0.412 | -0.636 |
| TAGLN2 | P37802 | 0.723 | -0.637 |
| IPO4 | Q8TEX9 | 0.704 | -0.638 |
| TMEM205 | Q6UW68 | 0.745 | -0.638 |
| ABI2 | F8WAL6 | 0.583 | -0.639 |
| MAPT | I3L170 | 0.746 | -0.639 |
| CAPG | P40121 | 0.822 | -0.64 |
| CDC20 | Q12834 | 0.463 | -0.64 |
| SMAD1 | Q15797 | 0.773 | -0.64 |
| DDX46 | A0A0C4DG89 | 0.808 | -0.641 |
| ISG20L2 | Q9H9L3 | 0.517 | -0.641 |
| MORC2 | Q9Y6X9 | 0.812 | -0.643 |
| PDCD6IP | Q8WUM4 | 0.639 | -0.643 |
| XPC | Q01831 | 0.775 | -0.643 |
| SYDE1 | Q6ZW31 | 0.762 | -0.644 |
| GLRX3 | O76003 | 0.781 | -0.645 |
| LIG1 | P18858 | 0.749 | -0.645 |
| WDR37 | Q9Y2I8 | 0.82 | -0.645 |
| HAT1 | O14929 | 0.748 | -0.647 |
| CAPN1 | P07384 | 0.761 | -0.649 |
| NCLN | Q969V3 | 0.662 | -0.649 |
| DCLRE1A | Q6PJP8 | 0.78 | -0.652 |
| PINX1 | Q96BK5 | 0.517 | -0.654 |
| SCD | O00767 | 0.715 | -0.654 |

| | | | |
|---|---|---|---|
| FKBP5 | Q13451 | 0.551 | -0.655 |
| SFMBT1 | Q9UHJ3 | 0.839 | -0.655 |
| ARHGAP4 | P98171 | 0.817 | -0.656 |
| PSPC1 | Q8WXF1 | 0.574 | -0.656 |
| KDM1A | O60341 | 0.685 | -0.657 |
| SNRNP40 | Q96DI7 | 0.73 | -0.657 |
| ZNF687 | Q8N1G0 | 0.84 | -0.657 |
| CLIC1 | O00299 | 0.841 | -0.658 |
| CTNNBL1 | A0A087WUB9 | 0.556 | -0.658 |
| NELFE | P18615 | 0.606 | -0.658 |
| PLEKHA7 | A0A1B0GTN9 | 0.719 | -0.658 |
| DNAJC17 | Q9NVM6 | 0.742 | -0.659 |
| RPF1 | Q9H9Y2 | 0.628 | -0.659 |
| ANKZF1 | Q9H8Y5 | 0.68 | -0.66 |
| SAPCD2 | Q86UD0 | 0.566 | -0.66 |
| MARCKSL1 | P49006 | 0.64 | -0.661 |
| PRDX6 | P30041 | 0.772 | -0.661 |
| UBE3C | Q15386 | 0.402 | -0.661 |
| NFIX.1 | C9JWJ8 | 0.702 | -0.662 |
| ERO1A | Q96HE7 | 0.824 | -0.663 |
| RYBP | Q8N488 | 0.62 | -0.663 |
| DNAAF5 | Q86Y56 | 0.8 | -0.664 |
| DROSHA | Q9NRR4 | 0.75 | -0.664 |
| RREB1 | Q92766 | 0.818 | -0.664 |
| CAPNS1 | A0A0C4DGQ5 | 0.741 | -0.667 |
| ZBTB14 | O43829 | 0.681 | -0.667 |
| CUL2 | A0A0A0MTN0 | 0.697 | -0.668 |
| TTLL5 | G3V2J9 | 0.686 | -0.669 |
| PAIP1 | Q9H074 | 0.723 | -0.67 |
| INPP5E | Q9NRR6 | 0.616 | -0.671 |
| CDK9 | P50750 | 0.747 | -0.673 |
| POMGNT1 | Q8WZA1 | 0.75 | -0.673 |
| SLC25A11 | Q02978 | 0.744 | -0.674 |
| PYCR1 | P32322 | 0.829 | -0.675 |
| PGD | P52209 | 0.805 | -0.676 |
| TERF2IP | Q9NYB0 | 0.525 | -0.678 |
| WNT16 | Q9UBV4 | 0.772 | -0.678 |
| DIP2B | Q9P265 | 0.675 | -0.679 |
| CYB5R3 | P00387 | 0.825 | -0.68 |
| RTF1 | Q92541 | 0.665 | -0.68 |
| ZGPAT | Q8N5A5 | 0.701 | -0.68 |
| HIVEP1 | P15822 | 0.792 | -0.681 |
| ISYNA1 | Q9NPH2 | 0.816 | -0.683 |
| MTM1 | Q13496 | 0.671 | -0.685 |
| ERCC4 | Q92889 | 0.811 | -0.687 |
| RHOT1 | Q8IXI2 | 0.606 | -0.687 |
| TANC2.1 | J3QQY7 | 0.789 | -0.687 |

| | | | |
|---|---|---|---|
| ARFGAP3 | Q9NP61 | 0.738 | -0.688 |
| RLIM | Q9NVW2 | 0.699 | -0.688 |
| KDM1B | Q8NB78 | 0.768 | -0.689 |
| NAA25 | Q14CX7 | 0.738 | -0.689 |
| TIA1 | C9JTN7 | 0.471 | -0.689 |
| QRICH1 | Q2TAL8 | 0.818 | -0.69 |
| EFTUD2 | Q15029 | 0.572 | -0.692 |
| NCOA5 | Q9HCD5 | 0.676 | -0.692 |
| KPNA4 | O00629 | 0.231 | -0.693 |
| SAP30 | O75446 | 0.748 | -0.693 |
| HDGF | P51858 | 0.302 | -0.694 |
| MICU2 | Q8IYU8 | 0.729 | -0.694 |
| QSOX2 | Q6ZRP7 | 0.412 | -0.694 |
| RBM27 | Q9P2N5 | 0.811 | -0.694 |
| SS18 | Q15532 | 0.808 | -0.694 |
| ANXA1 | P04083 | 0.801 | -0.695 |
| EXOSC7 | Q15024 | 0.645 | -0.696 |
| FANCD2 | Q9BXW9 | 0.515 | -0.697 |
| DDX10 | Q13206 | 0.705 | -0.698 |
| DDX42 | Q86XP3 | 0.493 | -0.698 |
| SLC4A7 | Q9Y6M7 | 0.824 | -0.698 |
| SMARCB1.1 | G5E975 | 0.808 | -0.698 |
| CD2BP2 | O95400 | 0.619 | -0.699 |
| JUNB | P17275 | 0.712 | -0.7 |
| AGL | P35573 | 0.689 | -0.701 |
| SREK1 | Q8WXA9 | 0.812 | -0.701 |
| AGK | Q53H12 | 0.477 | -0.702 |
| WDR53 | Q7Z5U6 | 0.722 | -0.702 |
| DLGAP5 | Q15398 | 0.434 | -0.703 |
| SARG | Q9BW04 | 0.69 | -0.704 |
| PPWD1 | Q96BP3 | 0.7 | -0.705 |
| TRERF1 | Q05GC8 | 0.831 | -0.706 |
| CNOT9 | Q92600 | 0.724 | -0.707 |
| HLTF | Q14527 | 0.801 | -0.707 |
| LMAN1 | P49257 | 0.812 | -0.709 |
| LSM14A | Q8ND56 | 0.305 | -0.709 |
| MRPL14 | Q6P1L8 | 0.661 | -0.709 |
| PAXBP1 | Q9Y5B6 | 0.788 | -0.709 |
| TUFM | P49411 | 0.792 | -0.709 |
| NRDC | B1AKJ5 | 0.787 | -0.711 |
| SLX4 | Q8IY92 | 0.8 | -0.711 |
| TADA3 | O75528 | 0.831 | -0.712 |
| TOMM70 | O94826 | 0.758 | -0.713 |
| RRP8 | O43159 | 0.805 | -0.714 |
| GATAD2B | Q8WXI9 | 0.679 | -0.715 |
| BRPF3 | Q9ULD4 | 0.696 | -0.717 |
| STAG2 | Q8N3U4 | 0.585 | -0.718 |

| | | | |
|---|---|---|---|
| TBC1D30 | Q9Y2I9 | 0.808 | -0.718 |
| METTL3 | Q86U44 | 0.779 | -0.721 |
| MLLT10 | P55197 | 0.629 | -0.721 |
| EIF2B3 | Q9NR50 | 0.687 | -0.722 |
| LARP7 | Q4G0J3 | 0.699 | -0.722 |
| NHP2 | Q9NX24 | 0.592 | -0.722 |
| HDAC2 | Q92769 | 0.386 | -0.724 |
| ELF1 | P32519 | 0.741 | -0.725 |
| PEPD | P12955 | 0.625 | -0.726 |
| SUCLA2 | Q9P2R7 | 0.399 | -0.726 |
| OXA1L | J3KNA0 | 0.511 | -0.727 |
| MUTYH | E5KP25 | 0.739 | -0.728 |
| CTU2 | H3BSW6 | 0.717 | -0.729 |
| GRB7 | Q14451 | 0.783 | -0.729 |
| PAPOLG | Q9BWT3 | 0.73 | -0.729 |
| RBM26.1 | A0A087X0H9 | 0.747 | -0.729 |
| GPRASP2 | Q96D09 | 0.631 | -0.73 |
| PPP1R8 | Q12972 | 0.676 | -0.731 |
| TAOK1 | Q7L7X3 | 0.763 | -0.732 |
| SYAP1 | Q96A49 | 0.757 | -0.733 |
| CDK5RAP3 | Q96JB5 | 0.488 | -0.735 |
| PCYT2 | Q99447 | 0.614 | -0.735 |
| APPL1 | Q9UKG1 | 0.77 | -0.736 |
| DRG1 | Q9Y295 | 0.57 | -0.737 |
| MCM8 | Q9UJA3 | 0.774 | -0.737 |
| PSMC5 | P62195 | 0.758 | -0.737 |
| PTPN6 | P29350 | 0.509 | -0.737 |
| FLNC | Q14315 | 0.581 | -0.739 |
| TFIP11 | Q9UBB9 | 0.765 | -0.74 |
| AFDN.1 | H0Y7R8 | 0.768 | -0.741 |
| MRPL3 | E7ETU7 | 0.519 | -0.741 |
| TTC27 | Q6P3X3 | 0.604 | -0.741 |
| NBN | O60934 | 0.365 | -0.742 |
| SEL1L | Q9UBV2 | 0.687 | -0.742 |
| MAZ | P56270 | 0.265 | -0.743 |
| VPS52 | Q8N1B4 | 0.746 | -0.744 |
| CHSY1 | Q86X52 | 0.633 | -0.745 |
| PIP4K2A | P48426 | 0.43 | -0.745 |
| TMEM214 | Q6NUQ4 | 0.364 | -0.748 |
| RAD54L2 | Q9Y4B4 | 0.724 | -0.749 |
| DPY19L1 | A0A1B0GW05 | 0.363 | -0.754 |
| ILKAP | Q9H0C8 | 0.749 | -0.754 |
| MT-CO2 | P00403 | 0.802 | -0.754 |
| PRDM16 | Q9HAZ2 | 0.685 | -0.754 |
| SMARCC2.1 | F8VXC8 | 0.392 | -0.754 |
| PRMT1 | Q99873 | 0.75 | -0.755 |
| CCDC47 | Q96A33 | 0.448 | -0.756 |

| | | | |
|---|---|---|---|
| NAP1L4 | Q99733 | 0.676 | -0.76 |
| CCPG1 | Q9ULG6 | 0.168 | -0.761 |
| EIF4G3 | A0A0U1RQK7 | 0.683 | -0.763 |
| ADAM15 | Q13444 | 0.78 | -0.764 |
| BRF1 | Q92994 | 0.583 | -0.765 |
| PREB | Q9HCU5 | 0.426 | -0.765 |
| COPS3 | Q9UNS2 | 0.73 | -0.767 |
| PNISR | Q8TF01 | 0.831 | -0.767 |
| CHD9 | Q3L8U1 | 0.505 | -0.771 |
| PHLDA1 | Q8WV24 | 0.606 | -0.771 |
| RBM26 | Q5T8P6 | 0.697 | -0.771 |
| TSR3 | Q9UJK0 | 0.689 | -0.771 |
| RBMX2 | Q9Y388 | 0.749 | -0.772 |
| FH | P07954 | 0.648 | -0.773 |
| RFC2 | P35250 | 0.516 | -0.775 |
| CDK13 | Q14004 | 0.795 | -0.777 |
| INTS3 | Q68E01 | 0.448 | -0.777 |
| SNX5 | Q9Y5X3 | 0.757 | -0.777 |
| STXBP1 | P61764 | 0.591 | -0.777 |
| NCAPG | Q9BPX3 | 0.422 | -0.778 |
| AK2 | P54819 | 0.63 | -0.779 |
| PPP1R10 | Q96QC0 | 0.74 | -0.779 |
| ERF | P50548 | 0.757 | -0.78 |
| ANP32B | Q92688 | 0.83 | -0.782 |
| BTAF1 | A0A0A0MTH9 | 0.296 | -0.783 |
| CASP8AP2 | Q9UKL3 | 0.798 | -0.783 |
| GNE | Q9Y223 | 0.712 | -0.783 |
| NT5E | P21589 | 0.804 | -0.784 |
| MGST1 | P10620 | 0.748 | -0.785 |
| NEMF | O60524 | 0.799 | -0.785 |
| RIF1 | Q5UIP0 | 0.818 | -0.785 |
| DOCK5 | Q9H7D0 | 0.168 | -0.786 |
| PITRM1 | Q5JRX3 | 0.688 | -0.786 |
| C17orf80 | Q9BSJ5 | 0.534 | -0.787 |
| GCLC | P48506 | 0.709 | -0.787 |
| LGALS1 | P09382 | 0.793 | -0.787 |
| OGFOD3 | Q6PK18 | 0.624 | -0.787 |
| STK39 | Q9UEW8 | 0.277 | -0.789 |
| WDR44 | Q5JSH3 | 0.743 | -0.789 |
| CXADR | P78310 | 0.579 | -0.79 |
| DNM2 | P50570 | 0.635 | -0.79 |
| PRKCSH | K7ELL7 | 0.775 | -0.79 |
| DCTN2 | Q13561 | 0.777 | -0.791 |
| ACAT1 | P24752 | 0.794 | -0.794 |
| PZP | P20742 | 0.611 | -0.794 |
| RANBP3 | Q9H6Z4 | 0.581 | -0.794 |
| UBAP2L | Q14157 | 0.304 | -0.795 |

| | | | |
|---|---|---|---|
| CORO6 | Q6QEF8 | 0.684 | -0.798 |
| TOE1 | Q96GM8 | 0.468 | -0.798 |
| EXOG | Q9Y2C4 | 0.519 | -0.799 |
| WARS | P23381 | 0.781 | -0.799 |
| SLC29A1 | Q99808 | 0.758 | -0.804 |
| PUS7 | Q96PZ0 | 0.217 | -0.805 |
| SLC4A7.1 | H7C3C4 | 0.765 | -0.805 |
| IDH3A | P50213 | 0.827 | -0.807 |
| ZBTB24 | O43167 | 0.736 | -0.809 |
| APEX2 | Q9UBZ4 | 0.356 | -0.81 |
| PCF11 | O94913 | 0.786 | -0.811 |
| SERPINB9 | P50453 | 0.626 | -0.811 |
| MRPL15 | Q9P015 | 0.737 | -0.812 |
| RAD21 | O60216 | 0.427 | -0.812 |
| PAPD5.1 | H3BQM0 | 0.472 | -0.813 |
| PHB | P35232 | 0.801 | -0.814 |
| PSME3 | P61289 | 0.723 | -0.815 |
| CASC4 | Q6P4E1 | 0.585 | -0.817 |
| FAM111B | Q6SJ93 | 0.775 | -0.817 |
| MAP10 | Q9P2G4 | 0.617 | -0.819 |
| MCM4 | P33991 | 0.465 | -0.82 |
| METTL16 | Q86W50 | 0.145 | -0.82 |
| PRR11 | Q96HE9 | 0.703 | -0.82 |
| UBQLN1 | Q9UMX0 | 0.808 | -0.82 |
| ATIC | P31939 | 0.763 | -0.823 |
| SP1 | P08047 | 0.648 | -0.824 |
| SWAP70 | Q9UH65 | 0.201 | -0.824 |
| TOX4 | O94842 | 0.54 | -0.825 |
| ZNF16 | P17020 | 0.8 | -0.825 |
| A0A0U1RQF3 | A0A0U1RQF3 | 0.806 | -0.828 |
| ASPSCR1 | Q9BZE9 | 0.692 | -0.829 |
| SCAMP3 | O14828 | 0.799 | -0.829 |
| SMARCD1 | Q96GM5 | 0.775 | -0.83 |
| TMEM165 | Q9HC07 | 0.288 | -0.83 |
| ARHGDIA | J3QQX2 | 0.749 | -0.831 |
| FRMD6 | Q96NE9 | 0.721 | -0.831 |
| SUGT1 | Q9Y2Z0 | 0.692 | -0.831 |
| IRAK1 | P51617 | 0.72 | -0.832 |
| CARMIL2 | Q6F5E8 | 0.353 | -0.833 |
| CS | O75390 | 0.766 | -0.834 |
| PPM1B | O75688 | 0.46 | -0.836 |
| FKBP10 | Q96AY3 | 0.326 | -0.839 |
| ETFA | P13804 | 0.242 | -0.841 |
| ATP6V0D1 | F5GYQ1 | 0.772 | -0.842 |
| EXOC8 | Q8IYI6 | 0.699 | -0.842 |
| CWC27 | Q6UX04 | 0.621 | -0.843 |
| SCAF8 | A0A0A0MT33 | 0.781 | -0.843 |

| | | | |
|---|---|---|---|
| DNAJB4 | Q9UDY4 | 0.723 | -0.845 |
| IPO5 | O00410 | 0.489 | -0.846 |
| MGAT2 | Q10469 | 0.315 | -0.846 |
| CLASP1 | Q7Z460 | 0.708 | -0.847 |
| ZKSCAN8 | Q15776 | 0.699 | -0.85 |
| OPA1 | O60313 | 0.583 | -0.851 |
| THOC3 | Q96J01 | 0.697 | -0.851 |
| HLA-C | Q07000 | 0.416 | -0.852 |
| TRIM26 | Q12899 | 0.619 | -0.852 |
| DNMT1.1 | K7ELB1 | 0.644 | -0.853 |
| PPP4R2 | Q9NY27 | 0.746 | -0.853 |
| A6NIZ1 | A6NIZ1 | 0.781 | -0.854 |
| THAP11 | Q96EK4 | 0.739 | -0.854 |
| SYMPK | Q92797 | 0.182 | -0.855 |
| MTMR2 | Q13614 | 0.71 | -0.856 |
| ELP4 | G5E9D4 | 0.666 | -0.859 |
| SLC4A1AP | Q9BWU0 | 0.144 | -0.859 |
| OSBPL8.1 | F8VUA7 | 0.695 | -0.86 |
| FOXJ3 | Q9UPW0 | 0.317 | -0.861 |
| PIK3R1 | P27986 | 0.575 | -0.861 |
| WARS2 | Q9UGM6 | 0.693 | -0.861 |
| ECH1 | Q13011 | 0.786 | -0.862 |
| MCM2 | P49736 | 0.476 | -0.862 |
| PIP5K1A | A6PW57 | 0.682 | -0.862 |
| U2SURP.1 | E7ET15 | 0.768 | -0.862 |
| MACF1.2 | E9PLY5 | 0.726 | -0.865 |
| NUDT5 | A6NFX8 | 0.775 | -0.865 |
| TOPBP1 | Q92547 | 0.728 | -0.865 |
| CTDP1 | Q9Y5B0 | 0.789 | -0.866 |
| MECOM | Q03112 | 0.717 | -0.866 |
| RAB6D | Q53S08 | 0.747 | -0.866 |
| STX12 | Q86Y82 | 0.638 | -0.866 |
| MAST3 | O60307 | 0.814 | -0.867 |
| FDXR | A0A0A0MT64 | 0.739 | -0.869 |
| GOT2 | P00505 | 0.75 | -0.869 |
| ALDH16A1 | Q8IZ83 | 0.791 | -0.871 |
| MSH2 | P43246 | 0.493 | -0.871 |
| FCHSD2 | E9PG19 | 0.153 | -0.872 |
| ARFIP1 | P53367 | 0.715 | -0.873 |
| GALNT2 | Q10471 | 0.25 | -0.873 |
| AMBRA1 | Q9C0C7 | 0.431 | -0.874 |
| CLASP2 | E7EW49 | 0.392 | -0.874 |
| RAB2A | P61019 | 0.589 | -0.874 |
| PFKL | P17858 | 0.672 | -0.875 |
| TCEA1 | A0A1W2PPZ5 | 0.675 | -0.875 |
| FBXO28 | Q9NVF7 | 0.684 | -0.876 |
| RBBP7 | Q16576 | 0.587 | -0.876 |

| | | | |
|---|---|---|---|
| RCN1 | Q15293 | 0.635 | -0.877 |
| ORC2 | Q13416 | 0.647 | -0.879 |
| RPLP2 | P05387 | 0.756 | -0.879 |
| SERPINE2 | P07093 | 0.81 | -0.88 |
| PAF1 | Q8N7H5 | 0.268 | -0.881 |
| CDC6 | Q99741 | 0.426 | -0.886 |
| ME1 | P48163 | 0.668 | -0.887 |
| ASF1B | Q9NVP2 | 0.725 | -0.889 |
| DNAJC2 | Q99543 | 0.293 | -0.89 |
| MAN1A1 | P33908 | 0.706 | -0.89 |
| NAA16 | Q6N069 | 0.602 | -0.891 |
| ITGB5 | P18084 | 0.25 | -0.892 |
| DDN | O94850 | 0.648 | -0.893 |
| SMARCAD1 | Q9H4L7 | 0.303 | -0.893 |
| GGNBP2 | Q9H3C7 | 0.774 | -0.895 |
| CPSF2 | Q9P2I0 | 0.729 | -0.897 |
| TONSL | Q96HA7 | 0.736 | -0.897 |
| WWP1 | Q9H0M0 | 0.463 | -0.897 |
| IRF2BPL | Q9H1B7 | 0.624 | -0.898 |
| MAT2A | P31153 | 0.774 | -0.899 |
| SNX17 | Q15036 | 0.516 | -0.899 |
| LAMB3 | Q13751 | 0.511 | -0.9 |
| WDR4 | P57081 | 0.543 | -0.9 |
| SEC13 | P55735 | 0.631 | -0.904 |
| EIF2D | P41214 | 0.308 | -0.905 |
| ATG3 | Q9NT62 | 0.664 | -0.907 |
| ARNTL | O00327 | 0.604 | -0.908 |
| ARHGAP18 | Q8N392 | 0.222 | -0.91 |
| SMC5 | Q8IY18 | 0.735 | -0.911 |
| DNTTIP1 | Q9H147 | 0.593 | -0.916 |
| PNP | P00491 | 0.781 | -0.916 |
| RAB5C | P51148 | 0.788 | -0.916 |
| SINHCAF | Q9NP50 | 0.767 | -0.917 |
| STK10 | O94804 | 0.6 | -0.918 |
| MRE11 | P49959 | 0.149 | -0.92 |
| CCNT2 | O60583 | 0.785 | -0.921 |
| E2F3 | O00716 | 0.736 | -0.921 |
| DDOST | P39656 | 0.581 | -0.922 |
| DNAH5 | Q8TE73 | 0.134 | -0.923 |
| HSPA14 | Q0VDF9 | 0.176 | -0.928 |
| B4GALT5 | O43286 | 0.0899 | -0.93 |
| PDHA1 | P08559 | 0.786 | -0.93 |
| CKB | P12277 | 0.591 | -0.932 |
| PHF23 | Q9BUL5 | 0.664 | -0.932 |
| UBE4B | O95155 | 0.526 | -0.932 |
| DHCR24 | Q15392 | 0.183 | -0.933 |
| SNX6 | A0A0A0MRI2 | 0.791 | -0.933 |

| | | | |
|---|---|---|---|
| ZMYND8.1 | A0A087WVZ6 | 0.758 | -0.934 |
| NRBP1 | F8W6G1 | 0.689 | -0.936 |
| SLC39A14 | Q15043 | 0.619 | -0.937 |
| SP100 | P23497 | 0.238 | -0.938 |
| RASA1 | P20936 | 0.626 | -0.939 |
| SUZ12 | Q15022 | 0.767 | -0.94 |
| UQCRC1 | P31930 | 0.627 | -0.94 |
| EEPD1 | Q7L9B9 | 0.726 | -0.941 |
| ZNF114 | Q8NC26 | 0.539 | -0.942 |
| EFL1 | Q7Z2Z2 | 0.641 | -0.944 |
| NFATC2IP | Q8NCF5 | 0.709 | -0.946 |
| AAMP | C9JEH3 | 0.648 | -0.948 |
| CXorf56 | Q9H5V9 | 0.681 | -0.949 |
| UFL1 | O94874 | 0.658 | -0.949 |
| ITGA6 | P23229 | 0.707 | -0.952 |
| PFKM | P08237 | 0.496 | -0.952 |
| ANXA4 | P09525 | 0.764 | -0.953 |
| BCL7C | I3L1Q2 | 0.786 | -0.955 |
| HSPBP1 | Q9NZL4 | 0.639 | -0.956 |
| MCAM | P43121 | 0.745 | -0.956 |
| HDAC3 | O15379 | 0.18 | -0.957 |
| ACSL4 | O60488 | 0.704 | -0.958 |
| MICALL2 | Q8IY33 | 0.179 | -0.958 |
| ACTR1A | P61163 | 0.654 | -0.963 |
| CLPB.1 | F5GX99 | 0.105 | -0.965 |
| PGRMC2 | O15173 | 0.543 | -0.966 |
| PTGES3 | A0A087WYT3 | 0.752 | -0.967 |
| ARMCX3 | Q9UH62 | 0.264 | -0.97 |
| BCL2L13 | Q9BXK5 | 0.705 | -0.97 |
| RPAP1 | Q9BWH6 | 0.37 | -0.972 |
| F2 | P00734 | 0.419 | -0.974 |
| LANCL2 | Q9NS86 | 0.356 | -0.974 |
| PSMD13 | Q9UNM6 | 0.668 | -0.974 |
| CHEK2 | O96017 | 0.468 | -0.976 |
| RAB10 | P61026 | 0.728 | -0.976 |
| ZNF142.1 | H7C414 | 0.52 | -0.977 |
| PRPF39 | Q86UA1 | 0.736 | -0.98 |
| RHBDF2 | Q6PJF5 | 0.597 | -0.984 |
| HLA-C.2 | A2AEA2 | 0.667 | -0.985 |
| RAD51AP1 | Q96B01 | 0.675 | -0.986 |
| TERF2 | Q15554 | 0.192 | -0.986 |
| NONO | Q15233 | 0.0663 | -0.987 |
| MRGBP | Q9NV56 | 0.267 | -0.989 |
| SSR3 | Q9UNL2 | 0.529 | -0.99 |
| MPG | P29372 | 0.539 | -0.991 |
| ALDH5A1 | P51649 | 0.296 | -0.994 |
| BMP1 | P13497 | 0.661 | -0.994 |

| | | | |
|---|---|---|---|
| SMYD3 | Q9H7B4 | 0.309 | -0.998 |
| DTD1 | Q8TEA8 | 0.355 | -0.999 |
| CMBL | Q96DG6 | 0.46 | -1 |
| DGKZ | Q13574 | 0.698 | -1 |
| HMGN5 | P82970 | 0.118 | -1 |
| MGEA5 | O60502 | 0.58 | -1 |
| ZC3H4 | Q9UPT8 | 0.67 | -1 |
| ASNA1 | O43681 | 0.684 | -1.01 |
| GC | D6RF35 | 0.746 | -1.01 |
| GPATCH2L | Q9NWQ4 | 0.784 | -1.01 |
| HLA-A | A0A0G2JIF2 | 0.565 | -1.01 |
| MYO1B | O43795 | 0.424 | -1.01 |
| MYC | Q16591 | 0.231 | -1.02 |
| OTUB1 | F5GYJ8 | 0.715 | -1.02 |
| VPS18 | Q9P253 | 0.625 | -1.02 |
| ANP32A | P39687 | 0.702 | -1.03 |
| ARF3 | P61204 | 0.739 | -1.03 |
| HGH1 | Q9BTY7 | 0.603 | -1.03 |
| HMBS | P08397 | 0.713 | -1.03 |
| PLEKHH3 | Q7Z736 | 0.662 | -1.03 |
| ETV6 | P41212 | 0.775 | -1.04 |
| MAVS | Q7Z434 | 0.786 | -1.04 |
| PRMT3 | O60678 | 0.605 | -1.04 |
| A2M | P01023 | 0.696 | -1.05 |
| CIZ1 | F5H2X7 | 0.619 | -1.05 |
| EIF2B1 | Q14232 | 0.0635 | -1.05 |
| GORASP2 | Q9H8Y8 | 0.689 | -1.05 |
| KMT2E | Q8IZD2 | 0.746 | -1.05 |
| PTPN1 | P18031 | 0.741 | -1.05 |
| ZNF189 | O75820 | 0.211 | -1.05 |
| NTSR1 | P30989 | 0.0455 | -1.06 |
| BRD2 | A0A0G2JK44 | 0.729 | -1.06 |
| GOLM1 | Q8NBJ4 | 0.0548 | -1.06 |
| IRF3 | Q14653 | 0.117 | -1.06 |
| NCEH1 | A0A0A0MTJ9 | 0.736 | -1.07 |
| SARS2 | M0QWZ7 | 0.663 | -1.07 |
| SEC22B | O75396 | 0.624 | -1.07 |
| SRM | P19623 | 0.75 | -1.07 |
| TSN | Q15631 | 0.742 | -1.07 |
| USP19.1 | A0A0A0MR08 | 0.184 | -1.07 |
| CDCA5 | B5MBX0 | 0.288 | -1.08 |
| CNN3 | Q15417 | 0.718 | -1.08 |
| NELFB | A0A0X1KG71 | 0.671 | -1.08 |
| PIGS | Q96S52 | 0.66 | -1.08 |
| ZNF512 | Q96ME7 | 0.469 | -1.08 |
| GLS.1 | B8ZZA8 | 0.592 | -1.09 |
| MTA3 | Q9BTC8 | 0.697 | -1.09 |

| | | | |
|---|---|---|---|
| CLN3 | A0A024QZB8 | 0.0364 | -1.1 |
| MYSM1 | Q5VVJ2 | 0.689 | -1.1 |
| POLR1C | O15160 | 0.108 | -1.1 |
| PSD3 | Q9NYI0 | 0.743 | -1.1 |
| RRN3 | Q9NYV6 | 0.458 | -1.1 |
| SMURF2 | Q9HAU4 | 0.655 | -1.1 |
| STAU2 | Q9NUL3 | 0.686 | -1.1 |
| TTI2 | Q6NXR4 | 0.416 | -1.1 |
| XPO4 | F2Z2X4 | 0.294 | -1.1 |
| NOL7 | Q9UMY1 | 0.574 | -1.11 |
| RIN1 | Q13671 | 0.381 | -1.11 |
| RPUSD2 | Q8IZ73 | 0.213 | -1.11 |
| UNG | P13051 | 0.664 | -1.11 |
| HMG20A | Q9NP66 | 0.197 | -1.12 |
| LRRC41 | Q15345 | 0.396 | -1.12 |
| PITPNB | A0A0A0MSW4 | 0.78 | -1.12 |
| SHCBP1 | Q8NEM2 | 0.615 | -1.12 |
| SIRT7 | Q9NRC8 | 0.248 | -1.12 |
| THUMPD3 | Q9BV44 | 0.461 | -1.12 |
| URGCP | Q8TCY9 | 0.119 | -1.12 |
| ARVCF | O00192 | 0.266 | -1.13 |
| ATE1 | O95260 | 0.357 | -1.13 |
| NAPRT | Q6XQN6 | 0.593 | -1.13 |
| TRAFD1 | O14545 | 0.432 | -1.13 |
| GNL2.1 | H0YG10 | 0.0932 | -1.14 |
| PRKRA | O75569 | 0.701 | -1.14 |
| RTN4 | Q9NQC3 | 0.758 | -1.14 |
| STUB1 | Q9UNE7 | 0.602 | -1.14 |
| UCK2 | Q9BZX2 | 0.603 | -1.14 |
| EML3 | Q32P44 | 0.501 | -1.15 |
| FAM208A | Q9UK61 | 0.616 | -1.15 |
| TRIOBP.1 | F6TR96 | 0.629 | -1.15 |
| USP27X | A6NNY8 | 0.132 | -1.15 |
| WASF2 | Q9Y6W5 | 0.0149 | -1.16 |
| BAG5 | Q9UL15 | 0.123 | -1.16 |
| RAVER2 | Q9HCJ3 | 0.481 | -1.16 |
| RIOX2 | Q8IUF8 | 0.609 | -1.16 |
| LAMB1 | G3XAI2 | 0.596 | -1.17 |
| ZBTB6 | Q15916 | 0.129 | -1.17 |
| ACADM | Q5T4U5 | 0.674 | -1.18 |
| AHI1 | Q8N157 | 0.642 | -1.18 |
| DFFA | O00273 | 0.504 | -1.18 |
| ADAP1 | O75689 | 0.368 | -1.19 |
| ATXN7L3 | Q14CW9 | 0.321 | -1.19 |
| EZH2 | Q15910 | 0.708 | -1.19 |
| GRHPR | Q9UBQ7 | 0.611 | -1.19 |
| KDM5A | P29375 | 0.773 | -1.19 |

| | | | |
|---|---|---|---|
| PKM.1 | H3BR70 | 0.717 | -1.19 |
| RANBP10 | R4GMX8 | 0.222 | -1.19 |
| SAV1 | Q9H4B6 | 0.161 | -1.19 |
| TSNAX | Q99598 | 0.388 | -1.19 |
| HECTD3 | Q5T447 | 0.559 | -1.2 |
| HSDL2 | Q6YN16 | 0.4 | -1.2 |
| TBC1D9B | Q66K14 | 0.43 | -1.2 |
| ACAA1 | P09110 | 0.291 | -1.21 |
| BRD9 | Q9H8M2 | 0.717 | -1.21 |
| DARS2 | Q6PI48 | 0.402 | -1.21 |
| GALNT12 | Q8IXK2 | 0.228 | -1.21 |
| FAF2 | Q96CS3 | 0.566 | -1.22 |
| FAM76B | Q5HYJ3 | 0.569 | -1.22 |
| NSFL1C | Q9UNZ2 | 0.672 | -1.22 |
| SYT7 | O43581 | 0.494 | -1.22 |
| ADK | P55263 | 0.762 | -1.23 |
| CEP250 | Q9BV73 | 0.598 | -1.23 |
| COPG1 | Q9Y678 | 0.329 | -1.23 |
| IRF2BP1 | Q8IU81 | 0.487 | -1.23 |
| LIPE | Q05469 | 0.579 | -1.23 |
| TARDBP | Q13148 | 0.235 | -1.23 |
| ABCC2 | Q92887 | 0.081 | -1.24 |
| ATP1B3 | P54709 | 0.647 | -1.24 |
| C1orf174 | Q8IYL3 | 0.185 | -1.24 |
| CRAT | P43155 | 0.621 | -1.24 |
| HEATR3 | Q7Z4Q2 | 0.0549 | -1.24 |
| MARK4 | Q96L34 | 0.409 | -1.24 |
| CHMP7 | Q8WUX9 | 0.592 | -1.25 |
| CUL4A | Q13619 | 0.335 | -1.25 |
| EFHD2 | Q96C19 | 0.665 | -1.25 |
| GALNT3 | Q14435 | 0.682 | -1.25 |
| PSMG2 | Q969U7 | 0.118 | -1.25 |
| TM9SF2 | Q99805 | 0.593 | -1.25 |
| TNFAIP2 | Q03169 | 0.162 | -1.25 |
| BCAP31 | P51572 | 0.736 | -1.26 |
| BUD31 | P41223 | 0.465 | -1.26 |
| FAF1 | Q9UNN5 | 0.175 | -1.26 |
| TBCC | Q15814 | 0.254 | -1.26 |
| TRPC4AP | Q8TEL6 | 0.153 | -1.26 |
| XPO6 | Q96QU8 | 0.0969 | -1.26 |
| SFPQ | P23246 | 0.00617 | -1.27 |
| USP6NL | X6RAB3 | 0.00906 | -1.27 |
| ZBTB46 | Q86UZ6 | 0.0286 | -1.27 |
| CCNL2 | Q96S94 | 0.688 | -1.27 |
| MIA3 | Q5JRA6 | 0.315 | -1.27 |
| RBL1 | P28749 | 0.378 | -1.27 |
| TMX2 | Q9Y320 | 0.517 | -1.27 |

| | | | |
|---|---|---|---|
| CENPC | Q03188 | 0.647 | -1.28 |
| PHF3 | Q92576 | 0.467 | -1.28 |
| TIMELESS | Q9UNS1 | 0.703 | -1.28 |
| ABL1.1 | R4GRW0 | 0.315 | -1.29 |
| ASTE1 | D6RG30 | 0.465 | -1.29 |
| HNRNPH3 | P31942 | 0.514 | -1.29 |
| RNH1 | P13489 | 0.441 | -1.29 |
| CPT1A | P50416 | 0.0116 | -1.3 |
| AGPAT5 | Q9NUQ2 | 0.0528 | -1.3 |
| MST1R | Q04912 | 0.417 | -1.3 |
| DENR | O43583 | 0.406 | -1.31 |
| MARVELD3 | Q96A59 | 0.074 | -1.31 |
| MUT | P22033 | 0.428 | -1.31 |
| PSMB3 | P49720 | 0.0724 | -1.31 |
| SEPHS1 | P49903 | 0.196 | -1.31 |
| BRD7 | Q9NPI1 | 0.713 | -1.32 |
| SUCO | Q9UBS9 | 0.492 | -1.32 |
| TAF2.1 | H0YEX2 | 0.646 | -1.32 |
| AHSG | P02765 | 0.715 | -1.33 |
| HSDL1 | Q3SXM5 | 0.307 | -1.33 |
| MPZL1 | O95297 | 0.0606 | -1.33 |
| CPT2 | A0A1B0GVF3 | 0.652 | -1.34 |
| PLEKHG4 | Q58EX7 | 0.659 | -1.34 |
| INTS13 | Q9NVM9 | 0.0416 | -1.35 |
| CKMT1A | P12532 | 0.643 | -1.35 |
| HSF1 | Q00613 | 0.596 | -1.35 |
| SNRPA | P09012 | 0.058 | -1.35 |
| CLP1 | E9PL17 | 0.00482 | -1.36 |
| LMF2 | Q9BU23 | 0.0467 | -1.36 |
| PHF12 | Q96QT6 | 0.747 | -1.36 |
| LTF | P02788 | 0.63 | -1.37 |
| PARS2 | Q7L3T8 | 0.0502 | -1.37 |
| VPS36 | Q86VN1 | 0.23 | -1.37 |
| MTERF3 | Q96E29 | 0.0369 | -1.38 |
| CHORDC1 | Q9UHD1 | 0.731 | -1.38 |
| PSRC1 | Q6PGN9 | 0.428 | -1.38 |
| RNF169 | Q8NCN4 | 0.613 | -1.38 |
| ZCCHC4 | Q9H5U6 | 0.387 | -1.38 |
| MCPH1 | Q8NEM0 | 0.472 | -1.39 |
| RNF138 | Q8WVD3 | 0.461 | -1.39 |
| ARHGAP45 | Q92619 | 0.538 | -1.4 |
| CDV3 | Q9UKY7 | 0.42 | -1.4 |
| HMGB3 | O15347 | 0.672 | -1.4 |
| PRDX2 | P32119 | 0.605 | -1.4 |
| SNRNP48 | Q6IEG0 | 0.235 | -1.4 |
| CORO7-PAM16 | A0A0A6YYL4 | 0.0189 | -1.41 |
| CC2D1B.1 | Q5T0F9 | 0.23 | -1.41 |

| | | | |
|---|---|---|---|
| ESRRA | P11474 | 0.52 | -1.41 |
| PRRC1 | Q96M27 | 0.318 | -1.41 |
| PSMA5 | P28066 | 0.683 | -1.41 |
| TLE3 | H0YL70 | 0.093 | -1.41 |
| HBA1 | P69905 | 0.312 | -1.42 |
| UBLCP1 | Q8WVY7 | 0.148 | -1.42 |
| ZNF496 | Q96IT1 | 0.107 | -1.42 |
| TJP1.2 | H0Y3R8 | 0.0176 | -1.43 |
| AIFM2 | Q9BRQ8 | 0.434 | -1.43 |
| RPUSD4 | Q96CM3 | 0.314 | -1.43 |
| TIAL1 | Q01085 | 0.564 | -1.43 |
| TRIM2 | Q9C040 | 0.502 | -1.43 |
| ARHGAP11A | Q6P4F7 | 0.0187 | -1.44 |
| DRAP1 | C9JCC6 | 0.211 | -1.44 |
| AKR1B1 | P15121 | 0.305 | -1.45 |
| FBXO22 | Q8NEZ5 | 0.503 | -1.45 |
| NCOR2 | Q9Y618 | 0.396 | -1.45 |
| TSSC4 | Q9Y5U2 | 0.0101 | -1.46 |
| CLPP | Q16740 | 0.176 | -1.46 |
| MAP4K3.1 | F5H5A3 | 0.17 | -1.46 |
| REXO4 | Q9GZR2 | 0.555 | -1.46 |
| TTF1 | Q15361 | 0.587 | -1.46 |
| DDRGK1 | Q96HY6 | 0.266 | -1.48 |
| PDCD10 | Q9BUL8 | 0.186 | -1.48 |
| BICD2 | Q8TD16 | 0.259 | -1.49 |
| GALE | Q14376 | 0.327 | -1.49 |
| ZNF579 | Q8NAF0 | 0.463 | -1.49 |
| MAGI1.1 | A0A087WT53 | 0.047 | -1.5 |
| ZFPL1 | O95159 | 0.00774 | -1.5 |
| PQBP1 | O60828 | 0.443 | -1.5 |
| STX7 | O15400 | 0.0609 | -1.5 |
| DEPDC7 | Q96QD5 | 0.00188 | -1.51 |
| FTSJ1 | Q9UET6 | 0.00757 | -1.51 |
| AP3M2 | P53677 | 0.468 | -1.51 |
| IFRD1 | O00458 | 0.0788 | -1.51 |
| MLLT1 | Q03111 | 0.625 | -1.51 |
| CBX8 | Q9HC52 | 0.511 | -1.52 |
| LSM14B.1 | Q5TBP9 | 0.0167 | -1.53 |
| B3GAT3 | O94766 | 0.063 | -1.53 |
| BAZ2A | Q9UIF9 | 0.0915 | -1.53 |
| TTC4 | O95801 | 0.329 | -1.53 |
| CCDC50 | Q8IVM0 | 0.448 | -1.54 |
| FBXO5 | Q9UKT4 | 0.0559 | -1.54 |
| GPATCH2 | Q9NW75 | 0.55 | -1.54 |
| PGM1 | P36871 | 0.468 | -1.55 |
| POLA2 | Q14181 | 0.0627 | -1.55 |
| SPINT1 | O43278 | 0.561 | -1.55 |

| | | | |
|---|---|---|---|
| GAN | Q9H2C0 | 0.304 | -1.56 |
| REST | Q13127 | 0.456 | -1.56 |
| ALG3 | Q92685 | 0.0186 | -1.57 |
| LIMK2 | P53671 | 0.0184 | -1.57 |
| ARHGEF2.1 | V9GYF0 | 0.151 | -1.57 |
| DYNC1I2.1 | E7EQL5 | 0.254 | -1.57 |
| OTUD6B | A0A087X0W9 | 0.00181 | -1.58 |
| BCKDK | O14874 | 0.363 | -1.58 |
| BZW2 | Q9Y6E2 | 0.169 | -1.58 |
| GNAQ | P50148 | 0.233 | -1.58 |
| RANBP9 | Q96S59 | 0.237 | -1.58 |
| ZNF827 | Q17R98 | 0.544 | -1.58 |
| NHLRC2 | Q8NBF2 | 0.244 | -1.59 |
| ZC3H7A | Q8IWR0 | 0.394 | -1.6 |
| GOLIM4 | O00461 | 0.0225 | -1.61 |
| KDELR1 | P24390 | 0.117 | -1.61 |
| RAF1 | P04049 | 0.0599 | -1.61 |
| NARS2 | Q96I59 | 0.219 | -1.62 |
| TGM5 | A0A087WVS3 | 0.0247 | -1.63 |
| GMCL1 | Q96IK5 | 0.135 | -1.63 |
| NOL8 | Q76FK4 | 0.647 | -1.63 |
| TCF25.1 | H3BQ53 | 0.154 | -1.63 |
| EPHB4 | P54760 | 0.211 | -1.64 |
| BZW1 | Q7L1Q6 | 0.0429 | -1.65 |
| GALNT1 | Q10472 | 0.00224 | -1.65 |
| INTS12 | Q96CB8 | 0.00133 | -1.65 |
| CENPV | Q7Z7K6 | 0.0314 | -1.66 |
| RAD54L | Q92698 | 0.041 | -1.66 |
| GPAM | Q9HCL2 | 0.23 | -1.66 |
| OCIAD1 | Q9NX40 | 0.314 | -1.66 |
| TIGD5 | Q53EQ6 | 0.077 | -1.66 |
| AIP | O00170 | 0.262 | -1.67 |
| NSRP1 | Q9H0G5 | 0.19 | -1.67 |
| EPB41L4B | Q9H329 | 0.00675 | -1.69 |
| PLA2G4A | P47712 | 0.035 | -1.69 |
| PLEKHA1 | Q9HB21 | 0.000875 | -1.69 |
| POU2F1 | P14859 | 0.0185 | -1.69 |
| C2CD2 | Q9Y426 | 0.492 | -1.69 |
| CTU1 | Q7Z7A3 | 0.0833 | -1.69 |
| LSS | P48449 | 0.216 | -1.69 |
| TUBAL3 | A6NHL2 | 0.0799 | -1.69 |
| FAM114A2 | Q9NRY5 | 0.277 | -1.7 |
| MFN1 | Q8IWA4 | 0.123 | -1.7 |
| NLN | Q9BYT8 | 0.557 | -1.7 |
| RBM23 | Q86U06 | 0.234 | -1.7 |
| RPRD1B | Q9NQG5 | 0.0507 | -1.71 |
| CSNK1A1.1 | D6REM4 | 0.0325 | -1.72 |

| | | | |
|---|---|---|---|
| HMOX2 | A0A087WT44 | 0.271 | -1.72 |
| NIPSNAP2 | O75323 | 0.254 | -1.72 |
| PSME4 | Q14997 | 0.0641 | -1.72 |
| SNX4 | O95219 | 0.15 | -1.72 |
| ALAS1 | P13196 | 0.128 | -1.73 |
| IST1 | P53990 | 0.148 | -1.73 |
| MTF2 | Q9Y483 | 0.124 | -1.73 |
| ZNF316 | A6NFI3 | 0.629 | -1.73 |
| MYCBP2.1 | H7C3U4 | 0.0119 | -1.74 |
| CHID1 | Q9BWS9 | 0.314 | -1.74 |
| HMGXB3 | Q12766 | 0.0547 | -1.74 |
| NDUFA9 | Q16795 | 0.186 | -1.74 |
| NFKB1 | P19838 | 0.272 | -1.74 |
| MAP2K1 | Q02750 | 0.418 | -1.75 |
| EXOC6 | Q8TAG9 | 0.0401 | -1.76 |
| TMX1 | Q9H3N1 | 0.639 | -1.76 |
| ACOX2 | Q99424 | 0.195 | -1.77 |
| DAD1 | P61803 | 0.264 | -1.77 |
| UQCRC2 | P22695 | 0.192 | -1.77 |
| PIN4 | Q9Y237 | 0.0171 | -1.78 |
| BCKDHB | P21953 | 0.428 | -1.78 |
| PRKACB | A0A0A0MS54 | 0.178 | -1.79 |
| TRMT5 | Q32P41 | 0.089 | -1.79 |
| BTF3.1 | D6RDG3 | 0.00974 | -1.8 |
| ZNF638.1 | A0A096LNQ0 | 0.0194 | -1.82 |
| DHRS7 | H0YJ66 | 0.0638 | -1.82 |
| HMGB2 | P26583 | 0.668 | -1.82 |
| SLBP | F8W8D3 | 0.000675 | -1.83 |
| HSPA12A | O43301 | 0.0681 | -1.83 |
| IDH3G | P51553 | 0.3 | -1.83 |
| PHF19 | A0A087X169 | 0.147 | -1.86 |
| NEO1 | Q92859 | 0.000409 | -1.87 |
| DPP4 | P27487 | 0.141 | -1.88 |
| PGP | A6NDG6 | 0.259 | -1.88 |
| PTGFRN | Q9P2B2 | 0.222 | -1.88 |
| ITPK1 | Q13572 | 0.0242 | -1.89 |
| P3H3 | Q8IVL6 | 3.03E-06 | -1.89 |
| SHANK1 | H9KV90 | 0.0384 | -1.89 |
| UBAP2L.1 | H0Y5H6 | 0.000272 | -1.89 |
| LMAN2 | Q12907 | 0.463 | -1.91 |
| RFX1 | P22670 | 0.364 | -1.92 |
| NOLC1.1 | S4R341 | 0.288 | -1.94 |
| KAT6A | Q92794 | 0.0735 | -1.95 |
| PARK7 | Q99497 | 0.484 | -1.95 |
| SAAL1 | E9PRZ1 | 0.0605 | -1.95 |
| ABRAXAS1 | Q6UWZ7 | 0.028 | -1.96 |
| DNAJB12 | J3KPS0 | 4.83E-05 | -1.96 |

| | | | |
|---|---|---|---|
| AP2A2.1 | A0A0G2JQM1 | 0.105 | -1.96 |
| ARIH1 | Q9Y4X5 | 0.226 | -1.96 |
| NUDCD3 | Q8IVD9 | 0.212 | -1.96 |
| LAMA3 | Q16787 | 0.0166 | -1.97 |
| LONP2 | Q86WA8 | 0.0356 | -1.97 |
| ESD | P10768 | 0.433 | -1.97 |
| PPME1 | Q9Y570 | 0.243 | -1.97 |
| INTS9 | Q9NV88 | 0.0101 | -1.98 |
| WDR74 | Q6RFH5 | 3.92E-05 | -1.98 |
| LAMC1 | P11047 | 0.148 | -1.98 |
| PPID | Q08752 | 0.182 | -1.98 |
| RPIA | P49247 | 0.0686 | -1.98 |
| LPCAT4 | Q643R3 | 0.000202 | -1.99 |
| PTPN2 | P17706 | 8.92E-06 | -2 |
| SPG7 | Q9UQ90 | 0.0197 | -2 |
| PPP1R16A | Q96I34 | 0.141 | -2.01 |
| SMC6 | Q96SB8 | 0.305 | -2.03 |
| FXR1.1 | E9PFF5 | 0.00474 | -2.04 |
| MAPK8IP3 | A0A087WYG2 | 0.00897 | -2.04 |
| PRDX3 | P30048 | 0.0255 | -2.04 |
| H0YHG0 | H0YHG0 | 0.306 | -2.04 |
| SETD5 | E7EWN3 | 0.424 | -2.04 |
| TMEM161A | Q9NX61 | 0.0178 | -2.05 |
| SF3B4 | Q15427 | 0.353 | -2.05 |
| RALA | P11233 | 0.272 | -2.06 |
| ESRP2 | Q9H6T0 | 0.0818 | -2.09 |
| SLC25A24 | Q6NUK1 | 0.223 | -2.09 |
| DGKD | Q16760 | 0.0292 | -2.1 |
| MAP2K7 | O14733 | 0.00983 | -2.1 |
| WIPF2 | Q8TF74 | 0.0144 | -2.1 |
| UBA3 | Q8TBC4 | 0.256 | -2.11 |
| CRYZ | Q08257 | 0.00835 | -2.12 |
| DHX32 | Q7L7V1 | 0.00911 | -2.14 |
| SCRN1 | Q12765 | 0.095 | -2.14 |
| UCHL5 | Q5LJA9 | 0.0297 | -2.15 |
| SP110 | Q9HB58 | 0.064 | -2.15 |
| LRR1 | Q96L50 | 0.00197 | -2.16 |
| TYMP | P19971 | 0.00521 | -2.16 |
| ZNF408 | Q9H9D4 | 0.291 | -2.16 |
| SMAD2 | Q15796 | 0.0383 | -2.19 |
| SLF2 | Q8IX21 | 0.0034 | -2.2 |
| PCBP2.1 | F8VRH0 | 0.153 | -2.2 |
| PCNP | Q8WW12 | 0.00212 | -2.21 |
| ACAT2 | Q9BWD1 | 0.02 | -2.22 |
| OSBPL7 | Q9BZF2 | 0.265 | -2.22 |
| UAP1 | Q16222 | 0.0533 | -2.22 |
| GLB1L2 | Q8IW92 | 0.000319 | -2.23 |

| | | | |
|---|---|---|---|
| SFN | P31947 | 0.359 | -2.23 |
| INTS5 | Q6P9B9 | 0.000477 | -2.24 |
| CLASP1.1 | H0Y5T1 | 0.426 | -2.24 |
| MBTPS1 | Q14703 | 0.224 | -2.24 |
| SHQ1 | Q6PI26 | 0.0054 | -2.25 |
| NUBP2 | H3BNS4 | 0.0419 | -2.27 |
| SAMD4B | Q5PRF9 | 0.0904 | -2.27 |
| TRIM3 | O75382 | 0.0904 | -2.27 |
| RETSAT | Q6NUM9 | 0.0395 | -2.28 |
| SMURF1 | Q9HCE7 | 0.000359 | -2.28 |
| MIPEP | Q99797 | 0.373 | -2.28 |
| TMCC1 | O94876 | 0.117 | -2.29 |
| LAMP2 | P13473 | 0.00172 | -2.3 |
| ANXA7 | P20073 | 0.377 | -2.3 |
| SHMT1 | P34896 | 0.0337 | -2.32 |
| FAM49B | Q9NUQ9 | 0.245 | -2.32 |
| RHOA | P61586 | 0.0958 | -2.33 |
| TAF1A | Q15573 | 8.00E-04 | -2.34 |
| RPP40 | O75818 | 0.0442 | -2.35 |
| NAF1 | Q96HR8 | 0.352 | -2.35 |
| CHST14 | Q8NCH0 | 0.000115 | -2.37 |
| UBA5 | Q9GZZ9 | 0.0215 | -2.4 |
| FAM110C | Q1W6H9 | 1.36E-06 | -2.46 |
| FAR1 | Q8WVX9 | 8.48E-05 | -2.48 |
| HS6ST2 | Q96MM7 | 2.13E-07 | -2.48 |
| UBXN8 | A0A087WT23 | 0.0278 | -2.48 |
| E4F1 | Q66K89 | 0.263 | -2.49 |
| SLC1A3 | P43003 | 0.00092 | -2.5 |
| BRMS1 | G5E9I4 | 0.000998 | -2.51 |
| TIGAR | Q9NQ88 | 0.0438 | -2.51 |
| LDB1 | Q86U70 | 4.60E-07 | -2.53 |
| PML | P29590 | 1.80E-05 | -2.53 |
| PER2 | O15055 | 0.00148 | -2.55 |
| PBK | Q96KB5 | 0.0151 | -2.56 |
| USP3 | Q9Y6I4 | 2.83E-07 | -2.56 |
| TBC1D15 | Q8TC07 | 0.0449 | -2.57 |
| ARID5B | Q14865 | 0.0246 | -2.58 |
| ABCD1 | P33897 | 2.42E-07 | -2.59 |
| CDC42EP1 | Q00587 | 0.000775 | -2.6 |
| SCML2 | Q9UQR0 | 2.60E-05 | -2.6 |
| APRT | P07741 | 0.0894 | -2.61 |
| GNA12 | Q03113 | 0.000589 | -2.62 |
| TBP | P20226 | 0.0168 | -2.62 |
| S100A8 | P05109 | 0.0205 | -2.64 |
| TMEM131L | A2VDJ0 | 0.0551 | -2.64 |
| COPS2 | P61201 | 0.166 | -2.65 |
| AAR2 | A2A2Q9 | 8.40E-07 | -2.66 |

| | | | |
|---|---|---|---|
| GPAT3 | Q53EU6 | 0.013 | -2.66 |
| HKDC1 | Q2TB90 | 0.00118 | -2.66 |
| MAP2K4 | P45985 | 0.0057 | -2.66 |
| ZRANB2 | O95218 | 0.00645 | -2.66 |
| RTCA | O00442 | 0.0366 | -2.68 |
| ERMP1 | Q7Z2K6 | 0.00723 | -2.7 |
| SAP30BP | J3QQJ0 | 0.0414 | -2.7 |
| CEP170.2 | E7EWM2 | 1.68E-05 | -2.72 |
| MAP2K2 | P36507 | 0.0685 | -2.72 |
| THBS1 | P07996 | 7.47E-05 | -2.74 |
| ZRANB3 | Q5FWF4 | 0.184 | -2.74 |
| CD97 | P48960 | 0.00789 | -2.76 |
| AUP1 | Q9Y679 | 0.00265 | -2.79 |
| FBXL19 | Q6PCT2 | 0.0494 | -2.8 |
| ACOX1 | Q15067 | 0.003 | -2.84 |
| MVD | P53602 | 0.0239 | -2.84 |
| ZC3H8 | A0A0C4DGZ1 | 1.08E-07 | -2.84 |
| GALNT7 | Q86SF2 | 0.011 | -2.85 |
| ABCC5 | O15440 | 0.0019 | -2.87 |
| CLU | P10909 | 0.0532 | -2.88 |
| FDXR.1 | P22570 | 0.000631 | -2.89 |
| TOLLIP | Q9H0E2 | 1.47E-05 | -2.89 |
| GSE1.1 | A0A1B0GUL3 | 1.27E-07 | -2.91 |
| SIRT1 | Q96EB6 | 0.00224 | -2.91 |
| POLR3F | Q9H1D9 | 8.00E-06 | -2.95 |
| PPP2R5E | Q16537 | 5.75E-06 | -2.99 |
| ZNF865 | P0CJ78 | 0.00915 | -2.99 |
| NCK1 | P16333 | 1.14E-10 | -3.06 |
| TTC26 | A0AVF1 | 7.37E-06 | -3.06 |
| SNX33 | Q8WV41 | 0.00517 | -3.09 |
| NARF | A0A088AWN8 | 0.000118 | -3.14 |
| PDHB | P11177 | 0.0456 | -3.15 |
| SMARCD3 | Q6STE5 | 0.000267 | -3.18 |
| UBXN7 | O94888 | 8.78E-05 | -3.19 |
| ANP32E | Q9BTT0 | 0.0633 | -3.21 |
| ORC4 | O43929 | 4.94E-05 | -3.25 |
| PPP1R7 | Q15435 | 4.70E-06 | -3.3 |
| POLD2 | A0A087WWF6 | 2.45E-05 | -3.33 |
| LAMB2 | P55268 | 0.00219 | -3.34 |
| CD46 | P15529 | 0.0171 | -3.54 |
| UROD | P06132 | 1.14E-05 | -3.59 |
| RAP1GDS1 | P52306 | 0.00408 | -3.63 |
| SURF6 | O75683 | 0.0303 | -3.67 |
| CDT1 | Q9H211 | 1.57E-05 | -3.69 |
| CBWD6 | Q4V339 | 4.53E-06 | -3.71 |
| HEXIM2 | Q96MH2 | 7.26E-08 | -3.79 |
| RNF213 | A0A0A0MTC1 | 8.11E-14 | -3.81 |

| | | | |
|---|---|---|---|
| HSP90AB4P | Q58FF6 | 9.01E-07 | -3.86 |
| CAPN6 | Q9Y6Q1 | 0.000241 | -3.94 |
| WBP4 | O75554 | 2.91E-12 | -3.96 |
| COPS5 | Q92905 | 0.000111 | -4.07 |
| ACACA.1 | A0A087WYK6 | 4.76E-07 | -4.22 |
| TUBD1 | Q9UJT1 | 8.11E-14 | -4.24 |
| CAVIN2 | O95810 | 8.29E-05 | -4.35 |
| F11R | Q9Y624 | 0.00216 | -4.8 |
