## Supplemental Table 3 for "Anchorage-independent cell proliferation promoted by fascin’s F-actin bundling"

### Table S3: Sequences of oligonucleotides used in this study

| Primer | Sequence (5' - 3') |
|---|---|
| FSCN1 gRNA #1 Fwd | CACCGTACCTGACGGCCGAGGCGTT |
| FSCN1 gRNA #1 Rev | AAACAACGCCTCGGCCGTCAGGTAC |
| FSCN1 gRNA #2 Fwd | CACCGCAACAAGTACCTGACGGCCG |
| FSCN1 gRNA #2 Rev | AAACCGGCCGTCAGGTACTTGTTGC |
| Fascin_BamH1_Fwd | attggatccACCGCCAACGGCACAGCCG |
| Fascin_XhoI_Rev | attctcgagGTACTCCCAGAGCGAGGCGG |
| FSCN1_homo_L | gccaacgagaggaacgtg |
| FSCN1_homo_R | ggcacactttttggtgtcg |
| FSCN2_homo_L | cactgggggctactggac |
| FSCN2_homo_R | aacatggtgttggcagaaact |
| FSCN3_homo_L | gatggccaatgtcatccttc |
| FSCN3_homo_R | gccataacgacctcgcaata |
| CMVipuro-miniTurbo_Fwd | ACTCTACTAGAGGATCTATTTCCgccaccATGATCCCGCTGCTGAACGCTAAACAGATTC |
| CMVipuro-miniTurbo_Fwd_extender | tagaagacaccgACTCTACTAGAGGATCTATTTCCgccaccATG |
| CMVipuro-miniTurbo_Rev | agcccccaccgctgccgctccgaagaccggaCTTTTCGGCAGACCGCAGACTGATTTCTC |
| CMVipuro-miniTurbo_Rev_extender | actccctgagccgcctgatgcagAGCCCCCACCGCTGCCGCTC |
| CMVipuro-miniTurbo_CMVipuro_Rev | cgggatccgcggccgctctagaactagtctcCTTTTCGGCAGACCGCAGACTGATTTCTC |
| 18-paxillin_Fwd | gcagcggtgggggctctgcatcaggcggctcagggagtGACGACCTCGACGCCCTGCTGG |
| 18-paxillin_CMVipuro_Rev | ggatccgcggccgctctagaactagtctcCTAGCAGAAGAGCTTGAGGAAGCAGTTCTGA |
